# Discovery of a molecular glue inhibitor that stabilises a non-productive Kalirin-Rac1 complex

**DOI:** 10.64898/2026.09.08.749156

**Authors:** Janine L. Gray, Mia C. Callens, Amelia Henley, Daniel Zaidman, Carmen Jimenez Antunez, Andrew P. Thompson, Zhihe Lei, Rachael Skyner, Michael Miller, Eleanor P. Williams, Anya Robinson, Raina Seupal, Matt T. Fry, Mihajlo Filep, Frank von Delft, Michael C. Willis, Annette R. von Delft, Nir London, Paul E. Brennan

## Abstract

Rho guanosine triphosphatases (GTPases) are molecular-switches implicated in neurodegenerative diseases, yet targeting them through competitive inhibition remains a challenge due to their high affinity for guanine nucleotides. Guanine nucleotide exchange factors (GEFs) catalyse GDP-to-GTP nucleotide exchange to activate GTPases, providing an alternative opportunity for GTPase modulation. Herein, we describe a complex-targeted strategy to inhibit nucleotide exchange with covalent molecular glues that engage the Kalirin-Rac1 GEF-GTPase complex at the nucleotide binding site and sequester the GEF Kalirin. Fragment hits were identified through XChem and in silico screening, and a fragment merging approach resulted in the generation of covalent inhibitors RS-009 and MC-278. Multiple analyses demonstrate our compounds inhibit nucleotide exchange both through competition with the nucleotides and by stabilising a ternary inhibitor-Rac1-Kalirin complex, thereby trapping Kalirin in a non-productive state and reducing GEF turnover. Biochemical selectivity screening and cellular activity-based protein profiling (ABPP) show that selectivity can be achieved across distinct GEF-GTPase complexes, which may result in improved spatiotemporal control over targeting the GTPase alone. This work provides evidence for targeting GTPase signalling via stabilization of the GEF-GTPase complex in a unique covalent molecular glue mechanism and provides the basis of a chemical probe or therapeutic.

## Introduction

The RAS superfamily of guanine triphosphatases (GTPases) are highly pleiotropic molecular switches.^[1]^ The larger family, containing over 150 members, is divided into five evolutionary conserved subfamilies based upon their structure and function: Ras, Rho, Ran, Rab and Arf.^[2]^ Gain-of-function missense mutations in RAS genes are reported to be found in ∼25% of human cancers, and thus have been of focus for new therapeutic approaches.^[1, 3]^ Specifically, the Rho GTPases are involved in the regulation of neuronal cytoskeletal dynamics and actin organization, and so are desirable targets for neurological diseases such as schizophrenia and Alzheimer’s disease.^[4]^

The GTPases cycle between a GDP-bound inactive state and a GTP-bound active state, in which both nucleotides exhibit high affinity (K_d_ = 10^-7^ – 10^-11^ M) for the guanine nucleotide binding pocket of the GTPase.^[5]^ The GTPase activation cycle is regulated by the GTPase activating proteins (GAPs), guanine nucleotide exchange factors (GEFs) and additionally guanine dissociation inhibitors (GDIs) for the Rho and Rab subfamilies (Figure 1). Respectively, these effectors are catalytically necessary for the hydrolysis of GTP to GDP to halt GTPase-mediated signalling, the exchange of the hydrolysed GDP for another GTP molecule to regenerate the active GTPase, and for the binding and sequestration of the inactive GTPase from cellular membranes.

**Figure 1.**
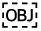
The GTPase activation cycle. GTPases exist in a GDP-bound inactive state and a GTP-bound active state. Nucleotide exchange is aided by guanine nucleotide exchange factors (GEFs), whilst hydrolysis from GTP to GDP is catalysed by GTPase activating proteins (GAPs). Guanine dissociation inhibitors (GDIs) modulate the Rho and Rab sub-families, maintaining the GTPase in an inactive state.

Downstream activation of the subsequent signalling pathways is mediated by the conformation of the switch I and II regions of the enzymes which modify the protein-protein interaction repertoire of the GTP and GDP bound states (Figure 1).^[1]^ Although it allows for tight regulation of GTPase signalling, the high affinity of the bound nucleotides has rendered the GTPases ‘undruggable’ due to the infeasibility of classical competitive inhibition. However, recent breakthrough compounds demonstrate the feasibility of modulating GTPase activity by covalently targeting the GDP-bound state. FDA approved drugs sotorasib and adagrasib are covalent inhibitors of KRas^G12C^, used in in the treatment of KRas^G12C^-positive non-small cell lung cancer (NSCLC).^[6]^ KRas^G12C^ is a mutated oncogene where the mutation inhibits GAP-mediated regulation of nucleotide exchange, resulting in aberrant RAS activity. These covalent GDP-uncompetitive drugs target the off-state and prevent the association of GTP whilst allowing GDP into the nucleotide binding site by reacting with the cysteine mutation and interacting with an adjacent cryptic pocket; this ultimately prevents the overactivation of KRas. Molecular glues are small molecules that bring two proteins together and induce a change in the function of one protein partner, most often degradation,^[7]^ but also inhibition.^[8]^ A promising recent example of a non-degrading molecular glue is the RAS inhibitor daraxonrasib which recruits cyclophilin A to inhibit RAS signaling.^[9]^ A covalent molecular glue EN450 has also been reported that reacts with a cysteine on the E2 UBE2D to degrade NFkB1.^[10]^

An alternative approach exploited here is to modulate GTPase activation by targeting their regulators.^[11]^ MRTX0902, a Ras:SOS1 protein-protein interaction (PPI) inhibitor, reached Phase I/II clinical trials for patients with advanced solid tumour malignancy, demonstrating the potential for the approach clinically.^[12]^ Nucleotide exchange, and thus activation of the GTPases, is controlled by GEFs in biological systems.^[13]^ Upon binding, GEFs induce extensive remodelling of the switch I and switch II regions of the GTPase that disrupts key interactions between the GTPase and the bound GDP (Figure 1). The conformational change reduces the affinity of the GDP to micromolar levels, and nucleotide exchange to GTP occurs at a much faster rate. The preference for reactivation of the GTPase by GTP binding is driven by the higher concentration of GTP in the cell (> 300 μM) compared to GDP (>30 μM).^[11]^

An attractive therapeutic approach could be to develop competitive inhibitors to compete with micromolar binding ligands of GEF/GTPase complex compared to picomolar binding to the GTPase in isolation.^[11]^ By pursuing covalent GEF/GTPase inhibitors, even high levels of GTP/GDP competition could be surmounted by irreversible binding. Such compounds would prevent GTPase activation and thus inhibit the associated signalling pathways. Additionally, for the Rho subfamily, GEFs outnumber the GTPases by three-fold, allowing for better spatiotemporal and cell specific control of downstream processes. This could allow for the design of an inhibitor that is selective for a subspecies of a single GTPase through targeting of a sole GEF/GTPase complex.

A GEF of pharmacological interest is Kalirin, a Rho Dbl GEF associated with synaptic plasticity and dendritic spine formation.^[14]^ Rho Dbl GEFs are characterised by the presence of a Dbl homology (DH) domain – the core catalytic domain responsible for promoting nucleotide exchange – and a tandem Pleckstrin homology (PH) domain – which is variable in its function depending on the GEF.^[13, 15]^ Kalirin is functionally interesting due to the presence of two GEF domains; the first (Kalirin-GEF(1)) is an activator of RAS-related C3 botulinum toxin substrate 1 (Rac1) activity, and the second (Kalirin-GEF(2)) targets RAS homolog family member A (RhoA). Kalirin7, an isoform containing the first GEF domain only, was associated with effects of anxiety in Kalirin7 knockout mice.^[14d]^ Kalirin7 mRNA was also found to be downregulated in the prefrontal cortex of schizophrenic patients.^[14a, 16]^ Contrastingly Kalirin9 mRNA, an isoform which contains both GEF domains, was found to be upregulated in the superior temporal cortex.^[14c, 16]^ These studies highlight the important role Kalirin has in neurodevelopment, but do not confirm that the observed phenotypic effects are due to the differing GEF activity of the Kalirin7/9 isoforms. The generation of a chemical probe for a Kalirin GEF/Rho GTPase complex would be invaluable to establish the role of the GEF activity of Kalirin in neurological signalling networks.^[17]^

Herein, we report the development of novel fragments that covalently inhibit Rac1 nucleotide exchange by targeting the Kalirin-Rac1 complex as a molecular glue. These fragments simultaneously block the nucleotide binding site and stabilise the Kalirin-Rac1 interaction, ultimately inhibiting Kalirin turnover and thus nucleotide exchange. The novel crystal structure of the holo-enzyme complex was solved to high resolution, and a variety of fragment and virtual screening techniques enabled initial hit identification. Covalent fragments were confirmed to engage a proximal cysteine residue within the nucleotide binding site, providing an effective method of preventing nucleotide binding. Subsequent synthesis and evaluation of related analogues allowed expansion of structure-activity relationships (SAR) and the identification of biochemically-active fragments. Notably, these inhibitors also promote stabilisation of the Kalirin-Rac1 complex upon binding, consistent with a glue-like mode of action. Together, this work establishes the basis of the development of chemical probes to study Kalirin-Rac1 biology and represents an opportunity for targeting the RAS superfamily through their regulatory complexes.

## Results and Discussion

### Crystallization of the Kalirin-Rac1 complex

Crystallization experiments for the complex were initiated through the expression of protein constructs derived from previously determined X-ray (Rac1) or NMR structures (Kalirin-GEF(1)) available in the PDB. Gratifyingly, the novel co-crystal structure of the Kalirin DH1 domain (1232-1411) in complex with Rac1 (1-177) with the flexible C-terminus truncated was solved to high 1.64 Å resolution, denoted herein as the Kalirin-Rac1 complex (PDB: 5O33). The Kalirin DH1 domain within the complex consists of a six α-helical bundle characteristic of the Dbl Rho GEF subfamily, whilst Rac1 exhibits a similar overall fold to other small GTPases containing a central β-sheet interspersed by α-helices (Figure 2A).^[15c]^ The two proteins largely interact via the flexible switch I and II regions of Rac1 (residues 26–45 and 59–74 respectively). The crystal structure of Kalirin-Rac1 shares a large similarity with the free solution NMR structure of Kalirin alone previously deposited in the PDB, suggesting that the conformation of Kalirin observed is energetically favourable (PDB: 2KR9).^[18]^ Notably GDP was present in high occupancy, despite not being added during the purification process, and thus must originate from the bacterial expression system (Figure 2B). The guanine base made key hydrogen bonds with D118 of Rac1, with phosphate groups interacting with the backbone of K16 and T17.

**Figure 2.**
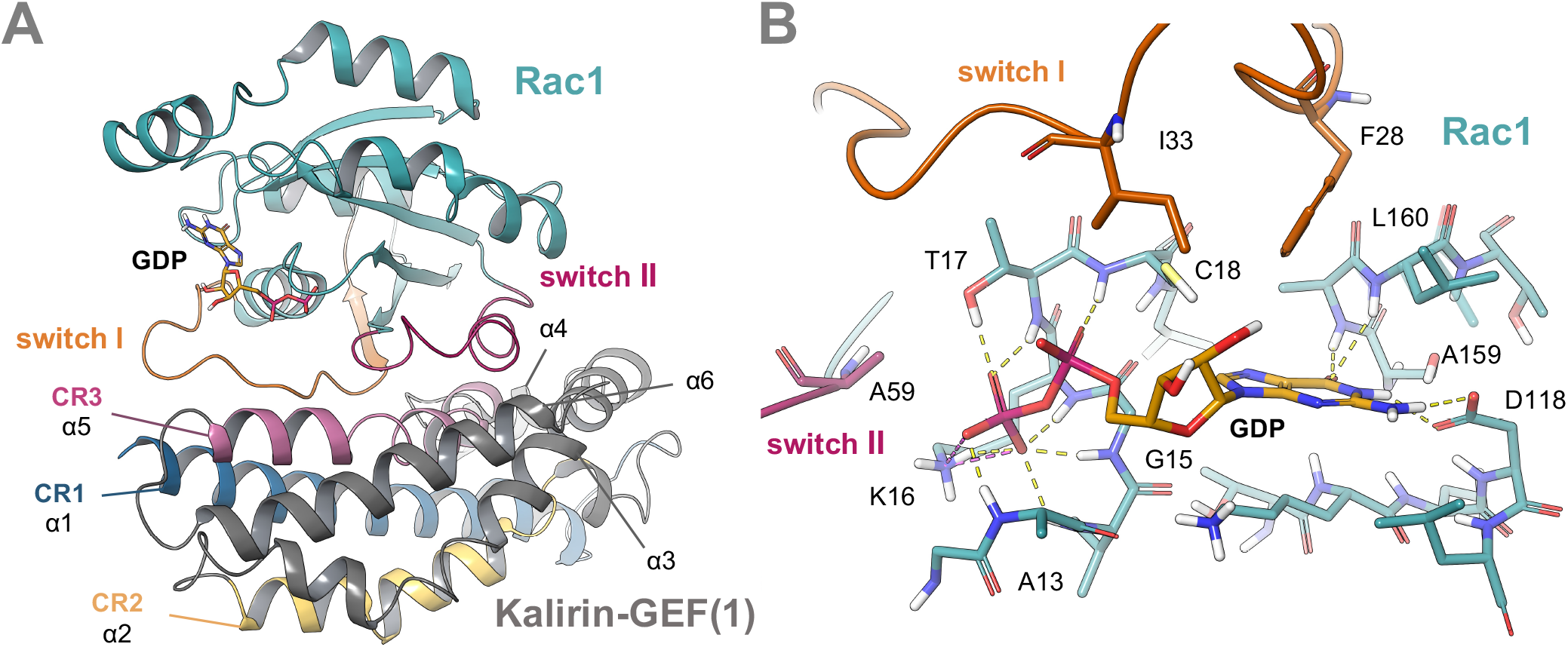
The novel GDP-bound Kalirin-GEF(1)-Rac1 crystal structure. A. The GDP-bound co-crystal structure of Rac1 and the DH domain of Kalirin-GEF(1) (PDB: 5O33). Rac1 (teal ribbons) contains the switch I (orange ribbons) and switch II (magenta ribbons) regions that form the nucleotide binding site for GDP (yellow sticks) and interact with the GEF domain. The DH domain of Kalirin-GEF(1) has a 6 a-helical structure containing conserved regions (CRs) CR1 (blue ribbons), CR2 (yellow ribbons) and CR3 (pink ribbons). CR1 and CR3 interact with Rac1 at the switch I and II regions. B The nucleotide binding site of Kalirin-GEF(1)/Rac1. GDP (yellow sticks) binds to Rac1 (teal sticks) through various interactions and is formed at the juncture of the switch I region (orange sticks) and switch II region (magenta sticks). Hydrogen bonding interactions (yellow dashes) and salt bridges (pink dashes) shown.

Mg^2+^ is an essential cofactor required for GTP binding and hydrolysis. In structures of Rac1 bound to GTP, the γ-phosphate of GTP as well as T17 and T35 coordinate to Mg^2+^ to form a high-affinity ‘active’ arrangement of the GTPase (PDB: 3TH5, Figure S1).^[19]^ However, electron density for Mg^2+^ was not observed in the Kalirin-Rac1 structure; we hypothesized that the conformation of Rac1 induced by Kalirin is unfavourable for Mg^2+^ binding, with key residue T35 of the switch I loop twisted out of position.^[20]^ The displacement of Mg^2+^ has been observed in the crystal structures of Rac1 bound to other Dbl family members (PDB: 2NZ8) and is thought to be responsible for reducing affinity to GDP and allowing nucleotide exchange. A59 of switch II also overlaps with the Mg^2+^ position observed in structures of Rac1 alone (PDB ID: 5N6O and 3TH5),^[21]^ suggesting the reorientation of this residue by Kalirin also reduces GDP affinity, preventing essential interactions between the phosphate groups and magnesium ion. We observed additional changes in the structure of Rac1 in the Kalirin-bound complex as opposed to Rac1 alone; key residues of the switch I (Y32 and I33) and II (A59 and Y64) loops move significantly as compared to the GTP-bound active form. We envisaged that this interface could be targeted by a small molecule for improved selectivity for the GEF-bound complex over the GTPase alone, in anticipation that targeting the low-affinity binding state of the guanine nucleotides would yield more potent competitive inhibitors. In addition, a cysteine residue (C18) was identified proximal to the GDP binding site, presenting an opportunity for covalent targeting.

The longer residence time of a covalent inhibitor and subsequent slower turnover of the Kalirin-Rac1 complex could result in enhanced potency and prevention of GTPase activation even in high cellular concentrations of guanine nucleotides.

### Initial hit identification

Crystallization conditions were optimized to yield Kalirin-Rac1 crystals that reproducibly diffracted with a resolution of 1.5-2.0 Å. Fragment screening was then initiated via X-ray crystallography at Diamond Light Source. Over 450 Kalirin-Rac1 crystals were soaked with a single fragment from the DSI-Poised^[22]^ or OxXChem^[23]^ fragment libraries and screened using the high-throughput X-ray crystallography platform at the XChem facility.^[24]^ Selected datasets from crystals that diffracted to >3 Å were submitted to the XChemExplorer pipeline^[25]^ for molecular replacement and refinement, before PanDDA^[26]^ was used to identify hits. Utilizing this workflow, isoxazole-urea fragment **1** was found to bind within the guanine nucleotide binding site with high occupancy, showing it was capable of displacing GDP from the pocket (PDB: 5QQD, Figure 3A). The urea moiety formed two key hydrogen bonds with D118, and the isoxazole interacted with the peptide backbone of A159, mimicking the hydrogen bond donor-donor-acceptor pattern observed in the GDP-bound structure. Changes in residue alignment were also observed in the GDP binding site and switch I loop (such as E31, Y32, and I33) compared to both the GDP-bound Kalirin-Rac1 and GDP/GTP-bound Rac1 structures. We anticipated that in the development of **1**, these position shifts could be exploited for selectivity over Rac1 alone. In pursuit of this, 1000 commercially available analogues were clustered and subsequently docked using Molsoft ICM-Pro. A total of 105 analogues, which were predicted to maintain the same binding pharmacophore as **1** but extended further into the pocket and the switch I region, were purchased and screened using XChem.

**Figure 3.**
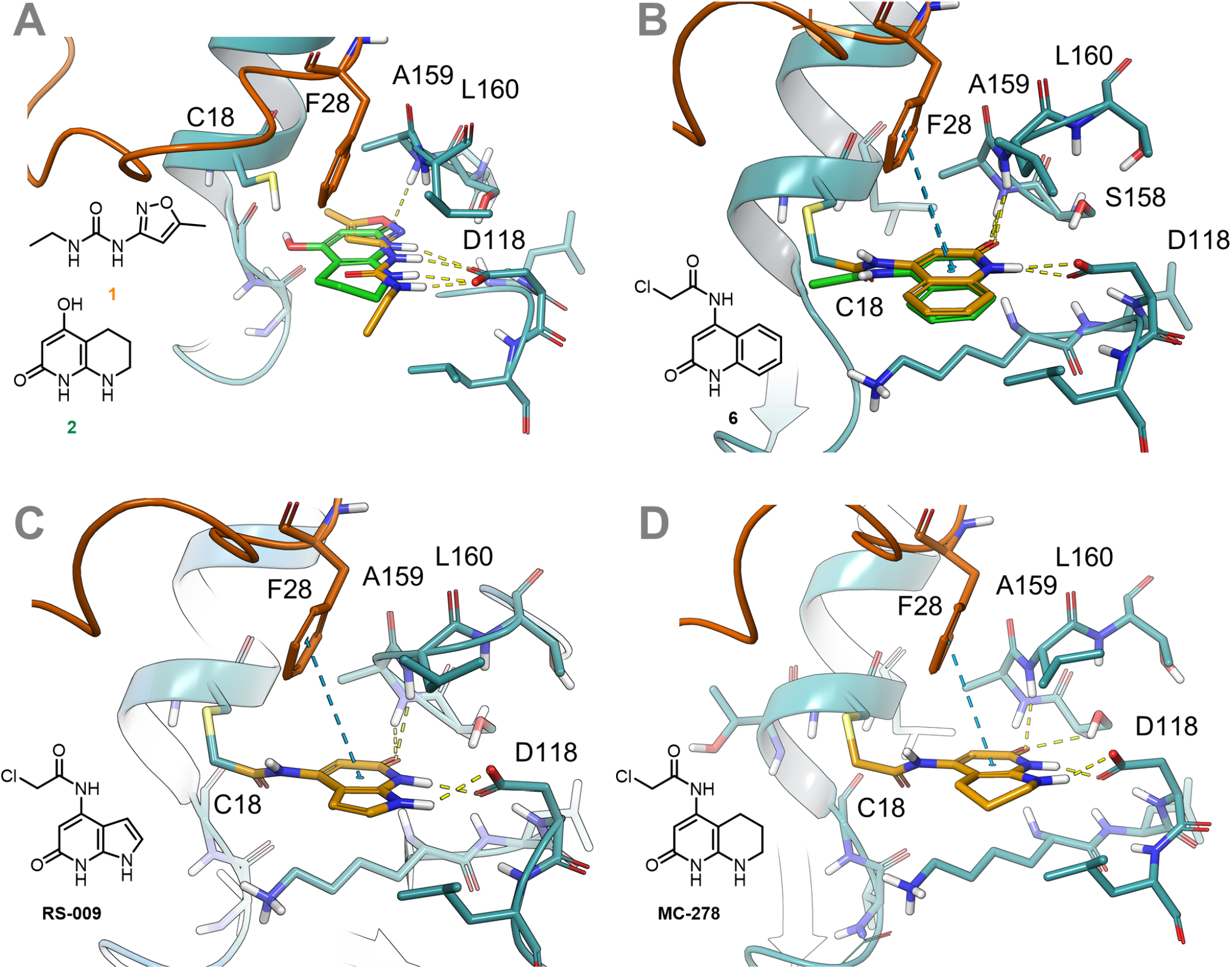
X-ray crystal structures of fragments and follow-on compounds bound to Kalirin-Rac1 at the nucleotide binding site. Rac1 (teal ribbons and sticks) and the switch I region (orange ribbons and sticks) are shown in each panel. A. Binding of isoxazole 1 (gold sticks) and piperidine 2 (green sticks) overlaid, showing dual hydrogen bonding interactions with D118. B. Binding of chloroacetamide 6 (gold sticks), overlaid with the predicted pose by DOCKovlanet (green sticks), covalently bound to C18. C. Binding of RS-009 (gold sticks) and D. MC-278 (gold sticks), designed through fragment merging.

From this, 10 compounds were identified using PanDDA; 9 of these compounds contained modifications to the ethyl chain (Figure S2). Analysis of the structures showed few additional interactions with the pocket, with the elaborated chains instead extending out into the solvent channel of the crystal. These compounds were not pursued further due to predicted poor ligand efficiency compared to **1**. As analogues of the urea moiety were not commercially available, bioisosteres were designed using SwissBioisostere^[27]^ and synthesised in house. It was anticipated that compounds which mimic the interactions of **1** but had accessible vectors for extension into the guanine nucleotide pocket would be identified. Despite the synthesis of 16 bioisosteres, no binding was identified when the compounds were soaked into Kalirin-Rac1 crystals (Figure S3).

The lack of improvement achieved with analogues of **1** was thought to be due to their small size and relatively few new interactions compared to the endogenous substrates. As such, an in silico search was conducted on the Enamine Real database for compounds with similar pharmacophores but diverse chemistry to GDP, fragmented GDP or non-covalent hit urea **1**. The commercially available options were docked using AutoDock^[28]^ and ranked using SuCos^[29]^ and pLIF.^[30]^ The top 300 compounds were purchased and screened using the XChem facility. Of the hits identified, pyridone **2** was of particular interest due to its interactions with D118 and similar binding pose to **1** (Figure 3A).

Concurrently, we envisaged that covalent analogues of **1** targeting C18 may show improved occupancy and binding in the Kalirin-Rac1 complex. Covalent docking using Molsoft ICM predicted that an electrophilic warhead installed on position 3 of the isoxazole ring of isoxazole **1** would bear the correct orientation to react with C18, whilst retaining the urea in position to interact with D118. Accordingly, we designed and docked three covalent analogues **3-5** with varying warheads, exampled here by chloroacetamide **3** (Table 1 and S4). Although the designed molecules were successfully synthesized, attempts to co-crystallize the compounds with Kalirin-Rac1 were unsuccessful, and no significant labelling was observed by intact mass spectrometry upon incubation with recombinant Kalirin and Rac1.

**Table 1.** IC_50_ values of compounds tested in the BODIPY-GTP nucleotide exchange assay. Dose responses were performed as 10-point 2-fold serial dilution with a high concentration of 500 μM. ^a^ Ambiguous due to precipitation. *ND* = not determined

| Compound | Structure | $IC_{50}$ (2h) (μM) | Compound | Structure | $IC_{50}$ (2h) (μM) |
| --- | --- | --- | --- | --- | --- |
| 1 |  | > 500 | 7 |  | ND |
| 2 | | <sup>a</sup> | RS-009 | | 11.9 ± 0.2 |
| 3 | | ND | MC-278 | | 10.8 ± 1.2 |
| 6 |  | > 500 | 8 |  | > 500 |

In an effort to more broadly explore covalent inhibition, 67,127 chloroacetamides were screened in silico using DOCKovalent^[31]^ against the GDP pocket of Rac1 in the Kalirin-Rac1 complex. Compounds were preferentially ranked based on their proposed capability to mimic the hydrogen bonding pharmacophore of GDP and isoxazole **1**. From this, 8 compounds were chosen and a dose-response assay was conducted, with labelling of Rac1 confirmed by intact mass spectrometry. Of these, chloroacetamides **6** and **7** were selected as hits due to their preferential labelling of Rac1 in the Kalirin-Rac1 complex over Rac1 alone. They also exhibited lower reactivity in a selectivity panel compared to other scaffolds with the chloroacetamide warhead, thus lowering the risk of poor selectivity associated with reactive covalent compounds in cells (Figure S5).^[32]^

To confirm the site of labelling, **6** and **7** were incubated with the Kalirin-Rac1 complex at a 4-fold higher concentration overnight at 4 °C to ensure > 80% labelling as confirmed by QTOF intact mass spectrometry. Crystal structures of the chloroacetamides bound to Kalirin-Rac1 were also obtained to 1.35 Å (PDB: 9IFK, Figure 3B) and 1.65 Å (PDB: 9IG1. Figure S6) respectively. Gratifyingly, electron density indicated that both compounds were covalently bound to C18 within the guanine nucleotide binding pocket and formed a hydrogen bond with D118, and the heterocyclic rings of both compounds were orientated in an analogous manner to the guanine ring of GDP. Moreover, the binding mode showed strong agreement with the corresponding poses originally predicted by DOCKovalent (Figure 3B and Figure S6). Chloroacetamide **6** was chosen for further elaboration as the orientation of the fused core with respect to D118 allowed for the introduction of another hydrogen bonding interaction, as well as offering a vector for extending further into the pocket in a synthetically accessible manner. In an attempt to extend the compound through fragment linking, an XChem fragment screen was conducted to identify fragments that bind near to **6** within the nucleotide binding pocket. In this aim, 176 co-crystals of **6** bound to Kalirin-Rac1 were soaked with individual fragments using the DSIPoised library. However, no fragments were identified within the nucleotide binding pocket.

By instead adopting a structure-based design approach, we anticipated that the second hydrogen bonding interaction observed in the co-crystal structures of Kalirin-Rac1 bound to GDP, isoxazole **1**, and piperidine **2** could be achieved by incorporating a hydrogen bond donor in the phenyl ring of chloroacetamide **6**. This strategy led to the design and synthesis of RS-009 and MC-278, using pyrrole and piperidine moieties respectively (Figure 3C, 3D). Co-crystallization experiments were performed and the structure of the RS-009-Kalirin-Rac1 complex solved to 1.68 Å, and MC-278-Kalirin-Rac1 to 1.48 Å. Both fragments were covalently bound to C18, with the core scaffolds shifting in comparison to chloroacetamide **6** to generate the second hydrogen bond interaction with D118. Comparison of RS-009 and MC-278 to GDP show that their heterocyclic rings overlap almost exactly, with the fragments exhibiting the desired modality proposed using structure-based design and covalent docking.

### Evaluation of compounds in a nucleotide exchange assay

To quantitatively assess the ability of the fragments to inhibit nucleotide binding, a nucleotide exchange assay was optimized for the Kalirin-Rac1 complex. Guided by established assays for RAS binders,^[33]^ we developed a biochemical fluorescence-based association assay to monitor the exchange of GDP for the fluorophore-containing nucleotide analogue BODIPY-GTP. Binding of BODIPY-GTP exhibits an increased fluorescence intensity, and nucleotide exchange inhibitors reduce the initial increase of fluorescence intensity upon addition of BODIPY-GTP.

Whilst chloroacetamide **6** was not active in the nucleotide exchange assay, RS-009 and MC-278 showed significant inhibition compared to the negative control after incubation (Table 1, Figure S7). Isoxazole **1** was also tested but did not show inhibition at 500 µM. This highlights the importance of the dual hydrogen bond donor pharmacophore, as well as the influence of the covalent bond with C18 to prolong the residence time of the compound. This interaction was further probed by synthesising and testing pyrazolopyridone **8**, which was predicted to be more synthetically accessible than RS-009 and MC-278. However, introducing an additional nitrogen atom at C2 was deleterious to activity, highlighting the importance of carefully tuning the hydrogen bond donor properties. The activity of pyridone **2** was also investigated, but a reproducible dose response curve could not be obtained unambiguously; this was suspected to be due to precipitation or aggregation. The importance of the covalent warhead at this stage of discovery was also emphasized by the lack of activity of non-covalent counterparts of MC-278, including the closely related acetamide (Table S1).

### Kinetic analysis of covalent inhibitors

The covalent nature of RS-009 and MC-278 was investigated further. Covalent fragments possess time-dependent inhibition, and so their true activity is best assessed as a kinetic parameter. Exploiting the method described by Krippendorf,^[34]^ both RS-009 (90 M^-1^s^-1^) and MC-278 (50 M^-1^s^-1^) were observed to inhibit Kalirin-Rac1 nucleotide exchange in a time-dependent manner, thus further demonstrating the covalent mechanism of action (Figure 4A, 4B). The covalent nature of RS-009 and MC-278 was investigated further. Covalent fragments possess time-dependent inhibition, and so their true activity is best assessed as a kinetic parameter. Exploiting the method described by Krippendorf,^[34]^ both RS-009 (90 M^-1^s^-1^) and MC-278 (50 M^-1^s^-1^) were observed to inhibit Kalirin-Rac1 nucleotide exchange in a time-dependent manner, thus further demonstrating the covalent mechanism of action (Figure 5A, 5B).

**Figure 4.**
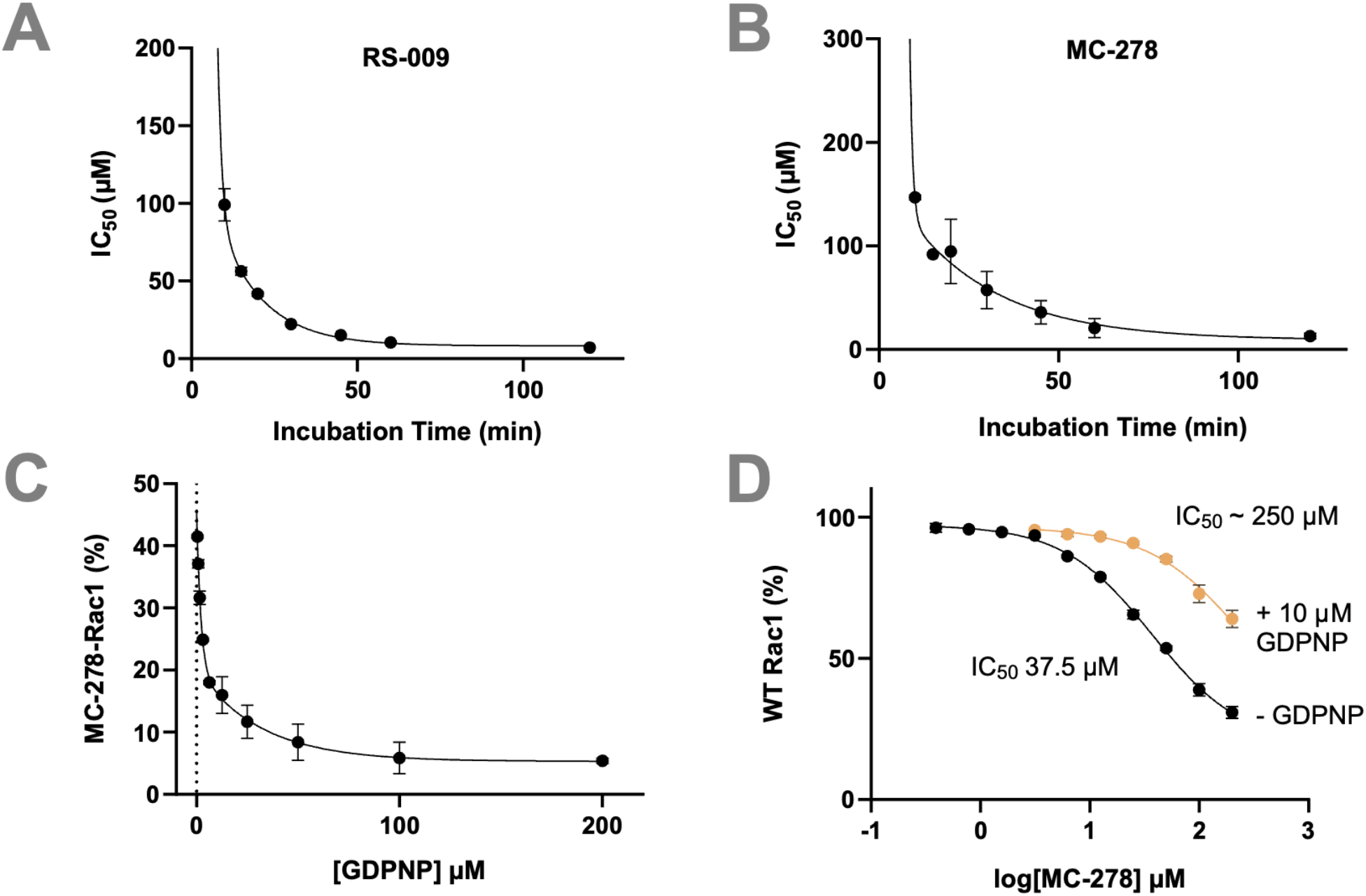
A. Time-dependent inhibition of Kalirin-Rac1 by RS-009. B. Time dependent inhibition of Kalirin-Rac1 by MC-278. C. Effect of [GDPNP] on Rac1 labelling by MC-278. MC-278-Rac1 (%) denotes the % conversion between MC-278 labelled Rac1 and WT Rac1. D. Effect of [GDPNP] on MC-278 IC_50_.

**Figure 5.**
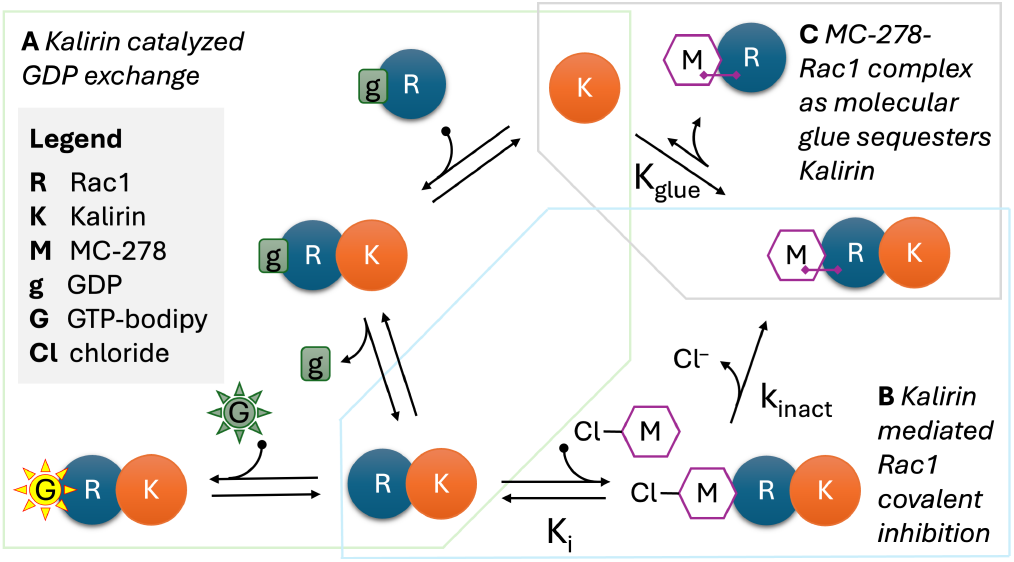
MC-278 stabilises Kalirin-Rac1 and reduces Kalirin turnover. A) Kalirin (K) binds to Rac1-GDP (gR) and g dissociates to give Rac1-Kalirin (RK) (light green box). BODIPY-GTP (green G) binds and fluoresces (yellow G). B) MC-278 (M) reversibly binds to the RK complex (Ki) and reacts with Cys18 (k_inact_) to give covalent MRK (blue box). C. MRK can theoretically dissociate to MR and free K, but MC-278 stabilises RK and K is prevented from re-entering the catalytic cycle (K_glue_ is large) (grey box).

### Competitive inhibition with endogenous ligands

To further corroborate the covalent and competitive nature of RS-009 and MC-278, a protein mass spectrometry (MS) assay was optimised. In studies herein, MC-278 was used as a representative compound on account of its superior synthetic accessibility. Here, the abundance of wild type (WT) Rac1 and MC-278 modified Rac1 (MC-278-Rac1) were measured under different conditions. Notably, when incubating in the absence of Kalirin, no MC-278-Rac1 was observed, demonstrating the selectivity of the compound for the complex (Table S2). Similarly, Kalirin was not observed to be MC-278 labelled in any experiment.

When Kalirin-Rac1 was pre-incubated with MC-278, the addition of GDPNP was unable to disrupt the covalent binding of MC-278 as expected (Figure S8). Finally, when incubating with both ligands simultaneously, GDPNP disrupted MC-278 binding before covalent modification, and thus increased the IC_50_ of MC-278 nearly 7-fold in the presence of 10 uM GDPNP (Figure 4C and 5D). This provided further evidence of covalent inhibition at the nucleotide binding site.

### MC-278 stabilises the Kalirin-Rac1 interaction

Complete inhibition of BODIPY-GTP binding in the biochemical assay was initially attributed to near-full occupancy of the Rac1 binding site by MC-278, facilitated by catalytic Kalirin. However, optimisation of the MS assay revealed that stoichiometric Kalirin:Rac1 was required to detect MC-278-labelled Rac1, indicating a distinct mechanism which is not catalytic with respect to Kalirin (Table S2).

This led us to hypothesize MC-278 functions as a covalent molecular glue, stabilising an MC-278-Rac1-Kalirin ternary complex (Figure 5). This stabilisation is proposed to sequester Kalirin in a non-productive state, thereby limiting catalytic turnover (Figure, 6C). Only a small fraction of Rac1 is modified before Kalirin is effectively abrogated, resulting in undetectable MC-278-Rac1 labelling by MS, but complete inhibition of nucleotide exchange biochemically. Molecules that influence the dissociation of native macromolecular interactions have been discussed previously, and could be an effective approach to modulate cellular processes.

We sought to investigate the molecular glue mechanism by exploring the proposed steps hypothesized in the model (Figure 5B and 5C).

The degree of Rac1 labelling by MS increased with both increased Kalirin concentration and time (Figure 6A). We then examined the Rac1 nucleotide exchange rate while increasing Kalirin concentration at a fixed concentration of MC-278 and Rac1. At low Kalirin concentrations, the initial rate of nucleotide exchange increased as expected for a catalyst. However, an inflection point was observed at 0.5 µM Kalirin, beyond which increasing the concentration paradoxically decreased the initial rate (Figure 6B). This behaviour is consistent with the MS data and supports the complex trapping mechanism (Figure 5C). At higher Kalirin concentrations, MC-278 appears to promote the formation of a non-productive MC-278-Rac1-Kalirin complex that sequesters Kalirin and prevents further GTPase exchange catalysis. As Kalirin concentration increases, a greater fraction of MC-278-Rac1 is formed, trapping more Kalirin in this complex, reducing the pool of active Kalirin and consequently decreasing the observed nucleotide exchange rate.

**Figure 6.**
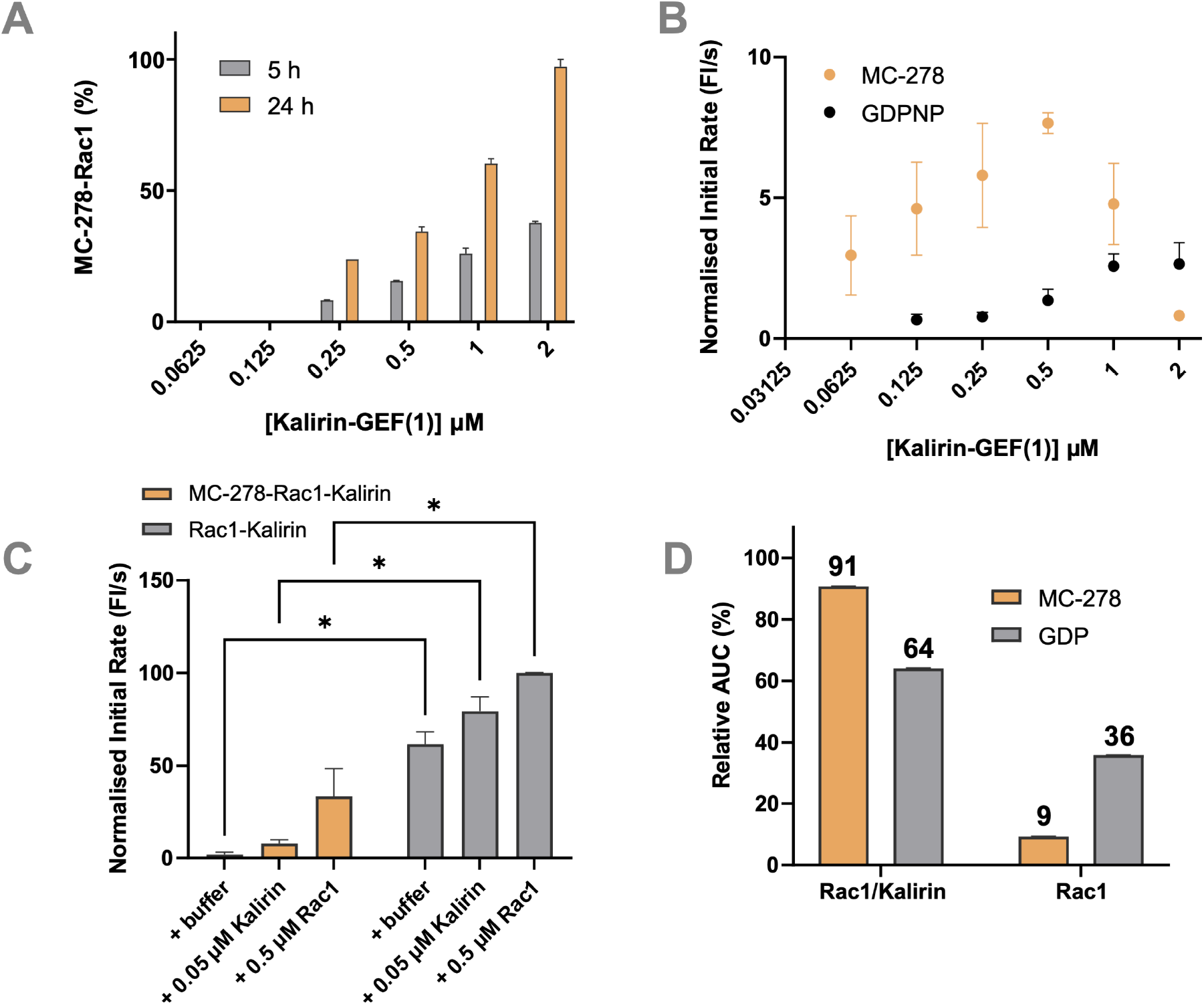
A. Time-dependent Rac1 (1 µM) labelling by MC-278 (100 µM) in the presence of varying concentrations of Kalirin after 5 h and 24 h incubation. MC-278–Rac1 (%) denotes the percentage conversion of WT Rac1 to MC-278-labelled Rac1. B. Kalirin-dependent inhibition of Rac1 (1 µM) by MC-278/GDPNP (100 µM) following 24 h incubation. C. Normalised initial rates measured after addition of 0.5 µM of either MC-278-Rac1_B_-Kalirin or Rac1_B_-Kalirin control complexes to buffer, Rac1_B_ (0.5 µM) or Kalirin (0.05 µM). Data from three independent experiments presented as mean ± SEM with significant increases (*p< 0.05) in normalised initial rates for Rac1_B_-Kalirin control compared to MC-278-Rac1_B_-Kalirin. D. Quantification of size-exclusion chromatography (SEC) peak areas from the elution 1 (E1) fraction obtained following Strep-Tactin purification of MC-278 incubated with biotinylated Rac1 (MC-278-Rac1_B_-Kalirin) or DMSO control incubated with biotinylated Rac1 (Rac1_B_-Kalirin). Peak integration shows that the MC-278 sample predominantly consists of the MC-278-Rac1_B_-Kalirin complex (91%), with only a minor fraction of free Rac1_B_ (9%). In contrast, the DMSO control contains a substantially lower proportion of Rac1_B_ -Kalirin complex (65%) and a higher fraction of free Rac1_B_ (35%).

To validate this interpretation and assess the stability of the complex, we attempted to isolate the MC-278-Rac1-Kalirin complex. However, purifying the MC-278-Rac1-Kalirin complex with sufficient purity by size exclusion chromatography (SEC) proved challenging due to co-elution with unbound Kalirin. To circumvent this limitation, a biotinylated Rac1 construct (Rac1_B_) was obtained and incubated with Kalirin in the presence of MC-278 or with no compound as a control.

The resulting complexes were isolated from free Kalirin using Strep-Tactin magnetic beads. Following two buffer washes to remove unbiotinylated Rac1 and unbound Kalirin, the MC-278-Rac1_B_-Kalirin and Rac1_B_-Kalirin complexes were eluted successfully (Figure S9).

The eluted MC-278-Rac1_B_-Kairin and Rac1_B_-Kalirin complexes were subsequently taken on to the fluorescent assay (Figure 6C). Firstly, the Rac1_B_-Kalirin complex demonstrated significantly increased initial rates compared to the MC-289-Rac1_B_-Kalirin complex. This increased rate suggested that MC-278 successfully inhibited Rac1, which is confirmed by the addition of Kalirin showing limited boost in activity. Critically, upon addition of 0.5 µM of Rac1, Rac1_B_-Kairin had significantly higher activity than MC278-Rac1_B_-Kalirin. We believed this to be due to MC-278 sequestering Kalirin in the Rac1_B_-Kalirin complex and thus not available to catalyse additional Rac1. On the contrary, adding more Rac1 to Rac1_B_-Kalirin leads to significantly faster exchange because Kalirin can more readily dissociate. Although this interpretation is not entirely robust as we were not able to keep total concentrations of the proteins consistent in this complex assay format and changes in rate could also be due to small differences in total protein concentrations. More definitive evidence came from SEC analysis: incubating with Kalirin with either GDP-Rac1_B_ or MC-278-Rac1_B_ shifted the equilibrium from the protein monomers toward the Rac1_B_-Kalirin complex, increasing the fraction of Kalirin-bound Rac1_B_ (GDP-complex) from 64% to 91% (Figures 7D and S10) confirming that MC-278 forms a molecular glue between Rac1 and Kalirin.

**Figure 7.**
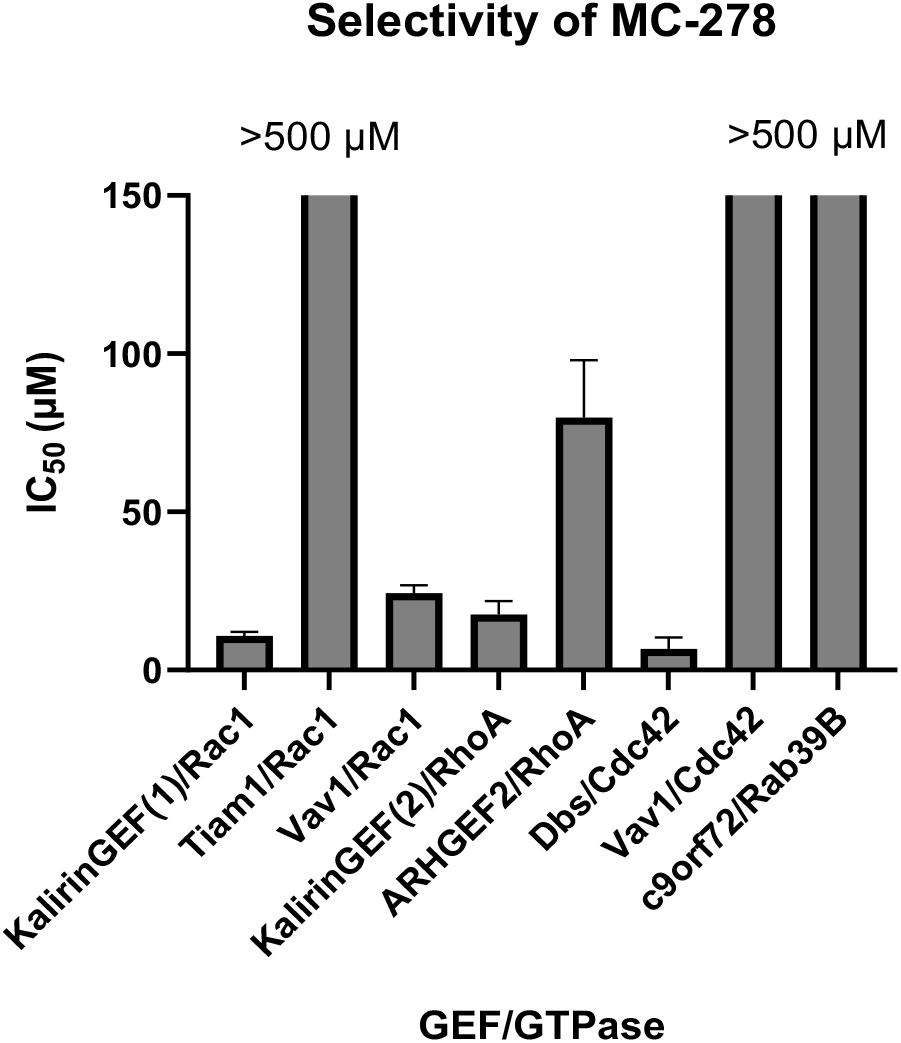
Selectivity of MC-278 for different GTPase/GEF pairs

### Cysteine and selectivity analysis

To explore the selectivity of MC-278 across the RAS superfamily, sequences from 153 members were aligned using ClusterX in JalView (Figure S11). The target cysteine position is predominantly occupied by serine, followed by cysteine in 21% of family members. It was expected that other residue positions in the canonical pocket may vary, allowing for selectivity of our compounds despite the cysteine presence. To investigate this, a selectivity panel comprising multiple Rho GEF-GTPase pairs, along with one Rab pair, was evaluated (Figure 7). All selected GTPases bore the corresponding C18 residue, and thus had the potential to covalently interact with MC-278.

Intriguingly, MC-278 did not uniformly inhibit nucleotide exchange across all GEF-GTPase pairs despite its fragment-like nature which renders it able to physically fit in all GTPase catalytic sites. This variation did not correlate with a specific GEF or GTPase alone, but with the specific pairs; inhibition is both GTPase dependent (Vav1/**Rac1** IC_50_ < 50 µM vs Vav1/**Cdc42** IC_50_ > 500 µM) and GEF dependent (**Kalirin(1)/**Rac1 and **Vav1**/Rac1 IC_50_ < 50 µM vs **Tiam**/Rac1 IC_50_ > 500 µM; **Kalirin(2)**/RhoA IC_50_ < 25 µM vs **ARHGEF2**/RhoA IC_50_ > 50 µM; **Dbs**/Cdc42 IC_50_ < 50 µM vs **Vav1**/Cdc42 IC_50_ > 500 µM).

Based on our earlier findings, these differences may arise from subtle changes in the binding pocket that influence MC-278 engagement, as well as differences in the stability of the resulting ternary complexes; MC-278 may diversely influence the dissociation rate of the GEF-GTPase complex in each case, and thus nucleotide exchange inhibition. The data demonstrates that selectivity between GEF-GTPase complexes in the same subfamily could be achieved, thus offering improved spatiotemporal control over inhibiting the GTPases alone.

A common issue with covalent fragments is that the warhead reacts non-selectively with other cysteine residues in cells. To profile the reactivity of these compounds, we performed an isoDTB experiment to quantitatively determine the labelling of cysteines within the human proteome.^[35]^ Whilst we were unable to identify Rac1-C18 in this experiment due to the theoretical tryptic peptide being undetectable by MS, we were able to identify Cdc42-C18 and RhoA-C20 (homologous with C18) in the top 2% and 4% of peptides enriched in the MC-278 and RS-009 samples compared to DMSO respectively. Interestingly, very few “C18” peptides were detected even with iodoacetamide labelling presumably due to the inability of even highly reactive electrophiles to compete against cellular GDP/GTP levels. Overall, the peptide labelling via MC-278 and RS-009 was minimal showing that despite their potential to non-specifically react with many Cys residues, the chloroacetamide was not generally reactive across the cysteinome. To further assess intrinsic thiol reactivity, a DTNB-based assay was performed (Figure S12).^[32]^ The initial second-order rate constant of MC-278 was determined from the consumption of TNB, monitored at 415 nm over 73 min, and was found to lie within the range reported for chloroacetamide-containing fragments, with iodoacetamide serving as a reference electrophile (Table S3). Consistent with this moderate reactivity, covalent adduct formation between MC-278 and TNB was only detected by HRMS after 24 h of incubation. Together, these results indicate that MC-278 possesses a relatively mild electrophilic character and does not exhibit broadly indiscriminate thiol reactivity.

Whilst alternative Cys profiling methods are required to show selective Rac1-C18 engagement in cells, these results are promising and correlate with the *in vitro* results that suggest that these compounds can bind to GEF-GTPase complexes that have reduced affinity for endogenous nucleotides.

## Conclusion

In summary, competitive inhibition of GTPase activity is still considered a challenge in modern drug discovery due to the tight binding of the nucleotides that dictate GTPase activity. This is despite considerable efforts to modulate the family due to their implication in diseases such as bipolar disorder and Alzheimer’s disease. In this complex-targeted approach, we have developed covalent inhibitors of the Kalirin-Rac1 GEF-GTPase complex that prevent nucleotide exchange through competitive inhibition and by trapping Kalirin in a stable ternary complex, preventing propagation of the GTPase cycle. The fragments, designed rationally based on hits from XChem and in silico screening, were characterized in fluorescence and mass spectrometry-based assays and the binding mode was confirmed by X-ray crystallography. The compounds preferentially bind to the complex over Rac1 alone, as predicted by our initial hypothesis that the reduced affinity of the GEF-GTPase complex for GDP-GTP would enable the development of guanine nucleotide competitive inhibitors. Comparisons of biochemical, MS and SEC experiments also support that the inhibitors stabilise the ternary complex, inhibiting the catalytic activity of Kalirin and thereby abrogating nucleotide exchange. The mechanism of how these covalent fragments function to glue Kalirin to Rac1 is not obvious as the fragments do not make any direct interactions with Kalirin base don the 3D structures. We hypothesize that RS-009 and MC-278 alter the structure of Rac1’s switch I region to strengthen the native interaction with Kalirin (Figures 2 and 3).

Selectivity studies show that our early-stage covalent fragments can achieve selective inhibition across GEF-GTPase complexes. This highlights the potential to develop inhibitors or probes that target specific GTPase cellular functions by optimising compounds for particular GEF-GTPase pairings, which may be more therapeutically relevant in certain diseases.

The mode of action of the covalent inhibitors offers the intriguing possibility of discovering selective GEF inhibitors via the pathway shown in Figure 5. As the catalyst of the GTP exchange, GEFs are generally present at lower levels than their GTPase/GDP substrates. Combining that with the behavior we observed – the GTPase/inhibitor complex is formed only in stoichiometric levels of GEF – in an endogenous cellular setting, excess GTPase would not be inhibited leaving it free for its pleiotropic functions and there would only be enough GTPase/inhibitor formed to inhibit the catalytic GEF through molecular glue sequestration. This mode of action of the covalent inhibitor would spare GTPases spatiotemporaly distal to the targeted GEF and has the potential to minimize sweeping on-target effects for an alternative GTPase inhibitor which is GEF independent. Or putting simply: the GEF only uses as much GTPase as needed to create its own inhibitor.

Structural and computational analysis of different GEF-GTPase complexes may provide an insight into the determinants of selectivity, enabling the rational design of truly selective compounds. The low micromolar activity fragments provide a promising starting point for the development of more potent and selective probes or drugs, exploiting a GEF-dependent mechanism for better spatiotemporal control of GTPase activity.

## Acknowledgements

This work was supported by Alzheimer’s Research UK. A.H was supported by King Abdulaziz University and University of Oxford Centre for Artificial Intelligence in Precision Medicine (KO-CAIPM). M.T.F. was supported by the EPSRC Centre for Doctoral Training in Synthesis for Biology and Medicine (EP/L015838/1). Research in the London lab is funded by the Abisch-Frenkel foundation, the Israel Science Foundation (1869/24), the Israel Cancer Research Fund (PG-25-1465171) the Honey and Dr. Barry Sherman Lab, the Abisch-Frenkel RNA Therapeutics Center, the Moross Integrated Cancer Center, the Goldhirsh-Yellin Foundation and Celia Zwillenberg-Fridman.

