## Supporting Information for "Discovery of a molecular glue inhibitor that stabilises a non-productive Kalirin-Rac1 complex"

Janine L. Gray†,[a,c,d] Mia C. Callens†,[a] Amelia Henley, [a] Daniel Zaidman,[b] Carmen Jimenez Antunez,[a] Andrew P. Thompson,[a,e], Zhihe Lei, [a,g,f] Rachael Skyner,[c] Michael Miller,[a] Eleanor P. Williams,[a] Anya Robinson,[a] Raina Seupal,[a] Matt T. Fry, [a] Mihajlo Filep,[b] Frank von Delft,[a,c] Michael C. Willis,[g] Annette R. von Delft,[a] Nir London,*[b] and Paul E. Brennan.*[a]

[a] The Centre for Medicines Discovery (CMD), Nuffield Department of Medicine, University of Oxford, Old Road Campus, Roosevelt Drive, Oxford, OX3 7FZ

[b] Dept. of Chemical and Structural Biology, Weizmann Institute of Science, 234 Herzl Street, Rehovot, Israel

[c] Diamond Light Source, Diamond House, Harwell Science and Innovation Campus, Fermi Ave, Didcot OX11 0DE

[d] Present address: Molecular Sciences Research Hub, Department of Chemistry, Imperial College London, Wood Lane, London W12 0BZ,UK

[e] Present address: Walter and Eliza Hall Institute of Medical Research (WEHI), 1G Royal Parade, Parkville, Victoria 3052, Australia

[f] Present address: Yusuf Hamied Department of Chemistry, University of Cambridge, Lensfield Road, Cambridge, CB2 1EW

[g] Department of Chemistry, University of Oxford, Chemistry Research Laboratory, Oxford, OX1 3TA, UK.

#### SI Figures and Tables

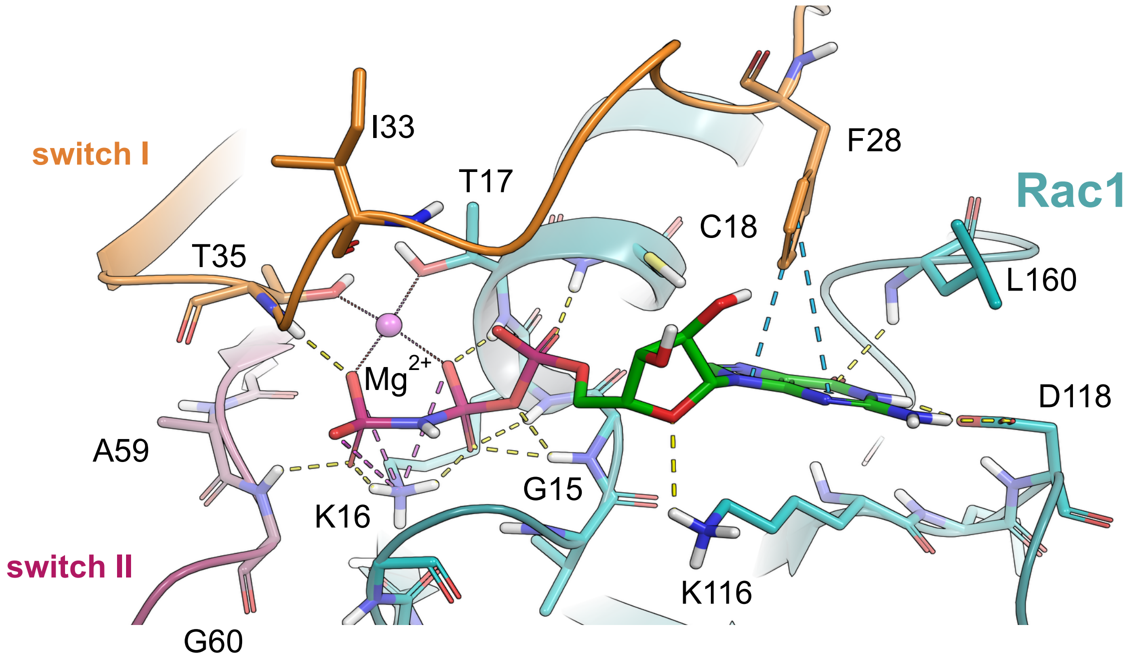

**Figure S1.** X-ray crystal structure of GTP bound to Rac1 (Ha, B.H., Boggon, T.J., PDB: 3TH5).

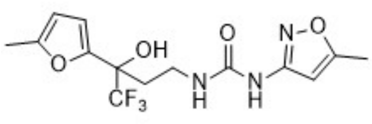

**S1**

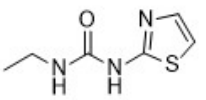

**S2**

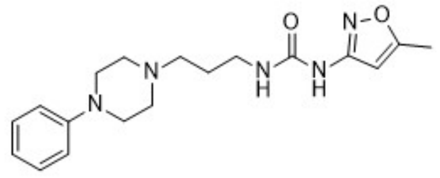

**S3**

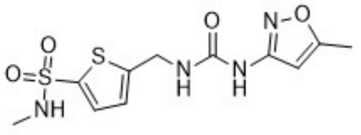

**S4**

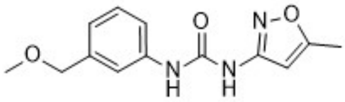

**S5**

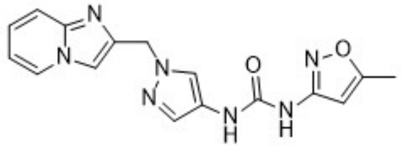

**S6**

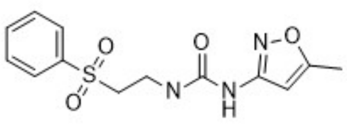

**S7**

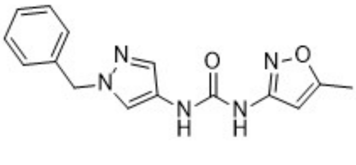

**S8**

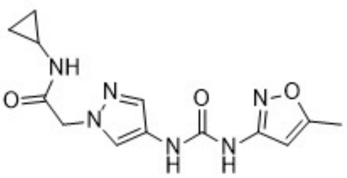

**S9**

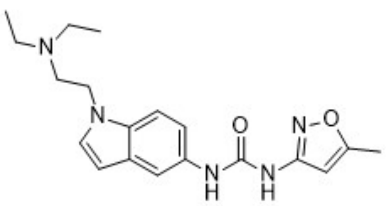

**S10**

**Figure S2.** Commercial compounds identified to bind to Kalirin-Rac1 in XChem screen.

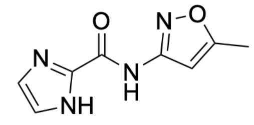

**S11**

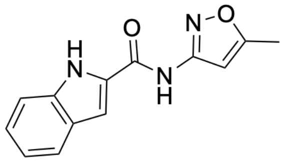

**S12**

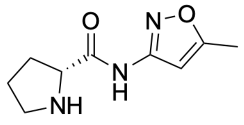

**S13**

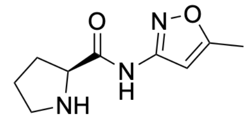

**S14**

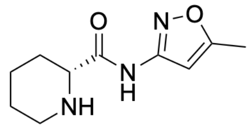

**S15**

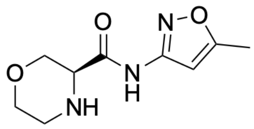

**S16**

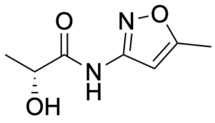

**S17**

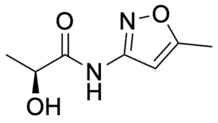

**S18**

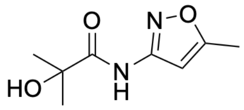

**S19**

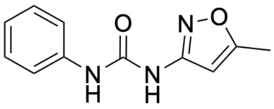

**S20**

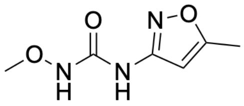

**S21**

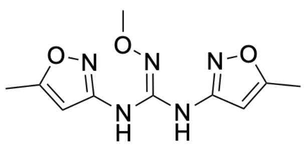

**S22**

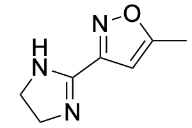

**S23**

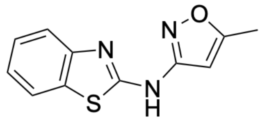

**S24**

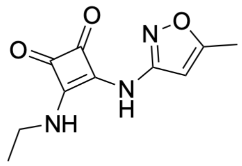

**S25**

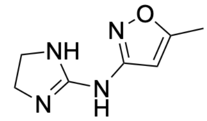

**S26**

**Figure S3.** Urea bioisosteres designed using SwissBioisostere and were synthesized in house.

| Compound | Structure |
|---|---|
| 3 | 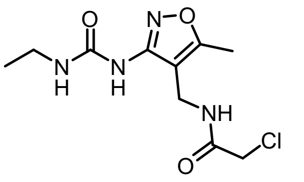 |
| 4 | 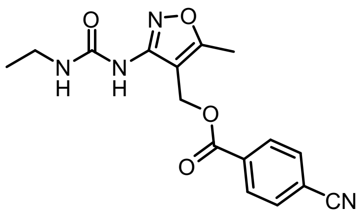 |
| 5 | 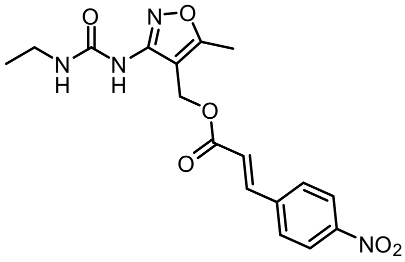 |
| 6 | 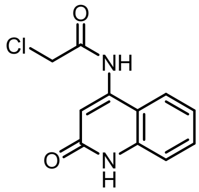 |
| 7 | 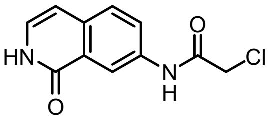 |

**Figure S4.** Covalent analogues explored in the initial hit stage of discovery.

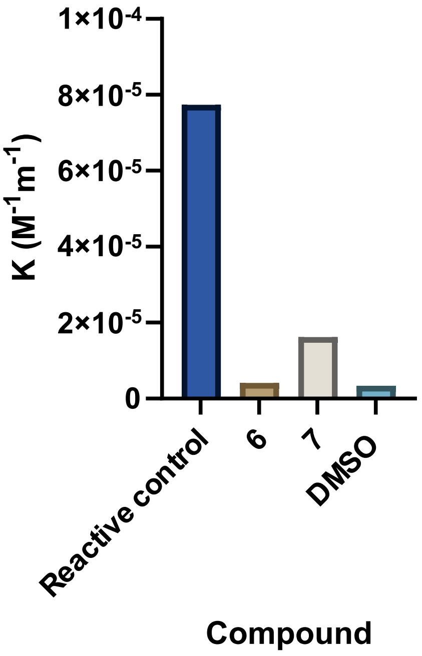

**Figure S5**. Reactivity of covalent analogues **6** and **7**.

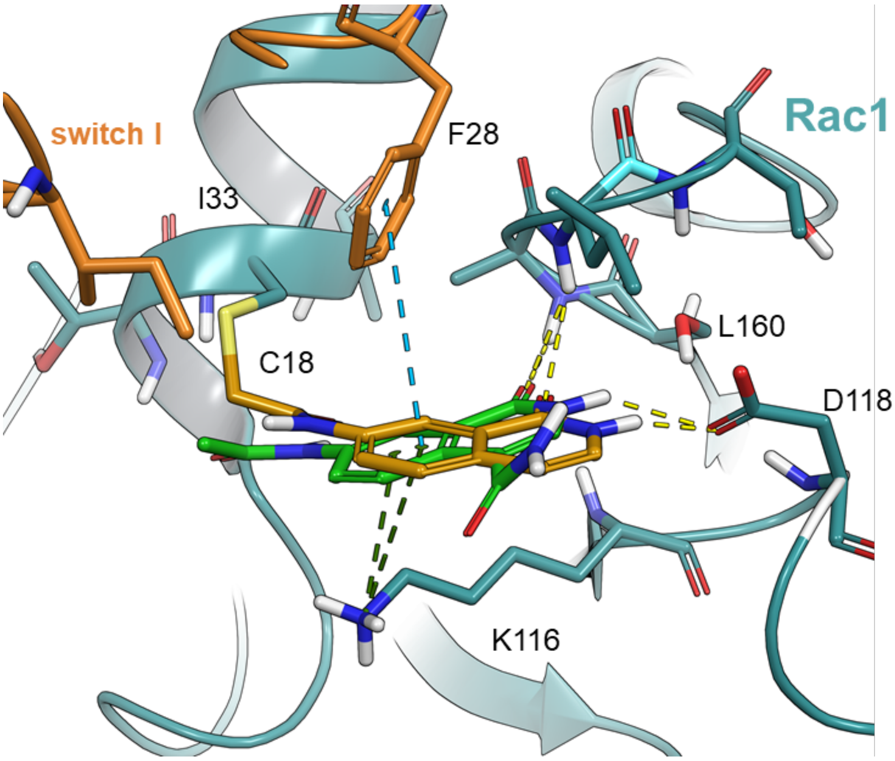

**Figure S6.** X-ray crystal structure of compound 7 (orange sticks) bound to the Kalirin-Rac1 complex (PDB: 9IG1), overlaid with the original DOCKovalent hit (green sticks).

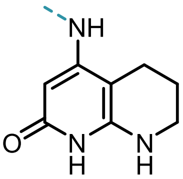

| compound | R |
| --- | --- |
| **S27** | 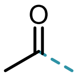 |
| **S28** | 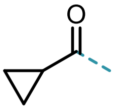 |
| **S29** | 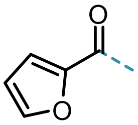 |
| **S30** | 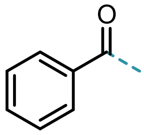 |
| **S31** | 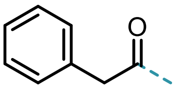 |
| **S32** | H |
| **S33** | Me |
| **S34** | 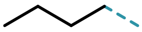 |
| **S35** | 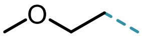 |

**Table S1**. Non-covalent analogues of MC-278 synthesised and tested against Kalirin-Rac1.

| Entry | Rac1 (μM) | Kalirin-GEF(1) (μM) | Buffer additions | Temp (°C) | Time (h) | MC-278-Rac1 (%) |
|---|---|---|---|---|---|---|
| **1** | 1.25 | 0.03 | BSA, DTT | r.t. | 2 | 0 |
| **2** | 1.25 | 0.03 | - | r.t. | 2 | 0 |
| **3** | 1.25 | 1.25 | - | r.t. | 2 | 0 |
| **4** | 1.25 | 0.03 | - | r.t. | 24 | 0 |
| **5** | 1.25 | 0.03 | - | 4 | 24 | 0 |
| **6** | 1.25 | 1.25 | - | 4 | 24 | 79 |
| **7** | 1.25 | 0 | - | 4 | 24 | 0 |

**Tables S2.** Optimisation of the protein mass spectrometry assay.

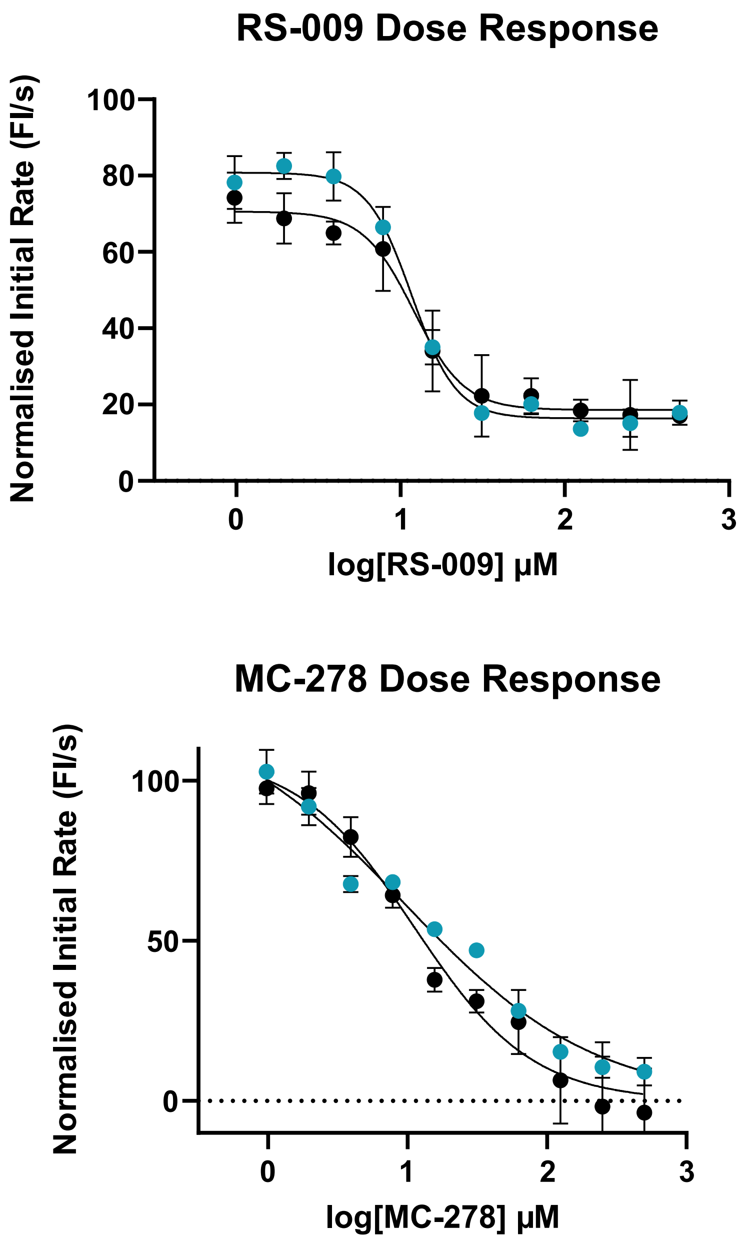

**Figure S7.** Dose response curves of RS-009 and MC-278.

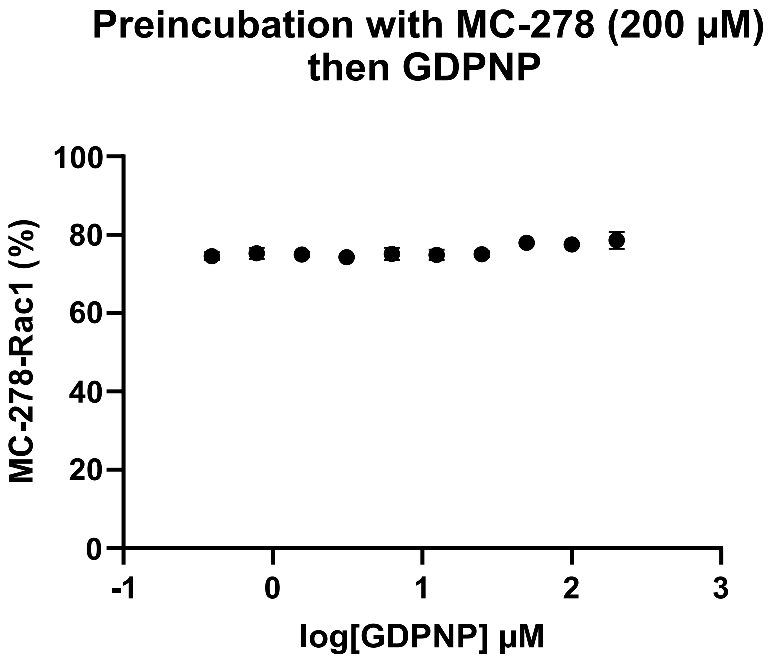

**Figure S8.** Percentage of MC-278 modified Rac1 after incubation of Kalirin and Rac1 with MC-278 (200 µM), followed by addition of GDPNP (n = 2).

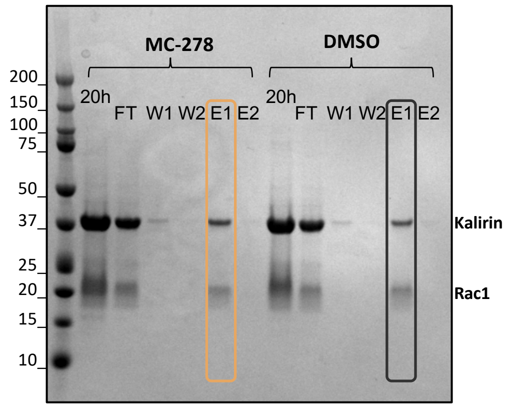

**Figure S9.** Analysis of reaction mixtures and Strep-Tactin purification by Coomassie-stained SDS-PAGE. MC-278 (200-500 µM) or DMSO was incubated with biotinylated RAC1 ($Rac1_B$, 35 µM) and Kalirin (50 µM) for 40 h, followed by purification using Strep-Tactin magnetic beads. Complexes, MC-278-$Rac1_B$-K and DMSO-$Rac1_B$-K, were washed with wash buffer twice (W1, W2) and eluted with 50 mM biotin in buffer twice (E1, E2).

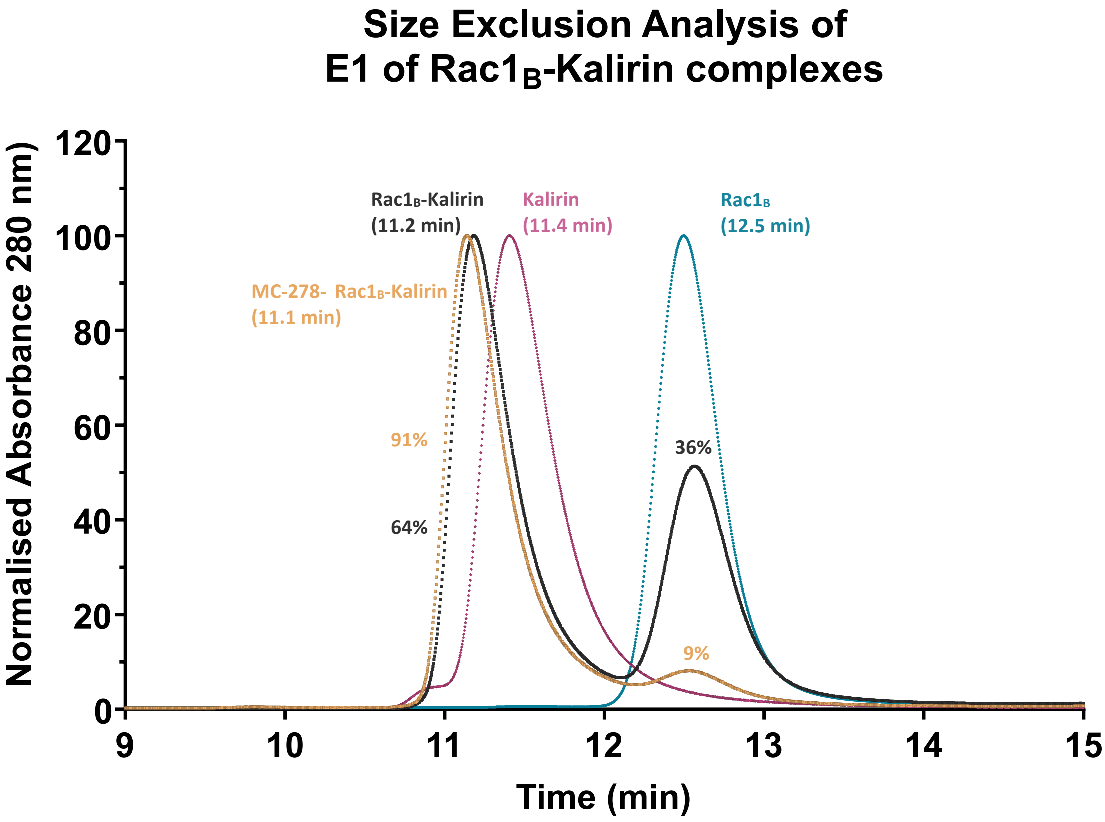

**Figure S10.** Overlay of the size-exclusion chromatograms (SEC) from free $Rac1_B$, Kalirin and the E1 fractions obtained following Strep-Tactin purification of MC-278-$Rac1_B$-Kalirin and $Rac1_B$-Kalirin complexes. Formation of higher-molecular-weight protein complexes is indicated by a left-shifted major peak retention time. The relative area under the curve (AUC) for the MC-278-$Rac1_B$-Kalirin complex (91%) vs free $Rac1_B$ (9%) exceeds that observed for the control $Rac1_B$-Kalirin complex (64%) vs free $Rac1_B$ (36%).

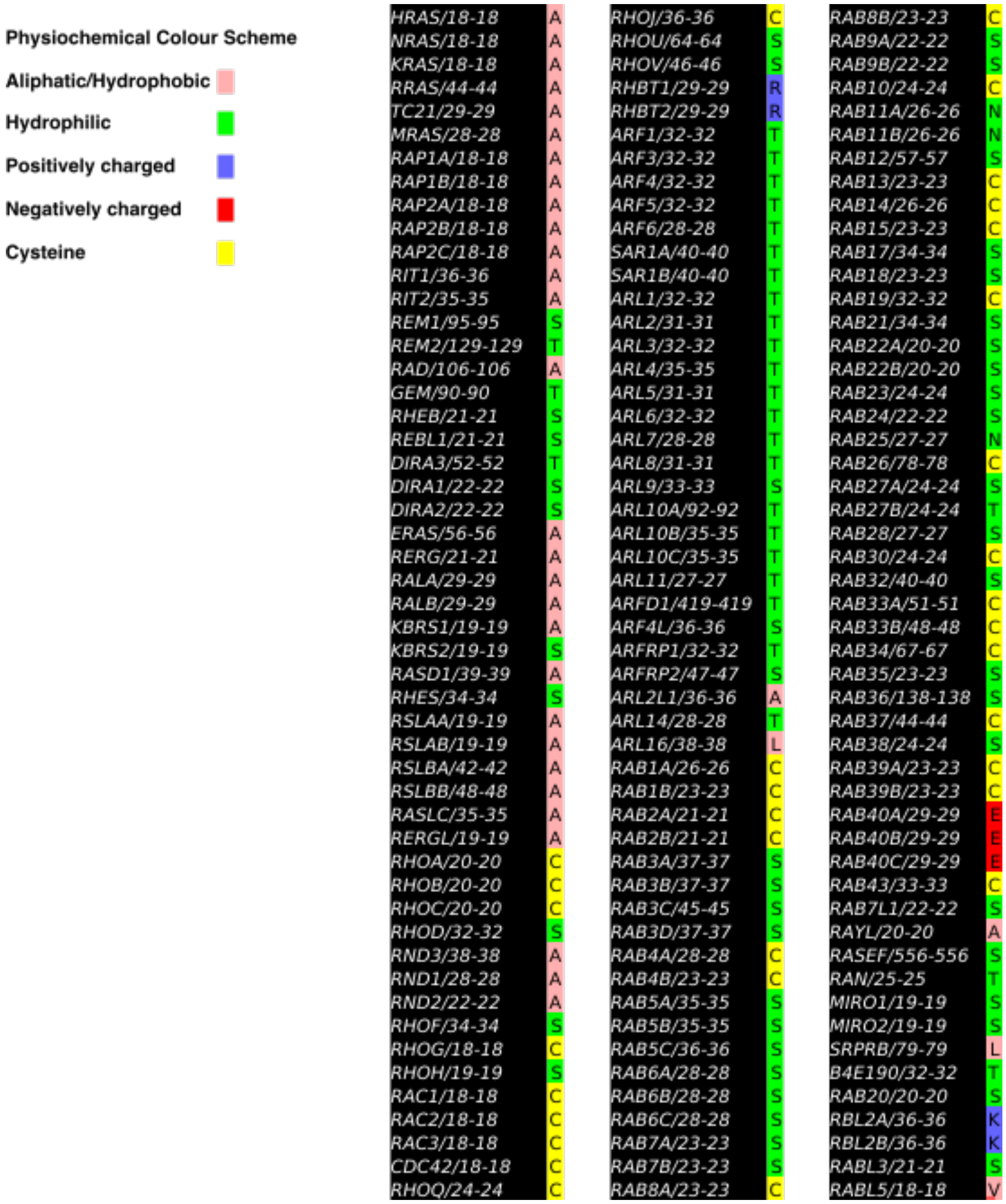

**Figure S11**. Sequence alignment of 153 members of the RAS superfamily of residue 18 (Rac1 numbering) using ClusterX in Jalview. The conserved residue in this position is serine (S), present in 32% of family members, followed by Cysteine (21%), Alanine (20%) and Threonine (18%).

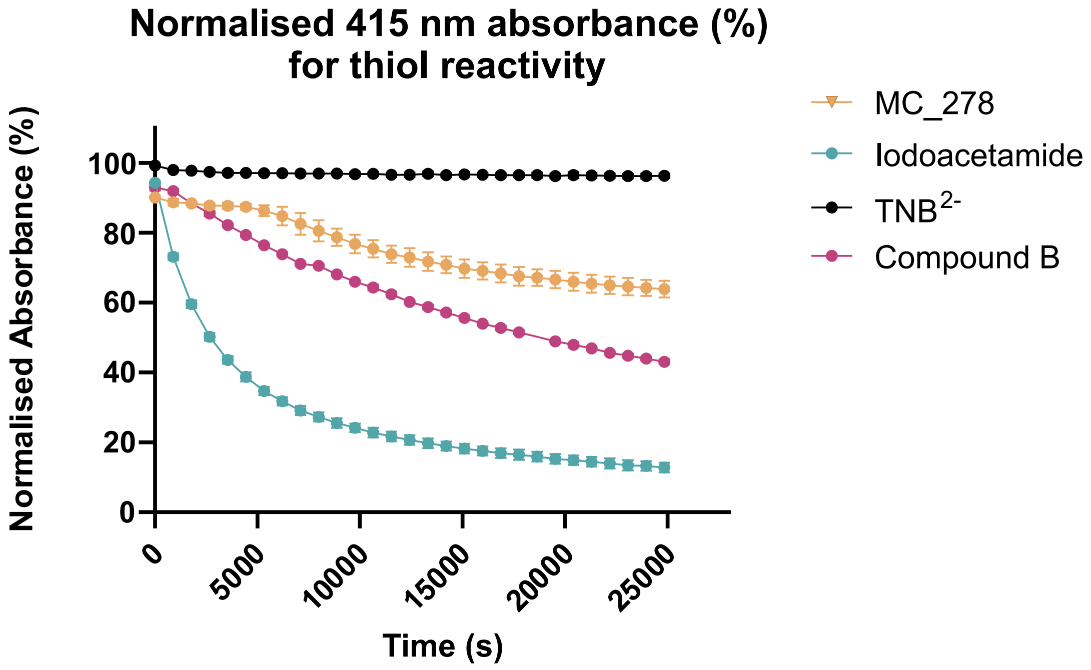

**Figure S12.** Example of normalised thiol-reactivity assay traces monitored at 415 nm for MC-278, control compound B (2-chloro-1-(4-methylpiperazin-1-yl)ethan-1-one) and iodoacetamide, with $TNB^{2-}$ alone serving as a control. MC-278 exhibited biphasic behaviour profile with an unexplained rate increase after 80 minutes. The data and was therefore fitted using a two-phase kinetic model with the initial rate used in comparison with controls (Table S3).

**Table S3.** Mean second-order rate constants ($k$; n = 3, and n = 1 for control compound B) and corresponding standard deviations (SD) obtained by linear regression of the kinetic traces. For MC-278, rate constants were determined separately for each of the two observed kinetic phases. Iodoacetamide exhibited approximately 50-fold higher reactivity than MC-278 in the initial phase.

| | **Initial rate constant** $k$ **($M^{-1}$ $s^{-1}$)** | | **Secondary rate constant** $k$ **($M^{-1}$ $s^{-1}$)** | |
|---|---|---|---|---|
| **Compound** | **Mean** | **SD** | **Mean** | **SD** |
| **MC-278** | $2.32\ x\ 10^{-6}$ | $1.97\ x\ 10^{-7}$ | $1.23\ x\ 10^{-5}$ | $2.94\ x\ 10^{-6}$ |
| **Iodoacetamide** | $9.39\ x\ 10^{-5}$ | $9.28\ x\ 10^{-6}$ | na | na |
| **Control B** | $1.85\ x\ 10^{-5}$ | $1.1\ x\ 10^{-7}$ | na | na |

### Experimental

#### Protein expression and purification

All synthetic DNA and primers were obtained from Eurofin Genomics. DNA constructs were amplified using PCR. Vectors and constructs were prepared for LIC cloning by creating suitable overhangs using T4 polymerase. A construct of the DH1 domain of rat Kalirin (UniProt P97924, residues 1232-1411) was cloned into a pNic28[1] vector (in-house) including a N-terminally fused with a deca His-tag followed by a Tobacco Etch Virus (TEV) recognition site. construct of the DH1-PH1 domain of human Kalirin (UniProt O60229, residues 1259-1590) was cloned into a pNic28 vector (in-house) including a N-terminally fused with a hexa His-tag followed by a Tobacco Etch Virus (TEV) recognition site. A C-terminally truncated construct of human Rac1 (residues 1-177) was cloned into pNic28-StIIT2 including a N-terminally fused Strep-II tag followed by a TEV recognition site.

For protein expression, plasmids were transformed into *E. coli* BL21(DE3)-R3-pRARE2 and stored in 50% glycerol at -80 °C until required. 10 mL overnight culture grown in LB containing 50 µg/mL kanamycin and 34 µg/mL chloramphenicol at 37 °C were used to inoculate 1 L of AIM-TB (ForMedium) supplemented to a final concentration with 20 g/L glucose, 0.01% Antifoam 204 (Sigma), 50 µg/mL kanamycin and 25 µg/mL chloramphenicol in a 2.5 L baffled flask (Ultrayield, Thomsen). The large scale cultures were grown for 4 h at 37 °C, 250 rpm before the temperature was reduced to 25 °C 250 rpm for 16 h. The cells were harvested by centrifugation at 4000 g for 20 min and the pellet stored at -20 °C. Cell pellets were resuspended in lysis buffer (10 mM HEPES pH 7.5, 500 mM NaCl, 5% glycerol, 0.5 mM TCEP, 0.5 mg/ml Lysozyme, 0.1 mg/ml benzonase and protease inhibitors [Calbiochem EDTA-free Protease Inhibitor Cocktail Set III]) to a final volume of 20 mL per 5 pellet. Triton X-100 to a final concentration of 2% was added and cells were lysed using three freeze-thaw cycles at -80 °C.

For purification of Kalirin DH1 and Kalirin DH1-PH1, imidazole was added to a final concentration of 20 mM post-lysis and centrifuged at 4000 g for 1 h. The clarified supernatant was loaded onto His GraviTrap columns (GE Healthcare) pre-equilibrated in wash buffer (10 mM HEPES pH 7.5, 500 mM NaCl, 5% Glycerol, 20 mM Imidazole, 0.5 mM TCEP) and washed extensively. The protein was eluted using elution buffer

(10 mM HEPES pH 7.5, 500 mM NaCl, 5% Glycerol, 500 mM Imidazole, 0.5 mM TCEP) before application and elution from a PD-10 desalting column (GE Healthcare) using wash buffer. TEV protease (1 mg per 10 mg eluted protein) was added and tag digestion was performed overnight at 4 °C. The protein, the affinity tag and TEV protease were separated using reverse IMAC. Relevant fractions were combined, concentrated to ~30 mg/mL and loaded onto a Yarra SEC-2000 (Phenomenex) gel filtration column pre-equilibrated in 10 mM HEPES pH 7.5, 500 mM NaCl, 5% Glycerol, 0.5 mM TCEP. The peaks corresponding to Kalirin were analysed by SDS-PAGE, pooled, concentrated to 10-20 mg/mL using a 10 kDa MWCO concentrator (Vivaspin, Cytiva), flash frozen in liquid nitrogen and stored at -80 °C.

For the purification of Rac1, lysed cells were clarified by centrifuging at 4000 g for 1 h. The clarified supernatant was loaded onto Strep-Tactin-XT columns (IBA Lifesciences) pre-equilibrated in wash buffer (10 mM HEPES pH 7.5, 500 mM NaCl, 5% Glycerol, 0.5 mM TCEP) and washed extensively. The protein was eluted using elution buffer (10 mM HEPES pH 7.5, 500 mM NaCl, 5% Glycerol, 0.5 mM TCEP, 50 mM D-Biotin). TEV protease (1 mg per 10 mg eluted protein) was added and tag digestion was performed overnight at 4 °C. The protein, the affinity tag and TEV protease were separated using reverse IMAC. Relevant fractions were combined, concentrated to ~30 mg/mL and loaded onto a Yarra SEC-2000 (Phenomenex) gel filtration column pre-equilibrated in 10 mM HEPES pH 7.5, 500 mM NaCl, 5% Glycerol, 0.5 mM TCEP. The peaks corresponding to Rac1 were analysed by SDS-PAGE, pooled, concentrated to 15 mg/mL using a 10 kDa MWCO concentrator, flash frozen in liquid nitrogen and stored at -80 °C.

To generate Kalirin-Rac1 complex, purified Kalirin and Rac1 were mixed in a 1:1 molar ratio and incubated on ice for 1 h. The complex was concentrated to 60 mg/mL using a 30 kDa MWCO concentrator (Vivaspin, Cytiva). For the GDP-bound complex, the protein was loaded onto a Yarra SEC-2000 (Phenomenex) gel filtration column pre-equilibrated in 10 mM HEPES pH 7.5, 500 mM NaCl, 5% Glycerol, 0.5 mM TCEP. The peaks corresponding to Kalirin-Rac1 were analysed by SDS-PAGE, pooled, concentrated to ~10 mg/mL using a 30 kDa MWCO concentrator, flash frozen in liquid nitrogen and stored at -80 °C. For the GDP-free complex, 40 µL of 10 mM β,γ-Methyleneguanosine 5'-triphosphate sodium salt (GppCp, Sigma) and 4 Units of Alkaline phosphatase (Roche) were added to the concentrated protein. 20 µL of the

10x exchange buffer was added (final concentration 2M $(NH_4)_3PO_4$, 10 µM ZnCl) and the mixture incubated at 4 °C for 3 h. The final concentration of Kalirin-Rac1 was 1 mM. Degradation of GDP to GMP and guanosine was monitored by HPLC (Agilent 1200, reverse-phase C-18) over 15 min in buffer containing 100 mM potassium phosphate pH 6.5, 10 mM tetrabutyl phosphonium bromide and 5% MeOH. Following complete degradation of GDP, 0.008 Units of snake venom phosphodiesterase I (Sigma) was added. The reaction was left overnight at 4 °C and the degradation of GppCp to GMP and guanosine monitored by HPLC. The protein was diluted to 30 mg/mL and loaded onto a Yarra SEC-2000 gel filtration column pre-equilibrated in 10 mM HEPES pH 7.5, 500 mM NaCl, 5% Glycerol, 0.5 mM TCEP. The peaks corresponding to Kalirin-Rac1 were analysed by SDS-PAGE, pooled, concentrated to 10.4 mg/mL using a 30 kDa MWCO concentrator, flash frozen in liquid nitrogen and stored at -80 °C.

Biotinylated Rac1 expression and purification was performed following the general procedure described above, with the following modifications. The Rac1 gene (1-177) was purchased from Twist Biosciences with an N-terminal Tev cleavable 6xHis and twin strep tag and a C-terminal Avi tag using a pNIC-CTH0 vector, omitting the plasmid tag using a stop codon. The plasmid was transformed into BL21(DE3)-R3-pRARE2-BirA Rosetta cells and expressed in Terrific Broth supplemented with 1 x TB salts, (50 mL), kanamycin (100 µL), chloroamphenicol (100 µL), 0.4% glycerol, 70mg/L biotin in 10mM bicine pH8.3, and induced with 0.4mM IPTG. Induced cultures were incubated overnight at 18 °C before harvesting by centrifugation and storing at -80 °C.

The pellet was suspended in base buffer (10 mM HEPES pH 7.5, 500 mM NaCl, 5% glycerol, 0.5 mM TCEP) with lysozyme (0.5 mg/mL), Benzonase (0.1 mg/mL), protease inhibitor cocktail (Merck), lysed by sonication and insoluble material excluded by centrifugation at 15000g . Purification was performed by Ni-affinity chromatography (Ni Sepharose FF Cytiva) and eluted in base buffer plus 500mM Imidazole. Eluted proteins were cleaved with Tev protease and dialysed overnight into base buffer. Cleaved protein was purified by Ni-affinity rebinding, concentrated using a spin concentrator (Amicon 10kDa MWCO) and further purified by size-exclusion chromatography using a S75 HiLoad 16/60 Superdex column equilibrated into base buffer. Protein was concentrated, aliquoted, flash frozen and stored at -80 °C. Intact mass spectrometry was used to confirm expected mass and purity was shown by SDS-PAGE.

**Crystallization**

Kalirin(1)-Rac1 GDP-bound structure (PDB: 5O33)

GDP-bound Kalirin-Rac1 crystals were grown by sitting-drop vapour diffusion at 20 °C with a 2:1 ratio of protein (15.6 mg/mL) to a reservoir solution consisting of 0.1 M citrate pH 4.2, 0.2 M NaCl and 18 % PEG 8K at 20 °C. Crystals were cryo-protected with 25% ethylene glycol and flash-cooled in liquid nitrogen. Diffraction data were collected at Diamond Light Source (DLS) Beamline I04-1 and indexed, integrated and scaled using the automated Xia2 pipeline.[3] Molecular replacement, using trio/Rac1 (PDB code: 2NZ8) as a search template, and subsequent model building and refinement was conducted using the PHENIX[4] software suite and manual model correction in Coot.[5]

**Co-crystallisation with covalent compounds**

**6** was incubated with nucleotide-free Kalirin-Rac1 (10.4 mg/mL) to a final concentration of 1 mM. Crystals were grown by mixing 50 nL of labelled Kalirin-Rac1 (10.4 mg/mL) with 100 nL of reservoir solution consisting of 0.1 M citrate pH 4.7, 0.8 M LiCl and 25% PEG 6K at 20 °C. The crystals were cryo-protected using 15% ethylene glycol. Diffraction data was collected at the I03 beamline at DLS.

**7** was incubated with nucleotide-free Kalirin-Rac1 (10.5 mg/mL) to a final concentration of 1 mM. Crystals were grown by mixing 75 nL of labelled Kalirin-Rac1 (10.5 mg/mL) with 75 nL of reservoir solution consisting of 0.1 M bis-tris pH 5.5, 0.2 M NaCl and 25% PEG 3350 at 20 °C. The crystals were cryo-protected using 20% DMSO. Diffraction data was collected at the I04-1 beamline at DLS.

MC-278 was incubated with Kalirin-Rac1 (15.0 mg/mL) and crystallisation grown by sitting-drop vapour diffusion with 100 nL of protein and 50 nL reservoir solution consisting of 0.1 M citrate pH 4.2, 0.2 M NaCl and 18 % PEG 8K at 20 °C. After 6 days, the droplet was added 13 µL reservoir solution, 6 µL ethylene glycol, and 1 µL MC-278 (100 mM in DMSO) to give a final concentration of 5 mM and left to incubate at ambient temperature for 2 h. The resulting crystals were flash cooled in liquid nitrogen. X-ray diffraction data was collected at Diamond Light Source on the I03-1 beamline.

RS-009 was incubated with nucleotide-free Kalirin-Rac1 (10.5 mg/mL) to a final concentration of 1 mM. Crystals were grown by mixing 75 nL of labelled Kalirin-Rac1 (10.5 mg/mL) with 75 nL of reservoir solution consisting of 0.1 M bis-tris pH 5.5, 0.1 M NaCl and 27% PEG 3350 at 20 °C. The crystals were cryo-protected using 20% DMSO.

For compound **6** molecular replacement was conducted using PHASER from PHENIX[4] software suite. Jligand from the CCP4 software suite was used to install the covalent linkage.[6] The structure was iteratively refined using the PHENIX software suite and manual model correction performed in Coot.[5] For compounds 7 and RS-009, molecular replacement was conducted using PHASER MR from CCP4 suite. ACEDRG[7] and REFMAC[8] from the CCP4 suite was used to install the covalent linkage. The structure was iteratively refined using REFMAC, and manual model correction performed in Coot.

For MC-278, the structure was phased and refined using PHENIX, the covalent bond modelled with REFMAC, and manual model correction performed in Coot.

#### XChem screening

##### Initial high-throughput screen

Hundreds of GDP-bound Kalirin-Rac1 crystals were mass produced using reservoir solutions of 0.1 M citrate pH 4.2, 0.1-0.4 M NaCl, and 16-26% PEG8K at 20 °C. For soaking, 37.5 nL of fragments from the DSI-Poised,[9] Covalent,[10] OxXChem,[11] Leeds3D[12] and Maybridge[13] libraries were added to each crystallisation drop using an ECHO acoustic liquid dispenser at DLS beamline I04-1, with a final concentration of 100 mM fragment and 20% DMSO. Crystals were soaked with compounds for 1 h before being harvested using SHIFTER[14] and flash-cooled in liquid nitrogen. Diffraction data was collected using the ‘automation-unattended’ mode and automatically processed on the I04-1 beamline. Crystals typically diffracted between 1.5-2 Å. The XChemExplorer[15] platform was used for initial structure solution and parallel molecular replacement was performed using DIMPLE with the GDP-bound

Kalirin-Rac1 (PDB code: 5O33) as a homology model, followed by subsequent electron density analysis and hit identification using PanDDA.[16] Model building and refinement was performed using Coot and REFMAC integrated within the XChemExplorer pipeline.

**Urea analogue screens**

GDP-bound Kalirin-Rac1 crystals were made with reservoir solution of 0.1M citrate pH 4.4, 0.8 M LiCl and 24% PEG 6K at 20 °C. Hundreds of nucleotide-free Kalirin-Rac1 crystals were made with reservoir solutions of 0.1 M bis-tris pH 5.5, 0-0.2 M NaCl, and 22-27% PEG 3350. 37.5 nL of each fragment from the purchased Molport library was dispensed, resulting in final concentration of 20% DMSO and 20 mM fragment per well. GDP-bound or nucleotide-free Kalirin-Rac1 was used as a homologous model in DIMPLE. The crystals typically diffracted between 1.7 and 2.5 Å.

**6-co-crystallisation screen**

**6**-labelled Kalirin-Rac1 crystals were made with reservoir solutions of 0.1 M bis-tris pH 5.5, 0-0.2 M NaCl, and 22-27% PEG 335 at 20 °C 0. 27.5 nL of each fragment from the DSI-Poised library was dispensed, resulting in final concentration of 15% DMSO and 62 mM fragment per well. **106** labelled Kalirin-Rac1 was used as a homologous model in DIMPLE. The crystals typically diffracted between 1.8 and 2.5 Å.

**Pharmacophore screen**

Nucleotide-free Kalirin-Rac1 crystals were made with reservoir solutions of 0.1 M bis-tris pH 5.5, 0-0.2 M NaCl, and 22-27% PEG 3350 at 20 °C. 27.5-37.5 nL of each fragment from the purchased Enamine library was dispensed, resulting in final concentration of 15-20% DMSO and 38.7 mM fragment per well. Nucleotide-free Kalirin-Rac1 was used as a homologous model in DIMPLE. The crystals typically diffracted between 1.8 and 2.5 Å.

**Crystallographic data tables**

GDP-bound Kalirin-Rac1

| PDB code | 5O33 |
|---|---|
| Beamline | DLS I04-1 |
| Wavelength (Å) | 0.92819 |
| Space group | P $6_5$ 2 2 |
| a, b, c (Å) | 63.255,63.255, 346.704 |
| a, b, g (°) | 90, 90, 120 |
| **Data collection** | |
| Resolution (Å) | 57.78-1.64 (1.67-1.64) |
| $R_{merge}$ | 0.0971 (2.643) |
| I/s(I) | 11.2 (1.2) |
| $CC_{1/2}$ | 0.999 (0.493) |
| Completeness (%) | 100 (99.7) |
| Redundancy | 18.5 (18.8) |
| No. reflections | 52137 (2571) |
| **Refinement** | |
| Resolution (Å) | 1.64 |
| No.reflections | 49371 |
| Rwork / Rfree | 0.2067/0.2398 |
| Average B-factor ($Å^2$) | 43.0 |
| RMSD: bonds (Å) | 0.009 |
| RMSD: bond angles (°) | 1.072 |
| Ramachandran favoured (%) | 97 |
| Ramachandran outliers (%) | 0.3 |

Urea analogues bound to Kalirin-Rac1

| PDB code | 5QQD | 5QQE | 5QQF |
|---|---|---|---|
| Ligand | **1** | **S1** | **S2** |
| Beamline | DLS I04-1 | DLS I04-1 | DLS I04-1 |
| Wavelength | 0.91587 | 0.91587 | 0.91587 |
| Space group | P $6_5$ 2 2 | P $6_5$ 2 2 | P $6_5$ 2 2 |
| a, b, c (Å) | 62.836, 62.836,342.580 | 62.807,62.807,342.65 | 62.49,62.49,341.271 |
| a, b, g (°) | 90, 90, 120 | 90,90,120 | 90, 90, 120 |
| **Data collection** | | | |
| Resolution | 57.10 - 1.91 | 57.11 – 1.95 | 56.88 – 2.26 |
| | (1.96 - 1.91) | (2.05 – 1.95) | (2.39 – 2.26) |
| $R_{merge}$ | 0.510 (2.691) | 0.275 (7.137) | 0.137 (1.575) |
| I/s(I) | 10.4 (1.6) | 8.8 (1.1) | 10.4 (1.4) |
| $CC_{1/2}$ | 0.998 (0.773) | 0.998 (0.433) | 0.998 (0.417) |
| Completeness (%) | 99.7 (98.0) | 97.2 (99.9) | 99.2 (98.1) |
| Redundancy | 33.9 (18.4) | 18.4 (18.5) | 10.7 (8.6) |
| No. reflections | 32694 (2338) | 30148 (4307) | 19514 (2696) |
| **Refinement** | | | |
| Resolution (Å) | 1.91 | 1.95 | 2.26 |
| No.reflections | 30853 | 28451 | 18445 |
| Rwork / Rfree | 0.2059/0.2586 | 0.218/0.284 | 0.1977/0.2699 |
| Average B-factor (Å$^2$) | 35.5 | 45.5 | 53.0 |
| RMSD: bonds (Å) | 0.009 | 0.009 | 0.012 |
| RMSD: bond angles (°) | 1.526 | 1.524 | 1.570 |
| Ramachandran favoured (%) | 96 | 95 | 96 |
| Ramachandran outliers (%) | 0.3 | 0.6 | 0.6 |
| Clashscore | 6 | 7 | 4 |

| | | | |
|---|---|---|---|
| RSRZ outliers (%) | 2.2 | 1.7 | 1.1 |
| Ligand RSCC | 0.86 | 0.88 | 0.88 |

| PDB code | 5QQG | 5QQH | 5QQI |
|---|---|---|---|
| Ligand | **S3** | **S4** | **S5** |
| Beamline | DLS I04-1 | DLS I04-1 | DLS I04-1 |
| Wavelength | 0.91587 | 0.91587 | 0.91587 |
| Space group | P $6_5$ 2 2 | P $6_5$ 2 2 | P $6_5$ 2 2 |
| a, b, c (Å) | 62.785,62.785, 344.573 | 62.659,62.659,342. 820 | 62.505,62.505,341. 300 |
| a, b, g (°) | 90, 90, 120 | 90,90,120 | 90, 90, 120 |
| **Data collection** | | | |
| Resolution | 57.43 – 2.23 | 57.14 – 2.09 | 56.88 – 2.08 |
| | (2.29 – 2.23) | (2.15 – 2.09) | (2.13 – 2.08) |
| $R_{merge}$ | 0.210 (3.115) | 0.373 (4.132) | 0.802 (2.454) |
| Mean I/s(I) | 11.2 (1.2) | 9.5 (1.7) | 10.2 (1.7) |
| $CC_{1/2}$ | 0.999 (0.333) | 0.998 (0.619) | 0.998 (0.745) |
| Completeness (%) | 100 (100) | 99.8 (100) | 99.7 (100) |
| Redundancy | 15.6 (13.7) | 36 (27.7) | 32.7 (18.7) |
| No. reflections | 20947 (2929) | 25043 (1740) | 25188 (1772) |
| **Refinement** | | | |
| Resolution (Å) | 2.23 | 2.09 | 2.08 |
| No. reflections | 19788 | 22660 | 23797 |
| Rwork / Rfree | 0.1970/0.2742 | 0.1995/0.2631 | 0.2188/0.2767 |
| Average B-factor ($Å^2$) | 52.7 | 41.3 | 41.4 |
| RMSD: bonds (Å) | 0.012 | 0.017 | 0.016 |
| RMSD: bond angles (°) | 1.572 | 1.763 | 1.738 |
| Ramachandran favoured (%) | 96 | 98 | 97 |
| Ramachandran outliers (%) | 0.6 | 0.3 | 0.3 |
| Clashscore | 4 | 6 | 5 |

| | | | |
|---|---|---|---|
| RSRZ outliers (%) | 0.3 | 0.3 | 5.3 |
| Ligand RSCC | 0.93 | 0.86 | 0.80 |

| PDB code | 5QQJ | 5QQK | 5QQL |
|---|---|---|---|
| Ligand | **S6** | **S7** | **S8** |
| Beamline | DLS I04-1 | DLS I04-1 | DLS I04-1 |
| Wavelength | 0.91587 | 0.91587 | 0.91587 |
| Space group | P $6_5$ 2 2 | P $6_5$ 2 2 | P $6_5$ 2 2 |
| a, b, c (Å) | 62.568,62.568,342.039 | 62.584,62.584,341.958 | 62.308,62.308,340.950 |
| a, b, g (°) | 90, 90, 120 | 90, 90, 120 | 90, 90, 120 |
| **Data collection** | | | |
| Resolution | 54.24 – 1.90 | 56.99 – 2.24 | 56.83 – 2.25 |
| | (1.95 – 1.90) | (2.36 – 2.24) | (2.39 – 2.25) |
| $R_{merge}$ | 0.197 (3.032) | 0.456 (4.818) | 0.297 (4.116) |
| Mean I/s(I) | 9.4 (1.6) | 9.2 (1.6) | 8.6 (1.1) |
| $CC_{1/2}$ | 0.998 (0.697) | 0.995 (0.517) | 0.998 (0.404) |
| Completeness (%) | 99.6 (100) | 100 (100) | 100 (99.9) |
| Redundancy | 18.2 (18.6) | 17.8 (18.4) | 18.4 (19.6) |
| No. reflections | 32844 (2340) | 20517 (2862) | 19877 (3096) |
| **Refinement** | | | |
| Resolution (Å) | 1.90 | 2.24 | 2.25 |
| No. reflections | 30971 | 19384 | 18781 |
| Rwork / Rfree | 0.2118/0.2687 | 0.1999/0.2656 | 0.1911/0.2702 |
| Average B-factor ($Å^2$) | 37.6 | 44.7 | 54.9 |
| RMSD: bonds (Å) | 0.008 | 0.014 | 0.012 |
| RMSD: bond angles (°) | 1.524 | 1.618 | 1.516 |
| Ramachandran favoured (%) | 94 | 95 | 95 |
| Ramachandran outliers (%) | 0.9 | 0.3 | 0.6 |
| Clashscore | 7 | 4 | 3 |

| | | | |
|---|---|---|---|
| RSRZ outliers (%) | 1.4 | 0.6 | 0.6 |
| Ligand RSCC | 0.82 | 0.87 | 0.922 |

| PDB code | 5QQM | 5QQN |
| --- | --- | --- |
| Ligand | **S9** | **S10** |
| Beamline | DLS I04-1 | DLS I04-1 |
| Wavelength | 0.91587 | 0.91587 |
| Space group | P $6_5$ 2 2 | P $6_5$ 2 2 |
| a, b, c (Å) | 62.498,62.498,342.286 | 62.492,62.492,341.798 |
| a, b, g (°) | 90, 90, 120 | 90,90,120 |
| **Data collection** | | |
| Resolution | 57.05 – 2.02 | 54.18 – 2.26 |
| | (2.08 – 2.02) | (2.38 – 2.26) |
| $R_{merge}$ | 0.342 (3.640) | 0.439 (3.63) |
| Mean I/s(I) | 10.8 (1.8) | 11.8 (2.8) |
| $CC_{1/2}$ | 0.946 () | 0.997 (0.830) |
| Completeness (%) | 99.8 (98.3) | 99.9 (99.9) |
| Redundancy | 34.6 (17.7) | 17.5 (19.4) |
| No. reflections | 27501 (2053) | 19972 (2787) |
| **Refinement** | | |
| Resolution (Å) | 2.02 | 2.26 |
| No. reflections | 25975 | 18877 |
| Rwork / Rfree | 0.2113/0.2731 | 0.2050/0.2786 |
| Average B-factor (Å$^2$) | 38.0 | 43.4 |
| RMSD: bonds (Å) | 0.017 | 0.008 |
| RMSD: bond angles (°) | 1.804 | 1.537 |
| Ramachandran favoured (%) | 96 | 97 |
| Ramachandran outliers (%) | 0.3 | 0.6 |
| Clashscore | 4 | 8 |

| | | |
|---|---|---|
| RSRZ outliers (%) | 0.6 | 1.4 |
| Ligand RSCC | 0.88 | 0.78 |

Co-crystallisation of covalent compounds to Kalirin-Rac1

| PDB code | 9IFK | 9IG1 |
|---|---|---|
| Ligand | **6** | **7** |
| Beamline | DLS I03 | DLS I04-1 |
| Wavelength | 0.97625 | 0.91587 |
| Space group | P $6_5$ 2 2 | P $6_5$ 2 2 |
| a, b, c (Å) | 62.48,62.48,339.60 | 62.28,62.28,339.67 |
| a, b, g (°) | 90, 90, 120 | 90, 90, 120 |
| **Data collection** | | |
| Resolution | 56.6 – 1.34 | 53.94 – 1.65 |
| | 1.36 – 1.34 | (1.68 – 1.65) |
| $R_{merge}$ | 0.196 (13.0) | 0.11 (1.789) |
| Mean I/s(I) | 19.4 (0.9) | 13.1 |
| $CC_{1/2}$ | 0.99 (0.352) | 0.998 (0.653) |
| Completeness (%) | 100 (100) | 100 (99.9%) |
| Redundancy | 72.0 (63.7) | 17.7 (13.7) |
| No. reflections | 90013 (4419) | 48.732 (2362) |
| **Refinement** | | |
| Resolution (Å) | 1.34 | 1.65 |
| No. reflections | 89977 | 48559 |
| Rwork / Rfree | 0.218/0.235 | 0.267/0.293 |
| Average B-factor ($Å^2$) | 23.0 | 30.54 |
| RMSD: bonds (Å) | 0.0054 | 0.0058 |
| RMSD: bond angles (°) | 0.7719 | 0.8214 |
| Ramachandran favoured (%) | 98 | 98 |
| Ramachandran outliers (%) | 0 | 0 |
| Clashscore | 1 | 1 |

| | | |
|---|---|---|
| RSRZ outliers (%) | 16.3 | 11.6 |
| Ligand RSCC | 0.94 | 0.95 |

| PDB code | 9HU8 | 9I1L |
| --- | --- | --- |
| Ligand | **MC-278** | **RS-009** |
| Beamline | DLS I03 | DLS I04-1 |
| Wavelength | 0.9763 | 0.9282 |
| Space group | P $6_5$ 2 2 | P $6_5$ 2 2 |
| a, b, c (Å) | 62.05,62.05,339.44 | 62.37,62.37,339.36 |
| a, b, g (°) | 90, 90, 120 | 90, 90, 120 |
| **Data collection** | | |
| Resolution | 56.57 – 1.48 | 56.56 – 1.68 |
| | 151 – 1.48 | 1.71 – 1.68 |
| $R_{merge}$ | 0.110 (2.207) | 0.121 (1.287) |
| Mean I/s(I) | 17.2 (0.4) | 12.0 (1.20) |
| $CC_{1/2}$ | 1.000 | 0.991 |
| Completeness (%) | 99.8 (93.4) | 100 (100) |
| Redundancy | 30.9 | 17.6 |
| No. reflections | 65984 (3016) | 46349 (2250) |
| **Refinement** | | |
| Resolution (Å) | 1.48 | 1.68 |
| No. reflections | 65303 | 46172 |
| Rwork / Rfree | 0.206/0.231 | 0.243/0.269 |
| Average B-factor (Å$^2$) | 23.0 | 31.0 |
| RMSD: bonds (Å) | 0.005 | 0.0065 |
| RMSD: bond angles (°) | 0.79 | 0.8571 |
| Ramachandran favoured (%) | 98.5 | 98 |
| Ramachandran outliers (%) | 0 | 0 |
| Clashscore | 1 | 2 |
| RSRZ outliers (%) | 6.5 | 9.8 |

| | | |
|---|---|---|
| Ligand RSCC | 0.93 | 0.96 |

#### Docking

*In silico* docking was performed using Molsoft ICM-Pro software.[288] The receptor was prepared from the structure of Kalirin-Rac1 in complex with **1** (PDB code: 5QQD). The protein was converted into an ICM object, the process of which deleted waters and optimised the orientation of Asn, Cys, Gln, His and Pro residues. The receptor pocket was defined surrounding the selected ligand. The original ligand was docked using the standard parameters in ICM to ensure the docking algorithm would produce valid poses. The Molsoft database was manually searched, and suitable compounds downloaded within an SDF file and loaded into ICM as a chemical table. The compounds clustered based on structural similarity and smaller molecular weight compounds were chosen to represent each subset. 3D stereoisomers were generated of the chosen compounds. A rigid docking protocol was performed using the standard parameters in ICM, with three conformations per compound. The compounds were ranked based on their RMSD from the original ligand and the compounds with RMSD < 9 were manually inspected and chosen for purchase.

#### Nucleotide exchange assay

Synthesised compounds were first tested at 500 µM or 250 µM final concentration using a nucleotide exchange assay. Dose-response curves were obtained using a 10-point 2-fold serial dilution with a high concentration of 500 µM. Compounds were dispensed using an ECHO acoustic dispenser in triplicate into a 384-polypropylene microplate (Grenier Bio-One), with a maximum final concentration of 2% DMSO (including control wells). The assay buffer (20 mM Tris pH 7.5, 50 mM NaCl, 1mM MgCl2, 100 µg/mL BSA and 1 mM DTT) was dispensed into sample and negative control wells (10 µL), and positive control wells added GDPNP (400 µM in assay buffer, 10 µL, Jena BioScience) to give a final concentration of 200 µM. The enzymes were mixed to a concentration of 4 µM GTPase and 0.1 µM GEF in assay buffer and dispensed into all wells (5 µL) to make a final concentration of 1 µM and 0.025 µM respectively. The plate was centrifuged at 2,000 rpm for 5 mins. After 2 h incubation at room temperature, BODIPYTM FL GTP (2 µM in assay buffer, 5 µL, ThermoFisher Scientific) was dispensed to all wells to give a final concentration of 0.5 µM. The plate was shaken for 10 seconds and the FI signal measured using the Pherastar FSX (BMG

Labtech, FI 485/520, Gain 300) every 0.5–2.0 min. The initial rate was calculated and plotted using GraphPad Prism 9, the values being normalised to the positive and negative controls. All values were plotted as normalised initial rate against each compound or concentration in the case of dose responses (FI/s). Kalirin-GEF(1) (amino acids 1273–1581) and full length Rac1 were used in the assay.

**Kalirin dose response**

The Kalirin dose response assay was performed following the general procedure described above, with the following modifications and using a 96-well plate format. MC-278 (400 μM) and Rac1 (4 μM) were prepared in assay buffer and dispensed into the appropriate wells (5 μL) in triplicate, giving final concentrations of 100 μM and 1 μM, respectively. A two-fold, six-point serial dilution of Kalirin was prepared in assay buffer with a maximum concentration of 8 μM, corresponding to a final maximum concentration of 2 μM in each well. The dilution series (5 μL) was dispensed in triplicate into the dose-response wells containing compound and GDPNP control wells (final concentration of 200 μM in assay buffer, 5 μL). Following dispensing, the plates were centrifuged at 2,000 rpm for 2 min and incubated at room temperature for 24 h before continuing with the assay as described in the general procedure.

**$k_{inact}/K_i$**

$k_{inact}/K_i$ was calculated using the equation reported by Krippendorf[17] and the FI nucleotide exchange assay. $IC_{50}$ values of both **RS-009** and **MC-278** were recorded at time periods of 10–120 min and plotted. The $k_{inact}$ and $K_i$ values were calculated in GraphPad Prism 9 in $min^{-1}$ $\mu M^{-1}$ and converted to $s^{-1}$ $M^{-1}$ prior to reporting.

**Selectivity panel**

The limited Rho selectivity panel was performed with the described BODIPY-GTP nucleotide exchange assay. Rac1, RhoA, Cdc42, Kalirin-GEF(1), Kalirin-GEF(2) and Rab39B were produced in-house; the GEF domains of Tiam1, Dbs, and Vav1 were purchased from Universal Biologicals; c9orf72 was purchased from Origene. All GTPases were used at a concentration of 1 μM, and GEFs at 25–250 nM. MC-278 was tested against each GEF/GTPase pair at 500 μM, and if activity was observed, a dose-response curve was obtained.

**GDPNP competition assay**

RapidFire experiments were performed in duplicate on an Agilent 6530 Accurate Mass QTOF and the data processed by the Agilent MassHunter Workstation software. Full length Rac1 (20 µL), Kalirin-GEF(1) (amino acids 1273–1581) (20 µL), MC-278 (20 µL) and GDPNP (20 µL) were incubated for 24 h at 4 °C in assay buffer (20 mM Tris PH 7.5, 50 mM NaCl, 1mM MgCl2) to give final concentrations of 1.25 µM, 1.25 µM, 200 µM and 0 – 200 µM respectively. The abundances of WT-Rac1 and MC-278-Rac1 by mass were calculated and used to determine the percentage of Rac1 labelling. For dose response curves, a 10-point 2-fold serial dilution was performed for **MC-278** with a high concentration of 200 µM. The measurement was repeated in the presence of GDPNP at a final concentration of 10 µM. Control wells absent of Kalirin-GEF(1) or GDPNP were displaced with buffer (20 µL).

**Kalirin dose response MS assay**

MC-278 (400 µM) and Rac1c007 (4 µM) were prepared in assay buffer and dispensed into the appropriate wells (5 µL) in 2 x triplicate in a 384-polypropylene microplate (Grenier Bio-One), giving final concentrations of 100 µM and 1 µM, respectively. A two-fold, six-point serial dilution of Kalirin-GEF(1) was then prepared with a maximum concentration of 8 µM in assay buffer (giving a final maximum Kalirin concentration of 2 µM in each well). The dilution series (5 µL) was dispensed in 2 x triplicate into the dose-response wells containing compound. Negative control wells in 2 x triplicate were prepared containing Rac1 (5 µL; final concentration 1 µM), MC-278 (5 µL; final concentration 100 µM), and assay buffer (5 µL). Plates were centrifuged at 2,000 rpm for 2 min and incubated for 5 h and 24 h at room temperature. Prior to MS analysis, samples were transferred to 384-well Greiner V-bottom plate (Greiner 781280) and quenched with Formic Acid (5 µL, 0.1% in Milli-Q® water). 20 µL of the sample was injected onto a 2.1x12.5 mm 5 µm particle size C3 reversed phase HPLC column in 0.1% formic acid, 10% methanol in water at 1.25 mL/min and eluted into the mass spectrometer (Agilent 6530 with Dual AJS ESI source) using a gradient of increasing methanol and acetonitrile concentration. Raw MS data were processed using MassHunter Workstation Software Qualitative Analysis (vB.07.00). The percentage conversion of WT Rac1 to MC-278-labelled was calculated and plotted using GraphPad Prism 11.

**Biotinylated Rac1-Kalirin isolation**

To 2 mL Eppendorf tubes, MC-278, biotinylated Rac1 (RB), and Kalirin were combined in assay buffer to give final concentrations of 200 µM, 35 µM, and 50 µM, respectively. A DMSO control was prepared in parallel with a final DMSO concentration of 0.2%. After incubation for 24 h at room temperature, additional MC-278 (200 µM) was added to give a final concentration of 500 µM. The mixtures were incubated for a total of 40 h, at which point mass spectrometry analysis confirmed quantitative labelling of RB by MC-278. The resulting MC-278-Rac1-Kalirin complex (MC-278-RB-Kalirin) and the control complex (RB-Kalirin) were purified using MagStrep® Strep-Tactin®XT beads (400 µL, IBA Lifesciences) pre-equilibrated with wash buffer (1 M Tris–HCl, 1.5 M NaCl, 10 mM EDTA, pH 8.0). The bead slurry was incubated separately with the MC-278-RB-Kalirin and RB-Kalirin mixtures for 1 h under gentle agitation. The beads were subsequently washed twice with wash buffer (2 mL) and the bound complexes were eluted twice with elution buffer (1 M Tris–HCl, 1.5 M NaCl, 10 mM EDTA, 500 mM biotin, pH 8.0; 500 µL per elution).

Size Exclusion Chromatography Purification

The first eluted fraction (E1) of MC-278-RB-Kalirin and RB-Kalirin from the Strep-Tactin®XT purification were purified by SEC in 1X PBS (SRT® SEC-100 300x7.8 mm, Sepax Technologies), alongside RB (500 µL, 106 µM) and Kalirin (500 µL, 94 µM).

**Thiol activity assay**

Thiol reactivity was assessed using a previously reported DTNB assay.[10] DTNB (50 µM) was reduced with TCEP (200 µM) in 20 mM sodium phosphate buffer (pH 7.5) containing 150 mM NaCl for 15 min at room temperature to generate $TNB^{2-}$. The reaction was initiated by addition of compound (final concentration 180–200 µM) and monitored in triplicate at 415 nm in a 96-well plate (Hidex spectrophotometer) at room temperature every 15 min for 7 h. Absorbance values were background-corrected for each compound under identical conditions and normalised prior to analysis.

Kinetic data were fitted to a second-order rate equation. The rate constant ($k$) was obtained from linear regression of $\ln([A][B_0]/[B][A_0])$, where $[A_0]$ and $[B_0]$ are the initial concentrations of compound and $TNB^{2-}$ (100 μM), respectively, and [A] and [B] are the corresponding time-dependent concentrations derived from spectrophotometric measurements. For MC-278, the initial linear region and the subsequent phase were analysed separately.

For HRMS analysis, 10 μL of reaction mixture was quenched with formic acid (10 μL, 0.1% in Milli-Q water) and transferred to a 384-well Greiner V-bottom plate. Samples (10 μL) were analysed by HRMS using a C18 column on an instrument configured for small-molecule detection. Raw data were processed using MassHunter Workstation Qualitative Analysis (vB.07.00).

### Synthetic procedures

#### General chemistry experimental

##### Solvents and reagents

Water was deionised by an Elga DV 25 system. All other solvents and reagents were purchased from commercial sources and used as supplied (analytical or HPLC grade) without prior purification. All reactions involving moisture sensitive reagents were carried out under a nitrogen atmosphere. Standard practices were employed when handling moisture- and air-sensitive compounds. Room temperature (r.t.) refers to ambient temperature. Temperatures of 0 °C were maintained using an ice-water bath.

##### Purification and chromatography

Reactions were monitored by thin layer chromatography (TLC) and/or liquid chromatography-mass spectrometry (LCMS). TLC analysis was performed on commercially prepared aluminium plates coated with 60 F254 silica gel. Plates were visualised using UV light (254 nm). Analytical LCMS was performed on the following system: Kinetex 5µ EVO C18 100A 100 x 3.0 mm column using a linear gradient of 5-95% of solvent A (93 % WATER, 5 % MeCN, 2 % of 0.5 M NH4OAc pH 6.0) in solvent B (18% WATER, 80% MeCN, 2% of 0.5 M NH4OAc pH 6.0), eluting at a flow rate of 2 mL/min. Compound formation and/or purity was assessed by either UV absorbance at 254 nm (Waters UV/Visible Detector 2489), ELSD signal (Waters ELS Detector 2424) or ESI+ TIC (SQ Detector 2). Flash column chromatography was performed on a Biotage Isolera One flash column chromatography platform. Pre-packed SNAP-KP-Sil or SNAP-Ultra columns were used unless otherwise stated (Biotage). Purification by HPLC was performed using a Waters SFO with 515 HPLC pump and Waters Binary Gradient 2545 device (5-95% solvent A [18% Water, 80% MeCN, 2% NH4OAc pH 6.0] in solvent B [93% Water, 5% MeCN, 2% NH4OAc pH 6.0]). Analytical pH 8 method: column: Kinetex 5 µM EVO C18 column (100 mm × 3.0 mm, 100 Å)eluent A: 88% water, 10% acetonitrile, and 2% of 0.5 M ammonium bicarbonate, eluent B: 18%water, 80% acetonitrile, and 2% of 0.5 M ammonium bicarbonate; gradient: 0-0.35 min 5% B, 0.35-1.35 min 5-95% B, 1.35-2.1 min 95% B, 2.1-2.2 min 95-5% B, 2.2-3 min 5% B; flowrate 2 ml/min; wavelength: 220 nm to 650 nm. Preparative pH 8 method: column: Kinetex 5 µM EVO C18 column (150 mm × 21.2 mm, 100 Å); eluent A: 88% water, 10% acetonitrile, and 2% of 0.5 M ammonium bicarbonate, eluent B: 18% water, 80% acetonitrile, and 2% of 0.5 M ammonium bicarbonate; gradient: 0-2.5 min 15% B, 2.5-5 min 15-90% B, 5-18 min 90% B, 18-18.5 min 90-15% B, 18.5-20 min 15% B; flow 20 ml/min; wavelength: 220 nm to 650 nm.

**Characterisation**

Melting points (Mpt) were recorded using a Stuart SMP40 apparatus and are reported in °C. Infrared (IR) spectra were recorded neat on a Thermo Scientific Nicolet iS5 with an iD7 ATR module. Selected absorption maxima (nmax) are reported in wavenumbers (cm-1). NMR spectra were recorded using a Bruker Avance 400 MHz spectrometer using the deuterated solvent stated. All NMRs were conducted at 298 K unless otherwise stated. The field was locked by external referencing to the relevant deuteron resonance. Chemical shifts (d) are quoted in parts per million (ppm) and referenced to the residual solvent peak. Proton shifts are quoted to the nearest 0.01 ppm and carbon shifts to the nearest 0.1 ppm. When peak multiplicities are reported, the following abbreviations are used: br = broadened, s = singlet, d = doublet, t = triplet, q = quartet, m = multiplet, dd = doublet of doublets, dt = doublet of triplets, dq = doublet of quartets, ddd = doublet of doublet of doublets, tt = triplet of triplets and qd = quartet of doublets. Coupling constants (J) are quoted in Hz and rounded to the nearest 0.1 Hz. Assignments of protons and carbons were supported by two-dimensional NMR experiments (COSY, HSQC, HMBC) or by analogy. Low resolution mass spectra were recorded by LCMS on a Water SQ Detector 2; data acquisition and processing were performed using Waters FractionLynx software. High Resolution Mass Spectrometry (HRMS) was performed on an Agilent 6530 Accurate Mass QTOF;

data acquisition and processing were performed using Agilent MassHunter Workstation software. Compound names were generated using ChemDraw v20.1 systematic naming. Atom numbering in structures is purely for the purposes of assignment and does not reflect IUPAC numbering conventions.

**RS-009**

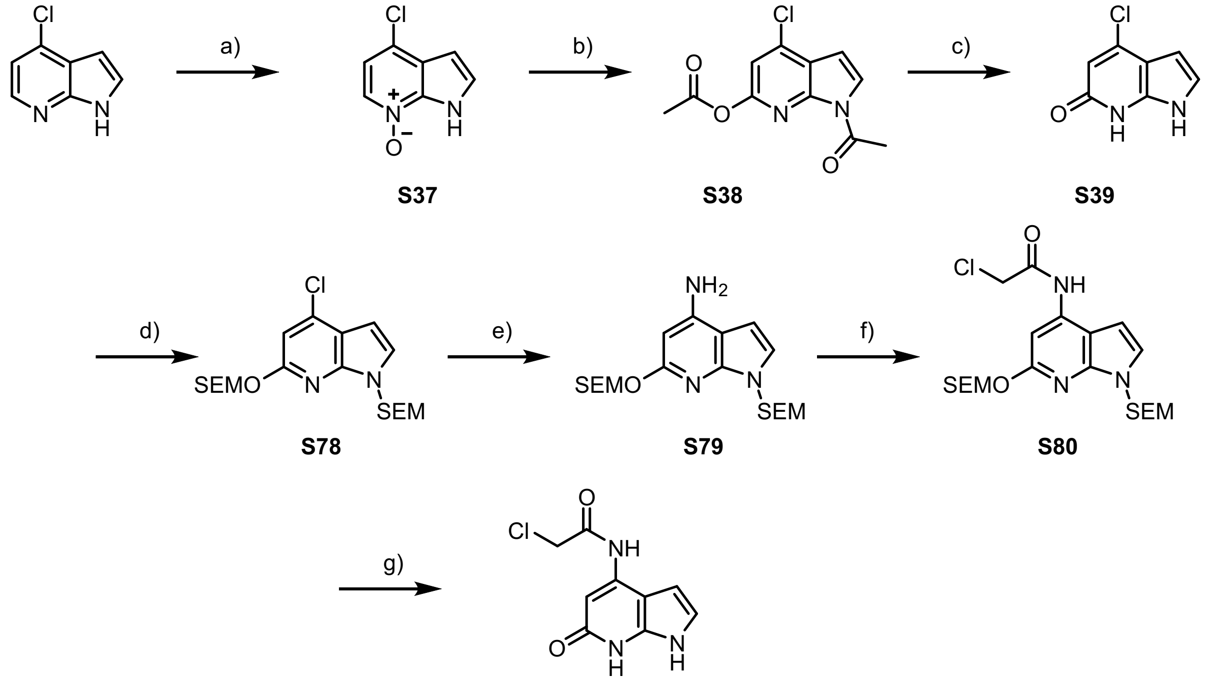

**RS-009**

#### 4-chloro-1H-pyrrolo[2,3-b]pyridine 7-oxide, S37

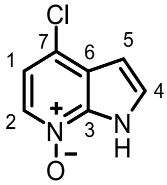

(a) To a solution of 4-chloro-1H-pyrrolo[2,3-b]pyridine (3.00 g, 19.7 mmol, 1.00 eq) in ethyl acetate (150 mL) was slowly added mCPBA (5.09 g, 29.5 mmol, 1.50 eq).

The resulting mixture was reacted at room temperature for 2 hours, filtered and dried overnight in a vacuum oven to afford **S37** (2.98 g, 17.7 mmol, 90%) as a pale pink solid.

**$^1$H NMR (400 MHz, CDCl$_3$) δ$_H$** 8.14 (d, *J* = 6.6 Hz, 1H, C(2)*H*), 7.44 (d, *J* = 3.4 Hz, 1H, C(4)*H*), 7.08 (d, *J* = 6.7 Hz, 1H, C(1)*H*), 6.63 (d, *J* = 3.4 Hz, 1H, C(9)*H*); **$^{13}$C NMR (101 MHz, CDCl$_3$) δ$_C$** 139.0 *C*(3), 132.0 *C*(2), 129.0 *C*(6), 128.0 *C*(4), 124.1 *C*(7), 115.9 *C*(1), 101.7 *C*(5); **LCMS** found [M+H]$^+$ 169.1, Rf = 1.07 min; **HRMS** (ESI$^+$) $C_7H_6ClN_2O^+$ ([M+H]$^+$) requires 169.0163; found 169.0155.

#### 1-acetyl-4-chloro-1H-pyrrolo[2,3-b]pyridin-6-yl acetate, S38

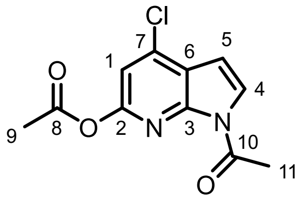

(b) **S37** (3.00 g, 17.8 mmol, 1.00 eq) was dissolved in acetic anhydride (30 mL) and refluxed at 150 °C overnight. The reaction mixture was extracted 3 times with DCM and washed 3 times with water dried over anhydrous sodium sulfate and concentrated under reduced pressure.

The product was purified by flash chromatography on silica gel (100 g, 2-15% ethyl acetate in cyclohexane) to afford **S38** (2.23 g, 8.82 mmol, 50%) as a white solid.

**$^1$H NMR (400 MHz, $CDCl_3$) $\delta_H$** 7.99 (d, *J* = 4.1 Hz, 1H, C(4)*H*), 7.01 (s, 1H, C(1)*H*), 6.71 (d, *J* = 4.1 Hz, 1H, C(5)*H*), 2.97 (s, 3H, C(11)$H_3$), 2.37 (s, 3H, C(9)$H_3$); **$^{13}$C NMR (101 MHz, $CDCl_3$) $\delta_C$** 169.1 *C*(8), 168.8 *C*(10), 153.5 *C*(2), 145.5 *C*(3), 138.7 *C*(7), 126.1 *C*(4), 121.6 *C*(6), 111.6 *C*(1), 103.8 *C*(5), 25.9 *C*(11), 21.3 *C*(9); **LCMS** found $[M\text{-}CH_3CO+H]^+$ 211.1, Rf = 1.63 min; **HRMS** ($ESI^+$) $C_{11}H_{10}ClN_2O_3^+$ ($[M+H]^+$) requires 253.0374; found 253.0376.

**4-chloro-1H-pyrrolo[2,3-b]pyridin-6-ol, S39**

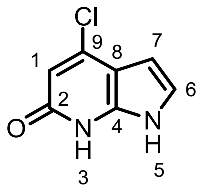

(c) **S38** (6.47 g, 25.6 mmol, 1.00 eq) was dissolved in degassed MeOH (129 mL). To the reaction flask was added a degassed solution of 3M aqueous $K_2CO_3$ (25.6 mL, 76.8 mmol, 3.00 eq). The reaction was stirred at room temperature for 1 hour.

The reaction mixture was quenched with 1 M aqueous HCl to afford a purple solid. The solid was collected by filtration, washed with MeOH and water and dried overnight in a vacuum oven to afford **S39** (3.01 g, 19.9 mmol, 70%) as a purple solid.

**$^1$H NMR (400 MHz, DMSO) $\delta_H$** 11.48 (s, 1H, N(*H*)), 10.86 (s, 1H, N(*H*)), 7.17 (d, *J* = 3.5 Hz, 1H, C(6)*H*), 6.43 (s, 1H, C(1)*H*), 6.31 (d, *J* = 3.5 Hz, 1H, C(7)*H*) **$^{13}$C NMR (101 MHz, DMSO) $\delta_C$** 160.2 *C*(2), 145.5 *C*(4), 136.7 *C*(9), 122.5 *C*(6), 111.6 *C*(8),

102.9 *C*(1), 98.4 *C*(7); **LCMS** found [M+H]$^+$ 169.1, Rf = 1.15 min; **HRMS** (ESI$^+$) $C_7H_6ClN_2O^+$ ([M+H]$^+$) requires 169.0163; found 169.0162.

**4-chloro-6-((2-(trimethylsilyl)ethoxy)methoxy)-1-((2-(trimethylsilyl)ethoxy)methyl)-1H-pyrrolo[2,3-b]pyridine, S78**

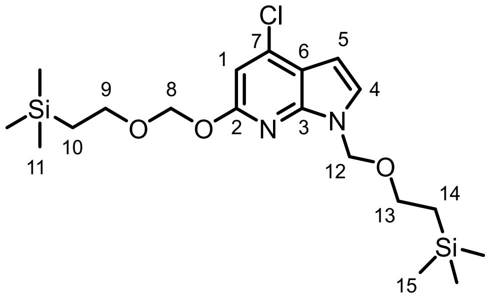

(d) **S39** (1.24 g, 7.37 mmol, 1.00 eq) was added to a round bottom flask and evacuated and back-filled 3 times with nitrogen, dissolved in DMF (50 mL) and cooled to 0 °C.

To the reaction was added a freshly prepared solution of 2M sodium bis(trimethylsilyl)amide in THF (11.1 mL, 22.1 mmol, 3.00 eq) dropwise by syringe pump (1 mL/min), followed by the dropwise addition (syringe pump 0.5 mL/min) of 2-(chloromethoxy)ethyl]trimethylsilane (3.90 mL, 22.1 mmol, 3.00 eq) at 0 °C. The reaction was stirred at room temperature for one hour.

The reaction was quenched with water and extracted 3 times with TBME. The combined organic layers were washed 3 times with water and 2 times with 1M aqueous LiCl solution. The organic layers were dried over anhydrous sodium sulfate and concentrated under reduced pressure.

The product was purified by flash chromatography on silica gel (50 g, 0-2.5% acetone in cyclohexane) to afford impure product as a pale purple oil. The product was purified again by flash chromatography on silica gel (25 g, 0-5% ethyl acetate in cyclohexane) to afford **S78** (1.32 g, 3.07 mmol, 42%) as a colourless oil.

**$^1$H NMR (400 MHz, CDCl$_3$) $\delta_H$** 7.16 (d, *J* = 3.6 Hz, 1H, C(4)*H*), 6.70 (s, 1H, C(1)*H*), 6.52 (d, *J* = 3.6 Hz, 1H, C(5)*H*), 5.59 (s, 2H, C(8)*H$_2$*), 5.55 (s, 2H, C(12)*H$_2$*), 3.85 – 3.76 (m, 2H, C(9)*H$_2$*), 3.58 – 3.48 (m, 2H, C(13)*H$_2$*), 1.03 – 0.95 (m, 2H, C(10)*H$_2$*), 0.92 – 0.86 (m, 2H, C(14)*H$_2$*), 0.01 (s, 9H, C(11)*H$_3$*), -0.06 (s, 9H, C(15)*H$_3$*); **$^{13}$C NMR (101 MHz, CDCl$_3$) $\delta_C$** 159.6 *C*(2), 146.0 *C*(3), 138.3 *C*(7), 125.7 *C*(4), 114.7 *C*(6),

104.7 *C*(1), 100.1 *C*(5), 91.1 *C*(8), 73.2 *C*(12), 67.4 *C*(9), 66.5 *C*(13), 18.3 *C*(10), 17.9 *C*(14), -1.3 *C*(11,15); **LCMS** found $[M+Na]^+$ 451.4, Rf = 1.88 min; **HRMS** ($ESI^+$) $C_{19}H_{34}ClN_2O_3Si_2^+$ ($[M+H]^+$) requires 429.1791; found 429.1791.

**6-((2-(trimethylsilyl)ethoxy)methoxy)-1-((2-(trimethylsilyl)ethoxy)methyl)-1H-pyrrolo[2,3-b]pyridin-4-amine, S79**

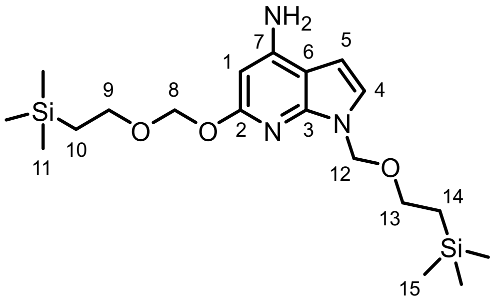

(e) A 2-necked round bottom flask was charged with tri-*tert*-butylphosphonium tetrafluoroborate (110 mg, 0.378 mmol, 0.30 eq) and tris(dibenzylideneacetone)dipalladium(0)) (87.0 mg, 0.094 mmol, 0.075 eq) and evacuated and back-filled with nitrogen three times.

A round bottom flask was dried under vacuum and evacuated and backfilled with nitrogen 3 times. In this flask toluene (15 mL) was degassed.

A round bottom flask containing **S78** (540 mg, 1.26 mmol, 1.00 eq) was evacuated and back-filled with nitrogen three times dissolved in degassed toluene (13 mL) and added to the reaction flask.

To the reaction mixture was added a 1M solution of lithium bis(trimethylsilyl)amide in THF (2.50 mL, 2.52 mmol, 2.00 eq). The reaction was stirred at 100 °C overnight.

The reaction was cooled to room temperature, diluted with TBME (10 mL) and aqueous 1 M HCl (10 mL) was added and the reaction was stirred at room temperature for 45 minutes. To the reaction mixture was added saturated aqueous $NaHCO_3$ solution and the reaction mixture was filtered over celite. The organic layer was taken and the aqueous layer was extracted twice more with TBME and the combined organic layers were washed twice with water and twice with brine and dried over anhydrous sodium sulfate and concentrated under reduced pressure.

The product was purified by flash chromatography on silica gel (25 g, 0-20% acetone in cyclohexane) to afford **S79** (290 mg, 0.566 mmol, 45%) as a dark brown oil.

**$^1$H NMR (400 MHz, CDCl$_3$) δ$_H$** 6.99 (d, *J* = 3.6 Hz, 1H, C(4)*H*), 6.32 (d, *J* = 3.7 Hz, 1H, C(5)*H*), 5.90 (s, 1H, C(1)*H*), 5.55 (s, 2H, C(8)*H$_2$*), 5.52 (s, 2H, C(12)*H$_2$*), 4.33 (broad s, 2H, (N)*H$_2$*), 3.83 – 3.76 (m, 2H, C(9)*H$_2$*), 3.57 – 3.50 (m, 2H, C(13)*H$_2$*), 1.01 – 0.96 (m, 2H, C(10)*H$_2$*), 0.92 – 0.87 (m, 2H, C(14)*H$_2$*), -0.00 (s, 9H, C(11)*H$_3$*), -0.07 (s, 9H, C(15)*H$_3$*); **$^{13}$C NMR (101 MHz, CDCl$_3$) δ$_C$** 161.3 *C*(2), 149.0 *C*(7), 146.9 *C*(3), 122.8 *C*(4), 104.5 *C*(6), 97.8 *C*(5), 90.7 *C*(8), 87.7 *C*(1), 72.9 *C*(12), 67.0 *C*(9), 66.2 *C*(13), 18.3 *C*(10), 17.9 *C*(14), -1.3 *C*(11,15); **LCMS** found $[M-H]^-$ 408.4, Rf = 2.37 min; **HRMS** (ESI$^+$) $C_{19}H_{36}N_3O_3Si_2^+$ ($[M+H]^+$) requires 410.2290; found 410.2291**.**

**2-chloro-N-(6-((2-(trimethylsilyl)ethoxy)methoxy)-1-((2-(trimethylsilyl)ethoxy)methyl)-1H-pyrrolo[2,3-b]pyridin-4-yl)acetamide, S80**

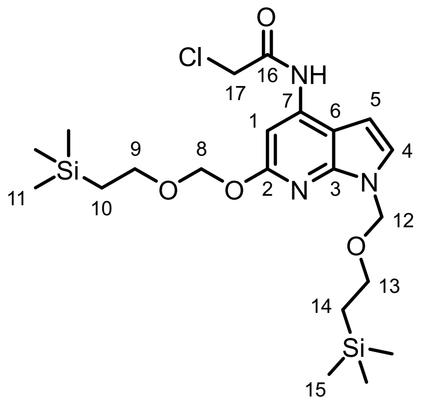

(f) A solution of **S79** (100 mg, 0.244 mmol, 1.00 eq) in DCM (2.44 mL) was cooled to 0 °C. Triethylamine (68.0 µL, 0.488 mmol, 2.00 eq) followed by 2-chloroacetyl chloride (29.2 µL, 0.366 mmol, 1.50 eq) was added at 0 °C. The reaction was stirred at room temperature for 30 minutes.

The reaction was quenched with water and extracted with DCM through a phase separator. The product was purified by flash chromatography on silica gel (5 g HC, 12 mL/min flow rate 2-20% acetone in cyclohexane) to afford **S80** (61 mg, 0.125 mmol, 51%) as a brown oil.

**$^1$H NMR (400 MHz, CDCl$_3$) δ$_H$** 8.63 (broad s, 1H, N(*H*)), 7.51 (s, 1H, C(1)*H*), 7.13 (d, *J* = 3.7 Hz, 1H, C(4)*H*), 6.40 (d, *J* = 3.7 Hz, 1H, C(5)*H*), 5.61 (s, 2H, C(8)*H$_2$*), 5.56 (s, 2H, C(12)*H$_2$*), 4.25 (s, 2H, C(17)*H$_2$*), 3.86 – 3.75 (m, 2H, C(9)*H$_2$*), 3.58 – 3.48 (m, 2H, C(13)*H$_2$*), 1.04 – 0.94 (m, 2H, C(10)*H$_2$*), 0.93 – 0.84 (m, 2H, C(14)*H$_2$*), 0.00 (s, 9H, C(11)*H$_3$*), -0.07 (s, 9H, C(15)*H$_3$*); **$^{13}$C NMR (101 MHz, CDCl$_3$) δ$_C$** 164.1 *C*(16), 160.7 *C*(2), 146.4 *C*(3), 138.7 *C*(7), 124.6 *C*(4), 106.1 *C*(6), 97.0 *C*(5), 94.6 *C*(1), 90.8 *C*(8),

73.0 *C*(12), 67.3 *C*(9), 66.4 *C*(13), 43.2 *C*(17), 18.3 *C*(10), 17.9 *C*(14), -1.3 *C*(11,15 overlapped); **LCMS** found $[M+H]^+$ 486.3, Rf = 2.60 min; **HRMS** ($ESI^+$) $C_{21}H_{37}ClN_3O_4Si_2^+$ ($[M+H]^+$) requires 486.2006; found 486.2115.

**2-chloro-N-(6-oxo-6,7-dihydro-1H-pyrrolo[2,3-b]pyridin-4-yl)acetamide, RS-009**

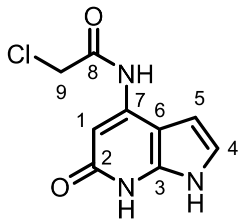

(g) **S80** (70.0 mg, 0.144 mmol, 1.00 eq) was dissolved in DCM (1.46 mL) and anisole (234.7 µL, 2.16 mmol, 15.0 eq) followed by TFA (554.7 µL, 7.20 mmol, 50.0 eq) was added and the reaction was stirred at 40 °C for 1 hour.

The reaction mixture was co-evaporated 4 times with DCM and concentrated under reduced pressure. The crude was dissolved in methanol (1.09 mL) and aqueous 1M $Na_2CO_3$ (360.0 µL, 0.360 mmol, 2.50 eq) was added and the reaction was stirred at room temperature for 30 minutes.

The reaction mixture was filtered and purified by reverse phase prep HPLC (pH 8 95-85% buffer A in buffer B) to afford **RS-009** (6.2 mg, 0.027 mmol, 19%) as a white solid. **$^1$H NMR (400 MHz, DMSO) $\delta_H$** 11.13 (broad s, 1H, N(*H*)), 10.14 (s, 1H, N(*H*)), 7.13 (s, 1H, C(1)*H*), 6.99 (dd, *J* = 3.5, 2.3 Hz, 1H, C(4)*H*), 6.60 (dd, *J* = 3.5, 2.0 Hz, 1H, C(5)*H*), 4.42 (s, 2H, C(9)$H_2$); **$^{13}$C NMR (101 MHz, DMSO) $\delta_C$** 165.8 *C*(8), 161.0 *C*(2), 145.4 *C*(3), 140.2 *C*(7), 119.8 *C*(4), 103.8 *C*(6), 98.4 *C*(5), 92.9 *C*(1), 43.5 *C*(9); **LCMS** found $[M+H]^+$ 226.1, Rf = 1.00 min; **HRMS** ($ESI^+$) $C_9H_9ClN_3O_2^+$ ($[M+H]^+$) requires 226.0378; found 226.0333.

**MC-278**

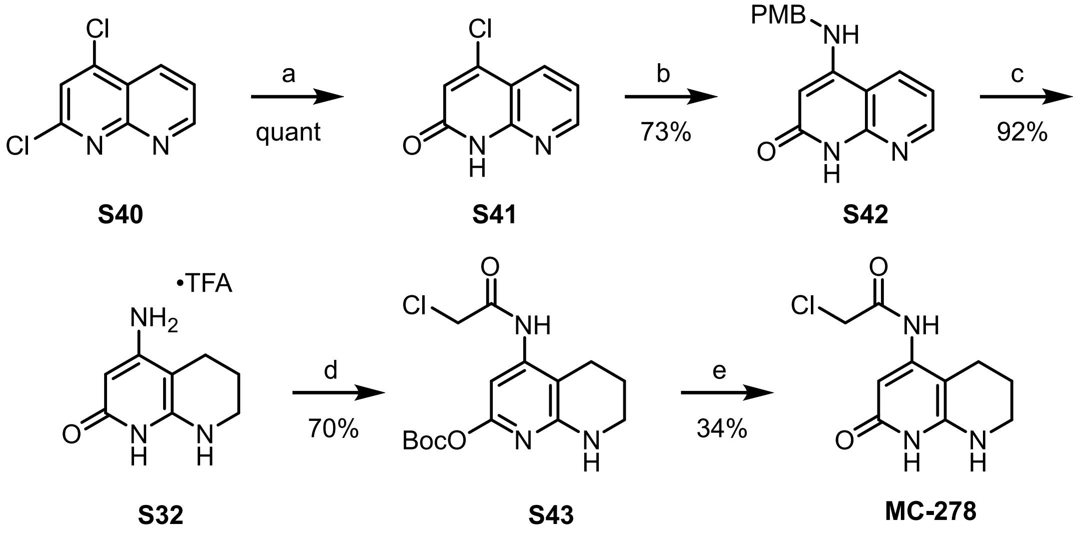

**4-chloro-1,8-naphthyridin-2-ol, S41**

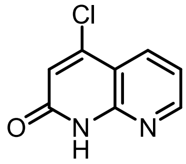

(a) To a solution of 2,4-dichloroquinoline, **S40** (500 mg, 2.51 mmol) in dioxane (5 mL) was added HCl (6N, 7.5 mL) dropwise at 0 °C. The mixture stirred at reflux for 17 h, cooled and water added. The resulting precipitate was filtered and dried under vacuum to give **S41** as a white solid (455 mg, quant).

**$^{1}$H NMR** (400 MHz, DMSO-$d_6$) δ 12.45 (s, 1H), 8.63 (dd, $J$ = 4.7, 1.7 Hz, 1H), 8.26 (dd, $J$ = 8.0, 1.7 Hz, 1H), 7.38 (dd, $J$ = 8.0, 4.7 Hz, 1H), 6.90 (s, 1H); **$^{13}$C NMR** (101 MHz, DMSO-$d_6$) δ 161.6, 152.5, 149.7, 143.4, 134.3, 122.8, 119.4, 113.4; **LR-ESI-MS** $C_8H_5ClN_2O$ [M+H]$^+$ $m/z$ found 181.2 cald 181.1; **HR-ESI-MS** $C_8H_5ClN_2O$ [M+H]$^+$ $m/z$ found 181.0161 cald 181.0169.

**4-((4-methoxybenzyl)amino)-1,8-naphthyridin-2(1H)-one, S42**

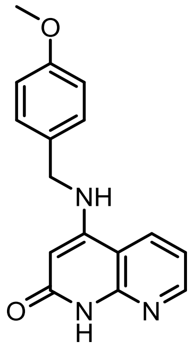

(b) **S41** (1.04 g, 5.6 mmol) was added 4-methoxybenzylamine (12.3 mL) and DMSO (26.1 mL) and stirred at 150 °C for 1 h. The reaction was cooled, water added, and the precipitate filtered. The resulting solid was washed with DCM/MeOH to give **S42** as a white solid (1.2 g, 73%).

**$^1$H NMR** (400 MHz, DMSO-*d*$_6$) δ 11.07 (s, 1H), 8.49–8.40 (m, 2H), 7.71 (t, *J* = 5.9 Hz, 1H), 7.33–7.25 (m, 2H), 7.20 (dd, *J* = 8.0, 4.7 Hz, 1H), 6.95–6.87 (m, 2H), 5.19 (d, *J* = 1.6 Hz, 1H), 4.37 (d, *J* = 5.7 Hz, 2H), 3.73 (s, 3H). **$^{13}$C NMR** (101 MHz, DMSO-*d*$_6$) δ 164.4, 159.2, 151.2, 150.9, 131.9, 131.0, 129.1, 117.8, 114.8, 110.1, 92.7, 55.9, 46.1; **LR-ESI-MS** $C_{16}H_{15}N_3O_2$ $[M+H]^+$ *m/z* found 282.4 cald 282.1; **HR-ESI-MS** $C_{16}H_{15}N_3O_2$ [M+H]+ m/z found 282.1245 calc. 282.1237.

**4-amino-5,6,7,8-tetrahydro-1,8-naphthyridin-2(1H)-one, S32**

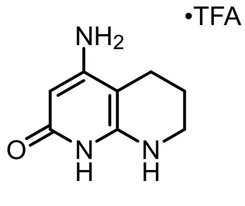

(c) **S42** (991 mg, 3.52 mmol) was dissolved in formic acid (50 mL) and MeOH (20 mL) and the mixture passed through a Pd/C CatCart (40 °C, 20 bar, 0.5 mL/min, H-cube midi). The resulting mixture was evaporated to dryness, TFA (12 mL) added, and stirred for 13 h at 60 °C. The resulting mixture was added DCM, washed with water, and the aqueous evaporated to dryness to give **S32** as an orange solid (900 mg, 92%).

**$^1$H NMR** (400 MHz, DMSO-*d*$_6$) 11.92 (s, 1H), 6.80 (s, 2H), 6.70 (s, 1H), 5.45 (s, 1H), 3.23 (t, *J* = 5.5 Hz, 2H), 2.27 (t, *J* = 6.3 Hz, 2H), 1.81–1.72 (m, 2H); **$^{19}$F NMR** (376 MHz, DMSO) δ -74.0; **$^{13}$C NMR** (101 MHz, DMSO-*d*$_6$) δ 158.5, 155.6, 148.0, 88.5, 81.9, 40.1*, 20.3, 19.5. **LR-ESI-MS** $C_8H_{11}N_3O$ $[M+H]^+$ *m/z* found 166.3 cald 166.1; **HR-ESI-MS** $C_8H_{11}N_3O$ $[M+H]^+$ *m/z* found 166.0981 cald.166.0975. **chemical shift determined from 2D NMR data.*

***tert*-butyl (4-(2-chloroacetamido)-5,6,7,8-tetrahydro-1,8-naphthyridin-2-yl) carbonate, S43**

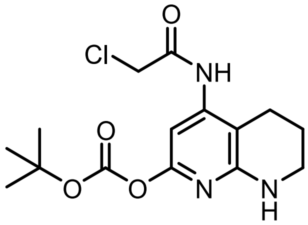

(d) **S32** (150 mg, 0.54 mmol) was added DMF (2.1 mL), TEA (300 µL, 2.15 mmol), and $Boc_2O$ (117 mg, 0.54 mmol) and stirred at r.t. for 16 h. The reaction was cooled to 0 °C and chloroacetyl chloride (43 µL, 0.54 mmol) in DMF (0.6 mL) was added dropwise. The reaction was warmed to rt, partitioned between DCM and water, and the organic extracted with water (3x). The organic was washed with brine, filtered through a phase separator and the solvent evaporated. The crude was purified by preparative HPLC to yield **S43** as an off-white solid (128 mg, 70%).

**$^1$H NMR** (400 MHz, DMSO-*d*$_6$) δ 9.58 (s, 1H), 6.78 (d, *J* = 2.6 Hz, 1H), 6.59 (s, 1H), 4.34 (s, 2H), 3.26–3.18 (m, 2H), 2.52 (d, *J* = 8.7 Hz, 2H), 1.76 (d, *J* = 5.9 Hz, 2H), 1.46 (s, 9H). **$^{13}$C NMR** (101 MHz, DMSO-*d*$_6$) δ 165.7, 156.2, 155.2, 151.3, 145.2, 104.1, 96.2, 83.3, 43.8, 40.4*, 27.7, 22.0, 20.6. **LR-ESI-MS** $C_8H_{11}N_3O$ $[M+H]^+$ *m/z* found 342.5/344.4 cald 342.1/344.1; **HR-ESI-MS** $C_{15}H_{20}ClN_3O_4$ $[M+H]^+$ found 342.1301 cald. 342.1215. **chemical shift determined from 2D NMR data.*

**2-chloro-N-(2-oxo-1,2,5,6,7,8-hexahydro-1,8-naphthyridin-4-yl)acetamide, MC-278**

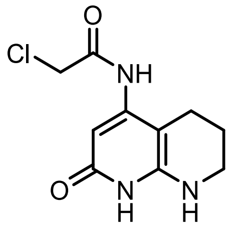

(e) **S43** (128 mg, 0.37 mmol) was added DCM (1.9 mL) and 1-methylpiperazine (50 μL, 0.45 mmol) and stirred for 2 h. The DCM was blown off with nitrogen and the residue purified by preparative HPLC to give **MC-278** as an off-white waxy solid (30.3 mg, 34%).

**$^1$H NMR** (400 MHz, DMSO-d$_6$) δ 10.31 (s, 1H), 9.19 (s, 1H), 5.97 (s, 1H), 5.88 (t, J = 2.7 Hz, 1H), 4.32 (s, 2H), 3.19 (dq, J = 6.0, 2.8 Hz, 2H), 2.36 (t, J = 6.4 Hz, 2H), 1.72 (p, J = 6.1 Hz, 2H). **$^{13}$C NMR** (101 MHz, DMSO-*d*$_6$) δ 165.5, 161.8, 149.6, 147.7, 94.0, 89.1, 43.9, 40.1*, 21.3, 20.6; **LR-ESI-MS** $C_{10}H_{12}ClN_3O_2$ $[M+H]^+$ *m/z* found 242.4/244.4 cald 242.1/244.1; **HR-ESI-MS** $C_{10}H_{12}ClN_3O_2$ $[M+H]^+$ *m/z* found 242.0695 cald 242.0696. **chemical shift determined from 2D NMR data.*

**Pyrazolopyridone 8**

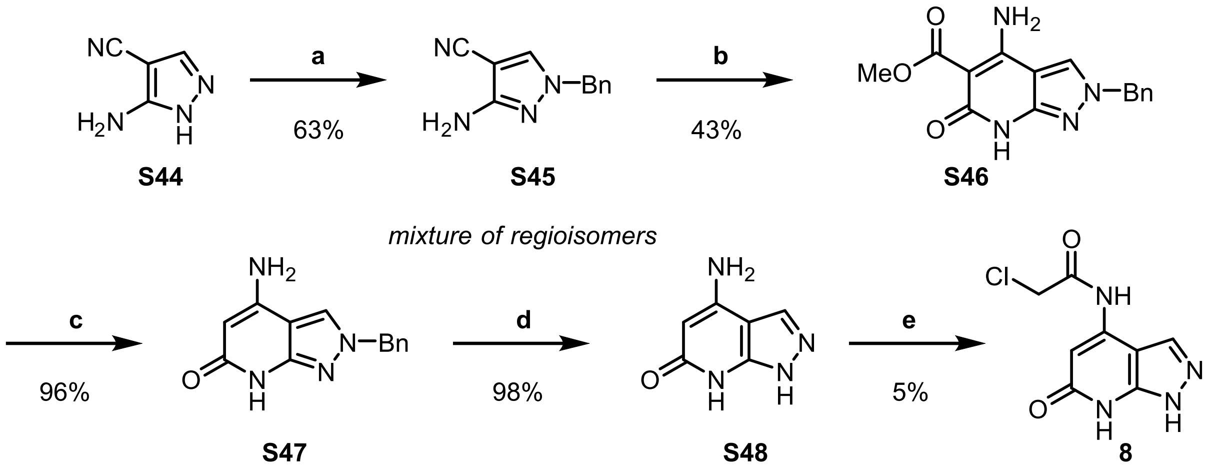

**5-amino-1-benzyl-1H-pyrazole-4-carbonitrile, S45**

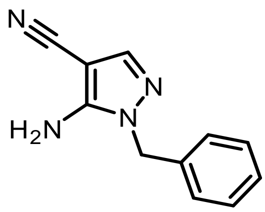

(**a**) 5-amino-1*H*-pyrazole-4-carbonitrile **S44** (500 mg, 4.5 mmol), $K_2CO_3$ (961 mg, 7.0 mmol), and anhydrous DMF (5 mL) were charged into a $N_2$ flushed flask, followed by the dropwise addition of BnBr (650 µL, 5.44 mmol). The reaction was stirred at room temperature for 22 h. The resulting mixture was diluted in water, extracted with EtOAc, and the organic layer washed with brine. The organic was dried ($Na_2SO_4$) and evaporated to dryness, and the final product was purified by flash column chromatography (0-50% EtOAc in cyclohexane) to give **S45** as a white solid and a mixture of regioisomers (2*N*-benzyl : 1*N*-benzyl 3:1) (568 mg, 63%).

**$^1$H NMR** (400 MHz, DMSO- *d*$_6$) 7.59 (s, 1H), 7.39 – 7.13 (m, 5H), 6.72 (s, 2H), 5.16 (s, 2H); **$^{13}$C NMR** (101 MHz, DMSO- *d*$_6$) δ 50.4, 72.7, 115.7, 127.9, 128.3, 128.9, 137.2, 141.0, 152.1; *N2-Bn* δ **$^1$H NMR** (400 MHz, DMSO- *d*$_6$) 8.24 (s, 1H), 7.39 – 7.13 (m, 5H), 5.57 (s, 2H), 5.08 (s, 2H); **$^{13}$C NMR** (101 MHz, DMSO- *d*$_6$) δ 55.3, 77.2, 115.1, 127.7, 128.2, 130.0, 136.3, 137.0, 158.3; **LCMS** found $[M+H]^+$ 199.3, Rf 1.32 min; **HR-ESI-MS** $C_{11}H_{10}N_4$ $[M+Na]^+$ *m/z* 221.0811 cald 221.0804.

**methyl 4-amino-2-benzyl-6-oxo-6,7-dihydro-1H-pyrazolo[3,4-b]pyridine-5-carboxylate, S46**

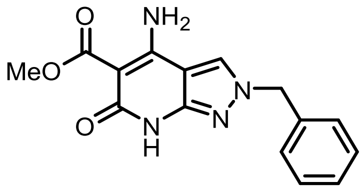

(**b**) To a nitrogen filled flask was charged MeOH (50.0 mL), NaOMe (25 wt. % in MeOH, 19.9 mL, 74.5 mmol) and dimethyl malonate (8.1 mL, 69.4 mmol) and stirred at room temperature for 15 mins. **S45** (11.00 g, 55.5 mmol) was charged and stirred at reflux for 2 days. The reaction was cooled, solid filtered, and the filtrate evaporated. To the filtrate was added MeOH and the solid filtered. The filtered solids were combined to give **S46** as a white solid that was used without further purification (7.12 g, 43%).

**$^{1}$H NMR** (400 MHz, DMSO-$d_6$) δ 10.73 (s, 1H), 8.35 (s, 1H), 8.28 (s, 2H), 7.43 – 7.21 (m, 5H), 5.34 (s, 2H), 3.66 (s, 3H); **$^{13}$C NMR** (101 MHz, DMSO-$d_6$) δ 170.0, 162.4, 155.6, 149.6, 136.9, 129.1, 128.5, 128.5, 128.2, 100.4, 89.9, 56.0, 51.0; **LCMS** found $[M+H]^+$ 299.3, Rf 1.21 min; **HR-ESI-MS** $C_{16}H_{15}N_3O_3$ found $[M+Na]^+$ 321.0078 cald 321.0964

**4-amino-2-benzyl-2,7-dihydro-6H-pyrazolo[3,4-b]pyridin-6-one, S47**

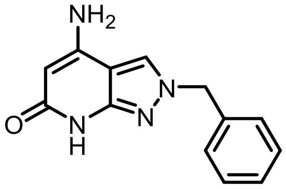

(**c**) **S46** (500 mg, 1.68 mmol) and 15% wt. NaOH solution (10.0 mL) were charged and stirred at reflux for 18 h. The reaction was cooled, acidified to pH 5, and the resulting solid was filtered and dried on high vacuum to give **S47** as an off-white that was used without further purification (385 mg, 96%).

**$^{1}$H NMR** (400 MHz, DMSO-$d_6$) δ 10.53 (s, 1H), 8.04 (s, 1H), 7.41 – 7.24 (m, 5H), 6.42 (s, 2H), 5.32 (s, 2H), 4.96 (s, 1H). **$^{13}$C NMR** (101 MHz, DMSO-$d_6$) δ 165.5, 150.7, 150.4, 137.4, 129.1, 128.6, 128.3, 125.4, 101.6, 89.1, 55.8; **LCMS** found $[M+H]^+$ 241.3, Rf 1.28 min; **HR-ESI-MS** $C_{13}H_{12}N_4O$ found $[M+H]^+$ 241.0556 cald 241.1090.

**4-amino-1,7-dihydro-6H-pyrazolo[3,4-b]pyridin-6-one, S48**

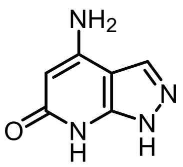

(**d**) **S47** (1.00 g, 4.15 mmol) and Pd/C (50% wetted, 800 mg) were charged and the flask was evacuated and filled with nitrogen three times. MeOH (100 mL) and 4M HCl (in dioxane, 20 mL) were then added consecutively by needle under nitrogen. The flask was evacuated and filled with nitrogen three times. The flask was evacuated and the $H_2$ balloon inserted and tap opened. The flask was then evacuated and back filled with $H_2$ twice before finally leaving the flask under $H_2$ for 3 days at room temperature.

The reaction was filtered through celite and evaporated to dryness to give the crude product **S48** as a green solid that was used without further purification (611 mg, 98%).

**$^1$H NMR** (400 MHz, DMSO-$d_6$) δ 8.09 (s, 1H), 6.57 (s, 2H), 4.98 (s, 1H); **$^{13}$C NMR** (101 MHz, DMSO-$d_6$) δ 162.7, 157.0, 147.5*, 127.4*, 102.1, 82.5; **LCMS** found [M+H]$^+$ 151.3, Rf 0.34 min; **HR-ESI-MS** $C_6H_6N_4O$ [M+H]$^+$ *m/z* found 151.0626 cald 151.0612. **chemical shift determined from 2D NMR data.*

**2-chloro-N-(6-oxo-6,7-dihydro-1H-pyrazolo[3,4-b]pyridin-4-yl)acetamide, 8**

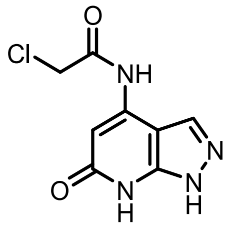

(**e**) To a solution of **S48** (50 mg, 0.33 mmol) and TEA (93 µL, 0.67 mmol) in DCM/DMF (1:1, 0.6 mL) was charged chloroacetyl chloride (106 µL, 1.33 mmol) and stirred at room temperature for 1 h. The reaction was evaporated, added MeOH (1.0 mL) and sat. $Na_2CO_3$ (0.5 mL) and stirred for 10 mins. The reaction was partitioned between DCM and water, and the organic washed with brine, dried ($Na_2SO_4$), and evaporated to dryness. The crude product was purified by preparative HPLC to give **8** as an off-white solid (3.6 mg, 5%).

**$^1$H NMR** (400 MHz, DMSO-$d_6$) δ 13.10 (s, 1H) 11.40 (s, 1H), 10.46 (s, 1H), 8.26 (s, 1H), 6.83 (s, 1H), 4.44 (s, 2H); **LCMS** found [M+H]$^+$ 227.3/229.3, Rf 0.77 min; **HR-ESI-MS** $C_8H_7ClN_4O_2$ [M+Na]$^+$ *m/z* found 249.0102 cald 249.0155.

**Note:** The acylation of **S48** to afford **8** proceeded in low yield and could not be reproducibly scaled to provide sufficient material for a satisfactory $^{13}$C NMR spectrum. Compound **8** was therefore characterised by $^1$H NMR and LCMS prior to biological evaluation. As **8** was inactive in the biochemical BODIPY assay, further resynthesis for additional spectroscopic characterisation was not pursued.

**Urea bioisosteres**

***N*-(5-methyl-1,2-oxazol-3-yl)-1*H*-imidazole-2-carboxamide, S11**

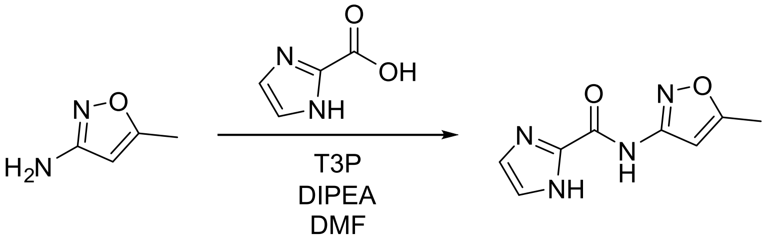

1*H*-imidazole-2-carboxylic acid (100 mg, 0.892 mmol), 3-amino-5-methylisoxazole (105 mg, 1.07 mmol) and anhydrous DIPEA (466 µL, 2.68 mmol) were dissolved in anhydrous DMF (4 mL) and T3P 50 wt% in EtOAc (789 µL, 1.34 mmol) added dropwise. The reaction was stirred at r.t. for 16 h. The reaction was diluted with DCM and the organic phase washed with water, brine dried using $Na_2SO_4$ and the solvent removed *in vacuo*. The crude product was purified using flash column chromatography eluting with a gradient of 20-80% EtOAc in Cy. Further purification using preparative HPLC and subsequent lyophilisation of fractions yielded **S11** (12.7 mg, 66.1 µmol, 7%) as a white solid.

**$^1$H NMR** (400 MHz, DMSO-*d*$_6$) δ 13.36 (s, 1H), 10.87 (s, 1H,), 7.41 (s, 1H), 7.16 (s, 1H), 6.64 (d, J = 0.9 Hz, 1H), 2.40 (d, J = 0.9 Hz, 3H); **$^{13}$C NMR** (101 MHz, DMSO-*d*$_6$) **δ** 169.4, 157.5, 156.4, 139.8, 129.6, 121.3, 96.8, 12.1; **LR-ESI-MS:** $C_8H_9N_4O_2$ [M+H]$^+$ *m/z* found 193.28 cald 193.07; **HR-ESI-MS:** $C_8H_9N_4O_2$ [M+H]$^+$ *m/z* found 193.0739 cald 193.0720.

***N*-(5-methylisoxazol-3-yl)-1*H*-indole-2-carboxamide, S12**

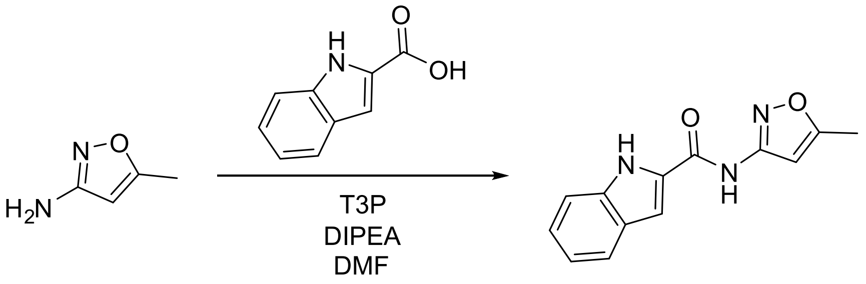

1*H*-indole-2-carboxylic acid (250 mg, 1.55 mmol), 3-amino-5-methylisoxazole (183 mg, 1.86 mmol) and DIPEA (811 µL, 4.65 mmol) were dissolved in anhydrous DCM (10 mL) and T3P 50 wt% in EtOAc (1.40 mL, 2.33 mmol) added dropwise before stirring at r.t. for 16 h. The reaction was washed with water, dried using $Na_2SO_4$ and solvent removed *in vacuo*. The crude product was purified using flash column chromatography eluting with a gradient of 0-100% EtOAc in Cy. Further purification using preparative HPLC and subsequent lyophilisation of fractions yielded **S12** (6.0 mg, 24.9 µmol, 2%) as a brown solid.

**$^1$H NMR** (400 MHz, DMSO-$d_6$) δ 11.80 (s, 1H), 11.33 (s, 1H), 7.66 (d, J = 8.2 Hz, 1H), 7.55 (s, 1H), 7.46 (d, J = 8.2 Hz, 1H), 7.29-7.19 (m, 1H), 7.11-7.02 (m, 1H), 6.79 (s, 1H), 2.43 (s, 3H); **$^{13}$C NMR** (101 MHz, DMSO-$d_6$) δ 169.4, 159.4, 158.4, 137.1, 130.1, 126.9, 124.3, 122.1, 120.1, 112.5, 105.4, 96.9, 12.1; **LR-ESI-MS:** $C_{13}H_{12}N_3O_2$ $[M+H]^+$ *m/z* found 242.28 cald 242.09; **HR-ESI-MS:** $C_{13}H_{11}N_3O_2Na$ $[M+Na]^+$ *m/z* found 264.0747 cald 264.0743.

##### (*R*)-*N*-(5-methylisoxazol-3-yl)pyrrolidine-2-carboxamide, S13

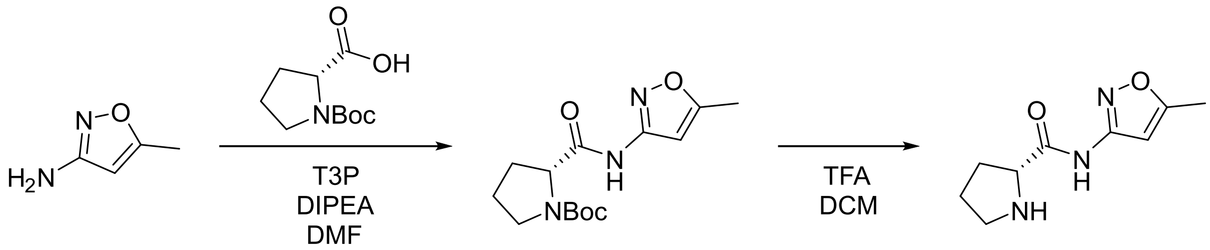

**S49** **S13**

*N*-Boc-D-proline (250 mg, 1.16 mmol), 3-amino-5-methylisoxazole (137 mg, 1.39 mmol) and DIPEA (607 µL, 3.48 mmol) were dissolved in anhydrous DCM (10 mL) and T3P 50 wt% in EtOAc (1.00 mL, 1.74 mmol) added dropwise. The reaction was stirred at r.t. for 16 h. The reaction was washed with water, brine, dried using $Na_2SO_4$ and the solvent removed *in vacuo*. The crude product was purified using flash column chromatography eluting with a gradient of 0-60% EtOAc in Cy. *tert*-butyl (*R*)-2-((5-methylisoxazol-3-yl)carbamoyl)pyrrolidine-1-carboxylate **S49** (205 mg, 0.697 mmol, 60%) was obtained as a white solid.

**$^1$H NMR** (400 MHz, 348 K, DMSO-$d_6$) δ 10.67 (s, 1H), 6.59-6.57 (m, 1H), 4.30 (dd, J = 8.2, 3.5 Hz, 1H), 3.46-3.31 (m, 2H), 2.37 (d, J = 0.8 Hz, 3H), 2.23-2.13 (m, 1H), 1.93-1.77 (m, 3H), 1.34 (s, 9H); **$^{13}$C NMR** (101 MHz, DMSO-$d_6$) δ 171.8, 171.3, 169.5, 169.4, 158.1, 158.1, 153.5, 153.0, 96.3, 96.2, 78.8, 78.6, 59.8, 59.5, 46.7, 46.5, 30.8,

30.0, 28.1, 27.9, 24.0, 23.4, 12.1; **LR-ESI-MS:** $C_{14}H_{22}N_3O_4$ $[M+H]^+$ *m/z* found 296.51 cald 296.16; **HR-ESI-MS:** $C_{14}H_{21}N_3O_4Na$ $[M+Na]^+$ *m/z* found 318.1259 cald 318.1424.

Reduced rotation of Boc group at 298 K resulted in two rotamer values per carbon in the $^{13}C$ NMR spectra

**S49** (123 mg, 0.416 mmol) was dissolved in DCM (2.4 mL). TFA (2.1 mL, 27.8 mmol) was added, and the reaction stirred at r.t. for 3 h. The volatile reagents were removed *in vacuo*. The crude oil was redissolved in minimal DCM and washed with saturated aq. $Na_2CO_3$. The organic phases were combined, dried using $Na_2SO_4$, and the solvent removed *in vacuo*. **S13** (61.2 mg, 0.313 mmol, 75%) was obtained as a yellow oil.

**$^1$H NMR** (400 MHz, DMSO-*d*$_6$) δ 6.64 (d, J = 1.0 Hz, 1H), 3.76 (dd, J = 8.8, 5.7 Hz, 1H), 2.88 (t, J = 6.6 Hz, 2H), 2.37 (d, J = 1.0 Hz, 3H), 2.09-2.00 (m, 1H), 1.79-1.71 (m, 1H), 1.68-1.61 (m, 2H); **$^{13}$C NMR** (101 MHz, DMSO-*d*$_6$) δ 173.3, 169.8, 157.6, 95.9, 60.3, 46.7, 30.2, 25.8, 12.1; **LR-ESI-MS:** $C_9H_{14}N_3O_2$ $[M+H]^+$ *m/z* found 196.26 cald 196.10; **HR-ESI-MS:** $C_9H_{14}N_3O_2$ $[M+H]^+$ *m/z* found 196.1099 cald 196.1081.

**(*S*)-*N*-(5-methylisoxazol-3-yl)pyrrolidine-2-carboxamide, S14**

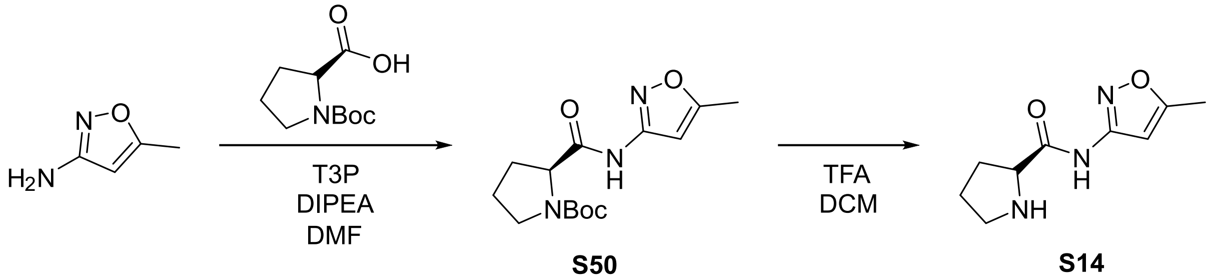

**S50**

**S14**

*N*-Boc-L-proline (250 mg, 1.16 mmol), 3-amino-5-methylisoxazole (137 mg, 1.39 mmol) and anhydrous DIPEA (607 mL, 3.48 mmol) were added to anhydrous DCM (10 mL) and T3P 50 wt% in EtOAc (1.00 mL, 1.74 mmol) was added dropwise. The reaction was stirred at r.t. for 16 h before the organic layer was washed with water, brine, dried using $Na_2SO_4$ and the solvent removed *in vacuo*. The crude product was purified using flash chromatography eluting with a gradient of 0-60% EtOAc in Cy. *tert*-butyl (*S*)-2-((5-methylisoxazol-3-yl)carbamoyl)pyrrolidine-1-carboxylate **S50** (168 mg, 0.568 mmol, 49%) was obtained as a white solid.

**$^1$H NMR** (400 MHz, 348 K, DMSO-$d_6$) δ 10.67 (s, 1H, 6.58 (d, J = 0.9 Hz, 1H), 4.30 (dd, J = 8.3, 3.5 Hz, 1H), 3.47-3.30 (m, 2H), 2.37 (d, J = 0.9 Hz, 3H), 2.23-2.14 (m, 1H), 1.93-1.76 (m, 3H), 1.34 (s, 9H); **$^{13}$C** NMR (101 MHz, DMSO-$d_6$) δ 171.8, 171.3, 169.5, 169.5, 158.1, 158.1, 153.6, 153.0, 96.3, 96.2, 78.8, 78.6, 59.8, 59.5, 46.7, 46.5, 30.8, 30.0, 28.1, 27.9, 24.0, 23.4, 12.1; **LR-ESI-MS:** $C_{14}H_{22}N_3O_4$ $[M+H]^+$ *m/z* found 296.45 cald 296.16; **HR-ESI-MS:** $C_{14}H_{21}N_3O_4Na$ $[M+Na]^+$ *m/z* found 318.1259 cald 318.1424.

Reduced rotation of Boc group at 298 K resulted in two rotamer values per carbon in the $^{13}$C NMR spectra

**S50** (86.7 mg, 0.294 mmol) was dissolved in anhydrous DCM (1.7 mL). TFA (1.5 mL, 19.7 mmol) was added, and the reaction stirred at r.t. for 1.5 h. The volatile reagents were removed *in vacuo*. The crude oil was redissolved in minimal DCM and washed with saturated aq. $Na_2CO_3$.The organic phases were combined, dried using $Na_2SO_4$, and the solvent removed *in vacuo*. **S14** (53.2 mg, 0.273 mmol, 93%) was obtained as a yellow oil.

**$^1$H NMR** (400 MHz, DMSO-$d_6$) δ 10.53 (s, 1H), 6.64 (d, J = 1.0 Hz, 1H), 3.75-3.70 (m, 1H), 2.86 (t, J = 6.5 Hz, 2H), 2.37 (d, J = 1.0 Hz, 3H), 2.08-1.97 (m, 1H), 1.78-1.69 (m, 1H), 1.67-1.60 (m, 2H); **$^{13}$C NMR** (101 MHz, DMSO-$d_6$) δ 173.6, 169.7, 157.5, 95.9, 60.4, 46.7, 30.2, 25.9, 12.1; **LR-ESI-MS:** $C_9H_{14}N_3O_2$ $[M+H]^+$ *m/z* found 196.26 cald 196.10; **HR-ESI-MS:** $C_9H_{14}N_3O$ $[M+H]^+$ *m/z* found 196.1098 cald 196.1081.

**(*R*)-*N*-(5-methylisoxazol-3-yl)piperidine-2-carboxamide, S15**

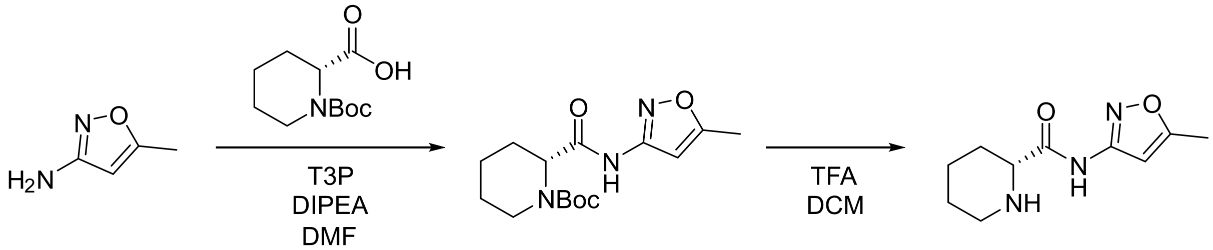

**S51** **S15**

(2*R*)-1-[(*tert*-butoxy)carbonyl]piperidine-2-carboxylic acid (257 mg, 1.12 mmol), 3-amino-5-methylisoxazole (132 mg, 1.35 mmol) and anhydrous DIPEA (586 µL, 3.36 mmol) were dissolved in anhydrous DCM (10 mL) and T3P 50 wt% in EtOAc (991 µL, 1.68 mmol) added dropwise. The reaction was stirred at r.t. for 72 h. Additional T3P (991 µL, 1.68 mmol was added and the reaction stirred at r.t. for 16 h. The crude reaction was diluted with water, and the organic phase was washed with water, brine, dried using $Na_2SO_4$ and the solvent removed *in vacuo*. The crude product was purified using flash column chromatography eluting with a gradient of 0-60% EtOAc in Cy to give *tert*-butyl (*R*)-2-((5-methylisoxazol-3-yl)carbamoyl)piperidine-1-carboxylate **S51** (34.3 mg, 0.111 mmol, 10%) as a white solid.

**$^1$H NMR** (400 MHz, 348 K, DMSO-$d_6$) **δ** 10.61 (s, 1H), 6.57 (d, J = 0.9 Hz, 1H), 4.69 (dd, J = 5.9, 2.3 Hz, 1H), 3.87-3.68 (m, 1H), 3.22 (td, J = 12.5, 3.4 Hz, 1H), 2.37 (d, J = 0.9 Hz, 3H), 2.14-1.95 (m, 1H), 1.81-1.49 (m, 3H), 1.38 (s, 9H), 1.35-1.21 (m, 2H); **$^{13}$C NMR** (101 MHz, DMSO-$d_6$) δ 171.7, 170.0, 158.5, 155.6, 96.8, 79.5, 55.3, 54.1, 42.4, 41.6, 28.5, 27.7, 24.7, 24.5, 19.9, 12.6; **LR-ESI-MS:** $C_{15}H_{24}N_3O_4$ $[M+H]^+$ *m/z* found 310.56 cald 310.17; **HR-ESI-MS:** $C_{15}H_{23}N_3O_4Na$ $[M+Na]^+$ *m/z* found 332.1409 cald 332.1581.

Reduced rotation of Boc group at 298 K resulted in two rotamer values per carbon in the $^{13}$C NMR spectra

**S51** (50.1 mg, 0.162 mmol) was dissolved in DCM (1 mL). TFA (836 □L, 10.9 mmol) was added, and the reaction was stirred at r.t. for 3 h. The volatile reagents were removed *in vacuo*. The crude oil was redissolved in minimal DCM and washed with saturated aq. $Na_2CO_3$. The organic phases were combined, dried using $Na_2SO_4$, and the solvent removed *in vacuo*. **S15** (30.9 mg, 0.148 mmol, 91%) was obtained as a yellow amorphous solid.

**$^1$H NMR** (400 MHz, DMSO-*d*6) δ 6.61 (d, J = 1.0 Hz, 1H), 3.33-3.25 (m, 1H), 2.97-2.87 (m, 1H), 2.57-2.49 (m, 1H), 2.36 (d, J = 1.0 Hz, 3H), 1.79-1.69 (m, 2H), 1.50-1.21 (m, 4H); **$^{13}$C NMR** (101 MHz, DMSO-*d*6) δ 172.8, 169.9, 158.3, 96.7, 59.8, 45.4, 29.7, 26.2, 24.2, 12.6; **LR-ESI-MS:** $C_{10}H_{16}N_3O_2$ $[M+H]^+$ *m/z* found 210.21 cald 210.12; **HR-ESI-MS:** $C_{10}H_{16}N_3O_2$ $[M+H]^+$ *m/z* found 210.1259 cald 210.1237.

**(*S*)-*N*-(5-methylisoxazol-3-yl)morpholine-3-carboxamide, S16**

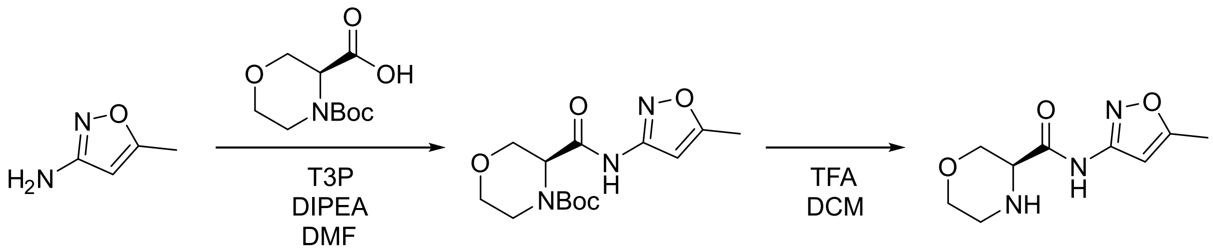

**S52** **S16**

(3*S*)-4-[(*tert*-butoxy)carbonyl]morpholine-3-carboxylic acid (250 mg, 1.08 mmol), 3-amino-5-methylisoxazole (127 mg, 1.30 mmol) and DIPEA (565 µL, 3.24 mmol) were dissolved in anhydrous DCM (10 mL) and T3P 50 wt% in EtOAc (956 µL, 1.62 mmol) added dropwise. The reaction was stirred for 72 h at r.t. and the solvent removed *in vacuo*. The crude product was purified using flash chromatography eluting with a gradient of 20-80% EtOAc in Cy. *tert*-butyl (*S*)-3-((5-methylisoxazol-3-yl)carbamoyl)morpholine-4-carboxylate **S52** (124 mg, 0.399 mmol, 37%) was obtained as a white solid.

**$^1$H** NMR (400 MHz, Methanol-*d*4) δ 6.60 (br s, 1H), 4.49 (br s, 1H), 4.25 (br s, 1H), 3.90 (br s, 1H), 3.76 (dd, J = 12.3, 4.2 Hz, 1H), 3.72-3.63 (m, 1H), 3.57-3.43 (m, 2H), 2.40 (s, 3H), 1.44 (br s, 9H); **$^{13}$C NMR** (101 MHz, Methanol-*d*4) δ 170.0, 169.5, 158.0, 156.1, 95.8, 80.7, 67.7, 67.2, 66.2, 55.8, 54.7, 42.0, 41.0, 27.1, 10.9; **LR-ESI-MS:** $C_{14}H_{22}N_3O_5$ $[M+H]^+$ *m/z* found 312.48 cald 312.15; **HR-ESI-MS:** $C_{14}H_{21}N_3O_5Na$ $[M+Na]^+$ *m/z* found 334.1200 cald 334.1373.

Reduced rotation of Boc group at 298 K resulted in two rotamer values per carbon in the $^{13}C$ NMR spectra

**S52** (124 mg, 0.399 mmol) was dissolved in DCM (2.3 mL). TFA (2.1 mL, 26.7 mmol) was added, and the reaction was stirred at r.t. for 1 h. The volatile reagents were removed *in vacuo*. The crude product was redissolved in minimal DCM and washed with saturated aq. $Na_2CO_3$. The organic phases were combined, dried using $Na_2SO_4$, and the solvent removed *in vacuo*. The crude product was purified using preparative HPLC and subsequent lyophilisation of fractions. **S16** (14.7 mg, 69.6 □mol, 17%) was obtained as a white solid.

**$^1$H NMR** (400 MHz, DMSO-$d_6$) δ 10.64 (s, 1H), 6.61 (s, 1H), 3.83-3.75 (m, 1H), 3.62-3.56 (m, 1H), 3.55-3.47 (m, 2H), 3.44-3.37 (m, 1H), 2.86-2.79 (m, 1H), 2.75-2.65 (m, 1H), 2.37 (s, 3H); **$^{13}$C NMR** (101 MHz, DMSO-$d_6$) δ 170.2, 170.1, 158.2, 96.7, 68.7, 67.3, 58.0, 44.0, 12.5; **LR-ESI-MS:** $C_9H_{14}N_3O_3$ $[M+H]^+$ *m/z* found 212.29 cald 212.10; **HR-ESI-MS:** $C_9H_{13}N_3O_3Na$ $[M+Na]^+$ *m/z* found 234.0889 cald 234.0849.

**(*R*)-2-hydroxy-*N*-(5-methylisoxazol-3-yl)propenamide, S17**

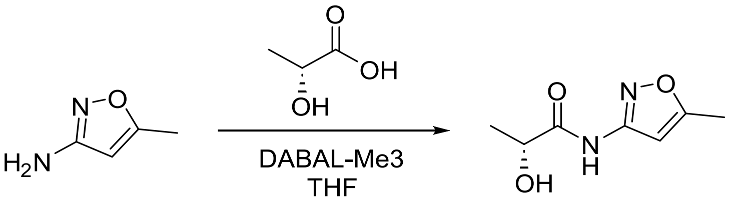

**S17**

3-amino-5-methylisoxazole (142 mg, 1.45 mmol) was added to a solution of DABAL-Me3 (370 mg, 1.45 mmol) in anhydrous THF (9 mL). The solution was stirred at 40 °C for 1.5 h before methyl (2*R*)-2-hydroxypropanoate (92.0 □L, 0.963 mmol) was added and the reaction refluxed for 16 h. The reaction was cooled to r.t., quenched by dropwise addition of aq. HCl (1 M, 10 mL) and extracted with EtOAc. The organic phase was washed with brine, dried using $Na_2SO_4$ and the solvent removed *in vacuo*. The crude product was purified using flash column chromatography eluting with a gradient of 20-70% EtOAc in Cy. Further purification using preparative HPLC and subsequent lyophilisation of fractions yielded **S17** (57.1 mg, 0.336 mmol, 35%) as a white solid.

**$^{1}$H NMR** (400 MHz, MeOD) δ 6.61 (d, J = 0.9 Hz, 1H), 4.27 (q, J = 6.8 Hz, 1H), 2.40 (d, J = 0.9 Hz, 3H), 1.41 (d, J = 6.8 Hz, 3H); **$^{13}$C NMR** (101 MHz, MeOD) δ 176.0, 171.5, 159.1, 97.1, 69.2, 21.0, 12.3; **LR-ESI-MS:** $C_7H_{11}N_2O_3$ $[M+H]^+$ *m/z* found 171.28 cald 171.07; **HR-ESI-MS:** $C_7H_{10}N_2O_3Na$ $[M+Na]^+$ *m/z* found 192.0848 cald 192.0584.

##### (*S*)-2-hydroxy-*N*-(5-methylisoxazol-3-yl)propenamide, S18

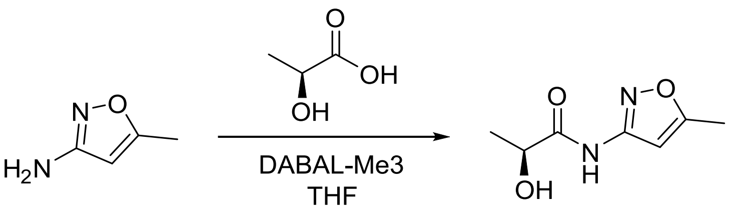

**S18**

3-amino-5-methylisoxazole (142 mg, 1.45 mmol) was added to a stirred solution of DABAL-Me3 (370 mg, 1.45 mmol) in anhydrous THF (9 mL). The solution was stirred at 40 °C for 1.5 h before methyl (2*S*)-2-hydroxypropanoate (92.0 µL, 0.963 mmol) was added and the solution refluxed for 16 h. The reaction was cooled to r.t., quenched by dropwise addition of aq. HCl (1 M, 10 mL) and extracted with EtOAc. The organic phase was washed with brine, dried using $Na_2SO_4$ and the solvent removed *in vacuo*. The crude product was purified using flash column chromatography eluting with a gradient of 20-70% EtOAc in Cy. Further purification using preparative HPLC and subsequent lyophilisation of fractions yielded **S18** (48.4 mg, 0.284 mmol, 30%) as a white solid.

**$^{1}$H NMR** (400 MHz, Methanol-*d*$_4$) δ 6.61 (d, J = 0.9 Hz, 1H), 4.27 (q, J = 6.8 Hz, 1H), 2.40 (d, J = 0.9 Hz, 3H), 1.41 (d, J = 6.8 Hz, 3H); **$^{13}$C NMR** (101 MHz, Methanol-*d*$_4$) δ 176.0, 171.5, 159.1, 97.1, 69.2, 21.0, 12.3; **LR-ESI-MS:** $C_7H_{11}N_2O_3$ $[M+H]^+$ *m/z* found 171.16 cald 171.07; **HR-ESI-MS:** $C_7H_{10}N_2O_3Na$ $[M+Na]^+$ *m/z* found 193.1090 cald 193.5840.

**2-hydroxy-2-methyl-*N*-(5-methylisoxazol-3-yl)propenamide, S19**

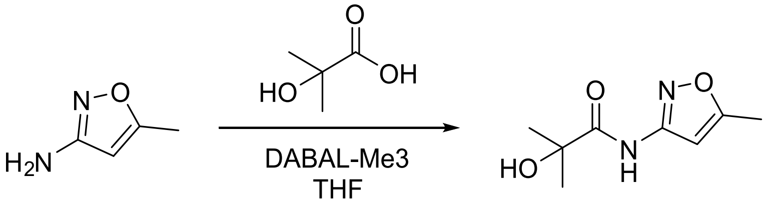

**S19**

3-amino-5-methylisoxazole (62.3 mg, 0.635 mmol) was added to a stirred solution of DABAL-$Me_3$ (163 mg, 0.635 mmol) in anhydrous THF (4 mL). The solution was stirred at 40 °C for 1 h before methyl 2-hydroxy-2-methylpropanoate (49.0 μL, 0.423 mmol) was added. The solution was refluxed for 16 h. The reaction was cooled to r.t., quenched by dropwise addition of aq. HCl (1 M, 10 mL) and extracted with EtOAc. The organic phase was washed with brine, dried using $Na_2SO_4$ and the solvent removed *in vacuo*. The crude product was purified using flash column chromatography eluting with a gradient of 10-60% EtOAc in Cy. **S19** (16.2 mg, 88.0 μmol, 21%) was obtained as a white solid.

**$^1$H NMR** (400 MHz, $CDCl_3$) δ 9.41 (s, 1H), 6.72 (d, J = 0.9 Hz, 1H), 2.40 (d, J = 0.9 Hz, 3H), 1.53 (s, 6H); **$^{13}$C NMR** (101 MHz, $CDCl_3$) δ 174.9, 170.3, 157.8, 96.2, 74.2, 27.9, 12.8; **LR-ESI-MS:** $C_8H_{13}N_2O_3$ $[M+H]^+$ *m/z* found cald 185.09; **HR-ESI-MS:** $C_8H_{12}N_2O_3Na$ $[M+Na]^+$ *m/z* found 207.0619 cald 207.0740.

**1-(5-methylisoxazol-3-yl)-3-phenylurea, S20**

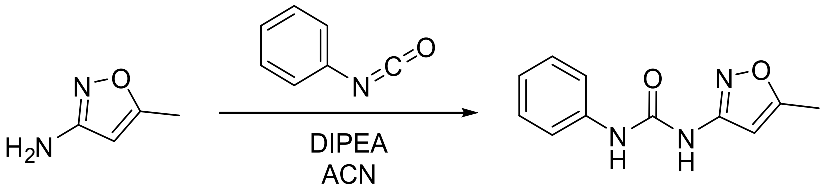

**S20**

Phenyl isocyanate (118 mg, 1.20 mmol), 3-amino-5-methylisoxazole (109 μL, 1.00 mmol) and anhydrous DIPEA (63.0 μL, 0.362 mmol) were dissolved in anhydrous MeCN (3 mL) and the reaction was stirred at 75 °C for 4 h. The reaction was quenched with water, extracted with DCM, the organic phase dried using $Na_2SO_4$ and the solvent removed *in vacuo*. The crude product was purified using flash column chromatography using a gradient of 0-4% [7N $NH_3$ in MeOH] in DCM. The crude product was further purified using preparative HPLC and subsequent lyophilisation of fractions. **S20** (39.8 mg, 0.183 mmol, 18%) was obtained as a white solid.

**$^{1}$H NMR** (400 MHz, MeOD) δ 7.46-7.41 (m, 2H), 7.33-7.27 (m, 2H), 7.06 (m, 1H), 6.40-6.38 (m, 1H), 2.38 (d, J = 0.8 Hz, 3H); **$^{13}$C NMR** (101 MHz, MeOD) δ 171.1, 160.3, 153.8, 139.8, 130.0, 124.6, 120.7, 96.4, 12.2; **LR-ESI-MS:** $C_{11}H_{12}N_3O_2$ $[M+H]^+$ *m/z* found 218.36 cald 218.09; **HR-ESI-MS:** $C_{11}H_{12}N_3O_2$ $[M+H]^+$ *m/z* found 218.0947 cald 218.0924.

##### 1-methoxy-3-(5-methylisoxazol-3-yl)urea (S21) & 2-methoxy-1,3-bis(5-methylisoxazol-3-yl)guanidine, S22

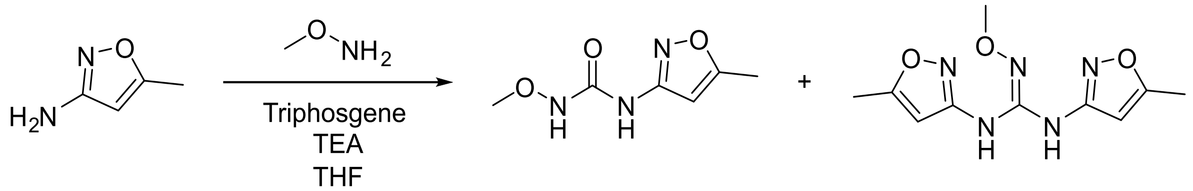

**S21** **S22**

3-amino-5-methylisoxazole (100 mg, 1.02 mmol) was dissolved in anhydrous THF (4 mL) and cooled to 0 °C. Triphosgene (151 mg, 0.510 mmol) was dissolved in anhydrous THF (1 mL) and added dropwise. After 5 mins, anhydrous TEA (426 µL, 3.06 mmol) was added dropwise, and the reaction was stirred at 0 °C for 30 mins. *O*-methylhydroxylamine hydrochloride (128 mg, 1.53 mmol) was dissolved in anhydrous 1:1 THF:DMSO (1 mL) and added dropwise. The reaction was stirred at r.t. for 16 h. The organic phase was washed with water and brine, dried using $Na_2SO_4$ and the solvent removed *in vacuo*. The crude products were purified by preparative HPLC. **S21** (14.9 mg, 87.1 µmol, 9%) was obtained as a yellow solid. **S22** (21.6 mg, 86.0 µmol, 8%) was obtained as a brown solid.

**S21: $^{1}$H NMR** (400 MHz, DMSO-*d*$_6$) δ 9.78 (s, 1H), 9.72 (s, 1H), 6.50 (d, J = 1.0 Hz, 1H), 3.58 (s, 3H), 2.34 (d, J = 1.0 Hz, 3H); **$^{13}$C NMR** (101 MHz, DMSO-*d*$_6$) δ 168.9, 158.5, 155.7, 96.2, 64.1, 12.1; **LR-ESI-MS:** $C_6H_{10}N_3O_3$ $[M+H]^+$ *m/z* found 172.19 cald 172.07; **HR-ESI-MS:** $C_6H_{10}N_3O_3$ $[M+H]^+$ *m/z* found 172.0840 cald 172.0717. **S22: $^{1}$H NMR** (400 MHz, DMSO-*d*$_6$) δ 10.85 (s, 2H), 6.59 (d, J = 1.0 Hz, 2H), 3.84 (s, 3H), 2.40 (d, J = 0.9 Hz, 6H); **$^{13}$C NMR** (101 MHz, DMSO-*d*$_6$) δ 170.1, 157.4, 150.34, 96.6, 64.6,

12.1; **LR-ESI-MS:** $C_{10}H_{14}N_5O_3$ $[M+H]^+$ *m/z* found 252.30 cald 252.11; **HR-ESI-MS:** $C_{10}H_{13}N_5O_3Na$ $[M+Na]^+$ *m/z* found 274.0763 cald 274.0911.

##### 3-(4,5-dihydro-1*H*-imidazol-2-yl)-5-methylisoxazole, S23

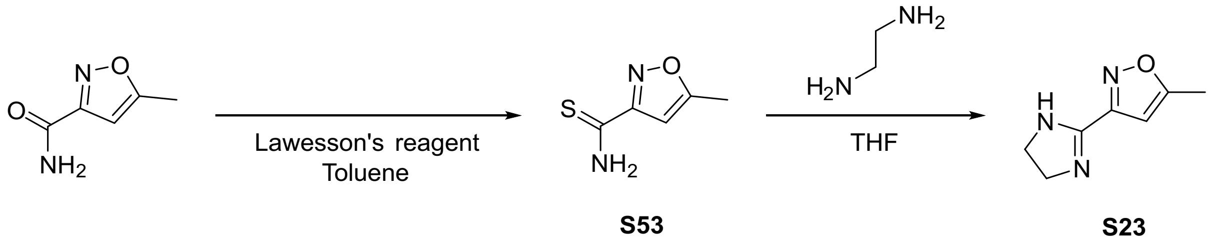

5-methylisoxazole-3-carboxamide (100 mg, 0.793 mmol) and Lawesson's reagent (353 mg, 0.872 mmol) were dissolved in anhydrous toluene (3 mL) and refluxed for 16 h. The reaction was quenched with water and the crude product extracted with EtOAc. The organic phase was washed with water, brine, dried using $Na_2SO_4$ and the solvent removed *in vacuo*. The crude solid was purified using flash column chromatography eluting with a gradient of 0-60% EtOAc in Cy**.** 5-methylisoxazole-3-carbothioamide **S53** was used directly without further purification.

**$^1$H NMR** (400 MHz, $CDCl_3$, crude) δ 6.63 (d, J = 1.1 Hz, 1H), 3.88 (m, 2H), 2.59-2.45 (m, 3H); **LR-ESI-MS:** $C_5H_7N_2OS$ $[M+H]^+$ *m/z* found 143.19 cald 143.02

**S53** (30.0 mg, 0.211 mmol) and ethane-1,2-diamine (20.0 µL, 0.299 mmol) were dissolved in anhydrous THF and refluxed for 16 h. The solvent was removed *in vacuo*. Water was added and the pH adjusted to 11 by the dropwise addition of aq. NaOH (1 M). The aqueous phase was extracted with DCM and EtOAc. The solvent was removed *in vacuo*, the crude product redissolved in minimal MeOH, and applied to an SCX column. The column was washed twice with 10 CV of MeOH and the product eluted using 2 M $NH_3$ in MeOH. The solvent was removed *in vacuo*. **S23** (16.3 mg, 0.108 mmol, 51%) was obtained as a white solid.

**$^1$H NMR** (400 MHz, $CDCl_3$) δ 6.42 (s, 1H), 3.72 (s, 4H), 2.40 (s, 3H); **$^{13}$C NMR** (101 MHz, $CDCl_3$) δ 170.8, 157.1, 155.3, 101.0, 55.2, 12.1; **LR-ESI-MS:** $C_7H_{10}N_3O$ $[M+H]^+$ *m/z* found 152.13 cald 152.08; **HR-ESI-MS:** $C_7H_{10}N_3O$ $[M+H]^+$ *m/z* found 152.0831 cald 152.0818.

**N-(benzo[d]thiazol-2-yl)-5-methylisoxazol-3-amine, S24**

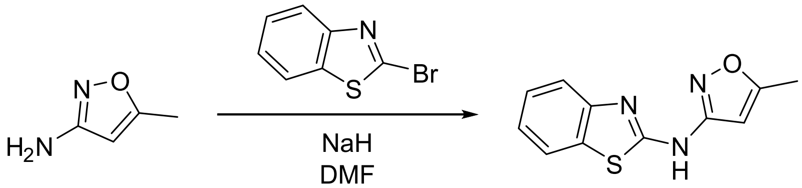

**S24**

NaH 60 wt% dispersion in mineral oil (48.9 mg, 1.22 mmol) was dissolved in anhydrous DMF (2 mL) and cooled to 0 °C. 3-amino-5-methylisoxazole (100 mg, 1.02 mmol) in anhydrous DMF (1.5 mL) was added and the solution stirred for 1 h at 0 °C. 2-bromo-1,3-benzothiazole (218 mg, 1.02 mmol) in anhydrous DMF (1.5 mL) was added and the reaction stirred at r.t. for 16 h. The reaction was quenched with water and extracted 3 times with DCM. The organic phase was washed with brine, dried with $Na_2SO_4$ and solvent removed *in vacuo*. The crude product was purified using flash column chromatography eluting with a gradient of 0-50% EtOAc in Cy. The product was further purified using preparative HPLC and subsequent lyophilisation of fractions. **S24** (35.5 mg, 0.154 mmol, 15%) was obtained as a white solid.

**$^1$H NMR** (400 MHz, DMSO-$d_6$) δ 11.35 (s, 1H), 7.89 (d, J = 7.8 Hz, 1H), 7.62 (d, J = 8.0 Hz, 1H), 7.49-7.30 (m, 1H), 7.30-7.06 (m, 1H), 6.34 (s, 1H), 2.40 (d, J = 0.7 Hz, 3H); **$^{13}$C NMR** (101 MHz, DMSO-$d_6$) δ 169.6, 160.6, 158.9, 149.3, 131.5, 126.0, 122.4, 121.5, 119.3, 95.3, 12.1; **LR-ESI-MS:** $C_{11}H_{10}N_3OS$ $[M+H]^+$ *m/z* found 232.22 cald 232.05; **HR-ESI-MS:** $C_{11}H_{10}N_3OS$ $[M+H]^+$ *m/z* found 232.0408 cald 232.0539.

**3-(ethylamino)-4-((5-methylisoxazol-3-yl)amino)cyclobut-3-ene-1,2-dione, S25**

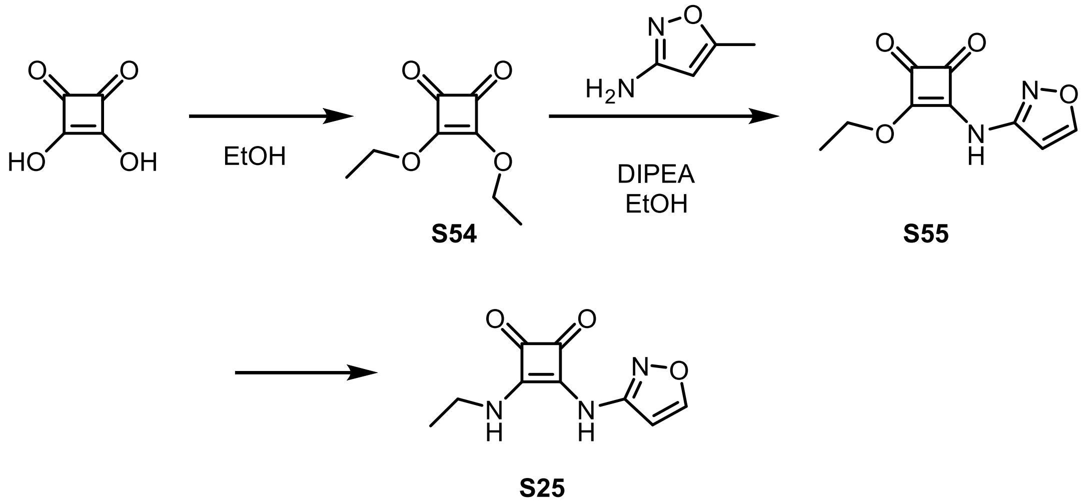

1,2-Dihydroxycyclobutenedione (500 mg, 4.38 mmol) was dissolved in EtOH (5 mL) and the mixture refluxed for 3 hours. The solvent was removed *in vacuo*, further EtOH (5 mL) added, and the mixture refluxed for another hour. This process was repeated 4 times. The solvent was removed *in vacuo* to give 3,4-diethoxycyclobutane-1,2-dione **S54** (686 mg, 4.03 mmol, 92%) as a pink oil.

**$^1$H NMR** (400 MHz, $CDCl_3$) δ 4.67 (q, J = 7.1 Hz, 4H), 1.41 (t, J = 7.1 Hz, 6H); **$^{13}$C NMR** (101 MHz, $CDCl_3$) δ 189.3, 184.2, 70.6, 15.5; **LR-ESI-MS:** $C_8H_{11}O_4$ $[M+H]^+$ *m/z* found 171.28 cald 171.06.

Analytical data consistent with that previously reported.

**S54** (295 µL, 1.99 mmol) and 3-amino-5-methylisoxazole (160 mg, 1.63 mmol) were dissolved in EtOH (15 mL). DIPEA (694 µL, 3.99 mmol) was added and the reaction was stirred for 120 h at 45 °C. The temperature was then increased to 55 °C for 4 h before being allowed to cool to r.t. The precipitate was collected by filtration, washed with ice cold EtOH and dried *in vacuo*. 3-ethoxy-4-((5-methylisoxazol-3-yl)amino)cyclobut-3-ene-1,2-dione **S55** (153 mg, 0.689 mmol, 42%) was obtained as a white solid.

**$^1$H NMR** (400 MHz, DMSO-$d_6$) δ 11.49 (s, 1H), 6.47 (s, 1H), 4.75 (q, J = 7.1 Hz, 2H), 2.38 (s, 3H), 1.40 (t, J = 7.1 Hz, 3H); **$^{13}$C NMR** (101 MHz, DMSO-$d_6$) δ 187.0, 185.2,

179.6, 170.5, 169.6, 158.4, 96.5, 69.8, 15.6, 12.2; **LR-ESI-MS:** $C_{10}H_{11}N_2O_4$ $[M+H]^+$ *m/z* found 223.28 cald 223.07; **HR-ESI-MS:** $C_{10}H_{10}N_2O_4Na$ $[M+Na]^+$ *m/z* found 245.0461 cald 245.0533.

**S55** (36.7 mg, 0.165 mmol) and ethylamine (17.0 µL, 0.330 mmol) were dissolved in EtOH (1.2 mL) and stirred at r.t. for 16 h. The precipitated solid was collected by filtration and washed with ice cold EtOH. **S25** (30.3 mg, 0.137 mmol, 83%) was obtained as a white solid.

**$^1$H NMR** (400 MHz, DMSO-$d_6$) δ 10.68 (s, 1H), 7.80 (t, J = 5.5 Hz, 1H), 6.37 (s, 1H), 3.69-3.57 (m, 2H), 2.36 (s, 3H), 1.18 (t, J = 7.2 Hz, 3H); **$^{13}$C NMR** (101 MHz, DMSO-$d_6$) δ 185.6, 180.8, 170.6, 169.2, 162.1, 158.9, 95.4, 38.7, 16.6, 12.1; **LR-ESI-MS:** $C_{10}H_{12}N_3O_3$ $[M+H]^+$ *m/z* found 222.25 cald 222.08; **HR-ESI-MS:** $C_{10}H_{12}N_3O_3$ $[M+H]^+$ *m/z* found 222.0901 cald 222.0873.

##### ***N*-(4,5-dihydro-1*H*-imidazol-2-yl)-5-methylisoxazol-3-amine, S26**

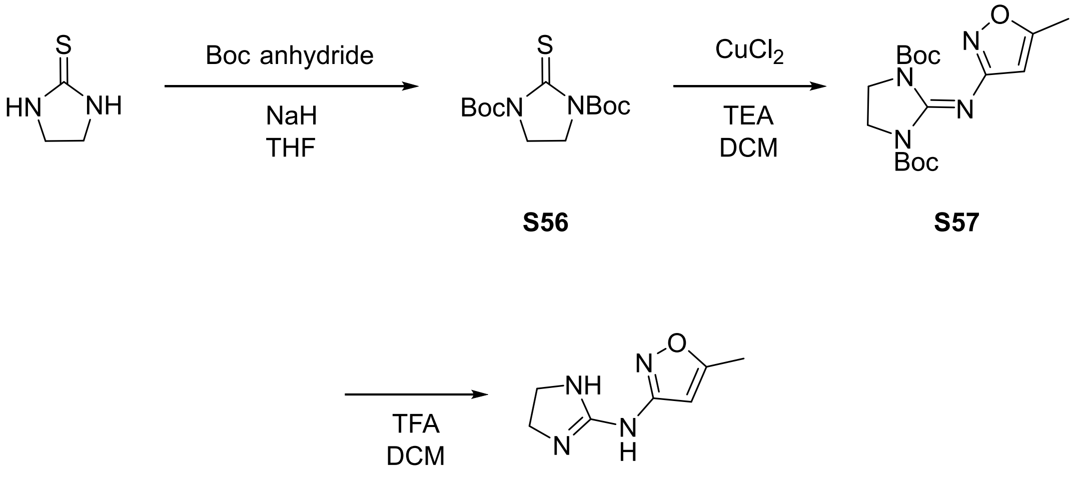

**S56** **S57** **S26**

Imidazolidine-2-thione (605 mg, 5.92 mmol) was dissolved in anhydrous THF (90 mL) and cooled to 0 °C before NaH 60 wt% dispersion in mineral oil (1.07 g, 26.6 mmol) was added. After 5 mins, the ice bath was removed, and the reaction stirred at r.t. for 10 mins. The reaction was then cooled to 0 °C and Boc anhydride (1.07 g, 13.0 mmol) added. The reaction was stirred at 0 °C for 30 minutes and then stirred at r.t. for 16 h. The reaction was quenched by the dropwise addition of saturated aq. $NaHCO_3$ (100 mL). The mixture was poured into water and extracted 3 times with EtOAc. The combined organic phases were washed with brine, dried using $Na_2SO_4$ and the solvent removed *in vacuo*. The crude product was purified using flash column chromatography eluting with a gradient of 0-60% EtOAc in Cy. Di-*tert*-butyl 2-thioxoimidazolidine-1,3-dicarboxylate **S56** (1.57 g, 5.19 mmol, 88%) was obtained as a yellow solid.

**$^1$H NMR** (400 MHz, $CDCl_3$) δ 3.90 (s, 4H), 1.54 (s, 18H); **$^{13}$C NMR** (101 MHz, $CDCl_3$) δ 175.6, 150.5, 84.1, 44.6, 28.2; **LR-ESI-MS:** $C_{13}H_{23}N_2O_4S$ $[M+H]^+$ *m/z* found 303.26 cald 303.13; **HR-ESI-MS:** $C_{13}H_{22}N_2O_4SNa$ $[M+Na]^+$ *m/z* found 325.1582 cald 325.1192.

**S56** (1.37 g, 4.53 mmol) and 3-amino-5-methylisoxazole (444 mg, 4.53 mmol) were dissolved in anhydrous DCM (23.4 mL) and anhydrous TEA (2.1 mL, 14.9 mmol) was added. The mixture was cooled to 0 °C and $CuCl_2$ (640 mg, 4.76 mmol) was added. The reaction was stirred at 0 °C for 30 mins and then stirred at r.t. for 16 h. The reaction was diluted with EtOAc and filtered twice through a pad of celite. Water was added, and the organic phase extracted, washed with brine, dried using $Na_2SO_4$ and the solvent removed *in vacuo*. The crude product was purified using flash column chromatography eluting with a gradient of 0-60% EtOAc in Cy. Di-*tert*-butyl 2-((5-methylisoxazol-3-yl)imino)imidazolidine-1,3-dicarboxylate **S57** (678 mg, 1.85 mmol, 41%) was obtained as a white solid.

**$^1$H NMR** (400 MHz, DMSO-$d_6$) δ 5.87 (d, J = 1.0 Hz, 1H), 3.76 (s, 4H, H-3), 2.27 (d, J = 1.0 Hz, 3H), 1.35 (s, 18H); **$^{13}$C NMR** (101 MHz, DMSO-$d_6$) δ 167.9, 165.7, 149.4, 144.0, 97.4, 82.1, 43.1, 27.5, 12.1; **LR-ESI-MS:** $C_{17}H_{27}N_4O_5$ $[M+H]^+$ *m/z* found 367.49 cald 367.19; **HR-ESI-MS:** $C_{17}H_{26}N_4O_5Na$ $[M+Na]^+$ *m/z* found 389.1582 cald 389.1191.

**S57** (99.2 mg, 0.271 mmol) was dissolved in anhydrous DCM (2.5 mL) and TFA (1.4 mL, 18.1 mmol) added dropwise. The reaction was stirred at r.t. for 16 h. The volatile reagents were removed *in vacuo*. The crude oil was redissolved in minimal DCM and washed with saturated aq. $Na_2CO_3$. The organic phases were combined, dried using $Na_2SO_4$, and the solvent removed *in vacuo*. **S26** (30.5 mg, 0.184 mmol, 68%) was obtained as a light brown solid.

**$^1$H NMR** (400 MHz, DMSO-$d_6$) δ 6.76 (s, 1H), 5.63 (d, J = 1.0 Hz, 1H), 3.42 (s, 4H), 2.24 (d, J = 1.0 Hz, 3H); **$^{13}$C NMR** (101 MHz, DMSO-$d_6$) δ 167.5, 166.9 (C-5), 160.6 (C-1), 99.6 (C-9), 42.0 (C-3), 12.0 (C-10); **LR-ESI-MS:** $C_7H_{11}N_4O$ $[M+H]^+$ *m/z* found 167.19 cald 167.09; **HR-ESI-MS:** $C_7H_{11}N_4O$ $[M+H]^+$ *m/z* found 167.0830 cald 167.0927.

**Non-covalent MC-278 analogues**

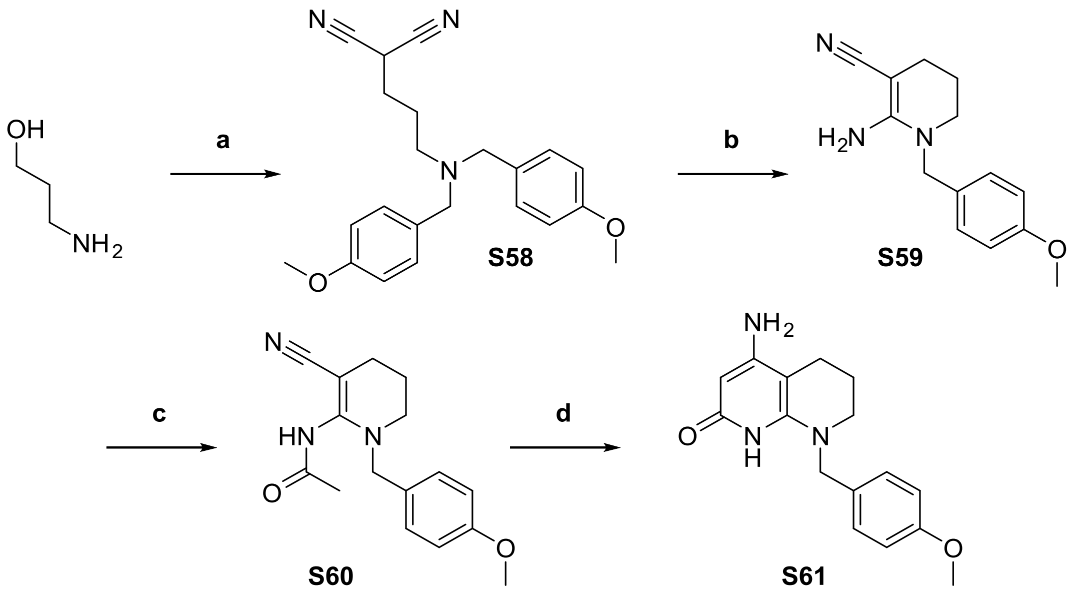

**2-(3-(bis(4-methoxybenzyl)amino)propyl)malononitrile, S58**

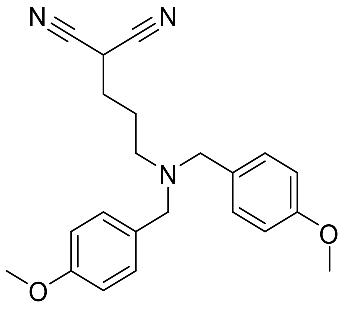

(**a**) To a flask containing $K_2CO_3$ (36.8 g, 266 mmol) and 3-aminopropan-1-ol (5.1 mL, 66.6 mmol) was added MeCN (100 mL) and PMB-Cl (9.0 mL, 66.4 mmol) and stirred at 50 °C for 22 h. The resulting mixture was cooled, filtered, and evaporated to dryness. The residue was partitioned between DCM and water, the organic washed with brine, dried ($Na_2SO_4$) and evaporated to dryness and the resulting residue was used without further purification in the next step.

In a separate oven-dried flask, $PPh_3$ (13.3 g, 50.8 mmol) in THF (100 mL) was cooled to 0 °C and DIAD (10.0 mL, 50.8 mmol) was added dropwise and stirred for 30 min. The residue was then added in one portion and stirred. After an additional 30 min, malononitrile (2.80 g, 42.3 mmol) was added and the reaction stirred at room temperature for 18 h. The mixture was evaporated to dryness and the product purified by flash column chromatography (0-70% EtOAc in Cyclohexane, Rf 50% EtOAc = 0.77) to give **S58** as a clear oil (13.1 g, 54%).

**$^1$H NMR** (400 MHz, DMSO-*d*$_6$) δ 7.27 – 7.19 (m, 4H), 6.93 – 6.85 (m, 4H), 4.70 (t, *J* = 6.7 Hz, 1H), 3.73 (s, 6H), 3.43 (s, 4H), 2.39 (t, *J* = 6.6 Hz, 2H), 1.96 – 1.86 (m, 2H), 1.63 (dq, *J* = 6.7, 14.3 Hz, 2H); **$^{13}$C NMR** (101 MHz, DMSO-*d*$_6$) δ 158.7, 131.4, 130.2, 114.9, 114.1, 114.0, 57.1, 57.0, 55.4, 51.4, 28.1, 24.0, 22.6, 22.4.; **LCMS** found $[M+H]^+$ 364.6, Rf 1.92 min; **HR-ESI-MS** $C_{22}H_{25}N_3O_2$ found $[M+Na]^+$ 386.1766 cald 386.1845.

**2-amino-1-(4-methoxybenzyl)-1,4,5,6-tetrahydropyridine-3-carbonitrile, S59**

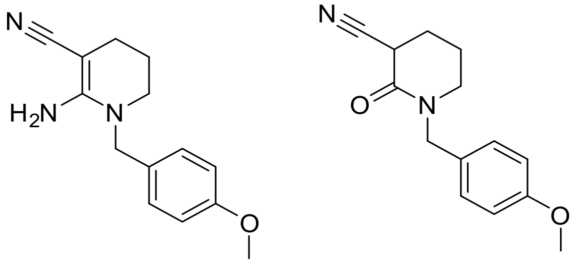

(**b**) **S58** (4.62 g, 12.7 mmol) was dissolved in anhydrous DCE (150 mL) and ACE-Cl (5.5 mL, 51.1 mmol) was added. The reaction was refluxed for 24 h, cooled, and the solvent evaporated. The residue was re-suspended in MeOH (150 mL) and refluxed for 24 h. The reaction was cooled, quenched with sat. $Na_2CO_3$ and added water and DCM. The aqueous was extracted with DCM (3x), and the combined extracts were dried ($Na_2SO_4$) and evaporated to dryness. A mixture of products **S59a** and **S59b** were obtained via flash column chromatography (0-5% MeOH/DCM, Rf 5% MeOH = 0.5) (2.11 g, 68%, 1:1 23/23b), which were not purified further.

**N-(3-cyano-1-(4-methoxybenzyl)-1,4,5,6-tetrahydropyridin-2-yl)acetamide, S60**

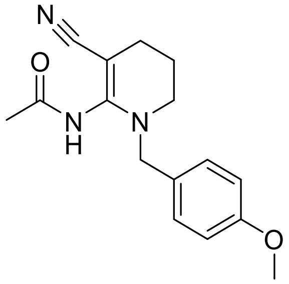

(**c**) The mixture of **S59a, 59b** (1.90 g) was dissolved in MeCN (16 mL) and acetic anhydride (1.2 mL, 1.58 mmol) was added. The reaction was stirred at 50 °C for 2 h. The reaction was cooled, partitioned between DCM and water, and the organic washed with brine, dried ($Na_2SO_4$) and evaporated to dryness. The product was purified by flash column chromatography (0-100% EtOAc in Cyclohexane, Rf 50% EtOAc = 0.21) to give the product **S60** as a red waxy solid (445 mg, 38%).

**$^1$H NMR** (400 MHz, DMSO-$d_6$) δ 9.78 (s, 1H), 7.22 – 7.15 (m, 2H), 6.94 – 6.84 (m, 2H), 4.19 (s, 2H), 3.74 (s, 3H), 3.06 (t, $J$ = 5.4 Hz, 2H), 2.19 (t, $J$ = 6.3 Hz, 2H), 1.98 (s, 3H), 1.55 (p, $J$ = 6.0 Hz, 2H); **$^{13}$C NMR** (101 MHz, DMSO-$d_6$) δ 169.4, 159.0, 148.7, 129.6, 129.5, 122.4, 114.3, 72.1, 55.5, 52.5, 46.6, 24.0, 23.1, 21.0; **LCMS** found $[M+H]^+$ 286.5, Rf 1.38 min.

**4-amino-8-(4-methoxybenzyl)-5,6,7,8-tetrahydro-1,8-naphthyridin-2(1H)-one, S61**

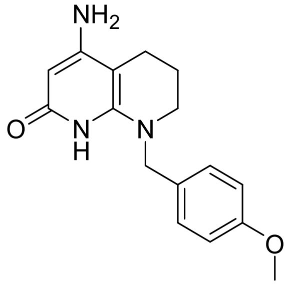

(**d**) **S60** (445 mg, 1.56 mmol) was dissolved in anhydrous THF (12 mL) and LiHMDS (1.0 M in THF, 9.4 mL, 9.36 mmol) was added. The reaction was stirred at 65 °C for 6 h. The reaction was cooled, poured onto water and DCM was added. The aqueous layer was extracted with DCM (3x), and the combined extracts were washed with brine, dried ($Na_2SO_4$) and evaporated to dryness to give the crude product **S52** as a red waxy solid which was used without further purification (373 mg, 84%).

$^{1}$**H NMR** (400 MHz, DMSO-$d_6$) $^{1}$**H NMR** (400 MHz, DMSO) δ 7.13 – 7.05 (m, 2H), 6.85 – 6.79 (m, 2H), 5.28 (br s, 2H), 5.26 (s, 1H), 4.62 (s, 2H), 3.70 (s, 3H), 3.06 (t, J = 5.6 Hz, 2H), 2.29 (t, J = 6.4 Hz, 2H), 1.80 – 1.71 (m, 2H). $^{13}$**C NMR** (101 MHz, DMSO) δ 161.5, 158.6, 156.0, 151.7*, 131.5, 129.2, 114.1, 89.1, 83.4, 55.4, 51.1, 47.3, 21.5, 21.1; **LR-ESI-MS** $C_{16}H_{19}N_3O_2$ $[M+H]^+$ *m/z* found 286.5 cald 286.4. **chemical shift determined from 2D NMR data. DMF present evidently present by NMR.*

**Note:** Due to poor synthetic accessibility and limited reproducibility of the synthetic route, namely the aforementioned cyclisation and acylation, the following compounds **S27–S31** were obtained only in small quantities. Consequently, low-intensity signals in the $^{13}$C NMR spectra, particularly those corresponding to quaternary carbon atoms, could not be observed. Only observed resonances are reported here. Further material was not prepared because these compounds showed no activity in the BODIPY biochemical assay, and repeated synthesis was not considered warranted given the poor reproducibility of the route. In each case, acylation, including regioselectivity, and deprotection was confirmed by $^{1}$H NMR and LCMS before testing.

**General procedure for amides S54 – S58**

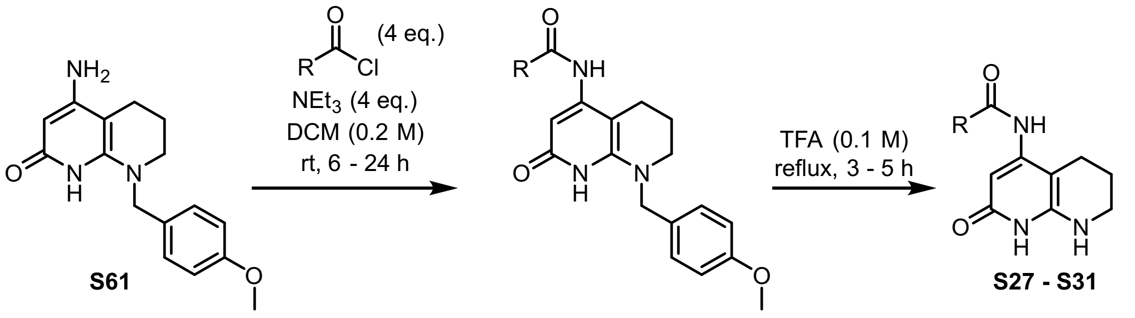

**Crude S52** was dissolved in DCM (0.2 M) and added TEA (4 eq.) and the corresponding acid chloride (4 eq.) consecutively. The reaction was stirred for 6-24 h at room temperature and partitioned between DCM and water. The organic was washed with water, brine, dried ($Na_2SO_4$) and evaporated to dryness. The crude was dissolved in TFA (0.1 M) and stirred at reflux for 3-5 h. The TFA was removed *in vacuo* and the products **S54-S58** were purified by preparative HPLC. All yields are reported over two steps.

**N-(2-oxo-1,2,5,6,7,8-hexahydro-1,8-naphthyridin-4-yl)acetamide, S27**

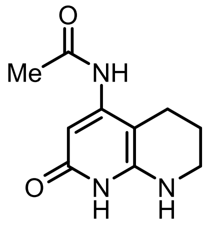

Synthesised as per the general procedure to give **S27** as a green waxy solid (1.3 mg, 10%).

**$^{1}$H NMR** (400 MHz, DMSO-$d_6$) δ 8.83 (s, 1H), 6.02 (s, 1H), 5.80 (s, 1H), 3.19 – 3.15 (m, 2H), 2.36 (t, *J* = 6.3 Hz, 2H), 2.05 (s, 3H), 1.81 – 1.66 (m, 2H); **LCMS** found $[M+H]^+$ 208.5, Rf 0.57 min; **HR-ESI-MS** $C_{10}H_{13}N_3O_2$ $[M+H]^+$ *m/z* found 208.0505 cald 208.1086.

**N-(2-oxo-1,2,5,6,7,8-hexahydro-1,8-naphthyridin-4-yl)cyclopropanecarboxamide, S28**

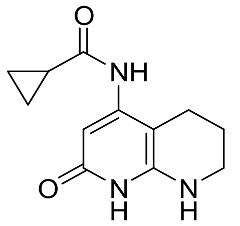

Synthesised as per the general procedure to give **S28** as a green waxy solid (5.1 mg, 13%)

**$^{1}$H NMR** (400 MHz, DMSO-$d_6$) δ 9.05 (s, 1H), 6.01 (s, 1H), 5.86 (br s, 1H), 3.16 (dt, *J* = 2.9, 6.3 Hz, 2H), 2.38 (t, *J* = 6.3 Hz, 2H), 2.03 – 1.92 (m, 1H), 1.73 (p, *J* = 6.3 Hz, 2H), 0.80 – 0.71 (m, 4H); **$^{13}$C NMR** (101 MHz, DMSO-$d_6$) δ 172.6, 161.9, 149.4*, 93.8*, 88.4*, 40.4*, 21.4, 20.6, 14.8, 8.0; **LCMS** found $[M-H]^-$ 232.1, Rf 1.05 min; **HR-ESI-MS** $C_{12}H_{15}N_3O_2$ $[M+H]^+$ *m/z* found 234.0588 cald 234.1243. **chemical shift determined from 2D NMR data.*

**N-(2-oxo-1,2,5,6,7,8-hexahydro-1,8-naphthyridin-4-yl)furan-2-carboxamide, S29**

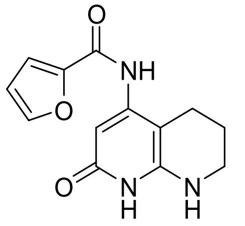

Synthesised as per the general procedure to give **S29** as a pink solid (2.2 mg, 12%)

**$^{1}$H NMR** (400 MHz, DMSO-$d_6$) δ 10.29 (br s, 1H), 9.21 (s, 1H), 7.92 (d, $J$ = 1.7 Hz, 1H), 7.30 (d, $J$ = 3.5 Hz, 1H), 6.70 (dd, $J$ = 3.5, 1.7 Hz, 1H), 5.95 (br s, 1H), 5.86 (s, 1H), 3.20 (dt, $J$ = 2.9, 6.3 Hz, 2H), 2.39 (t, $J$ = 6.3 Hz, 2H), 1.72 (p, $J$ = 6.3 Hz, 2H); **$^{13}$C NMR** (101 MHz, DMSO-d$_6$) δ 161.8, 156.4, 147.7, 146.3, 115.6, 112.7, 95.2, 40.5*, 21.4, 20.7; **LCMS** found [M-H]$^-$ 260.4, Rf 1.07 min; **HR-ESI-MS** $C_{13}H_{13}N_3O_2$ [M+H]$^+$ *m/z* found 260.0310 cald 260.1035. **chemical shift determined from 2D NMR data.*

**N-(2-oxo-1,2,5,6,7,8-hexahydro-1,8-naphthyridin-4-yl)benzamide, S30**

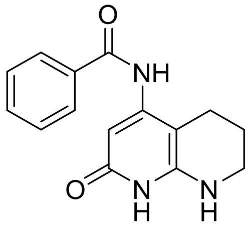

Synthesised as per the general procedure to give **S30** as an off-white solid (1.9 mg, 14%)

**$^{1}$H NMR** (400 MHz, DMSO-$d_6$) δ 10.39 (br s, 1H), 9.48 (s, 1H), 7.94 – 7.87 (m, 2H), 7.63 – 7.55 (m, 1H), 7.55 – 7.47 (m, 2H), 5.97 (s, 1H), 5.79 (s, 1H), 3.20 (dt, $J$ = 2.9, 6.3 Hz, 2H), 2.40 (t, $J$ = 6.3 Hz, 2H), 1.71 (p, $J$ = 6.3 Hz, 2H); **$^{13}$C NMR** (101 MHz, DMSO-d$_6$) δ 165.8, 161.8, 134.9, 132.2, 128.9, 128.2, 95.6*, 40.5*, 21.4, 21.0; **LCMS** found [M+H]$^+$ 270.4, Rf 1.17 min; **HR-ESI-MS** $C_{15}H_{15}N_3O_2$ found [M+H]$^+$ 270.0490 cald 270.1243. **chemical shift determined from 2D NMR data.*

**N-(2-oxo-1,2,5,6,7,8-hexahydro-1,8-naphthyridin-4-yl)-2-phenylacetamide, S31**

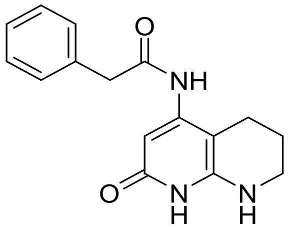

Synthesised as per the general procedure to give **S31** as an off-white solid (2.6 mg, 13%).

**$^{1}$H NMR** (400 MHz, DMSO-$d_6$) δ 10.16 (br s, 1H), 8.99 (s, 1H), 7.36 – 7.15 (m, 5H), 5.99 (s, 1H), 5.87 (br s, 1H), 3.69 (s, 2H), 3.17 (dt, $J$ = 2.9, 6.3 Hz, 2H), 2.32 (t, $J$ = 6.3 Hz, 2H), 1.71 (p, $J$ = 6.3 Hz, 2H); **$^{13}$C NMR** (101 MHz, DMSO-$d_6$) δ 170.0, 161.9, 149.4, 148.4, 136.4, 129.6, 128.8, 127.0, 94.0, 88.9*, 43.5, 40.3, 21.4, 20.6; **LCMS** found [M-H]$^-$ 282.2, Rf 1.19 min; **HR-ESI-MS** $C_{16}H_{17}N_3O_2$ found [M+H]$^+$ 284.0607 cald 284.1399. ***chemical shift determined from 2D NMR data.***

**N-alkyl MC-278 analogues**

**4-chloro-1,8-naphthyridin-2-ol, S62**

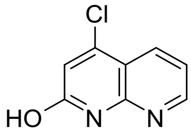

(**a**) To a solution of 2,4-dichloroquinoline (5.146 g, 25.9 mmol) in dioxane (51.7 mL) was added HCl (6N, 76.2 mL) dropwise at 0 °C. The mixture was refluxed overnight, cooled and water added. The resulting precipitate was filtered and dried under vacuum to give **S62** as a white solid (3.62 g, 78%).

**$^{1}$H NMR** (400 MHz, DMSO-$d_6$) δ 12.45 (s, 1H), 8.63 (dd, $J$ = 4.7, 1.7 Hz, 1H), 8.26 (dd, $J$ = 8.0, 1.7 Hz, 1H), 7.38 (dd, $J$ = 8.0, 4.7 Hz, 1H), 6.90 (s, 1H); **$^{13}$C NMR** (101 MHz, DMSO-$d_6$) δ 161.6, 152.5, 149.7, 143.4, 134.3, 122.8, 119.4, 113.4; **LC-ESI-MS** found [M+H]$^+$ 181.22, Rf 1.12 min; **HR-ESI-MS** $C_8H_5N_2OCl$ found [M+H]$^+$ 181.0161 cald 181.0169.

**4-(methylamino)-1,8-napthyridin-2-ol, S63**

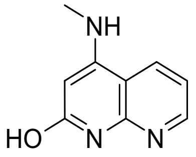

(**b**) To a solution of naphthyridine **S62** (572 mg, 3.17 mmol) in DMSO (8.0 mL) was charged methylamine solution (30% w/w in EtOH, 7.0 mL) and the reaction stirred at 60 $^{o}$C for 16 h. The resulting mixture was cooled and the precipitate filtered to give **S63** as a white solid (545 mg, 98%).

**$^{1}$H NMR** (400 MHz, DMSO-*d*$_6$) δ 11.04 (s, 1H), 8.45 (dd, *J* = 4.7, 1.7 Hz, 1H), 8.29 (dd, *J* = 8.0, 1.7 Hz, 1H), 7.18 (d, *J* = 4.6 Hz, 1H), 7.16 (dd, *J* = 8.0, 4.7 Hz, 1H), 5.23 (s, 1H), 2.80 (d, *J* = 4.6 Hz, 3H); **$^{13}$C NMR** (101 MHz, DMSO-*d*$_6$) δ 164.1, 151.9, 150.7, 150.5, 131.2, 117.3, 109.5, 90.9, 29.9; **LC-ESI-MS** found [M+H]$^+$ 176.47, Rf 0.80 min; **HR-ESI-MS** $C_9H_9N_3O$ found [M+H]$^+$ 176.0812 cald 176.0824.

**4-(butylamino) -1,8-napthyridin-2(*1H*)-one, S64**

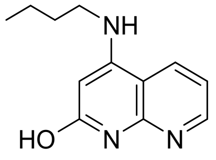

(**b**) To a solution of naphthyridine **S62** (64 mg, 0.34 mmol) in DMSO (1.6 mL) was added n-butylamine (1.0 mL, 10.6 mmol) and the reaction stirred at 45 $^{o}$C for 4 h. The reaction was cooled to 0 $^{o}$C and the resulting precipitate collected by vacuum filtration to give **S64** as a white solid (41 mg, 53%)

**$^{1}$H NMR** (400 MHz, DMSO-*d*$_6$) δ 11.03 (s, 1H), 8.44 (dd, J = 4.7, 1.6 Hz, 2H), 8.40 (d, J = 7.9 Hz, 1H), 7.17 (dd, *J* = 8.0, 4.7 Hz, 1H), 6.99 (t, *J* = 5.3 Hz, 1H), 5.28 (d, *J* = 6.8 Hz, 1H), 3.16 (q, *J* = 6.8 Hz, 2H), 1.62 (p, *J* = 7.5 Hz, 2H), 1.40 (dq, *J* = 14.6, 7.3 Hz, 2H), 0.94 (t, *J* = 7.3 Hz, 3H); **$^{13}$C NMR** (101 MHz, DMSO-*d*$_6$) δ 164.1, 151.0, 150.7, 150.5, 131.5, 117.2, 109.6, 91.0, 42.8, 30.1, 20.3, 14.2; **LC-ESI-MS** found [M+H]$^+$ 218.33, Rf 1.21 min; **HR-ESI-MS** $C_{12}H_{15}N_3O$ found [M+H]$^+$ 218.1288 cald 218.1293.

**4-((2-methoxyethyl)amino)-1,8-napthyridin-2-ol, S65**

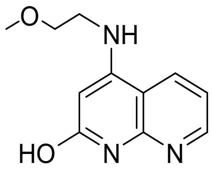

(**c**) To a solution of naphthyridine **S62** (62 mg, 0.34 mmol) in DMSO (1.6 mL) was added 2-methoxyethylamine (1.2 mL, 13.7 mmol) and the reaction stirred at 90 °C for 4 h. The reaction was cooled, partitioned between DCM and water, and the organic evaporated to dryness. The final product was purified by Isolera Biotage LPLC (0-5% MeOH in DCM) to give **S65** as a white solid (53 mg, 71%).

**$^1$H NMR** (400 MHz, DMSO-$d_6$) δ 11.07 (s, 1H), 8.45 (dd, *J* = 4.7, 1.6 Hz, 1H), 8.40 (dd, *J* = 8.2, 1.7 Hz, 1H), 7.18 (dd, *J* = 8.1, 4.6 Hz, 1H), 7.05 (t, *J* = 5.4 Hz, 1H), 5.36 – 5.32 (m, 1H), 3.57 (t, *J* = 5.6 Hz, 2H), 3.37 (t, *J* = 5.5 Hz, 2H), 3.30 (s, 3H); **$^{13}$C NMR** (101 MHz, DMSO-$d_6$) δ 164.1, 150.9, 150.7, 150.5, 131.5, 117.3, 109.5, 91.3, 69.8, 58.6, 42.8; **LC-ESI-MS** found $[M+H]^+$ 220.90, Rf 0.99 min; **HR-ESI-MS** $C_{11}H_{13}N_3O_2$ found $[M+H]^+$ 220.1079 cald 220.1086.

#### General procedure for the reduction of naphthyridinols to generate S33 – S35.

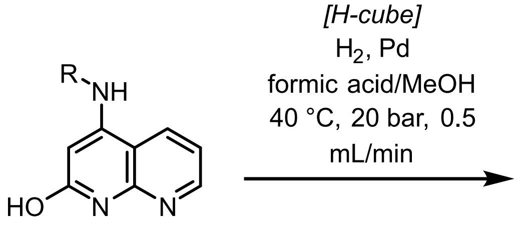

**S63 - S65** → **S33 - S35**

Napthyridinols **S60 – S67** were dissolved in MeOH (0.18 M) and formic acid (0.07 M) and subjected to flow hydrogenation with a Pd cartridge until complete conversion by LCMS (40 °C, 20 bar, 0.5 mL/min). The resulting solutions were evaporated to dryness and purified by preparative HPLC to give **S68 – S74**.

##### 4-(methylamino)-5,6,7,8-tetrahydro-1,8-napthyridin-2(*1H*)-one, S35

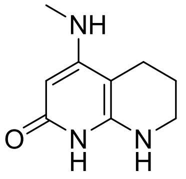

Synthesised as per the general procedure to give **S35** as a red solid (116 mg, 69%).

**$^1$H NMR** (400 MHz, DMSO-$d_6$) δ 5.58 (q, *J* = 4.9 Hz, 1H), 5.33 (s, 1H), 4.52 (s, 2H), 3.10 (q, *J* = 2.2 Hz, 2H), 2.60 (d, *J* = 4.7 Hz, 3H), 2.15 (t, J = 6.4 Hz, 2H), 1.73 (p, *J* = 6.2 Hz, 2H); **$^{13}$C NMR** (101 MHz, DMSO-$d_6$) δ 161.8, 157.2, 145.5, 81.5, 80.4, 40.0*, 29.2, 21.3, 19.0; **LC-ESI-MS** found [M+H]$^+$ 180.31, Rf 0.82 min; **HR-ESI-MS**; C9H13N3O found [M+H]+ 180.1121 cald 180.1137. **chemical shift determined from 2D NMR data.*

##### 4-(butylamino)-5,6,7,8-tetrahydro-1,8-napthyridin-2(*1H*)-one, S34

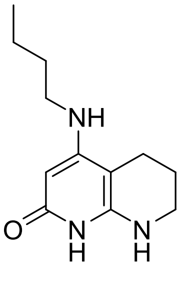

Synthesised as per the general procedure to give **S34** as an off-white solid (30 mg, 55%).

**$^1$H NMR** (400 MHz, DMSO-$d_6$) δ 9.29 (s, 1H), 5.37 (s, 1H), 5.29 (s, 1H), 4.56 (s, 1H), 3.11 (s, 2H), 2.96 (q, *J* = 6.6 Hz, 2H), 2.16 (t, *J* = 6.4 Hz, 2H), 1.73 (dd, *J* = 6.0, 5.3 Hz, 2H), 1.49 (p, *J* = 7.3 Hz, 2H), 1.30 (dt, *J* = 14.6, 7.3 Hz, 2H), 0.90 (q, J = 7.6 Hz, 3H); **$^{13}$C NMR** (101 MHz, DMSO-$d_6$) δ 162.3, 156.9, 146.2, 82.5, 80.8, 42.3, 40.9, 31.0, 21.7, 20.2, 19.5, 14.3; **LC-ESI-MS** found $[M+H]^+$ 222.37, Rf 1.23 min; **HR-ESI-MS** $C_{12}H_{19}N_3O$ found [M+H]+ 222.1598 cald 222.1606.

**4-((2-methoxyethyl)amino)-5,6,7,8-tetrahydro-1,8-napthyridin-2(*1H*)-one, S35**

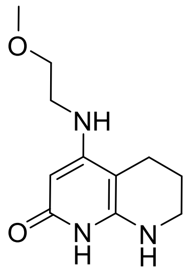

Synthesised as per the general procedure to give **S35** as a purple wax (16 mg, 49%).

**$^1$H NMR** (400 MHz, DMSO-$d_6$) δ 9.31 (br s, 1H), 5.35 (m, 2H), 5.33 (t, *J* = 5.5 Hz, 1H), 4.59 (s, 1H), 3.43 (t, *J* = 5.9 Hz, 2H), 3.26 (s, 3H), 3.16 (t, *J* = 5.9 Hz, 2H), 3.15 – 3.06 (m, 2H), 2.16 (t, *J* = 6.4 Hz, 2H), 1.74 (dt, *J* = 6.4, 4.2 Hz, 2H); **$^{13}$C NMR** (101 MHz, DMSO-$d_6$) δ 161.7, 156.1, 81.4*, 80.5, 70.0, 58.0, 41.7, 39.7*, 21.2, 19.0; **LC-ESI-MS** found $[M+H]^+$ 224.36, Rf 1.03 min; **HR-ESI-MS** $C_{11}H_{17}N_3O_2$ found [M+H]+ 224.1393 cald 224.1399. **chemical shift determined from 2D NMR data.*

#### Covalent urea isosteres

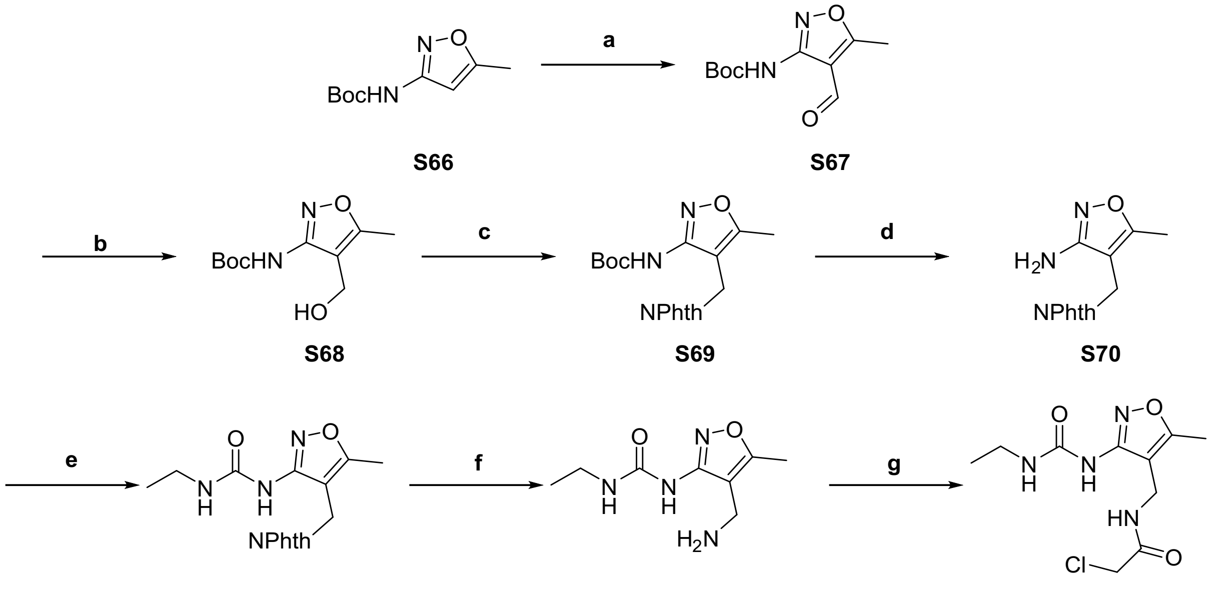

S66 S67 S68 S69 S70 S71 S72 3

##### *tert*-butyl (4-formyl-5-methylisoxazol-3-yl)carbamate, S67

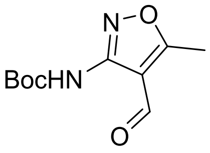

(a) **S66** (3.00 g, 15.1 mmol) was dissolved in anhydrous THF (75 mL) and cooled to -78 °C. *n*-BuLi 2.5 M in hexane (14.5 mL, 36.3 mmol) was added dropwise and stirred at -78 °C for 5 mins, before being allowed to warm to r.t. and stirred for 30 mins. The reaction was once again cooled to -78 °C and anhydrous DMF (2.3 mL, 30.3 mmol) was added dropwise. The reaction was stirred at r.t. for 16 h. The reaction was quenched by the dropwise addition of saturated aq. $NH_4Cl$ and stirred for 30 mins. The crude product was extracted 3 times with EtOAc. The combined organic phases were washed with brine, dried using $Na_2SO_4$ and the solvent removed *in vacuo*. The crude product was purified using flash column chromatography eluting with a gradient of 0-50% EtOAc in Cy. **S67** (2.04 g, 9.01 mmol, 60%) was obtained as a yellow oil.

**$^1$H NMR (400 MHz, $CDCl_3$)** δ 9.89 (s, 1H), 8.38 (s, 1H), 2.67 (s, 3), 1.53 (s, 9H, H); **$^{13}$C NMR (101 MHz, $CDCl_3$)** δ 184.6, 176.8, 156.5, 150.3, 109.2, 82.7, 28.2, 12.0; **LR-**

**ESI-MS:** $C_{10}H_{15}N_2O_4$ $[M+H]^+$ *m/z* found 227.59 cald 227.10; **HR-ESI-MS:** $C_{10}H_{14}N_2O_4Na$ $[M+Na]^+$ *m/z* found 249.0712 cald 249.0846.

***tert*-butyl (4-(hydroxymethyl)-5-methylisoxazol-3-yl)carbamate, S68**

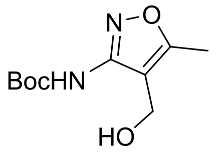

(b) **S67** (1.86 g, 8.22 mmol) was dissolved in anhydrous MeOH (40 mL) and cooled to 0 °C. $NaBH_4$ (373 mg, 9.87 mmol) was added, and the reaction stirred at r.t. for 1 h. The reaction was quenched with water and extracted 3 times with DCM. The combined organic phases were washed with brine, dried using $Na_2SO_4$ and the solvent removed *in vacuo*. The crude product was purified using flash column chromatography eluting with a gradient of 10-60% EtOAc in Cy. **S68** (946 mg, 4.14 mmol, 50%) was obtained as a white solid.

**$^1$H NMR (400 MHz, $CDCl_3$) δ** 7.00 (s, 1H), 4.37 (d, J = 6.8 Hz, 2H), 3.79 (t, J = 6.8 Hz, 1H), 2.41 (s, 3H), 1.51 (s, 9H); **$^{13}$C NMR (101 MHz, $CDCl_3$) δ** 169.3, 156.9, 153.3, 109.6, 82.7, 53.3, 28.1, 11.4; **LR-ESI-MS:** $C_{10}H_{17}N_2O_4$ $[M+H]^+$ *m/z* found 229.28 cald 229.11; **HR-ESI-MS:** $C_{10}H_{16}N_2O_4Na$ $[M+Na]^+$ *m/z* found 251.0868 cald 251.1002.

***tert*-butyl (4-((1,3-dioxoisoindolin-2-yl)methyl)-5-methylisoxazol-3-yl)carbamate, S69**

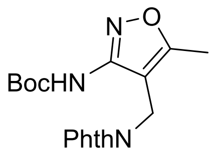

(c) **S68** (736 mg, 3.23 mmol), phthalimide (617 mg, 4.19 mmol) and DPPE (771 mg, 1.94 mmol) were dissolved in anhydrous THF (29 mL) and cooled to 0 °C. DtBAD (965 mg, 4.19 mmol) was added, and the reaction stirred at r.t. for 16 h. The reaction was filtered, water added to the filtrate and extracted 3 times with DCM. The combined organic phases were washed with brine, dried using $Na_2SO_4$ and the solvent removed *in vacuo*. The crude product was purified using flash column chromatography eluting with a gradient of 0-40 % EtOAc in Cy. **S69** (401 mg, 1.12 mmol, 35%) was obtained as a white solid.

**$^1$H NMR (400 MHz, CDCl$_3$) δ** 8.04 (s, 1H), 7.91-7.82 (m, 2H), 7.80-7.70 (m, 2H), 4.55 (s, 2H), 2.51 (s, 3H), 1.56 (s, 9H); **$^{13}$C NMR (101 MHz, CDCl$_3$) δ** 169.0, 168.6, 156.9, 151.4, 134.6, 131.9, 123., 103.8, 81.7, 29.1, 28.3, 11.5; **LR-ESI-MS:** $C_{18}H_{20}N_3O_5$ $[M+H]^+$ *m/z* found 358.37 cald 358.14; **HR-ESI-MS:** $C_{18}H_{19}N_3O_5Na$ $[M+Na]^+$ *m/z* found 380.1026 cald 380.1217.

**2-((3-amino-5-methylisoxazol-4-yl)methyl)isoindoline-1,3-dione, S70**

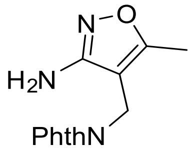

(d) **S69** (229 mg, 0.639 mmol) was dissolved in anhydrous DCM (5.9 mL) and TFA (3.3 mL, 42.8 mmol) was added dropwise. The reaction was stirred at r.t. for 1.5 h. The volatile reagents were removed *in vacuo*. The crude oil was redissolved in minimal DCM and washed with saturated aq. $Na_2CO_3$. The orga0ic phases were combined, dried using $Na_2SO_4$, and the solvent removed *in vacuo*. **S79** (164 mg, 0.638 mmol, 100%) was obtained as a light brown solid and used directly without further purification

**LR-ESI-MS:** $C_{13}H_{12}N_3O_3$ $[M+H]^+$ *m/z* found 258.31 cald 258.08; **HR-ESI-MS:** $C_{13}H_{11}N_3O_3Na$ $[M+Na]^+$ *m/z* found 280.0546 cald 280.0693.

**1-(4-((1,3-dioxoisoindolin-2-yl)methyl)-5-methylisoxazol-3-yl)-3-ethylurea, S71**

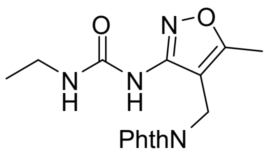

(e) **S70** (262 mg, 1.02 mmol) was dissolved in anhydrous DCM (17 mL) and cooled to 0 °C. Triphosgene (151 mg, 0.508 mmol) was dissolved in anhydrous DCM (1 mL) and added dropwise. The reaction was stirred for 5 mins before anhydrous TEA (425 μL, 3.05 mmol) was added dropwise. The reaction was stirred at 0 °C for 20 mins and ethylamine (763 µL, 1.53 mmol) was added dropwise. The reaction was allowed to warm to r.t. and stirred for 2 h. Additional ethylamine (112 µL, 0.383 mmol) was added and stirred at r.t for 1 h. The reaction was poured into ice water and diluted with DCM. The organic phase was extracted, washed with brine, dried using $Na_2SO_4$ and the solvent was removed *in vacuo*. The crude product was purified by trituration in EtOH. **S80** was used directly without further purification.

**LR-ESI-MS:** $C_{16}H_{17}N_4O_4$ $[M+H]^+$ *m/z* found 329.38 cald 329.12.

**1-(4-(aminomethyl)-5-methylisoxazol-3-yl)-3-ethylurea, S72**

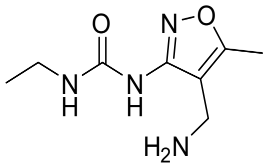

(f) **S71** (118 mg, 0.360 mmol) was suspended in EtOH (5 mL) and hydrazine monohydrate (118 µL, 1.95 mmol) added. The reaction was stirred at r.t. for 16 h. The reaction was filtered, the filtrate collected, and the solvent removed *in vacuo*. **S81** was used directly without further purification.

**LR-ESI-MS:** $C_8H_{15}N_4O_2$ $[M+H]^+$ *m/z* found 199.26 cald 199.12.

**2-chloro-*N*-((3-(3-ethylureido)-5-methylisoxazol-4-yl)methyl)acetamide, S3**

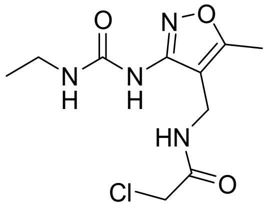

(g) **S81** (72.3 mg, 0.365 mmol) was dissolved in anhydrous DCM (3 mL) and anhydrous DMF (6 mL) and anhydrous TEA (102 µL, 0.729 mmol) added. The solution was cooled to 0 °C and 2-chloroacetyl chloride (35.0 µL, 0.438 mmol) added. The solution was stirred at r.t. for 2 h. The solvent was removed *in vacuo* and the crude product purified using preparative HPLC and subsequent lyophilisation of fractions. **3** (71.0 mg, 0.258 mmol, 71%) was obtained as a yellow solid.

**$^1$H NMR (400 MHz, DMSO-$d_6$) δ** 8.93 (s, 1H, H-9), 8.58 (t, J = 5.8 Hz, 1H), 7.25 (t, J = 5.4 Hz, 1H), 4.13 (s, 2H), 4.04 (d, J = 5.8 Hz, 2H), 3.18 (qd, J = 7.2, 5.4 Hz, 2H), 2.36 (s, 3H), 1.07 (t, J = 7.2 Hz, 3H); **$^{13}$C NMR (101 MHz, DMSO-$d_6$) δ** 167.3, 166.5, 157.9, 153.6, 105.5, 42.5, 34.4, 30.5, 15.2, 10.9; **LR-ESI-MS:** $C_{10}H_{16}ClN_4O_3$ $[M+H]^+$ *m/z* found 275.27 cald 275.08; **HR-ESI-MS:** $C_{10}H_{15}ClN_4O_3Na$ $[M+Na]^+$ *m/z* found 297.0571 cald 297.0725.

**Synthesis of (3-(3-ethylureido)-5-methylisoxazol-4-yl)methyl 4-cyanobenzoate (4)**

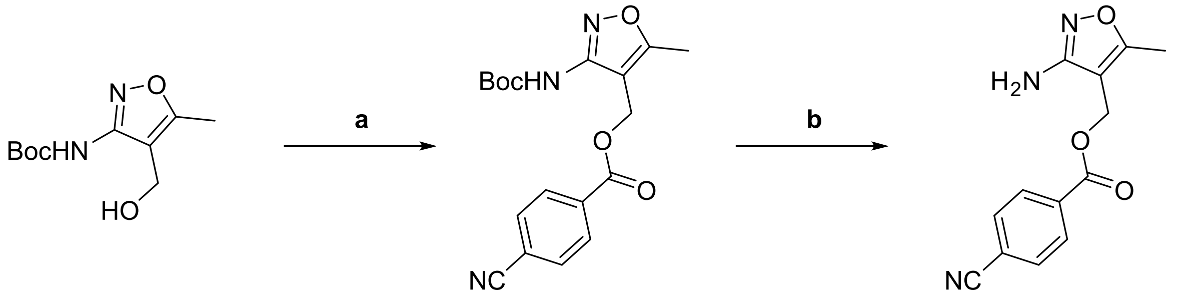

**S73** **S74** **S75**

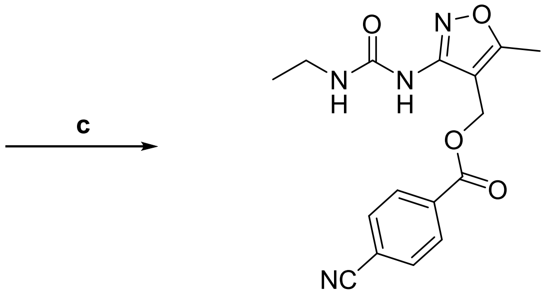

**4**

**(3-((*tert*-butoxycarbonyl)amino)-5-methylisoxazol-4-yl)methyl 4-cyanobenzoate, S74**

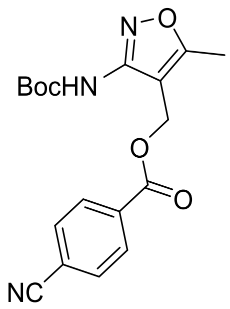

(a) **S73** (216 mg, 0.944 mmol), 4-cyanobenzoic acid (125 mg, 0.850 mmol) and DMAP (20.8 mg, 0.170 mmol) were dissolved in anhydrous DMF (8.9 mL) and cooled to 0 °C. EDCI (195 mg, 1.02 mmol) was added, and the reaction was stirred at r.t. for 16 h. The reaction was quenched with water and extracted 3 times with DCM. The combined organic phases were washed with brine, dried using $Na_2SO_4$ and solvent removed *in vacuo*. The crude product was purified using flash column chromatography eluting with a gradient of 0-50% EtOAc in Cy. **S74** (153 mg, 0.427 mmol, 50%) was obtained as a white solid.

**$^1$H NMR (400 MHz, $CDCl_3$) δ** 8.14-8.10 (m, 2H), 7.77-7.73 (m, 2H), 5.24 (s, 2H), 2.47 (s, 3H), 1.50 (s, 9H); **$^{13}$C NMR (101 MHz, $CDCl_3$) δ** 170.2, 165.5, 157.0, 151.7, 133.5, 132.4, 130.4, 117.9, 117.0, 104.8, 82.2, 56.8, 28.3, 11.8; **LR-ESI-MS:** $C_{18}H_{20}N_3O_5$ $[M+H]^+$ *m/z* found 358.39 cald 358.14; **HR-ESI-MS:** $C_{18}H_{19}N_3O_5Na$ $[M+Na]^+$ *m/z* found 380.1024 cald 380.1217.

**(3-amino-5-methylisoxazol-4-yl)methyl 4-cyanobenzoate, S75**

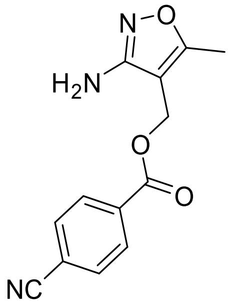

(b) **S74** (78.2 mg, 0.219 mmol) was dissolved in DCM (1.4 mL) and TFA (1.1 mL, 14.7 mmol) added dropwise. The reaction was stirred at r. t. for 1 h. The volatile reagents were removed *in vacuo*. The crude oil was redissolved in minimal DCM and washed with saturated aq. $Na_2CO_3$. The organic phases were combined, dried using $Na_2SO_4$, and the solvent removed *in vacuo*. **S75** (51.3 mg, 0.199 mmol, 91%) was obtained as a white gum.

**$^1$H NMR (400 MHz, MeOD) δ** 8.21-8.12 (m, 2H), 7.92-7.82 (m, 2H), 5.19 (s, 2H), 2.38 (s, 3H); **$^{13}$C NMR (101 MHz, MeOD) δ** 170.2, 166.6, 165.0, 135.2, 133.6, 131.3, 118.9, 117.8, 107.7, 57.1, 11.2; **LR-ESI-MS:** $C_{13}H_{12}N_3O_3$ $[M+H]^+$ *m/z* found 258.15 cald 258.08; **HR-ESI-MS:** $C_{13}H_{11}N_3O_3Na$ $[M+Na]^+$ *m/z* found 280.0543 cald 280.0693.

**(3-(3-ethylureido)-5-methylisoxazol-4-yl)methyl 4-cyanobenzoate, 4**

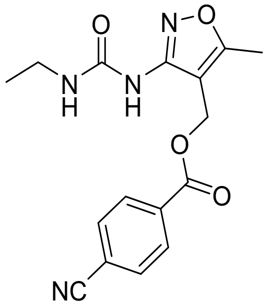

(c) **S75** (51.3 mg, 0.199 mmol) was dissolved in anhydrous THF (5 mL) and DMA (1 mL) and cooled to 0 °C. Triphosgene (29.6 mg, 0.100 mmol) was dissolved in anhydrous THF (0.5 mL) and added dropwise. The reaction was stirred for 5 mins before anhydrous TEA (83.0 µL, 0.598 mmol) was added dropwise. The reaction was stirred at 0 °C for 30 mins. Ethylamine (150 µL, 0.299 mmol) was added dropwise. The reaction was allowed to warm to r.t. and stirred for 2 h. The reaction was poured into ice water and diluted with DCM. The organic phase was extracted, washed with brine, dried using $Na_2SO_4$ and the solvent was removed *in vacuo*. MeOH was added, the precipitated solid collected by filtration, washed with ice cold MeOH, and dried *in vacuo*. **4** (24.1 mg, 73.4 µmol, 37%) was obtained as a white solid.

**$^1$H NMR (400 MHz, DMSO-*d*$_6$) δ** 9.20 (s, 1H), 8.10-8.03 (m, 2H), 8.02-7.97 (m, 2H), 7.08 (t, J = 5.6 Hz, 1H), 5.21 (s, 2H), 3.14 (qd, J = 7.2, 5.6 Hz, 2H), 2.43 (s, 3H), 1.03 (t, J = 7.2 Hz, 3H); **$^{13}$C NMR (101 MHz, DMSO-*d*$_6$) δ** 169.1, 164.4, 158.1, 153.7, 133.7, 132.8, 129.8, 118.0, 115.5, 103.2, 56.3, 34.4, 15.6, 11.0; **LR-ESI-MS:** $C_{16}H_{17}N_4O_4$ $[M+H]^+$ *m/z* found 329.40 cald 329.12; **HR-ESI-MS:** $C_{16}H_{16}N_4O_4Na$ $[M+Na]^+$ *m/z* found 351.0878 cald 351.1064.

**Synthesis of (3-(3-ethylureido)-5-methylisoxazol-4-yl)methyl 4-cyanobenzoate, 5**

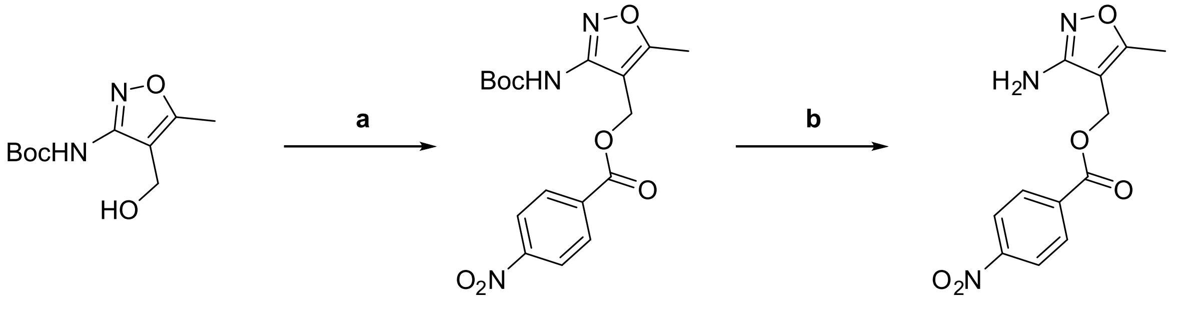

**S73** **S76** **S77**

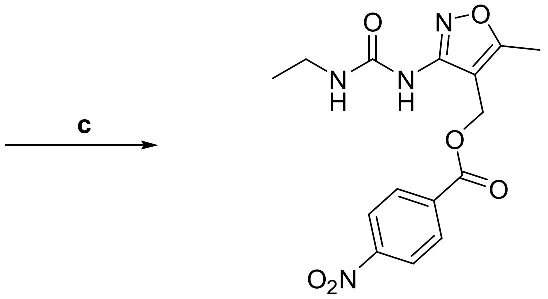

**5**

**(3-((*tert*-butoxycarbonyl)amino)-5-methylisoxazol-4-yl)methyl (*E*)-3-(4-nitrophenyl)acrylate, S76**

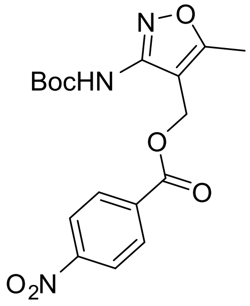

(a) **S73** (216 mg, 0.944 mmol), 3-(4-nitrophenyl)prop-2-enoic acid (164 mg, 0.850 mmol) and DMAP (30.8 mg, 0.252 mmol) were dissolved in anhydrous DMF (8.9 mL) and cooled to 0 °C. EDCI (195 mg, 1.02 mmol) was added and the reaction was stirred at r.t. for 16 h. The reaction was quenched with water and extracted 3 times with EtOAc. The organic phase was washed with 5% LiCl, brine, dried using $Na_2SO_4$ and the solvent removed *in vacuo*. The crude product was purified using flash column chromatography eluting with a gradient of 0-50% EtOAc in Cy. **S76** (241 mg, 0.597 mmol, 70%) was obtained as a white solid.

**$^1$H NMR (400 MHz, DMSO-*d*$_6$) δ** 9.70 (s, 1H), 8.27-8.22 (m, 2H), 8.03-7.99 (m, 2H), 7.75 (d, J = 16.1 Hz, 1H), 6.86 (d, J = 16.0 Hz, 1H), 5.10 (s, 2H), 2.46 (s, 3H), 1.42 (s, 9H); **$^{13}$C NMR (101 MHz, DMSO-*d*$_6$) δ** 169.4, 165.5, 157.3, 152.3, 148.1, 142.2, 140.4, 129.5, 124.0, 122.1, 105.6, 80.1, 55.7, 27.9, 11.2; **LR-ESI-MS:** $C_{19}H_{20}N_3O_7$ [M-H]$^-$ *m/z* found 402.20 cald 402.14; **HR-ESI-MS:** $C_{19}H_{21}N_3O_7Na$ [M+Na]$^+$ *m/z* found 426.1054 cald 426.1272.

**(3-amino-5-methylisoxazol-4-yl)methyl (*E*)-3-(4-nitrophenyl)acrylate, S77**

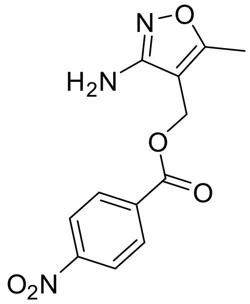

(b) **S76** (80.5 mg, 0.200 mmol) was dissolved in DCM (1.2 mL) and TFA (1.0 mL, 13.4 mmol) added dropwise. The reaction was stirred at r.t. for 1.5 h. The volatile reagents were removed *in vacuo*. The crude oil was redissolved in minimal DCM and washed with saturated aq. $Na_2CO_3$. The organic phases were combined, dried using $Na_2SO_4$, and the solvent removed *in vacuo*. **S77** (60.4 mg, 0.199 mmol, 100%) was obtained as a light brown solid.

**$^1$H NMR (400 MHz, DMSO-*d*$_6$) δ** 8.24 (d, J = 8.9 Hz, 2H), 8.02 (d, J = 8.9 Hz, 2H), 7.77 (d, J = 16.1 Hz, 1H), 6.87 (d, J = 16.1 Hz, 1H), 5.60 (s, 2H), 4.99 (s, 2H), 2.29 (s, 3H); **$^{13}$C NMR (101 MHz, DMSO-*d*$_6$) δ** 167.5, 165.9, 163.1, 148.1, 142.1, 140.5, 129.5, 124.0, 122.3, 101.6, 55.2, 11.0; **LR-ESI-MS:** $C_{14}H_{14}N_3O_5$ [M+H]$^+$ *m/z* found 304.34 cald 304.09; **HR-ESI-MS:** $C_{14}H_{14}N_3O_5$ [M+H]$^+$ *m/z* found 304.0762 cald 304.0928.

**(3-(3-ethylureido)-5-methylisoxazol-4-yl)methyl (*E*)-3-(4-nitrophenyl)acrylate, 5**

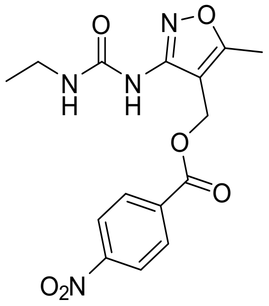

(c) **S77** (55.4 mg, 0.183 mmol) was dissolved in anhydrous THF (5 mL) and DMA (1 mL) and cooled to 0 °C. Triphosgene (27.1 mg, 0.0913 mmol) was dissolved in anhydrous THF (0.5 mL) and added dropwise. The reaction was stirred for 5 mins at 0 °C before anhydrous TEA (76.0 µL, 0.548 mmol) was added dropwise. The reaction was stirred at 0 °C for 30 mins. Ethylamine (137 µL, 0.274 mmol) was added dropwise. The reaction was stirred at r.t. for 2 h. The reaction was poured onto ice and extracted 3 times with DCM. The organic phase was washed with brine, dried using $Na_2SO_4$ and solvent removed *in vacuo*. The crude product was purified using preparative HPLC and subsequent lyophilisation of fractions. **5** (5.2 mg, 13.9 µmol, 8%) was obtained as a white solid.

**$^1$H NMR (400 MHz, DMSO-*d*$_6$) δ** 9.13 (s, 1H), 8.27-8.19 (m, 2H), 8.06-7.98 (m, 2H), 7.76 (d, J = 16.1 Hz, 1H), 7.09 (t, J = 5.6 Hz, 1H), 6.87 (d, J = 16.1 Hz, 1H), 5.10 (s, 2H), 3.16 (qd, J = 7.2, 5.6 Hz, 2H), 2.41 (s, 3H), 1.06 (t, J = 7.2 Hz, 3H); **$^{13}$C NMR (101 MHz, DMSO-*d*$_6$) δ** 168.9, 165.5, 158.0, 153.5, 148.0, 142.0, 140.3, 129.4, 123.8, 122.1, 103.3, 55.0, 34.3, 15.1, 10.8; **LR-ESI-MS:** $C_{17}H_{19}N_4O_6$ $[M+H]^+$ *m/z* found 375.36 cald 375.13; **HR-ESI-MS:** $C_{17}H_{18}N_4O_6Na$ $[M+Na]^+$ *m/z* found 397.0912 cald 397.1119.

### NMR spectra

## S37

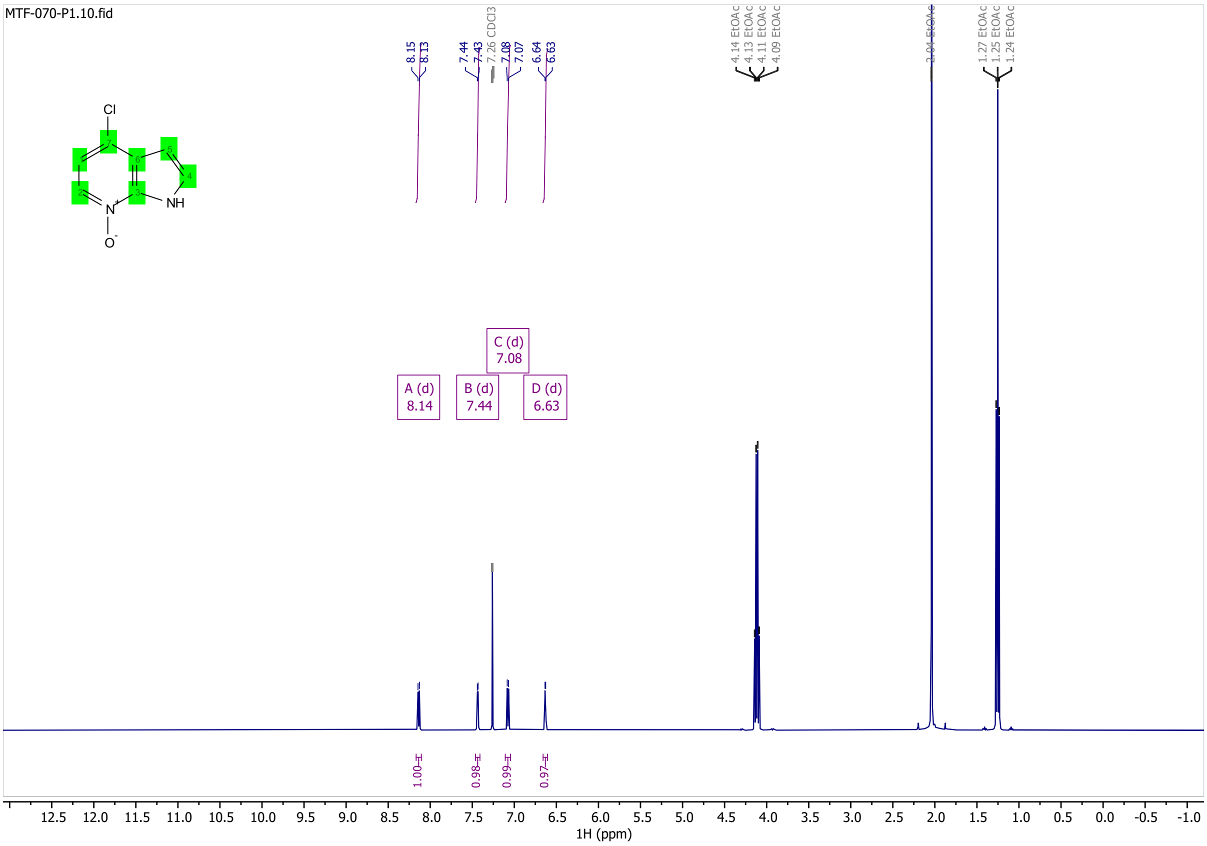

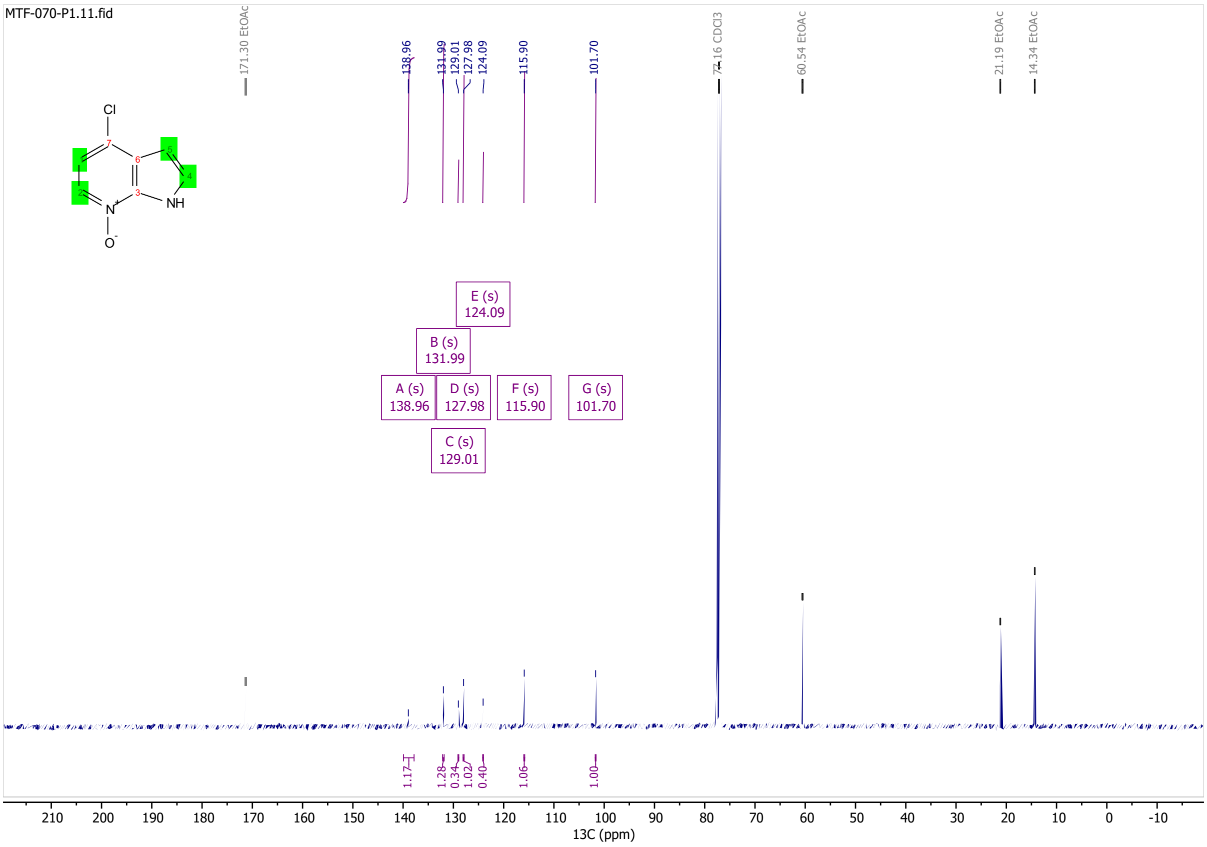

# S38

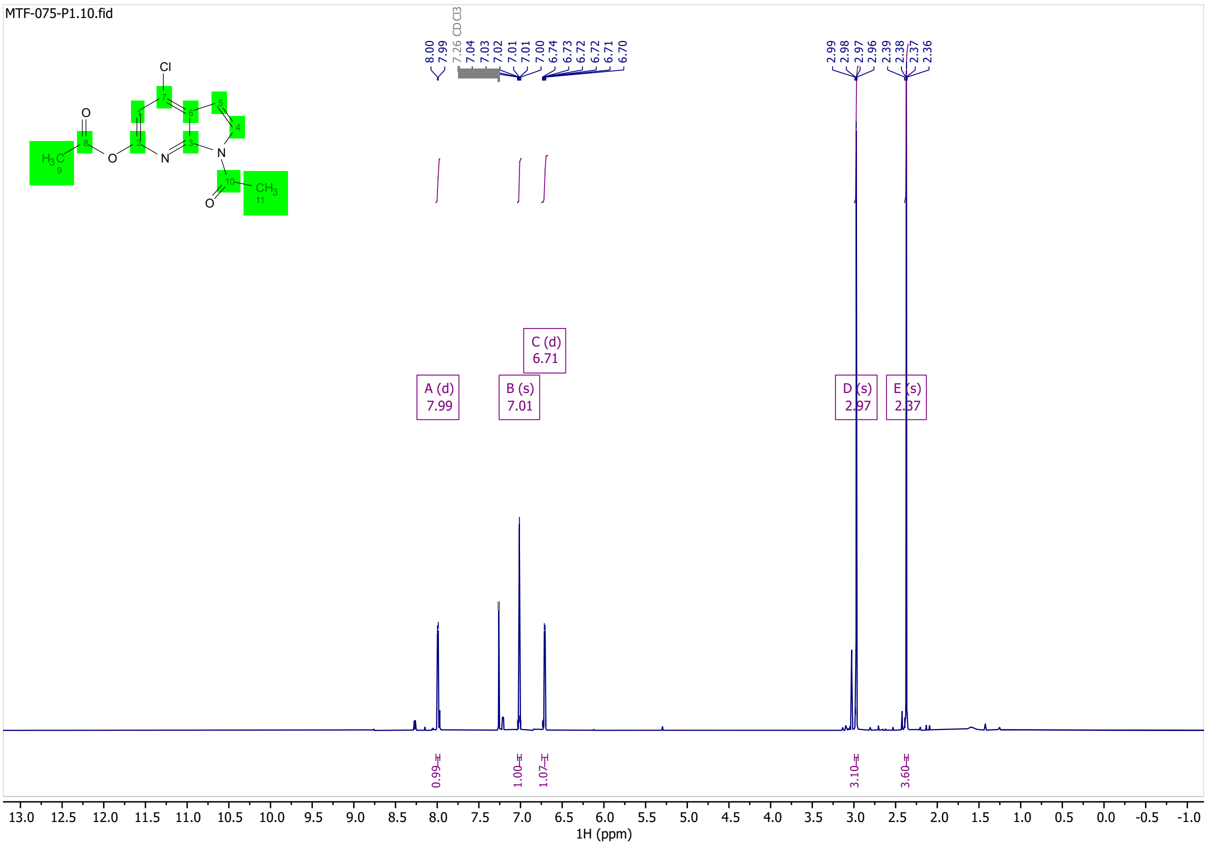
MTF-075-P1.10.fid
8.00
7.99
7.26 CDCl3
7.04
7.03
7.02
7.01
7.01
7.00
6.74
6.73
6.72
6.72
6.71
6.70
2.99
2.98
2.97
2.96
2.39
2.38
2.37
2.36
A (d)
7.99
B (s)
7.01
C (d)
6.71
D (s)
2.97
E (s)
2.37
0.99
1.00
1.07
3.10
3.60
1H (ppm)

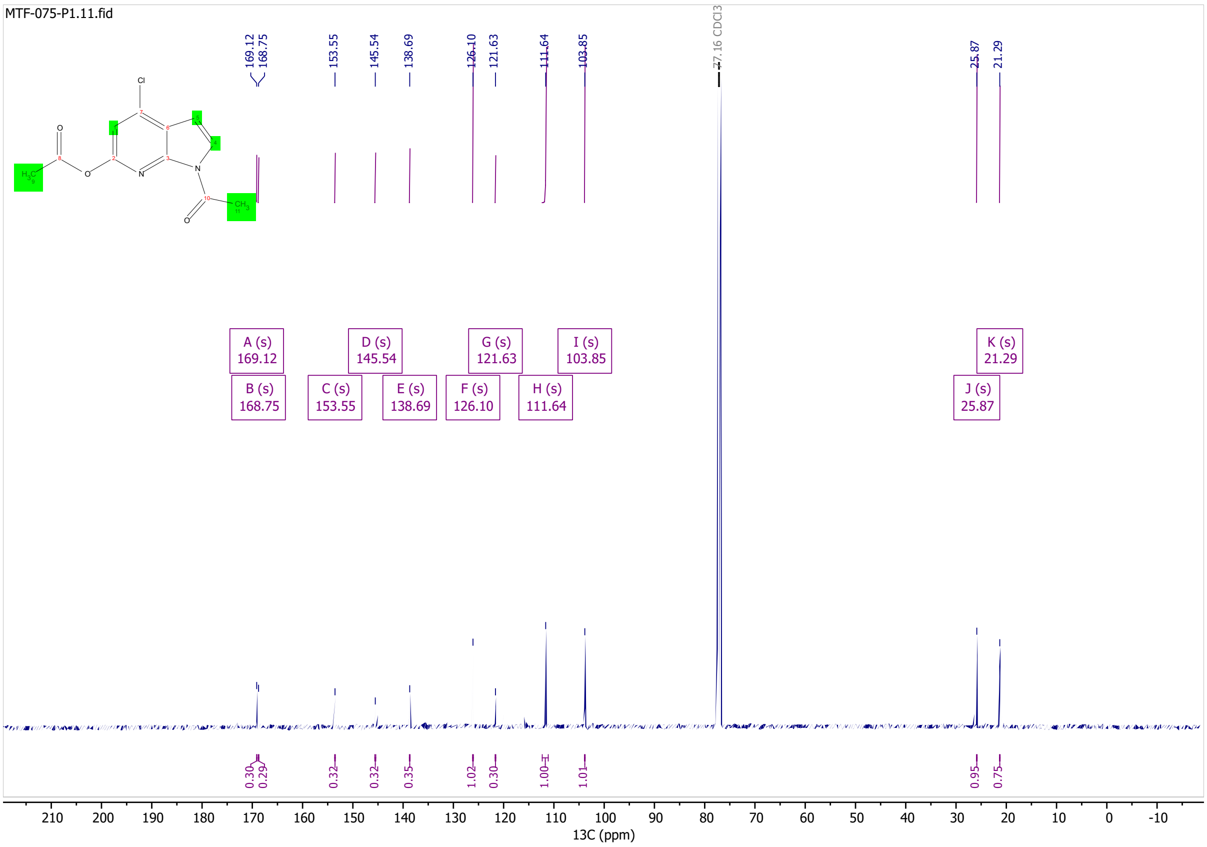
MTF-075-P1.11.fid
169.12
168.75
153.55
145.54
138.69
126.10
121.63
111.64
103.85
77.16 CDCl3
25.87
21.29
A (s)
169.12
B (s)
168.75
C (s)
153.55
D (s)
145.54
E (s)
138.69
F (s)
126.10
G (s)
121.63
H (s)
111.64
I (s)
103.85
J (s)
25.87
K (s)
21.29
13C (ppm)

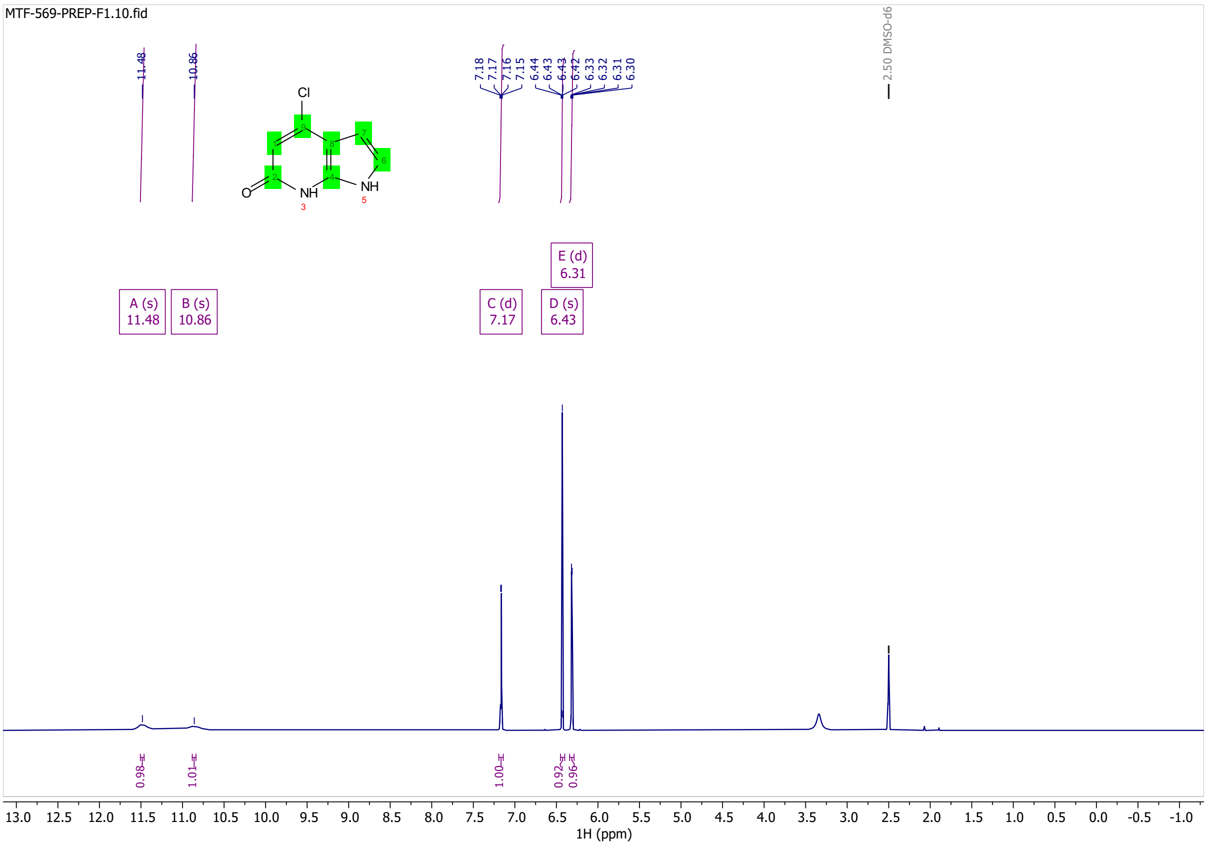
MTF-569-PREP-F1.10.fid
11.48
10.86
7.18
7.17
7.16
7.15
6.44
6.43
6.42
6.33
6.32
6.31
6.30
2.50 DMSO-d6
Cl
NH
NH
O
A (s)
11.48
B (s)
10.86
C (d)
7.17
D (s)
6.43
E (d)
6.31
0.98
1.01
1.00
0.92
0.96
1H (ppm)

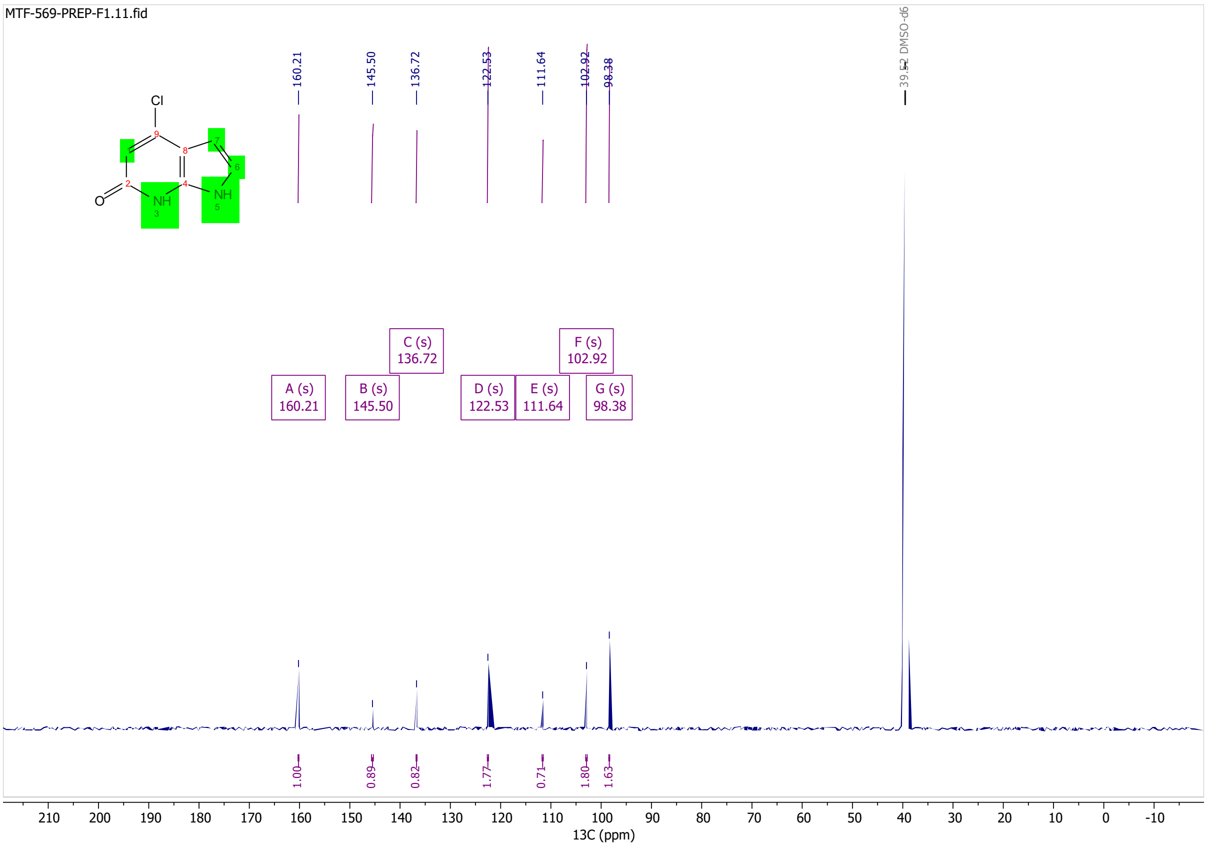
MTF-569-PREP-F1.11.fid
160.21
145.50
136.72
122.53
111.64
102.92
98.38
39.52 DMSO-d6
Cl
NH
NH
O
A (s)
160.21
B (s)
145.50
C (s)
136.72
D (s)
122.53
E (s)
111.64
F (s)
102.92
G (s)
98.38
1.00
0.89
0.82
1.77
0.71
1.80
1.63
13C (ppm)

# S78

MTF-077-P1-C3-F2-6.10.fid

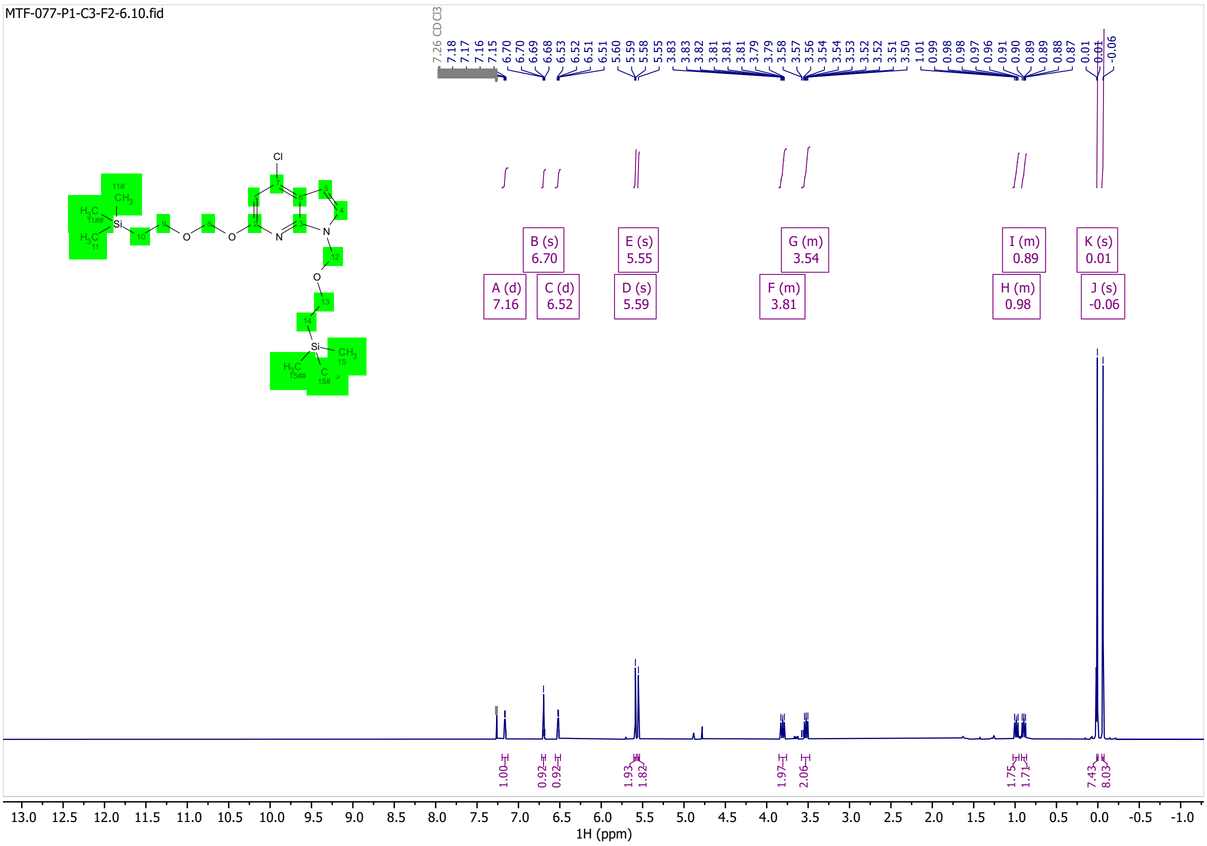

MTF-077-P1-C3-F2-6.11.fid

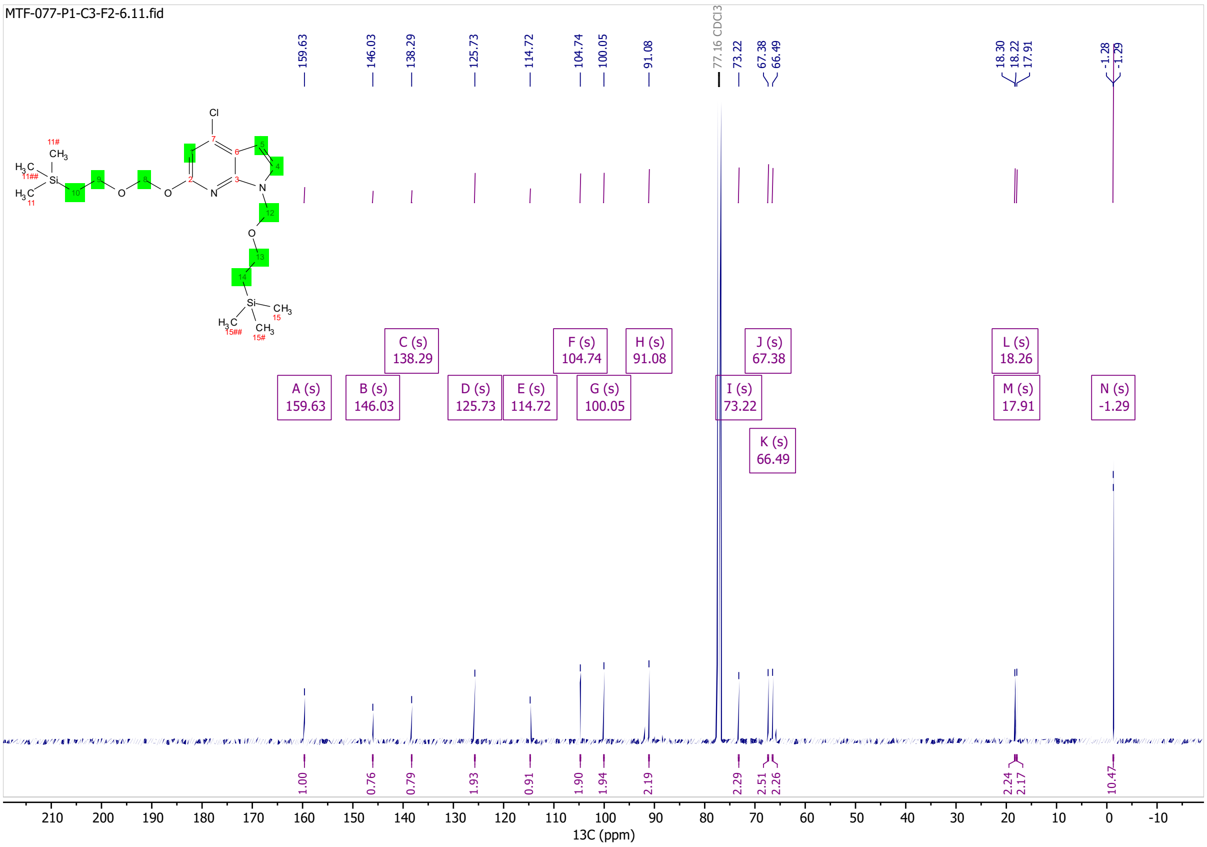

S79

MTF-110-C1-F1.10.fid

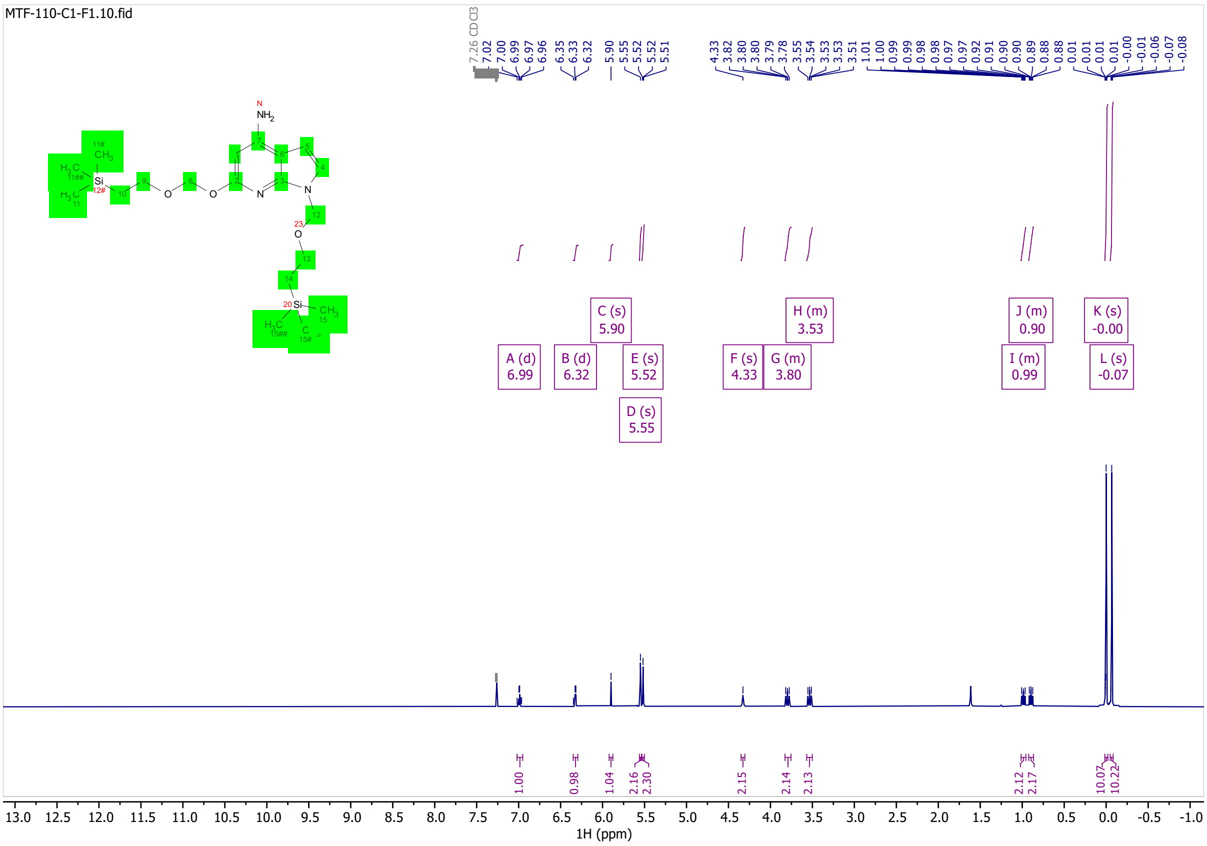

MTF-110-C1-F1.11.fid

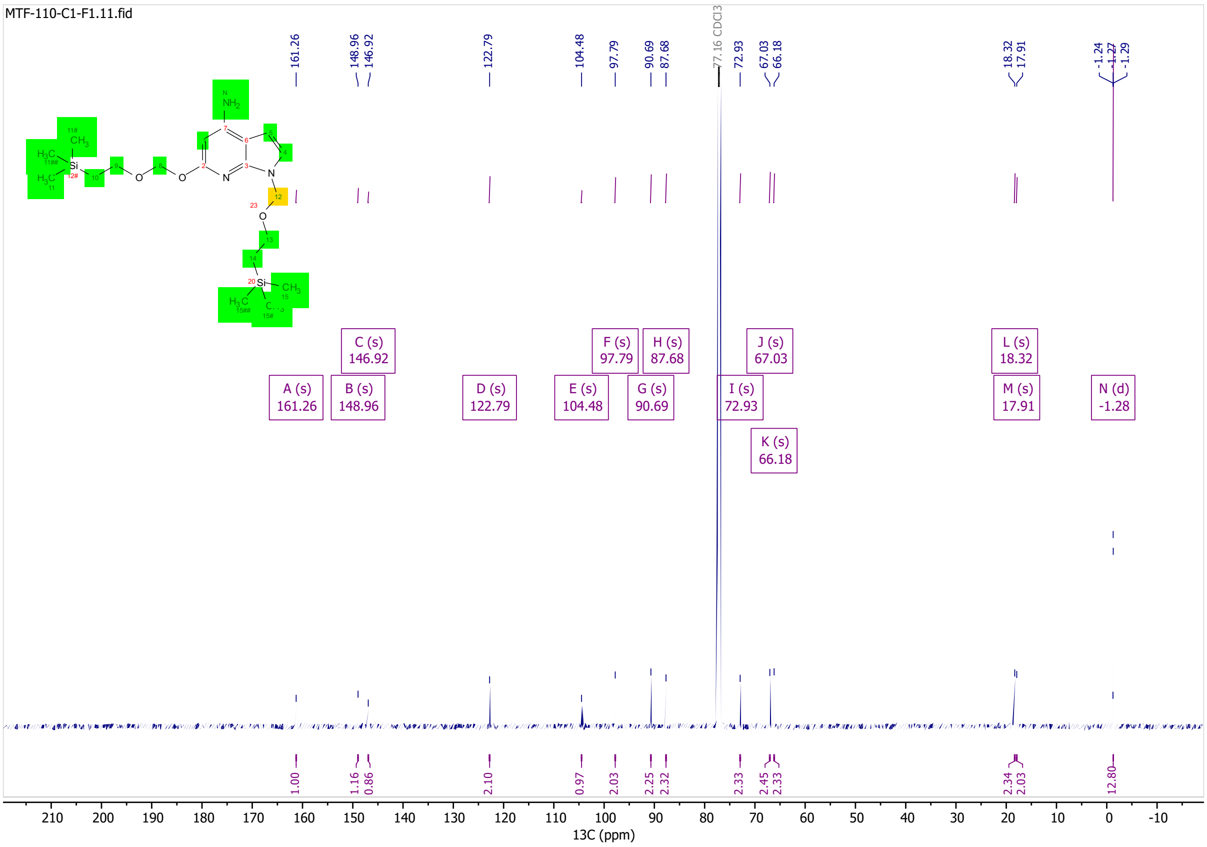

# S80

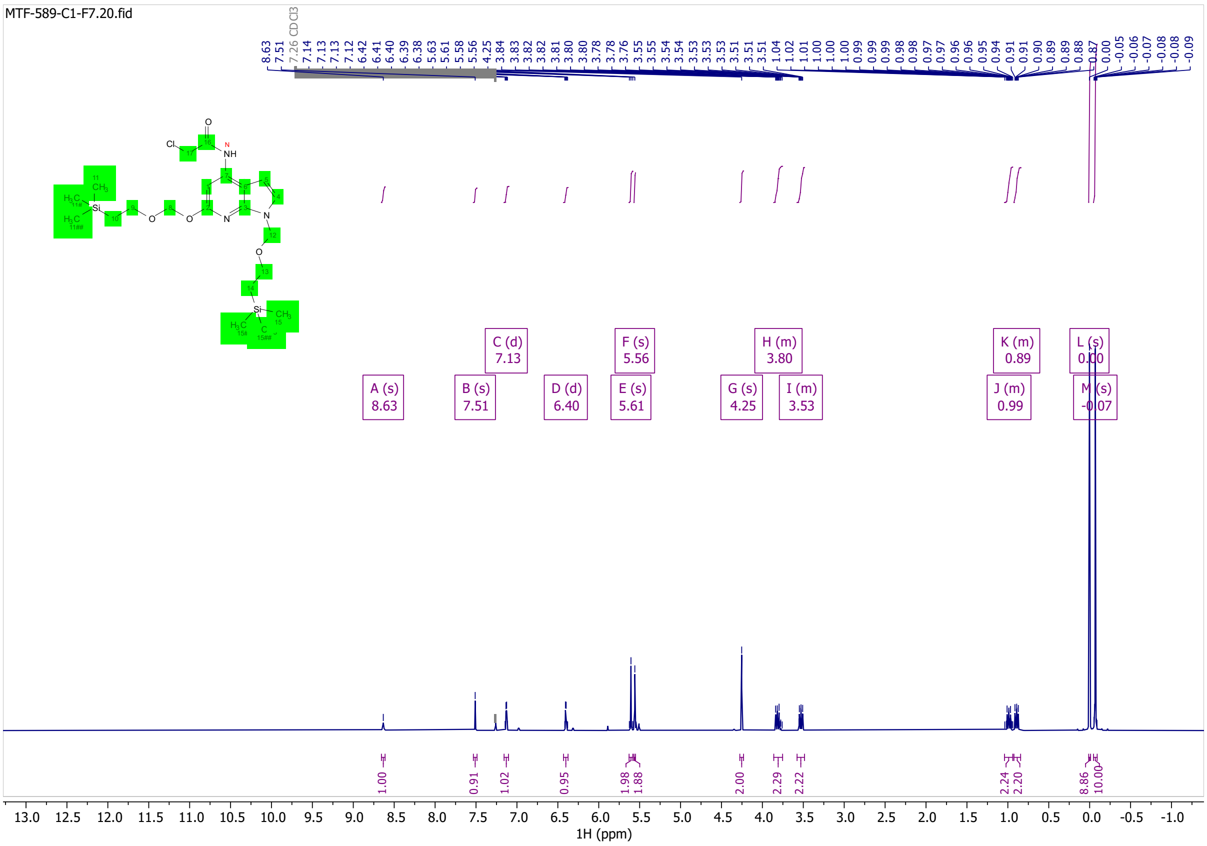
MTF-589-C1-F7.20.fid
A (s) 8.63
B (s) 7.51
C (d) 7.13
D (d) 6.40
E (s) 5.61
F (s) 5.56
G (s) 4.25
H (m) 3.80
I (m) 3.53
J (m) 0.99
K (m) 0.89
L (s) 0.00
M (s) -0.07
1H (ppm)

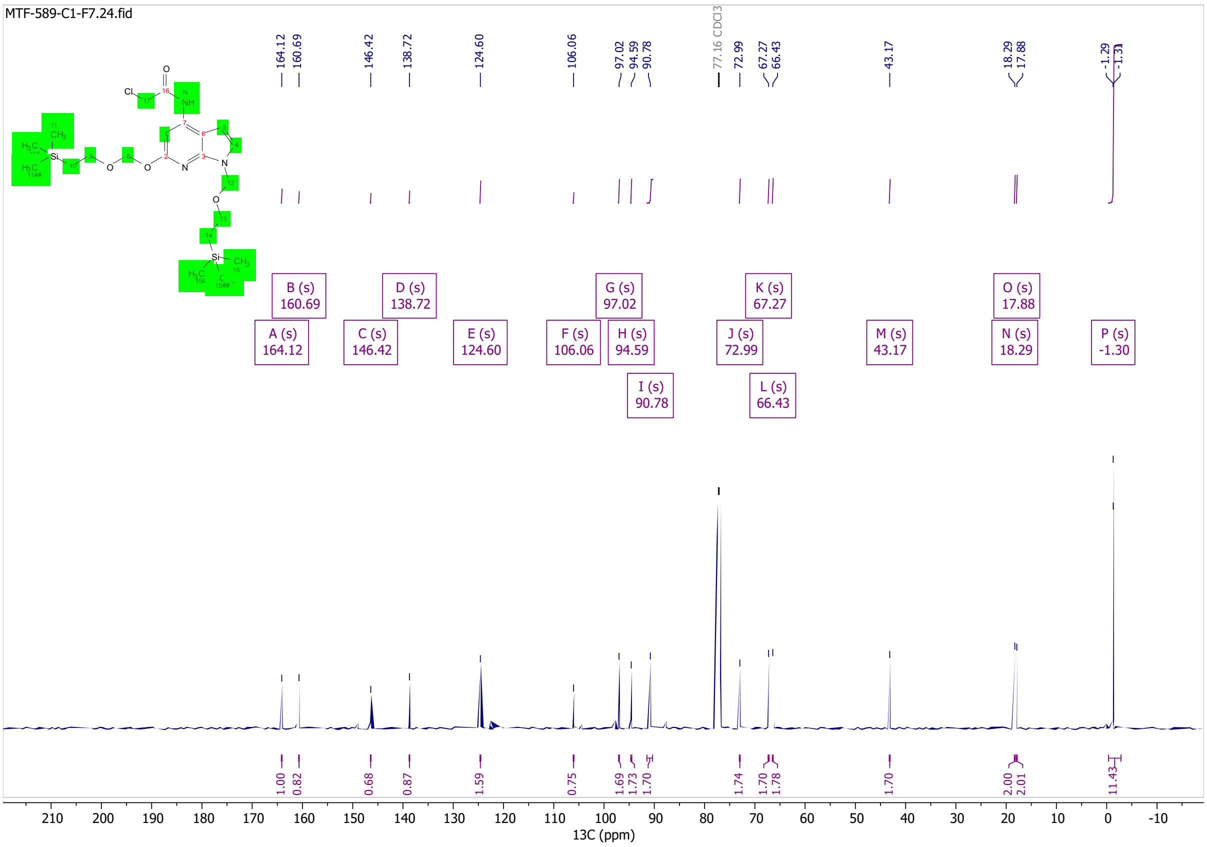
MTF-589-C1-F7.24.fid
A (s) 164.12
B (s) 160.69
C (s) 146.42
D (s) 138.72
E (s) 124.60
F (s) 106.06
G (s) 97.02
H (s) 94.59
I (s) 90.78
J (s) 72.99
K (s) 67.27
L (s) 66.43
M (s) 43.17
N (s) 18.29
O (s) 17.88
P (s) -1.30
13C (ppm)

# RS-009

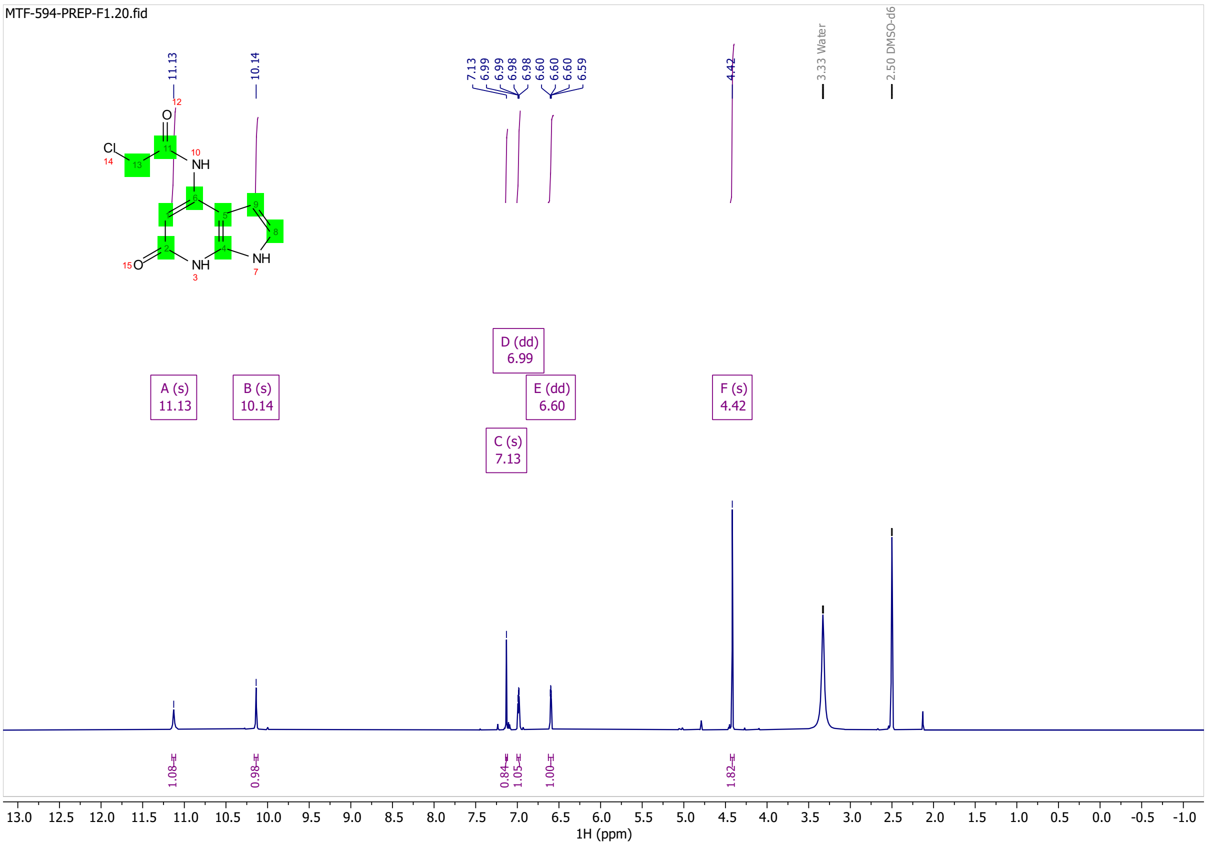

MTF-594-PREP-F1.20.fid
11.13
10.14
7.13
6.99
6.99
6.98
6.98
6.60
6.60
6.59
4.42
3.33 Water
2.50 DMSO-d6
A (s) 11.13
B (s) 10.14
C (s) 7.13
D (dd) 6.99
E (dd) 6.60
F (s) 4.42
1.08
0.98
0.84
1.05
1.00
1.82
1H (ppm)

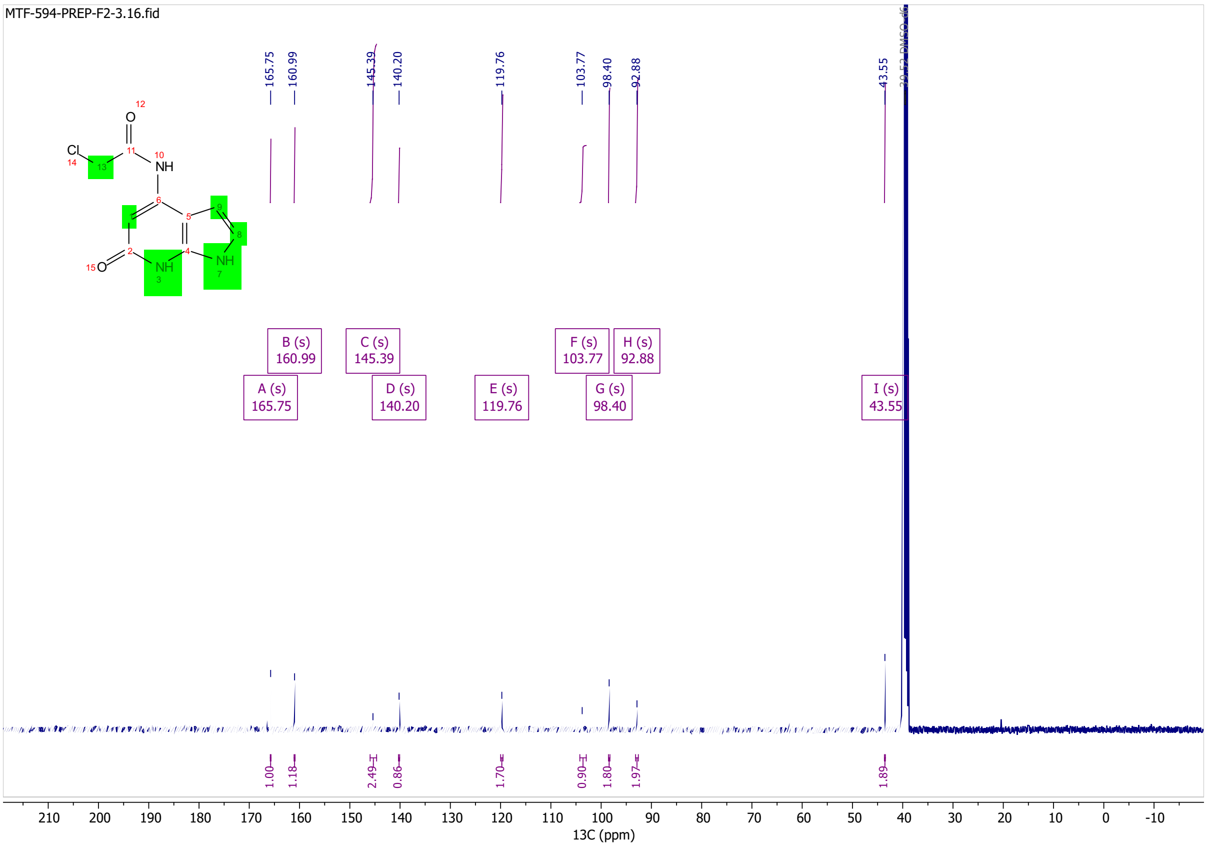

MTF-594-PREP-F2-3.16.fid
165.75
160.99
145.39
140.20
119.76
103.77
98.40
92.88
43.55
A (s) 165.75
B (s) 160.99
C (s) 145.39
D (s) 140.20
E (s) 119.76
F (s) 103.77
G (s) 98.40
H (s) 92.88
I (s) 43.55
1.00
1.18
2.49
0.86
1.70
0.90
1.80
1.97
1.89
13C (ppm)

S41

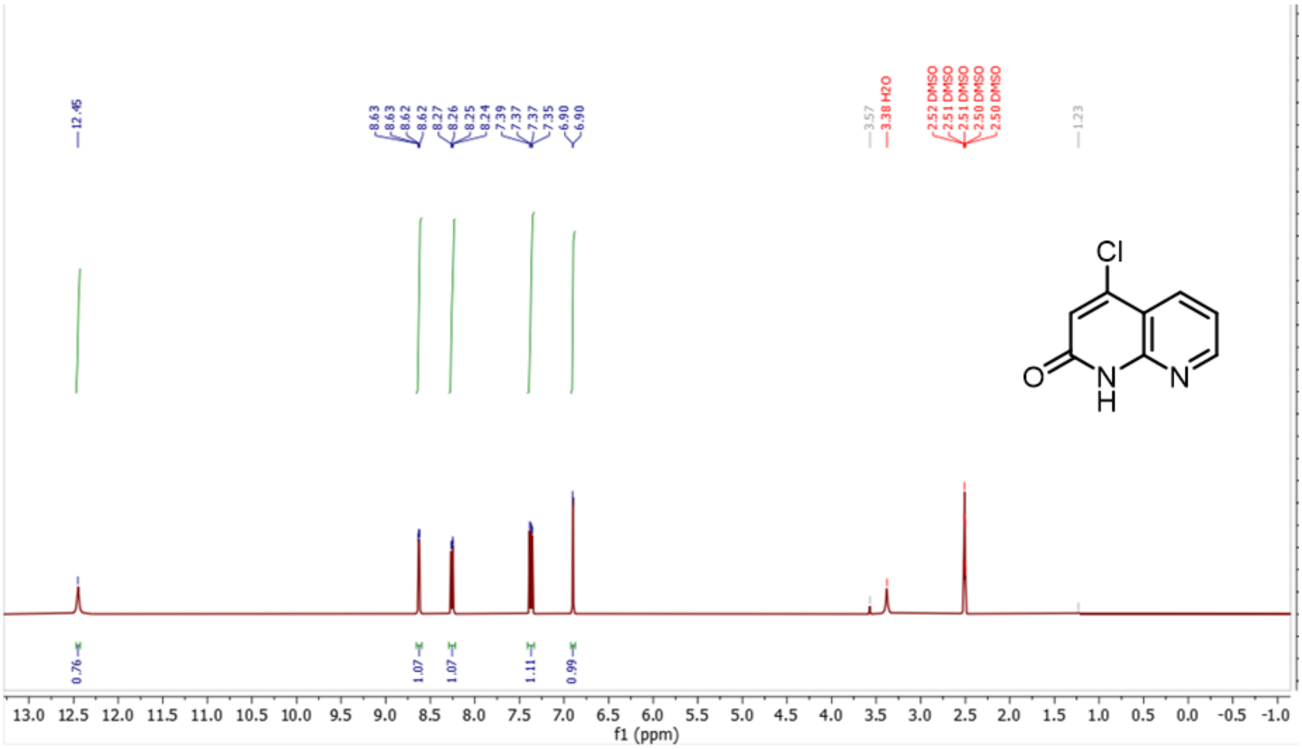
12.45
8.63
8.63
8.62
8.62
8.27
8.26
8.25
8.24
7.39
7.37
7.37
7.35
6.90
6.90
3.57
3.38 H2O
2.52 DMSO
2.51 DMSO
2.51 DMSO
2.50 DMSO
2.50 DMSO
1.23
Cl
O
N
H
N
0.76
1.07
1.07
1.11
0.99
f1 (ppm)

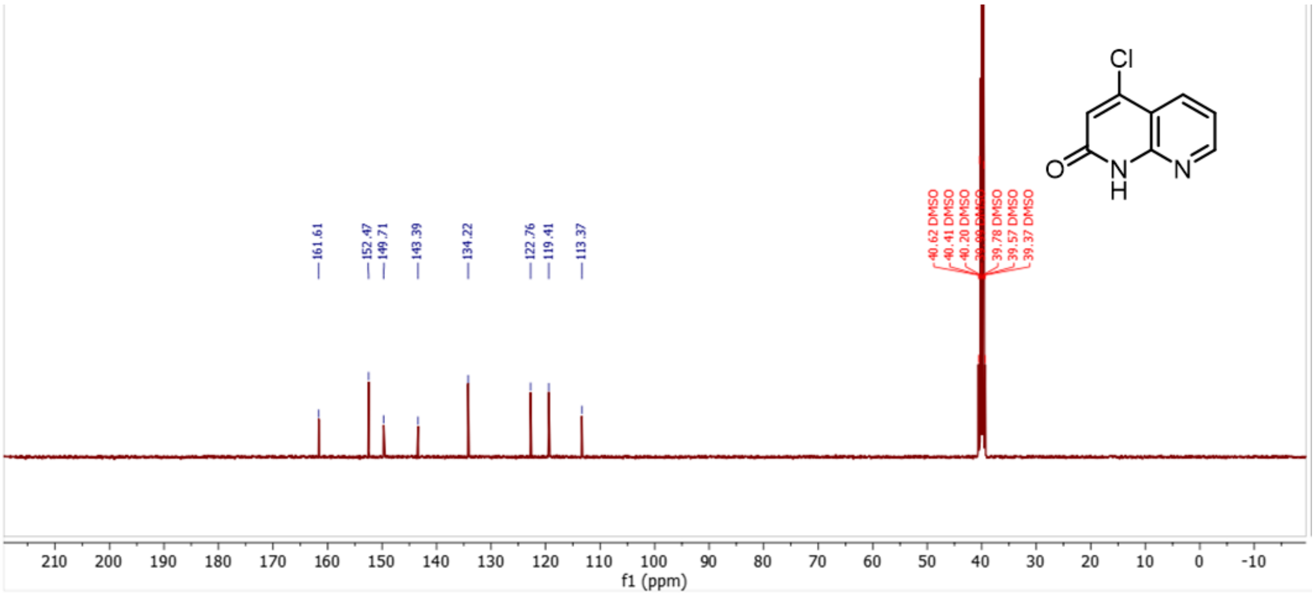
Cl
O
N
H
N
161.61
152.47
149.71
143.39
134.22
122.76
119.41
113.37
40.62 DMSO
40.41 DMSO
40.20 DMSO
39.78 DMSO
39.57 DMSO
39.37 DMSO
f1 (ppm)

# S42

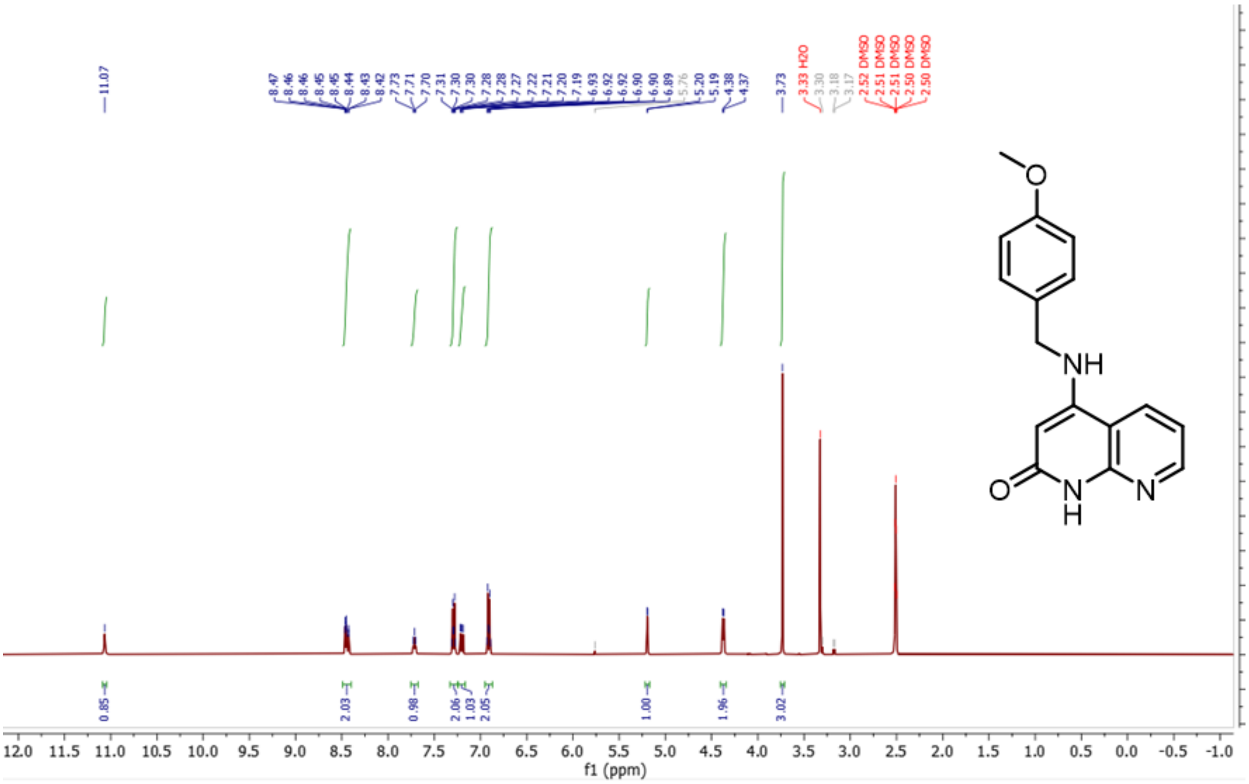

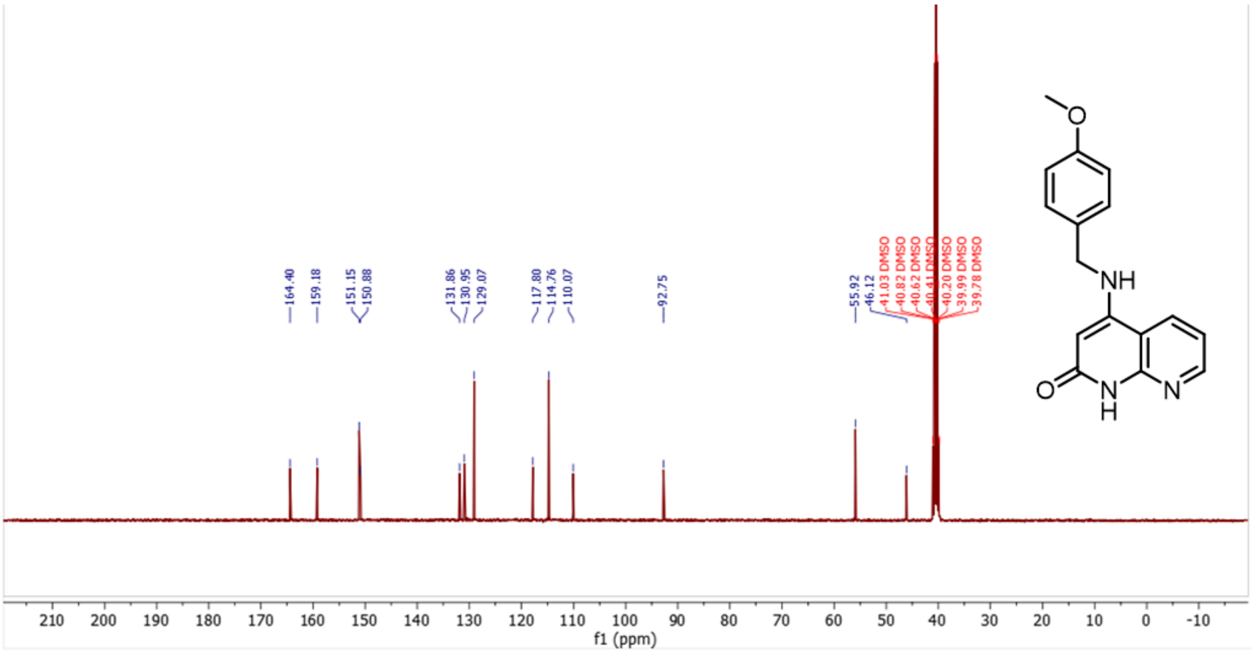

**S32**

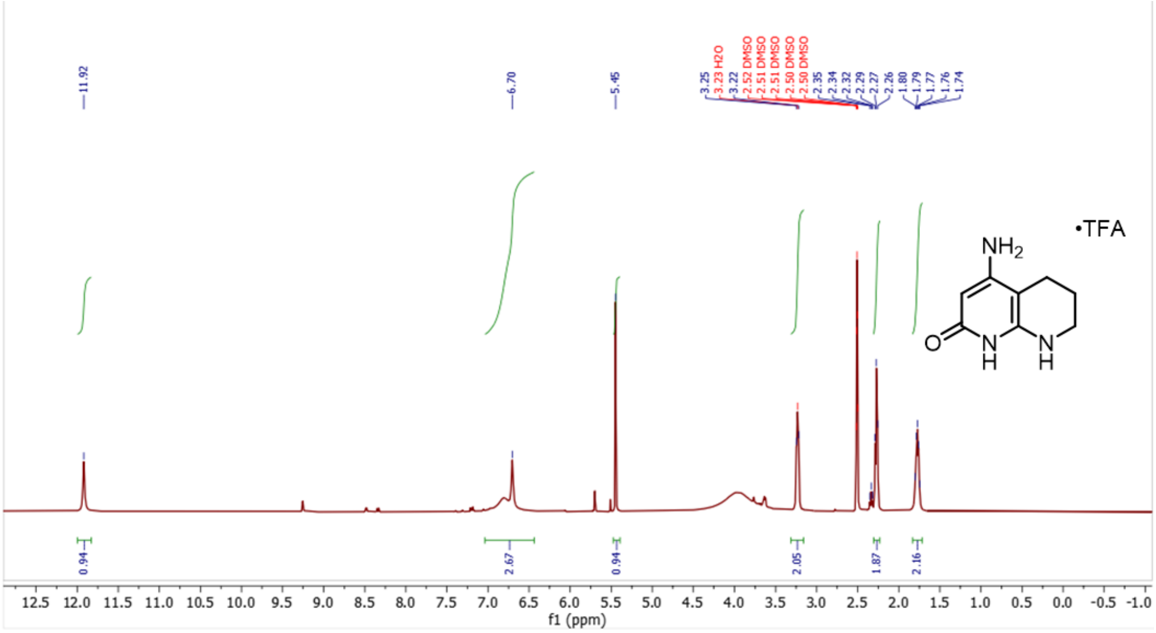
11.92
6.70
5.45
3.25
3.23 H2O
3.22
2.52 DMSO
2.51 DMSO
2.51 DMSO
2.50 DMSO
2.50 DMSO
2.35
2.34
2.32
2.29
2.27
2.26
1.80
1.79
1.77
1.76
1.74
•TFA
NH2
O
N H
N H
0.94
2.67
0.94
2.05
1.87
2.16
12.5 12.0 11.5 11.0 10.5 10.0 9.5 9.0 8.5 8.0 7.5 7.0 6.5 6.0 5.5 5.0 4.5 4.0 3.5 3.0 2.5 2.0 1.5 1.0 0.5 0.0 -0.5 -1.0
f1 (ppm)

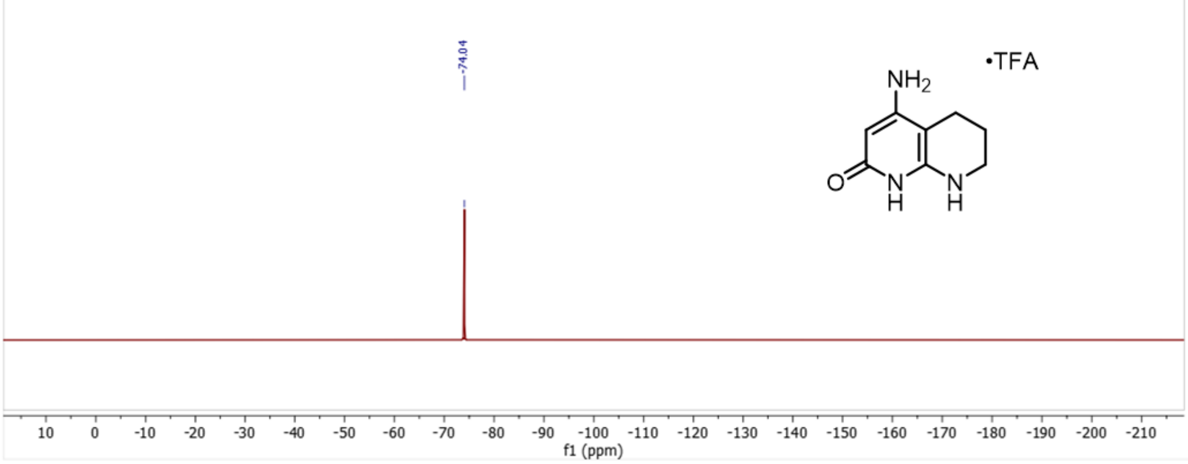
-74.04
•TFA
NH2
O
N H
N H
10 0 -10 -20 -30 -40 -50 -60 -70 -80 -90 -100 -110 -120 -130 -140 -150 -160 -170 -180 -190 -200 -210
f1 (ppm)

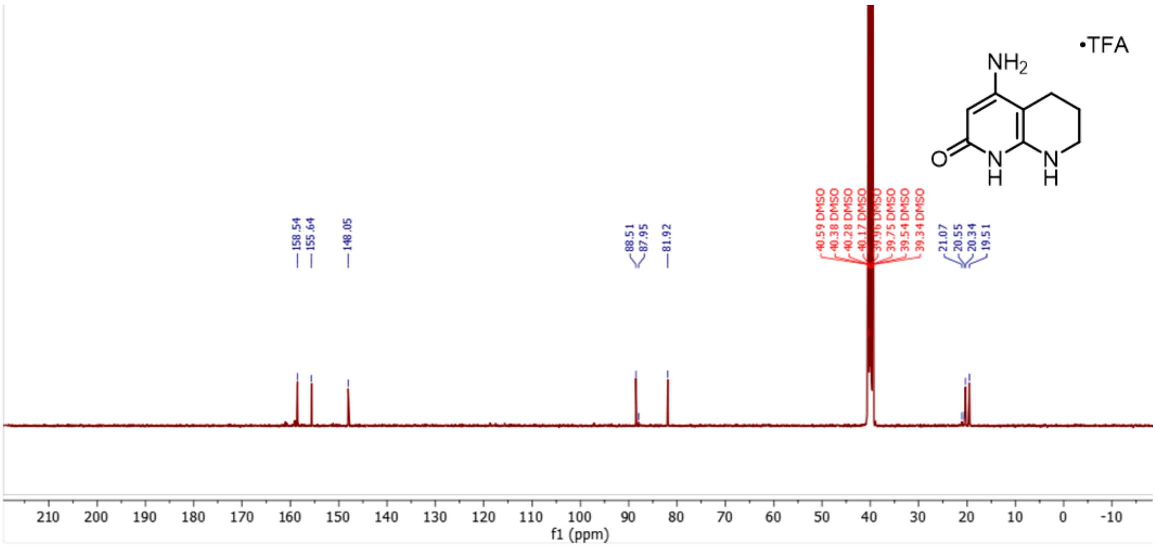
•TFA
NH2
O
N H
N H
158.54
155.64
148.05
88.51
87.95
81.92
40.59 DMSO
40.38 DMSO
40.28 DMSO
40.17 DMSO
39.96 DMSO
39.75 DMSO
39.54 DMSO
39.34 DMSO
21.07
20.55
20.34
19.51
210 200 190 180 170 160 150 140 130 120 110 100 90 80 70 60 50 40 30 20 10 0 -10
f1 (ppm)

**S43**

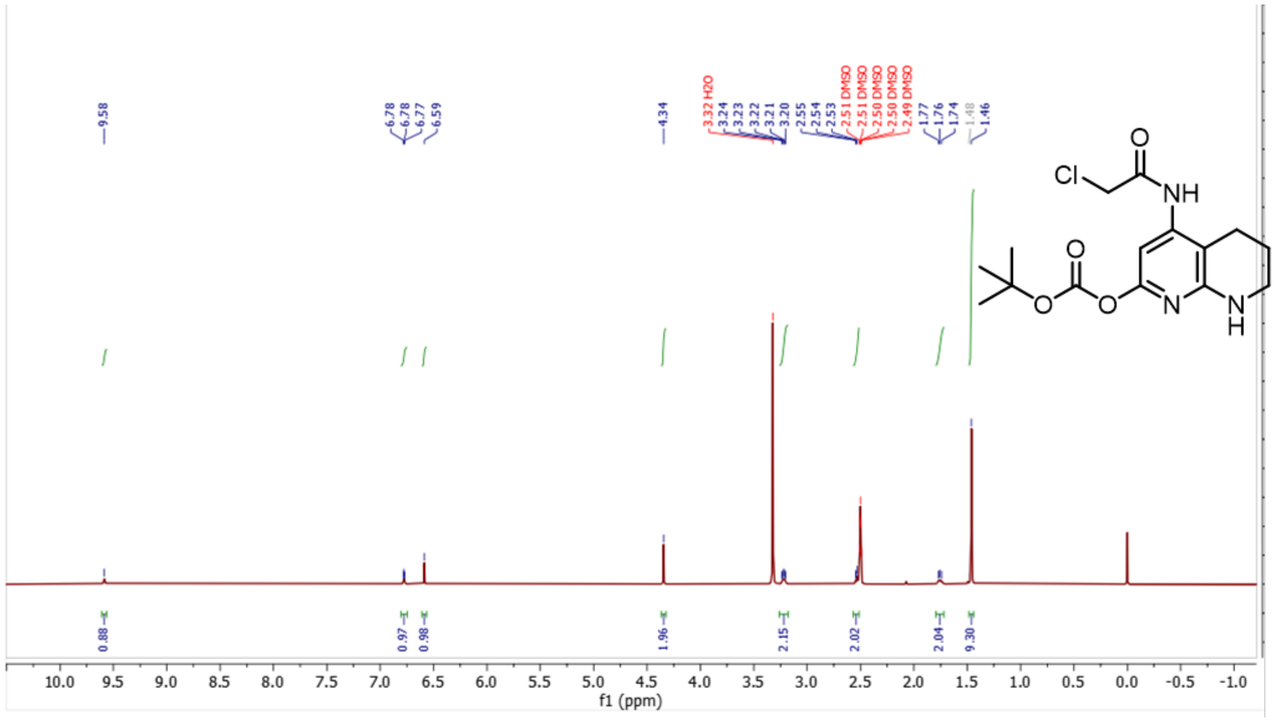
9.58
6.78
6.78
6.77
6.59
4.34
3.32 H2O
3.24
3.23
3.22
3.21
3.20
2.55
2.54
2.53
2.51 DMSO
2.51 DMSO
2.50 DMSO
2.50 DMSO
2.49 DMSO
1.77
1.76
1.74
1.48
1.46
0.88
0.97
0.98
1.96
2.15
2.02
2.04
9.30
f1 (ppm)

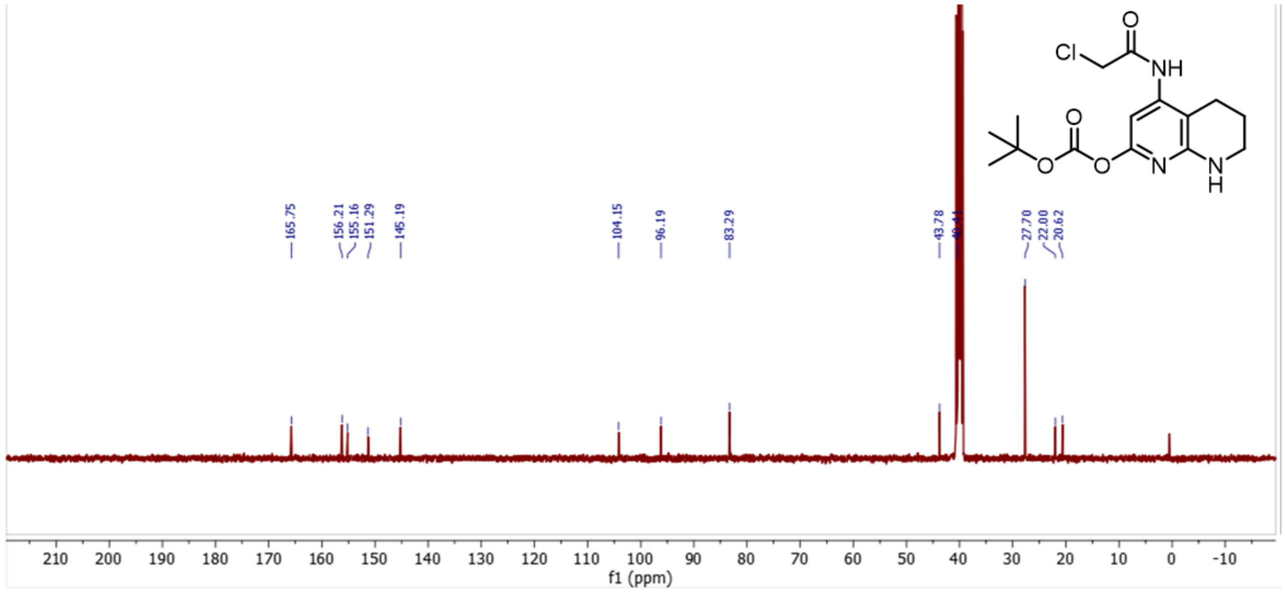
165.75
156.21
155.16
151.29
145.19
104.15
96.19
83.29
43.78
27.70
22.00
20.62
f1 (ppm)

**MC-278**

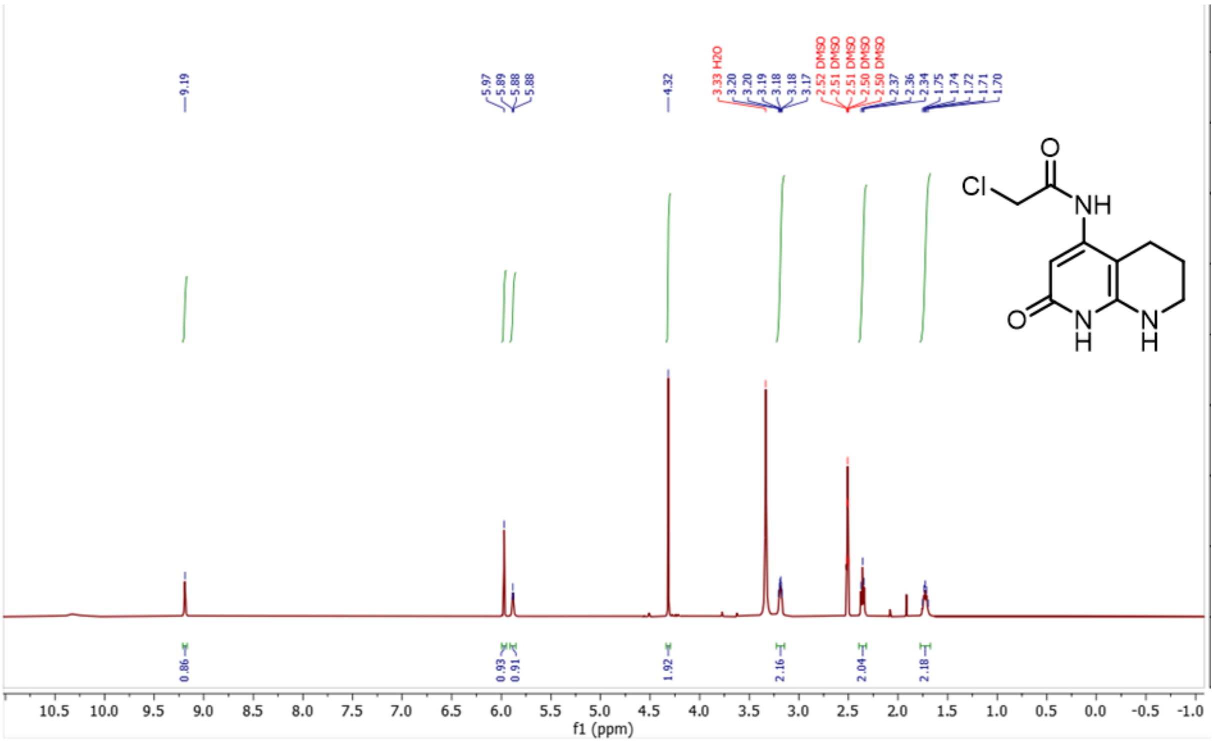

9.19
5.97
5.89
5.88
5.88
4.32
3.33 H2O
3.20
3.20
3.19
3.18
3.18
3.17
2.52 DMSO
2.51 DMSO
2.51 DMSO
2.50 DMSO
2.50 DMSO
2.37
2.36
2.34
1.75
1.74
1.72
1.71
1.70
Cl
O
NH
O
N
H
N
H
0.86
0.93
0.91
1.92
2.16
2.04
2.18
f1 (ppm)

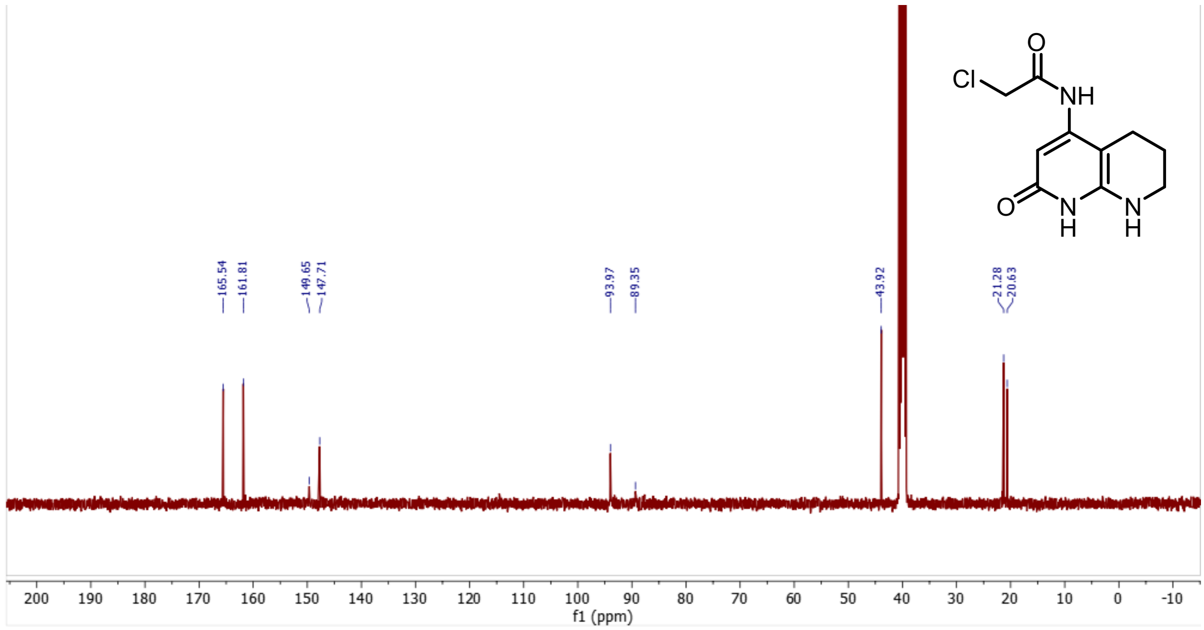

Cl
O
NH
O
N
H
N
H
165.54
161.81
149.65
147.71
93.97
89.35
43.92
21.28
20.63
f1 (ppm)

S45

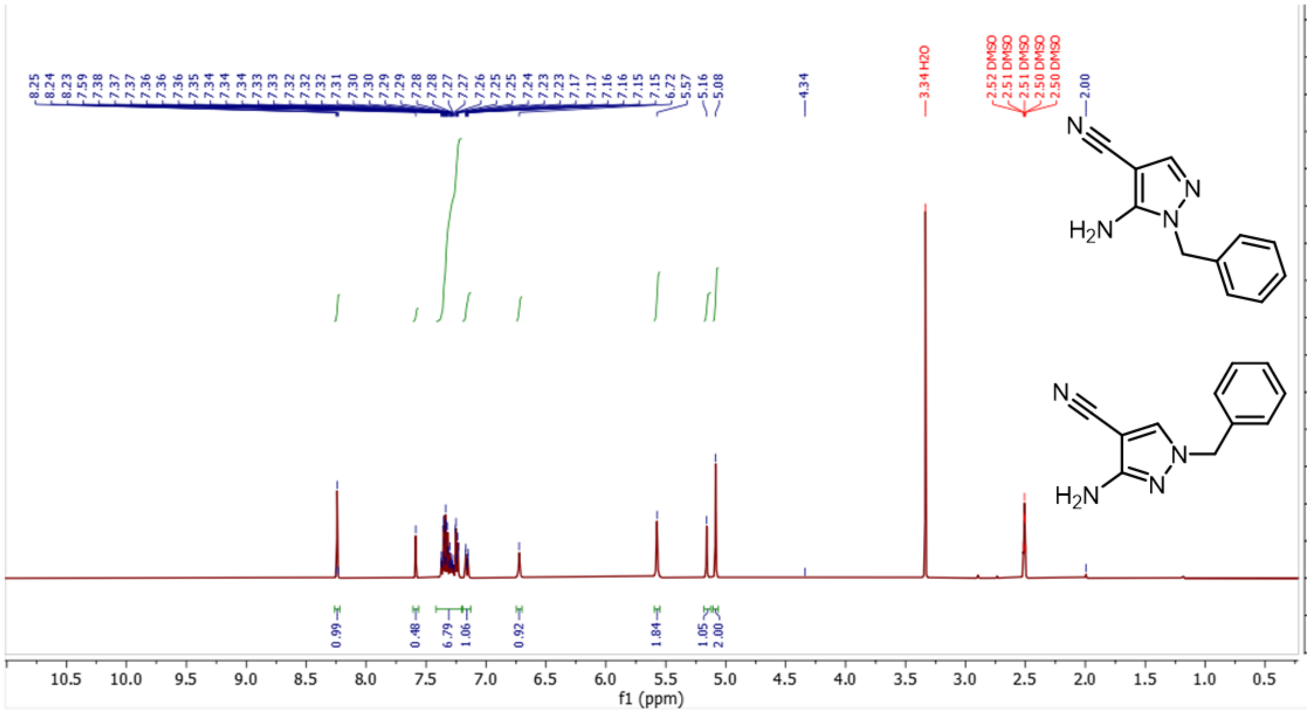

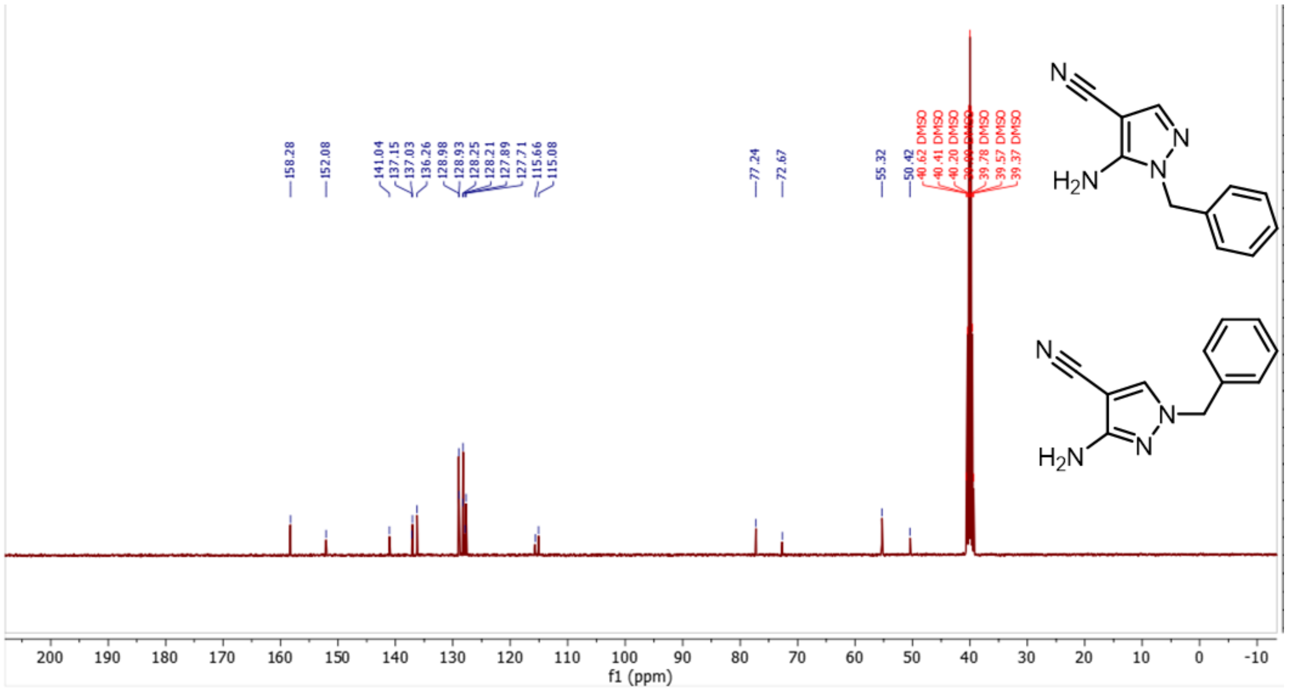

**S46**

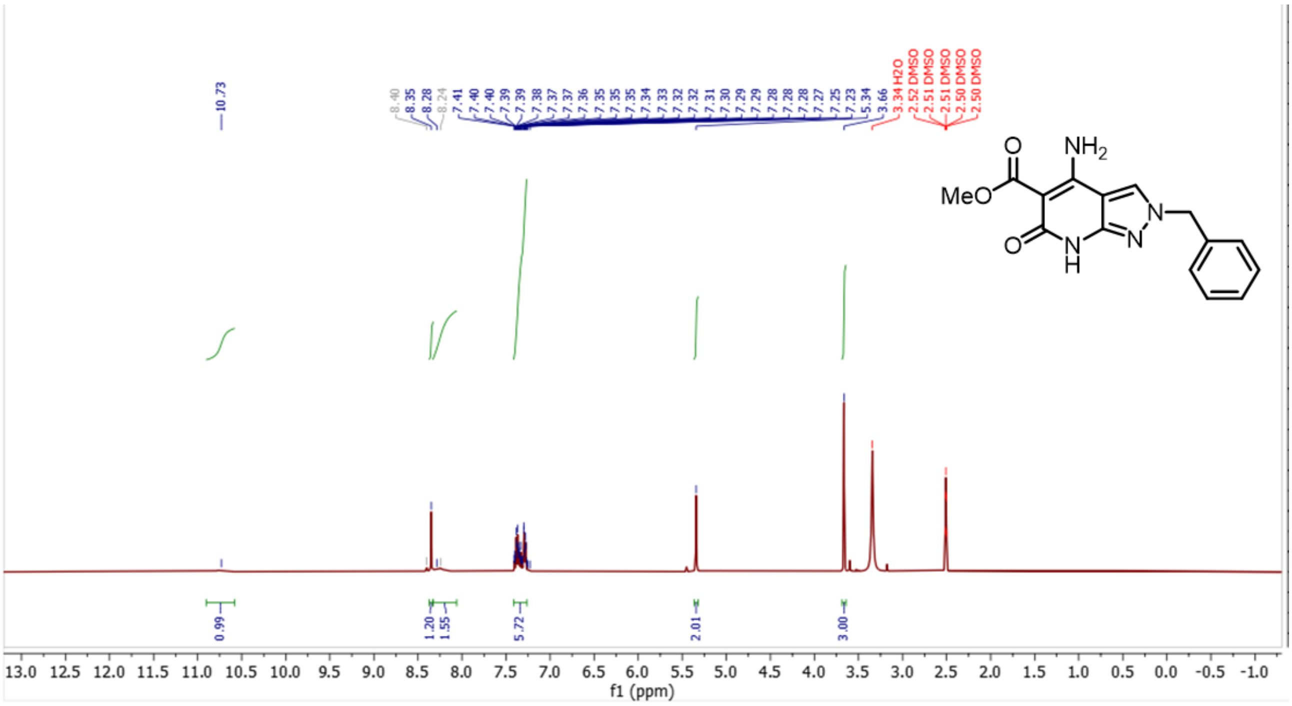

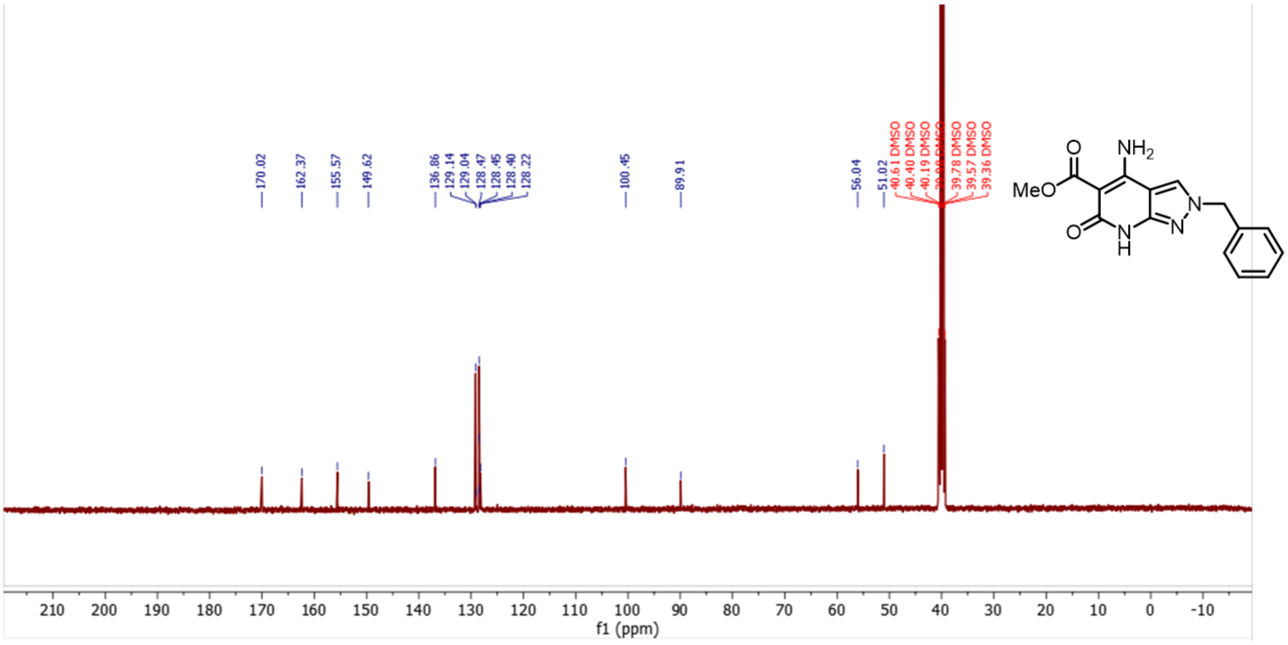

S47

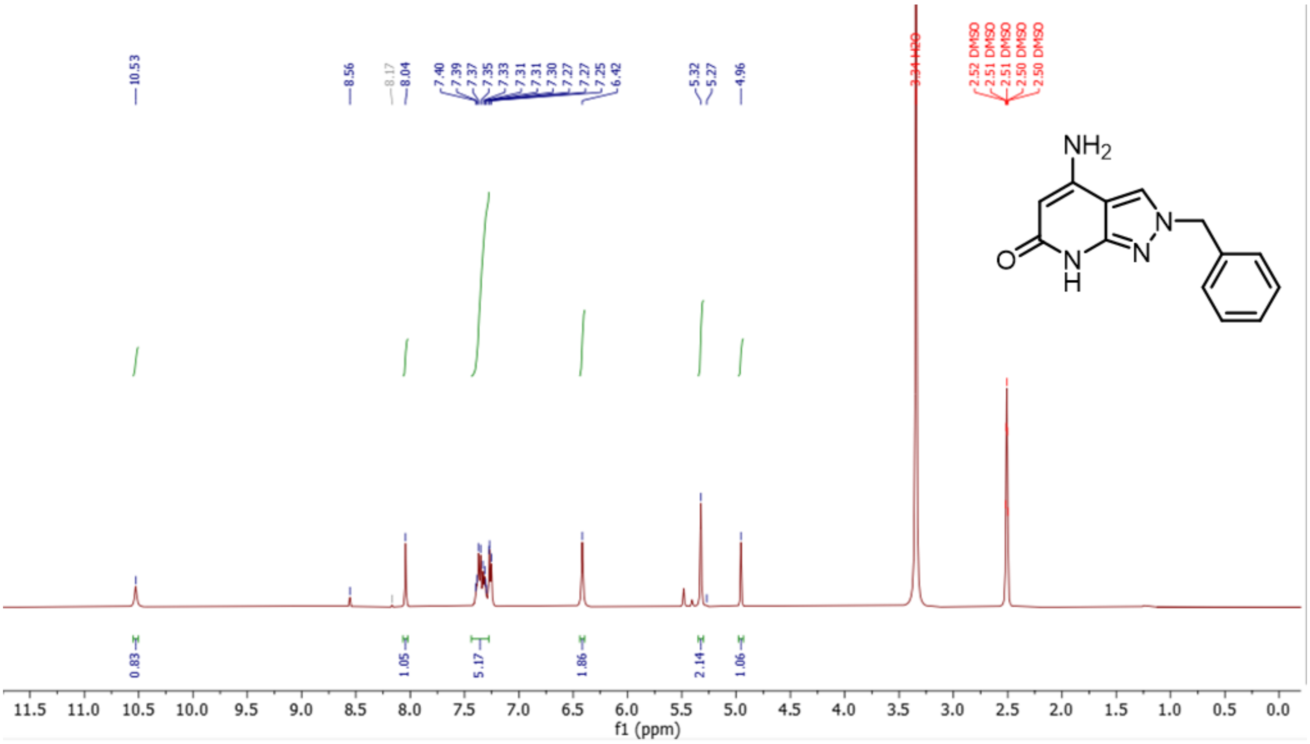

10.53
8.56
8.17
8.04
7.40
7.39
7.37
7.35
7.33
7.31
7.31
7.30
7.27
7.27
7.25
6.42
5.32
5.27
4.96
3.34 H2O
2.52 DMSO
2.51 DMSO
2.51 DMSO
2.50 DMSO
2.50 DMSO
NH2
O
N
H
N
N
0.83
1.05
5.17
1.86
2.14
1.06
11.5 11.0 10.5 10.0 9.5 9.0 8.5 8.0 7.5 7.0 6.5 6.0 5.5 5.0 4.5 4.0 3.5 3.0 2.5 2.0 1.5 1.0 0.5 0.0
f1 (ppm)

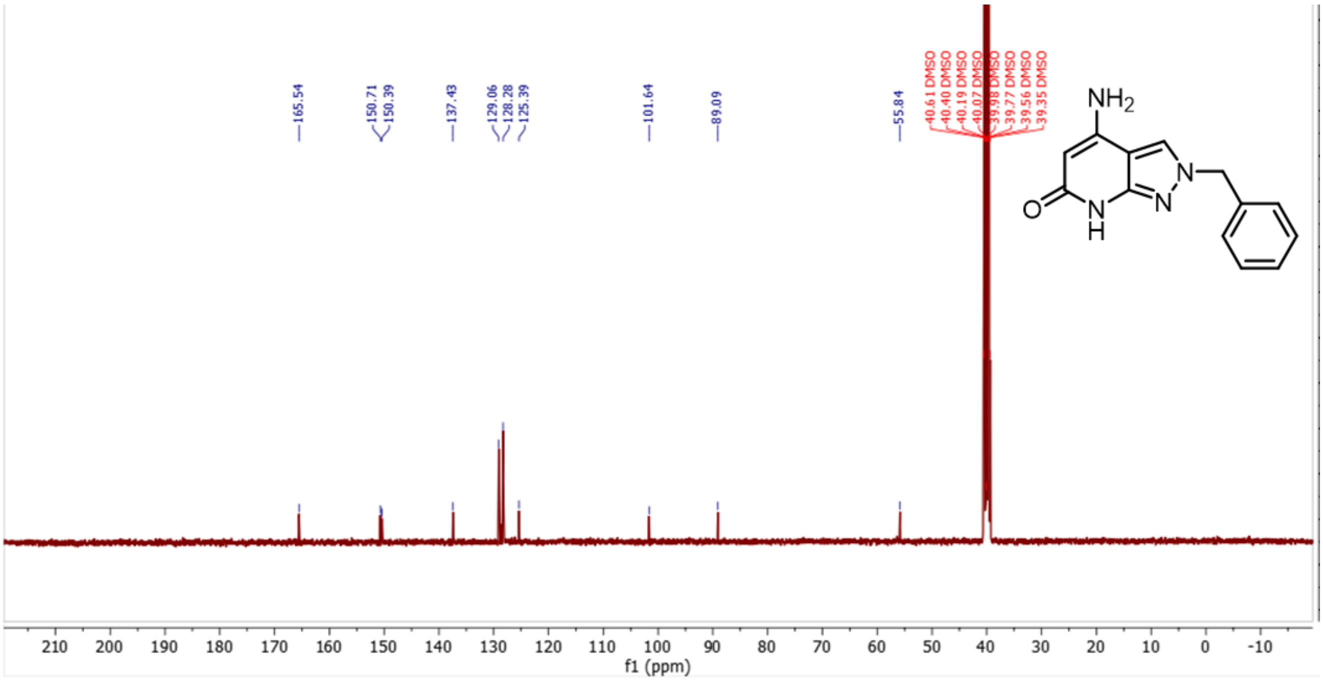

165.54
150.71
150.39
137.43
129.06
128.28
125.39
101.64
89.09
55.84
40.61 DMSO
40.40 DMSO
40.19 DMSO
40.07 DMSO
39.98 DMSO
39.77 DMSO
39.56 DMSO
39.35 DMSO
NH2
O
N
H
N
N
210 200 190 180 170 160 150 140 130 120 110 100 90 80 70 60 50 40 30 20 10 0 -10
f1 (ppm)

# S48

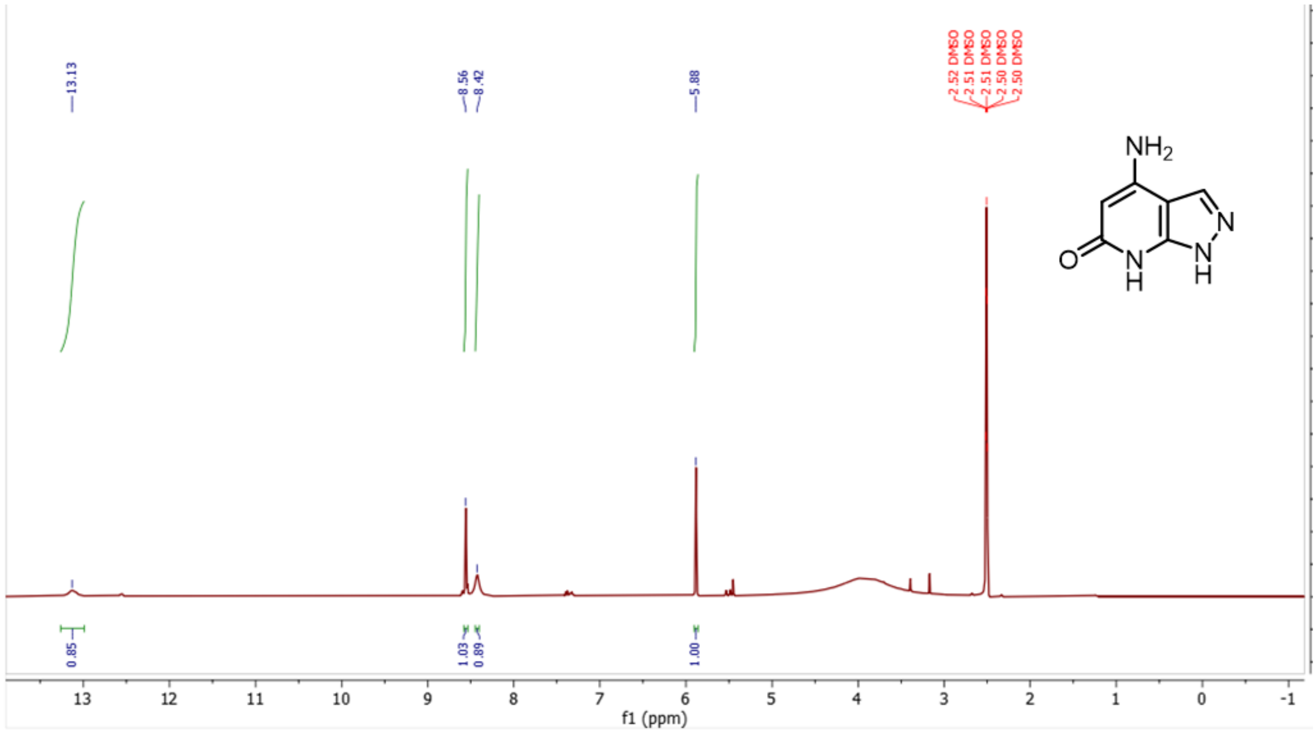
13.13
8.56
8.42
5.88
2.52 DMSO
2.51 DMSO
2.51 DMSO
2.50 DMSO
2.50 DMSO
NH2
N
N
H
N
H
O
0.85
1.03
0.89
1.00
f1 (ppm)

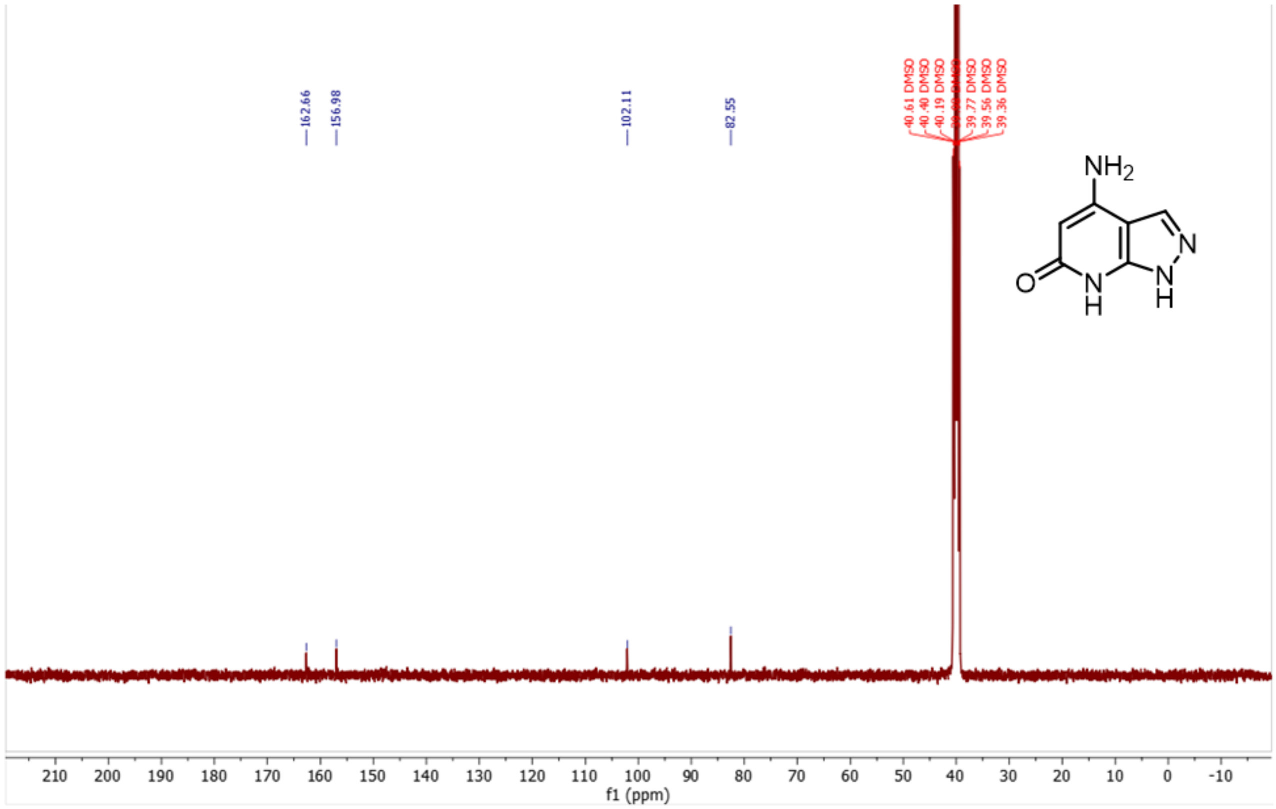
162.66
156.98
102.11
82.55
40.61 DMSO
40.40 DMSO
40.19 DMSO
39.77 DMSO
39.56 DMSO
39.36 DMSO
NH2
N
N
H
N
H
O
f1 (ppm)

**8**

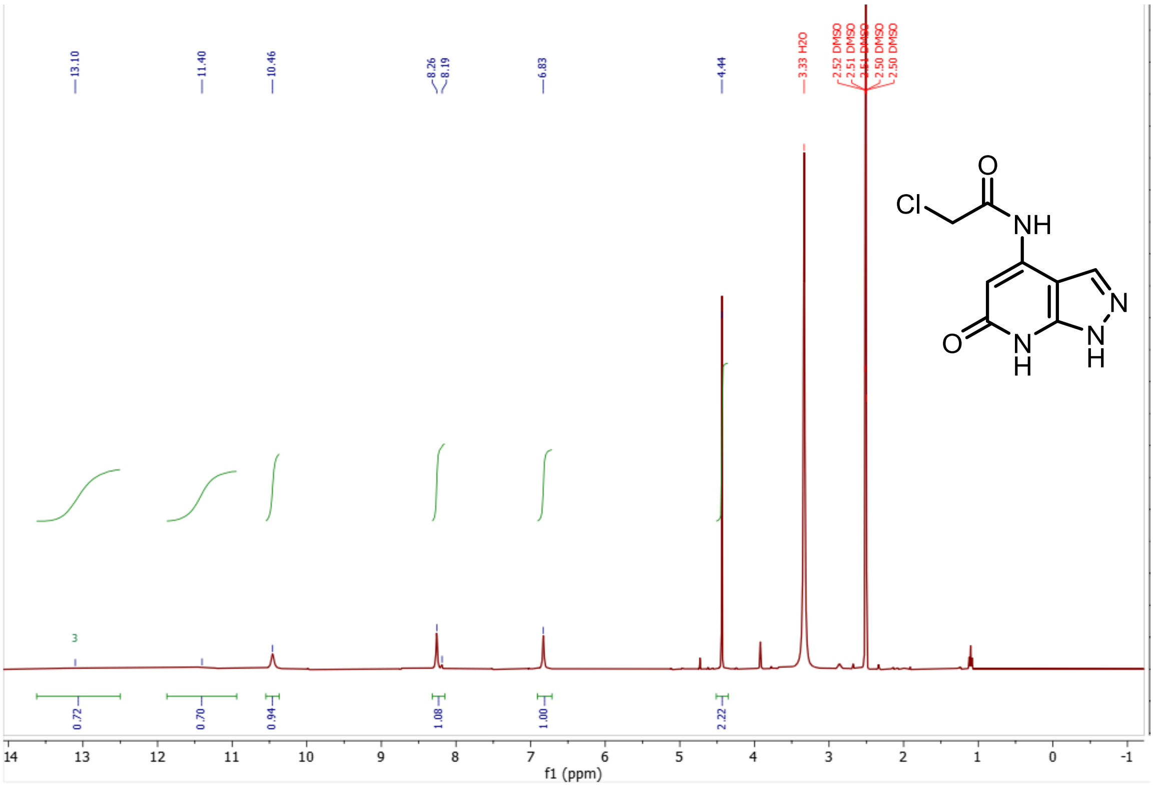
13.10
11.40
10.46
8.26
8.19
6.83
4.44
3.33 H2O
2.52 DMSO
2.51 DMSO
2.51 DMSO
2.50 DMSO
2.50 DMSO
0.72
0.70
0.94
1.08
1.00
2.22
14
13
12
11
10
9
8
7
6
5
4
3
2
1
0
-1
f1 (ppm)
Cl
O
NH
N
N
H
O
N
H

# S11

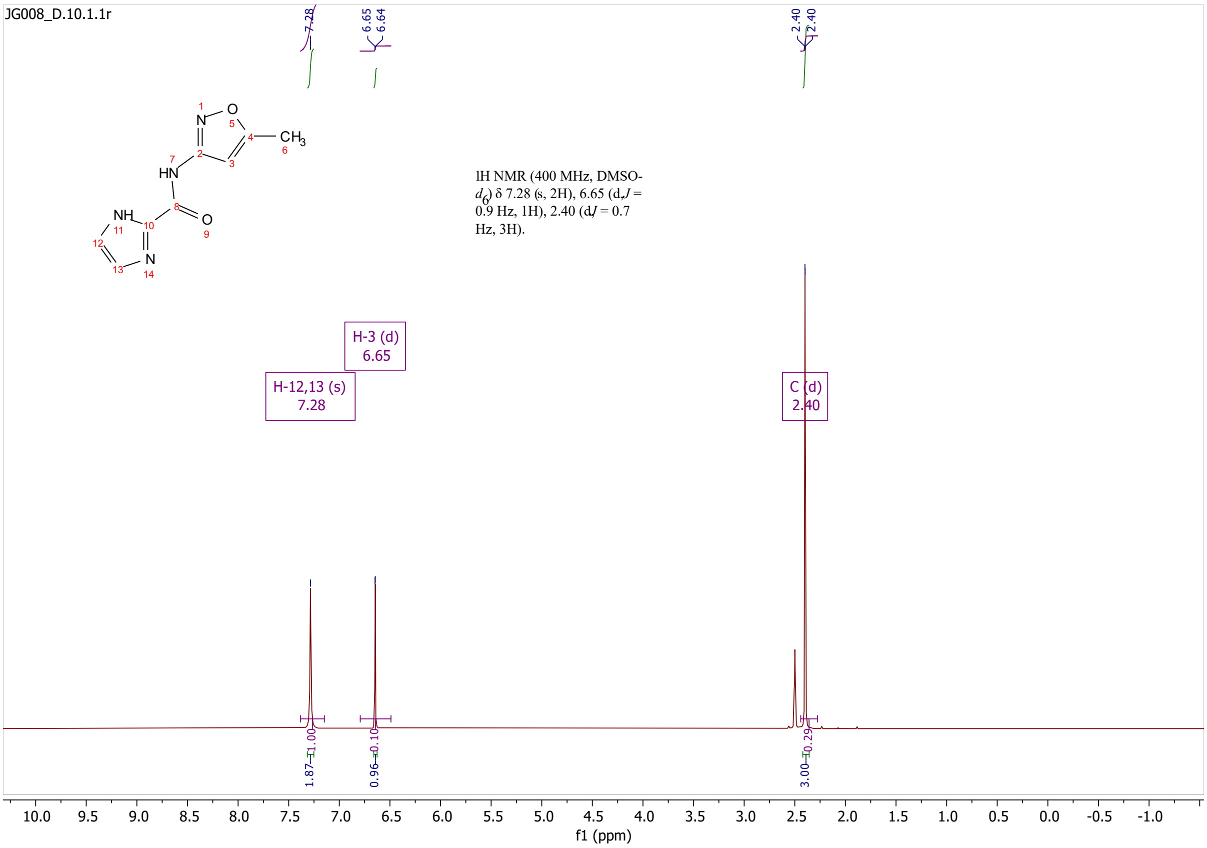
JG008_D.10.1.1r
1H NMR (400 MHz, DMSO-$d_6$) δ 7.28 (s, 2H), 6.65 (d, $J$ = 0.9 Hz, 1H), 2.40 (d, $J$ = 0.7 Hz, 3H).
H-3 (d) 6.65
H-12,13 (s) 7.28
C (d) 2.40
7.28
6.65
6.64
2.40
2.40
1.87
0.96
3.00
f1 (ppm)

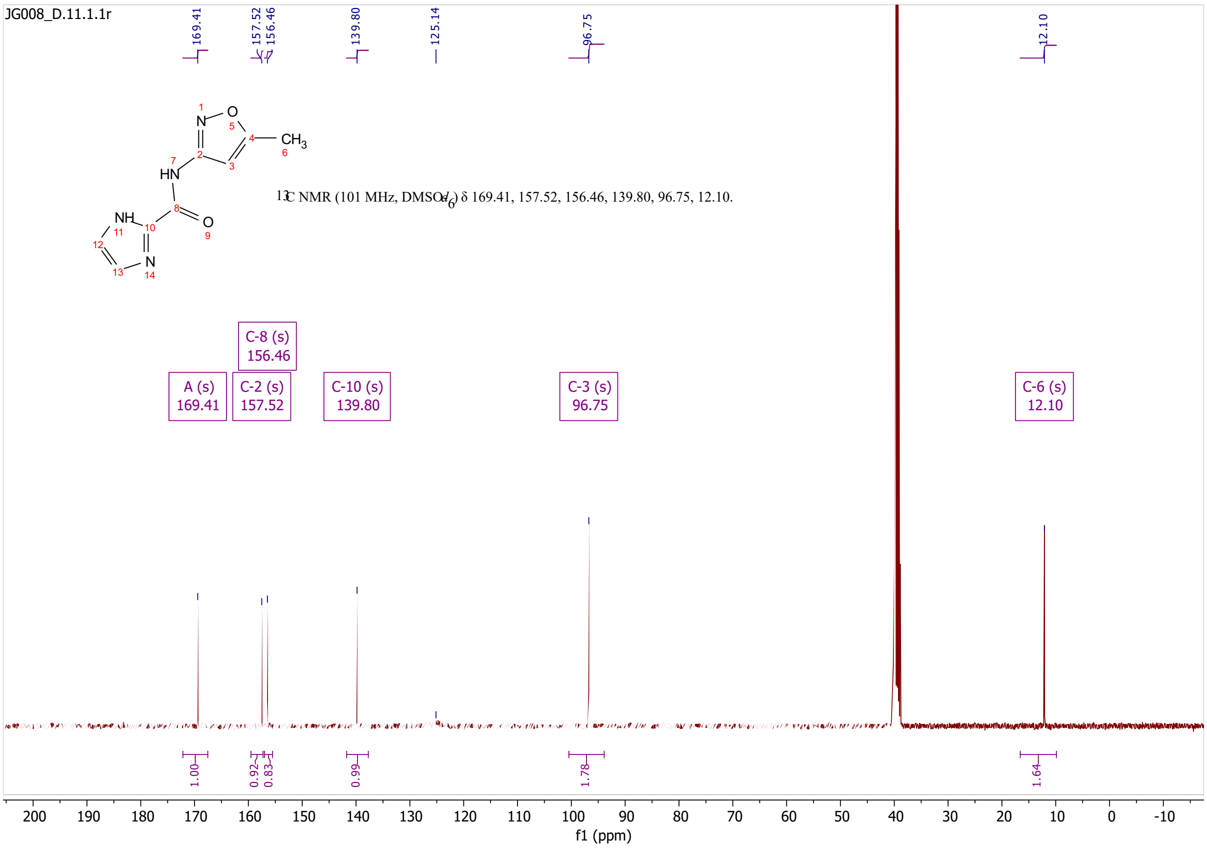
JG008_D.11.1.1r
13C NMR (101 MHz, DMSO-$d_6$) δ 169.41, 157.52, 156.46, 139.80, 96.75, 12.10.
169.41
157.52
156.46
139.80
125.14
96.75
12.10
C-8 (s) 156.46
A (s) 169.41
C-2 (s) 157.52
C-10 (s) 139.80
C-3 (s) 96.75
C-6 (s) 12.10
1.00
0.92
0.83
0.99
1.78
1.64
f1 (ppm)

# S12

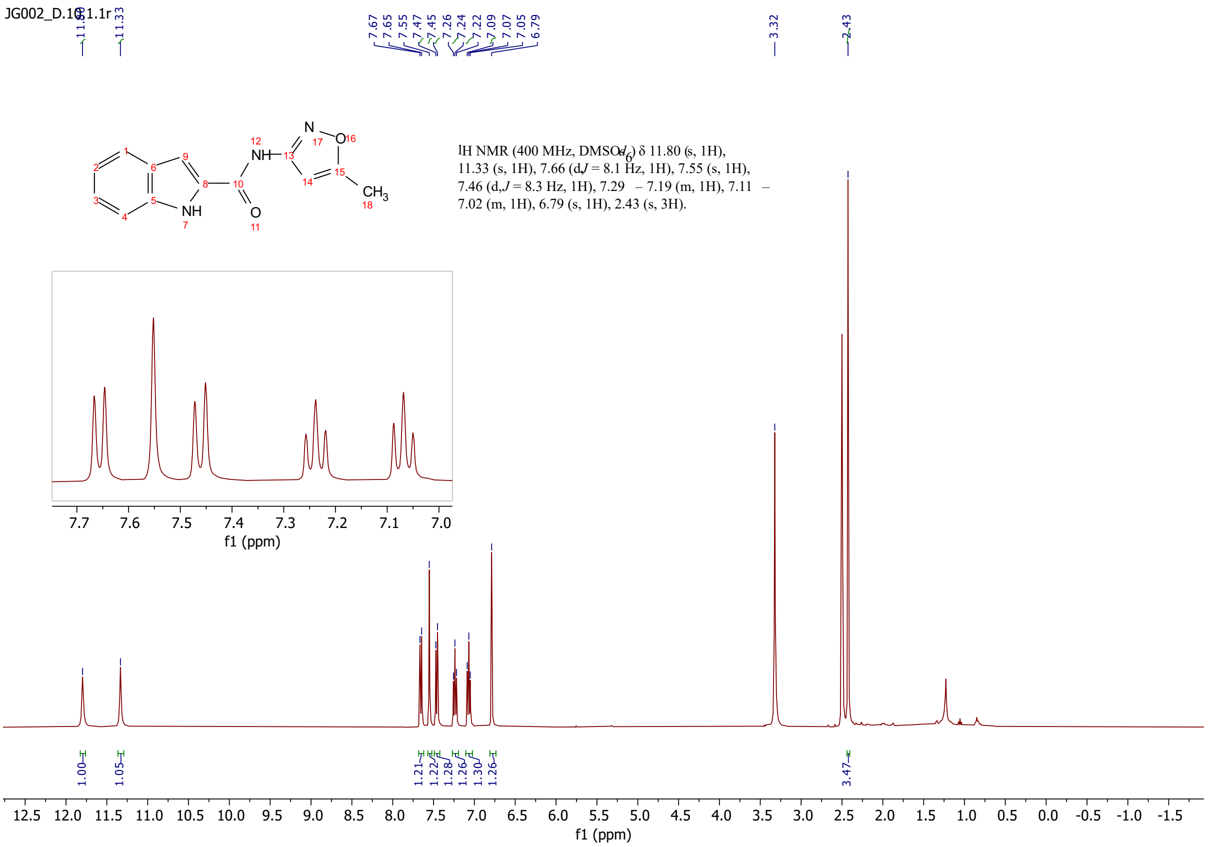
1H NMR (400 MHz, DMSO-$d_6$) δ 11.80 (s, 1H), 11.33 (s, 1H), 7.66 (d, J = 8.1 Hz, 1H), 7.55 (s, 1H), 7.46 (d, J = 8.3 Hz, 1H), 7.29 – 7.19 (m, 1H), 7.11 – 7.02 (m, 1H), 6.79 (s, 1H), 2.43 (s, 3H).
f1 (ppm)

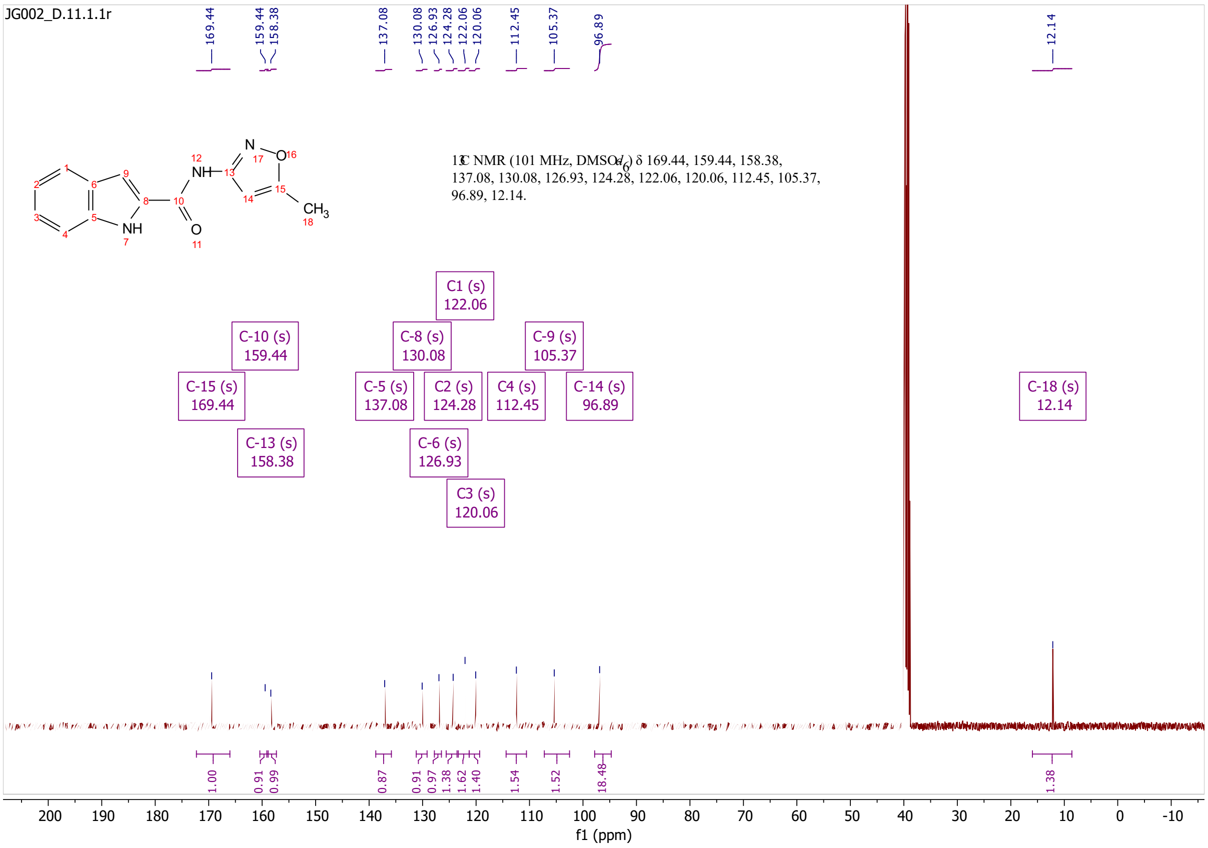
JG002_D.11.1.1r
13C NMR (101 MHz, DMSO-$d_6$) δ 169.44, 159.44, 158.38, 137.08, 130.08, 126.93, 124.28, 122.06, 120.06, 112.45, 105.37, 96.89, 12.14.
C1 (s) 122.06
C-10 (s) 159.44
C-8 (s) 130.08
C-9 (s) 105.37
C-15 (s) 169.44
C-5 (s) 137.08
C2 (s) 124.28
C4 (s) 112.45
C-14 (s) 96.89
C-18 (s) 12.14
C-13 (s) 158.38
C-6 (s) 126.93
C3 (s) 120.06
f1 (ppm)

JG006_T3P.12.fid

$^{1}$H NMR (400 MHz, DMSO-$d_6$) δ 10.67 (s, 1H), 6.59 – 6.57 (m, 1H), 4.30 (dd, $J$ = 8.2, 3.5 Hz, 1H), 3.46 – 3.31 (m, 2H), 2.37 (d, $J$ = 0.8 Hz, 3H), 2.23 – 2.13 (m, 1H), 1.93 – 1.77 (m, 3H), 1.34 (s, 9H).

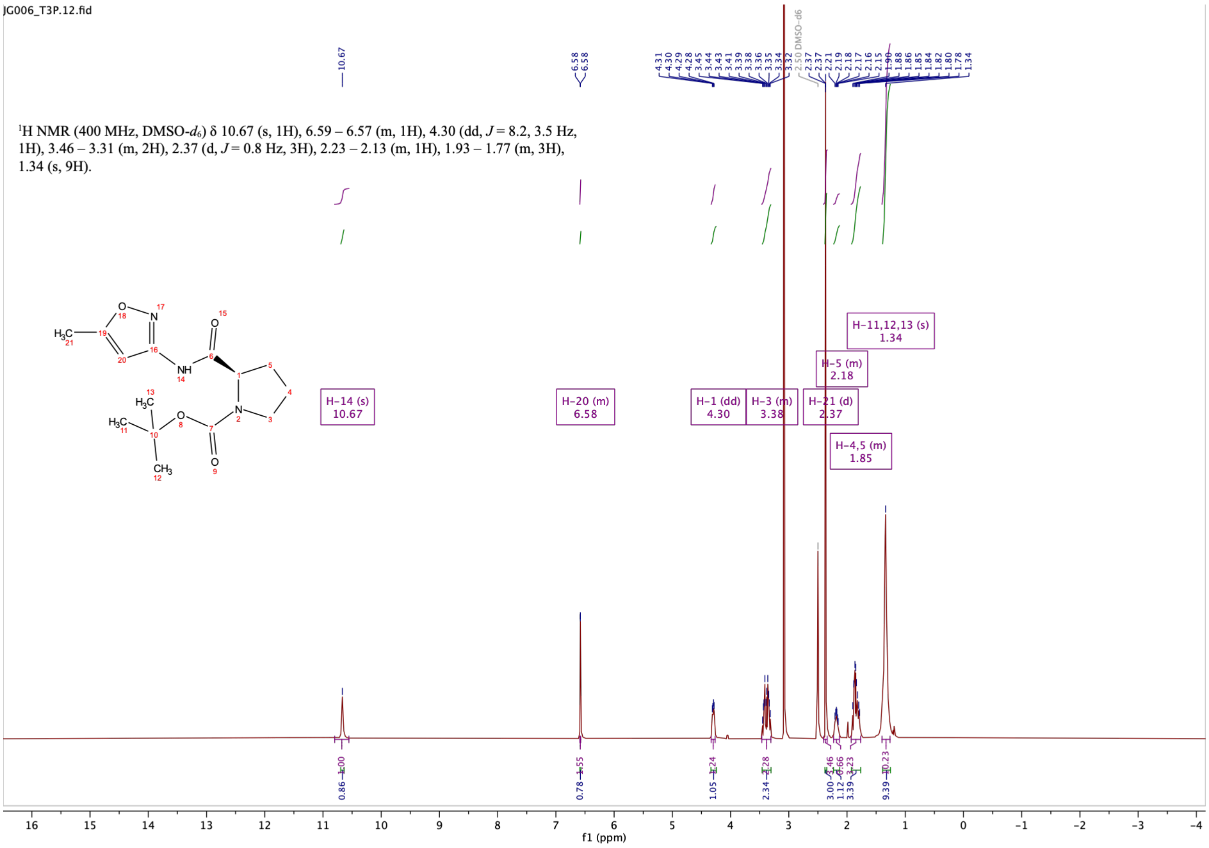

JG006_T3P.13.fid

$^{13}$C NMR (101 MHz, DMSO-$d_6$) δ 171.83, 171.30, 169.52, 169.44, 158.10, 158.07, 153.54, 152.96, 96.27, 96.15, 78.76, 78.60, 59.79, 59.50, 46.68, 46.46, 30.80, 30.02, 28.13, 27.91, 23.95, 23.36, 12.10.

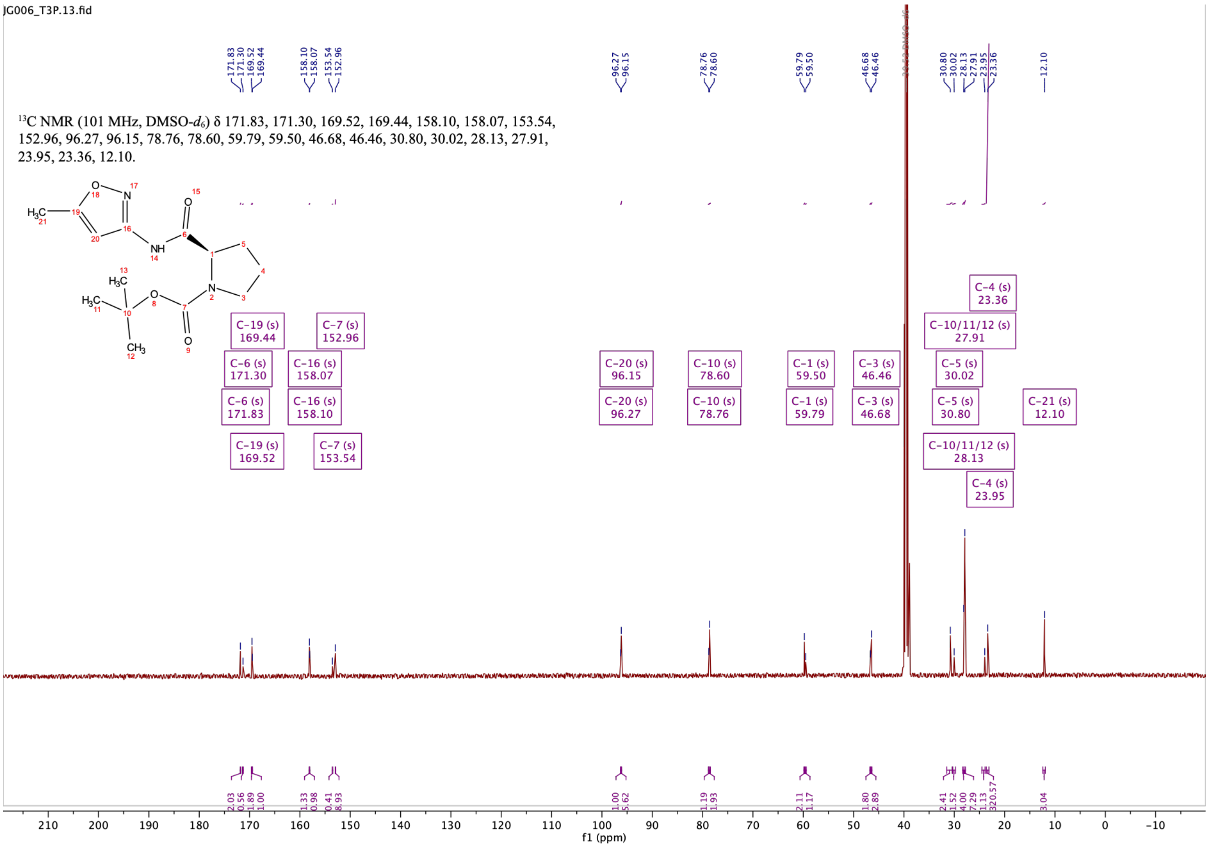

# S13

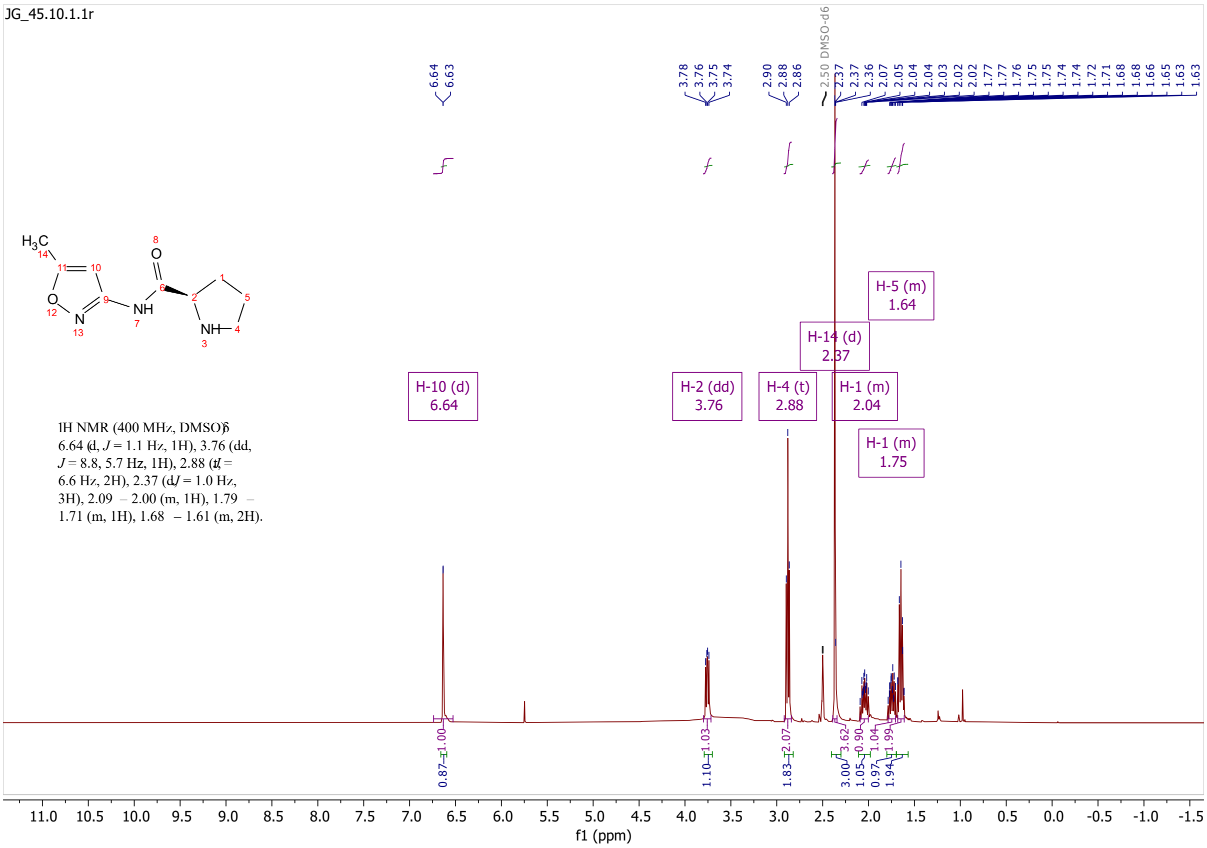

JG_45.10.1.1r
H-10 (d) 6.64
H-2 (dd) 3.76
H-4 (t) 2.88
H-14 (d) 2.37
H-1 (m) 2.04
H-1 (m) 1.75
H-5 (m) 1.64
1H NMR (400 MHz, DMSO) δ 6.64 (d, J = 1.1 Hz, 1H), 3.76 (dd, J = 8.8, 5.7 Hz, 1H), 2.88 (t, J = 6.6 Hz, 2H), 2.37 (d, J = 1.0 Hz, 3H), 2.09 – 2.00 (m, 1H), 1.79 – 1.71 (m, 1H), 1.68 – 1.61 (m, 2H).
f1 (ppm)

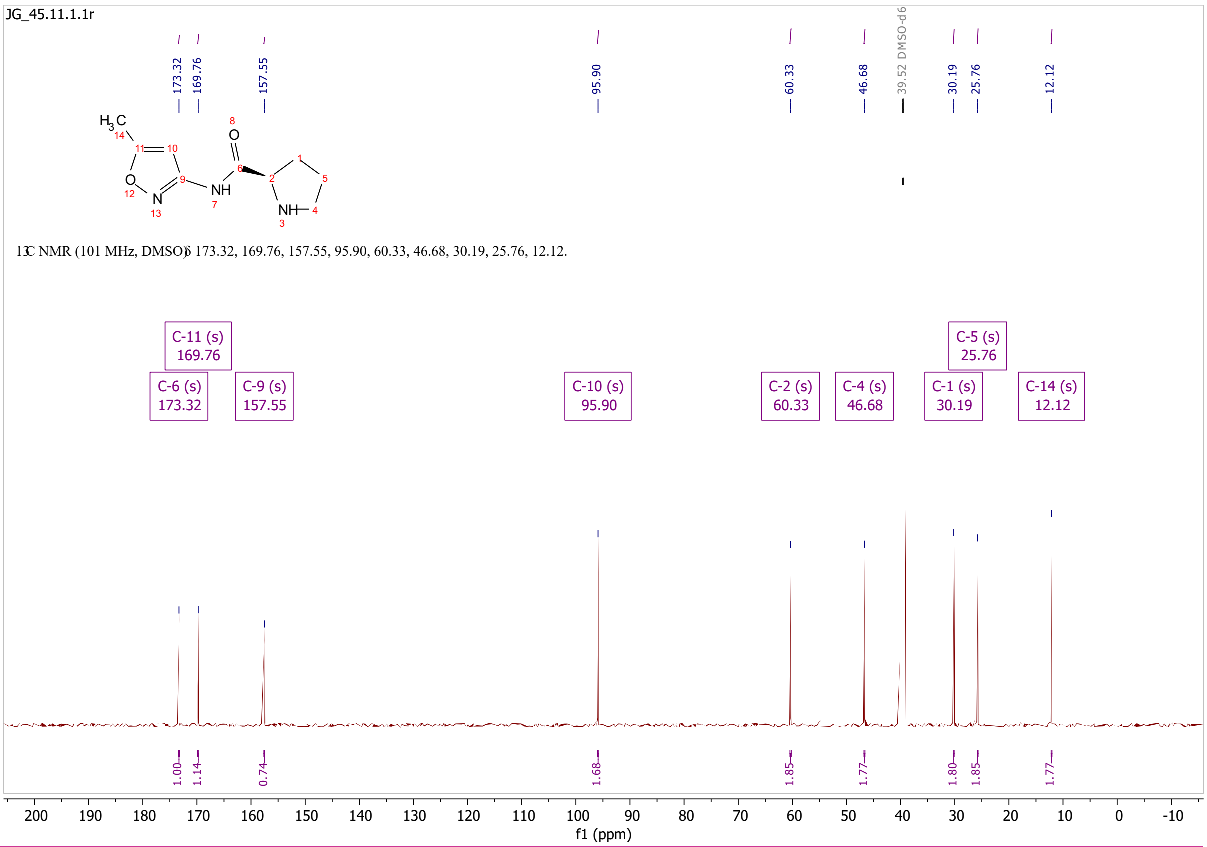

JG_45.11.1.1r
173.32
169.76
157.55
95.90
60.33
46.68
39.52 DMSO-d6
30.19
25.76
12.12
13C NMR (101 MHz, DMSO) δ 173.32, 169.76, 157.55, 95.90, 60.33, 46.68, 30.19, 25.76, 12.12.
C-11 (s) 169.76
C-6 (s) 173.32
C-9 (s) 157.55
C-10 (s) 95.90
C-2 (s) 60.33
C-4 (s) 46.68
C-1 (s) 30.19
C-5 (s) 25.76
C-14 (s) 12.12
f1 (ppm)

# S50

JG005_T3P.10.1.1r

[1]H NMR (400 MHz, DMSO-$d_6$) δ 10.67 (s, 1H), 6.58 (d, $J$ = 0.9 Hz, 1H), 4.30 (dd, $J$ = 8.3, 3.5 Hz, 1H), 3.47 – 3.30 (m, 2H), 2.37 (d, $J$ = 0.9 Hz, 3H), 2.23 – 2.14 (m, 1H), 1.93 – 1.76 (m, 3H), 1.34 (s, 9H).

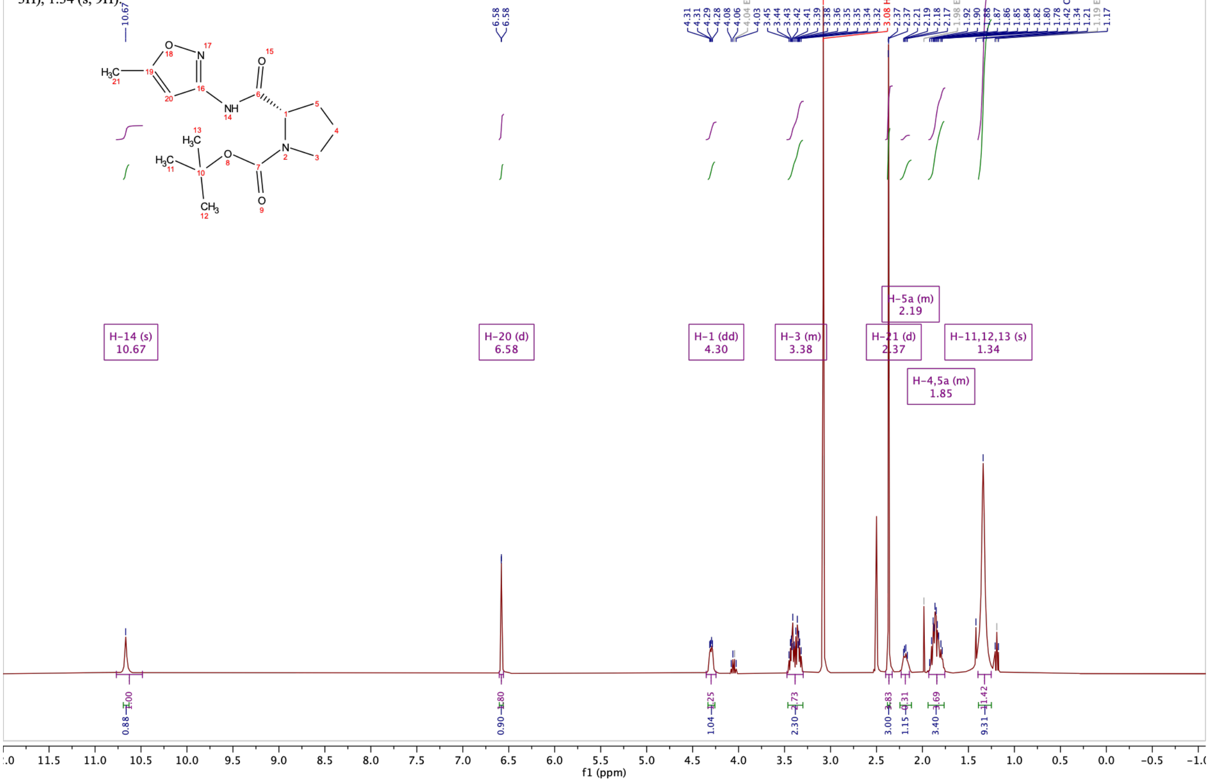

JG005_T3P.11.1.1r

[13]C NMR (101 MHz, DMSO-$d_6$) δ 171.84, 171.31, 169.53, 169.45, 158.12, 158.08, 153.55, 152.97, 96.28, 96.16, 78.76, 78.60, 59.79, 59.50, 46.68, 46.46, 30.80, 30.02, 28.12, 27.90, 23.95, 23.35, 12.10.

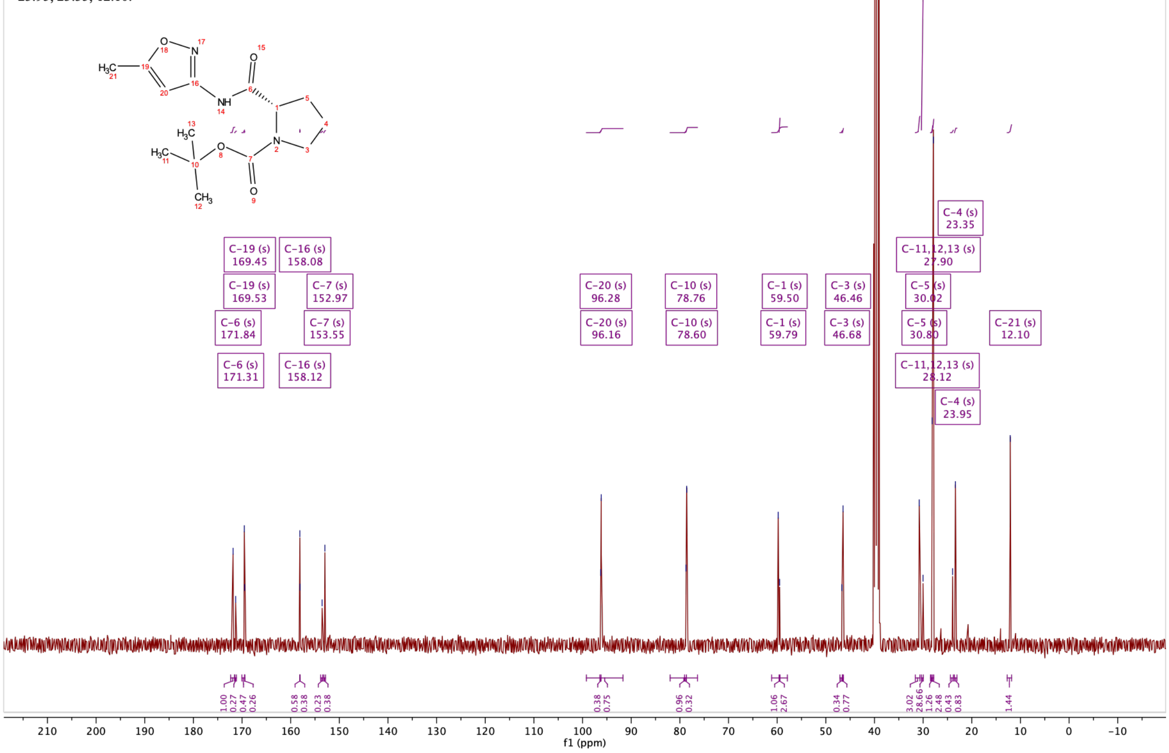

# S14

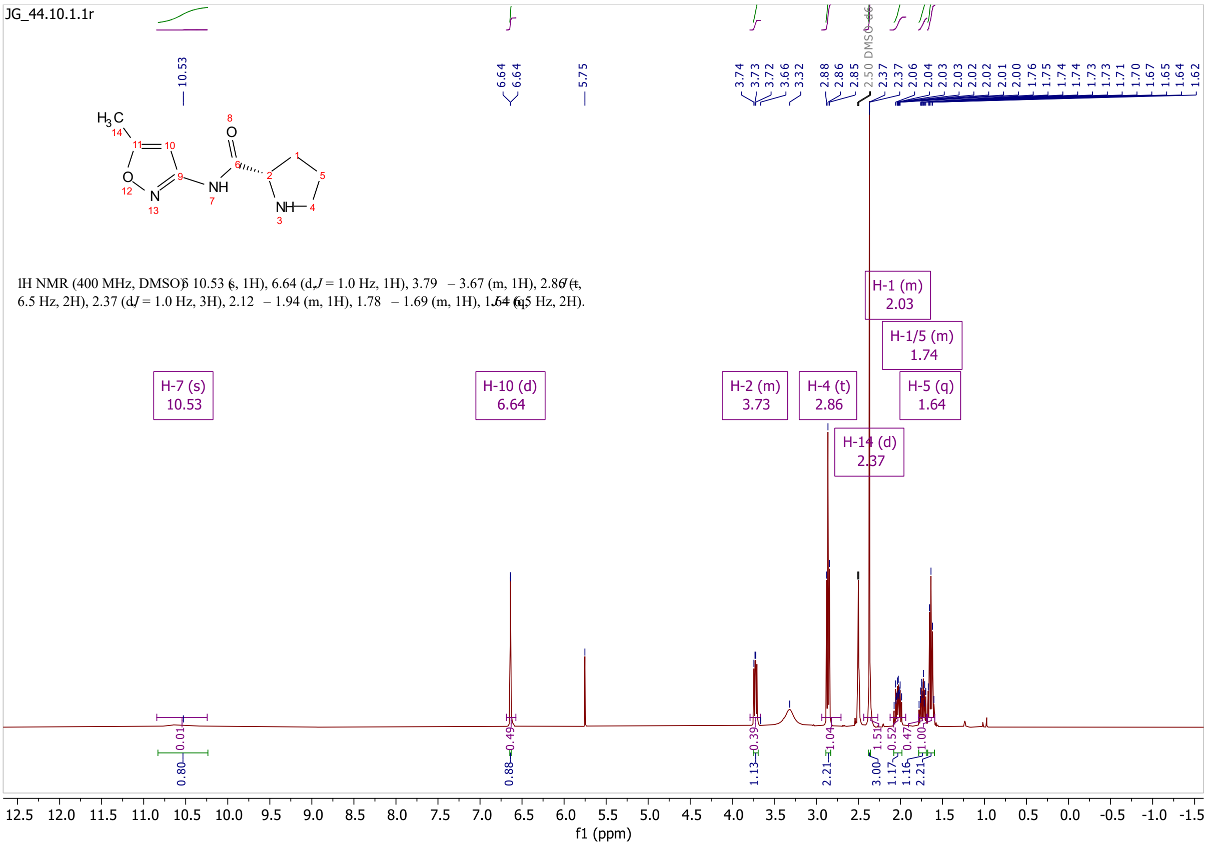

JG_44.10.1.1r
1H NMR (400 MHz, DMSO) δ 10.53 (s, 1H), 6.64 (d, J = 1.0 Hz, 1H), 3.79 – 3.67 (m, 1H), 2.86 (t, J = 6.5 Hz, 2H), 2.37 (d, J = 1.0 Hz, 3H), 2.12 – 1.94 (m, 1H), 1.78 – 1.69 (m, 1H), 1.64 (q, J = 6.5 Hz, 2H).
H-7 (s) 10.53
H-10 (d) 6.64
H-2 (m) 3.73
H-4 (t) 2.86
H-14 (d) 2.37
H-1 (m) 2.03
H-1/5 (m) 1.74
H-5 (q) 1.64
2.50 DMSO-d6
f1 (ppm)

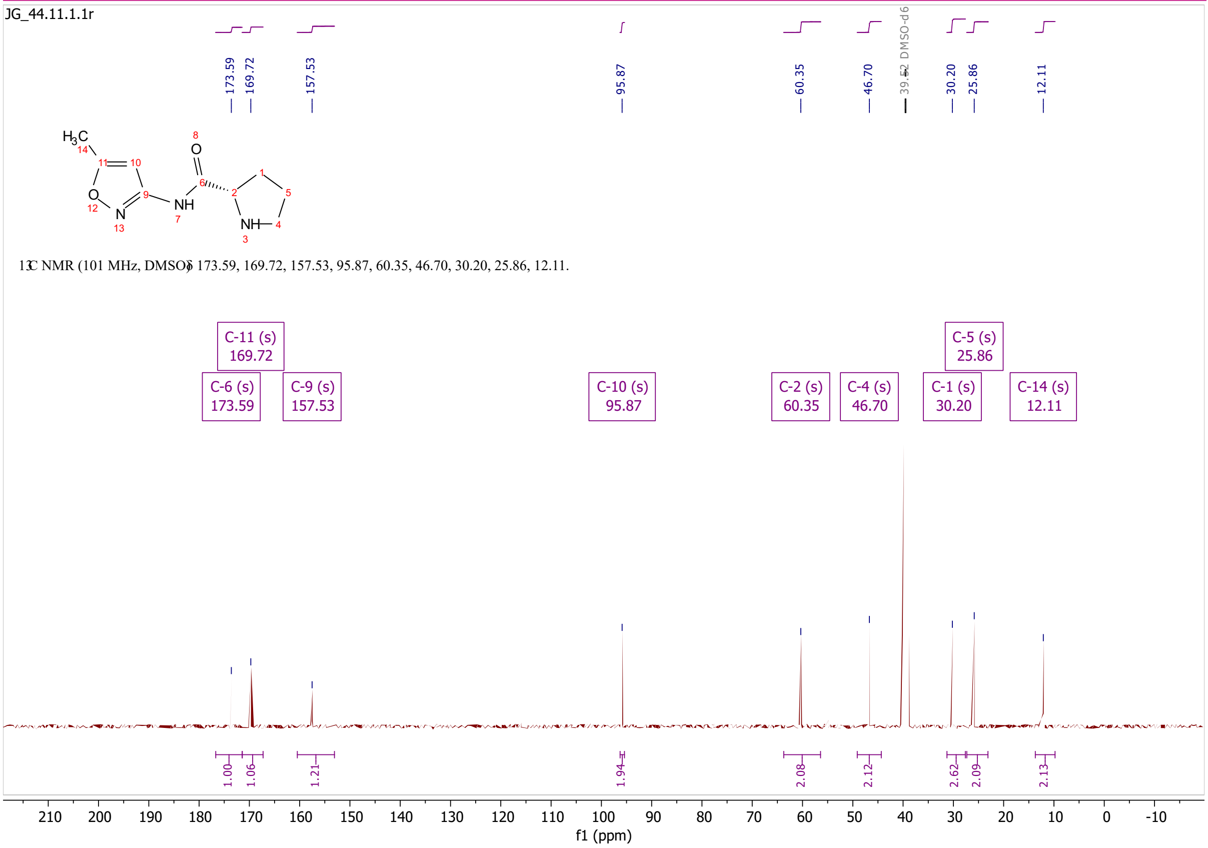

JG_44.11.1.1r
13C NMR (101 MHz, DMSO) δ 173.59, 169.72, 157.53, 95.87, 60.35, 46.70, 30.20, 25.86, 12.11.
C-11 (s) 169.72
C-6 (s) 173.59
C-9 (s) 157.53
C-10 (s) 95.87
C-2 (s) 60.35
C-4 (s) 46.70
C-1 (s) 30.20
C-5 (s) 25.86
C-14 (s) 12.11
39.52 DMSO-d6
f1 (ppm)

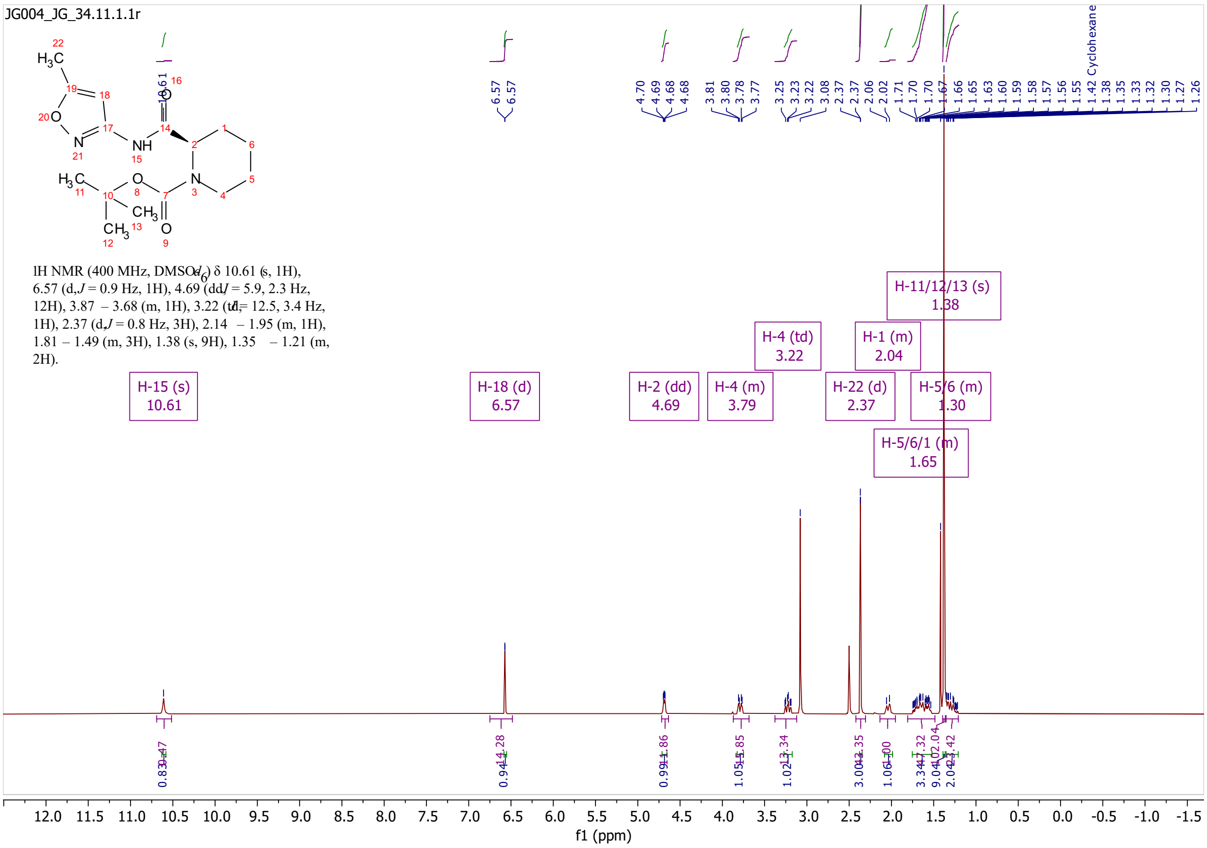
JG004_JG_34.11.1.1r
1H NMR (400 MHz, DMSO-$d_6$) δ 10.61 (s, 1H), 6.57 (d, $J$ = 0.9 Hz, 1H), 4.69 (dd, $J$ = 5.9, 2.3 Hz, 12H), 3.87 – 3.68 (m, 1H), 3.22 (td, $J$ = 12.5, 3.4 Hz, 1H), 2.37 (d, $J$ = 0.8 Hz, 3H), 2.14 – 1.95 (m, 1H), 1.81 – 1.49 (m, 3H), 1.38 (s, 9H), 1.35 – 1.21 (m, 2H).
H-15 (s) 10.61
H-18 (d) 6.57
H-2 (dd) 4.69
H-4 (m) 3.79
H-4 (td) 3.22
H-22 (d) 2.37
H-1 (m) 2.04
H-5/6/1 (m) 1.65
H-11/12/13 (s) 1.38
H-5/6 (m) 1.30
Cyclohexane
f1 (ppm)

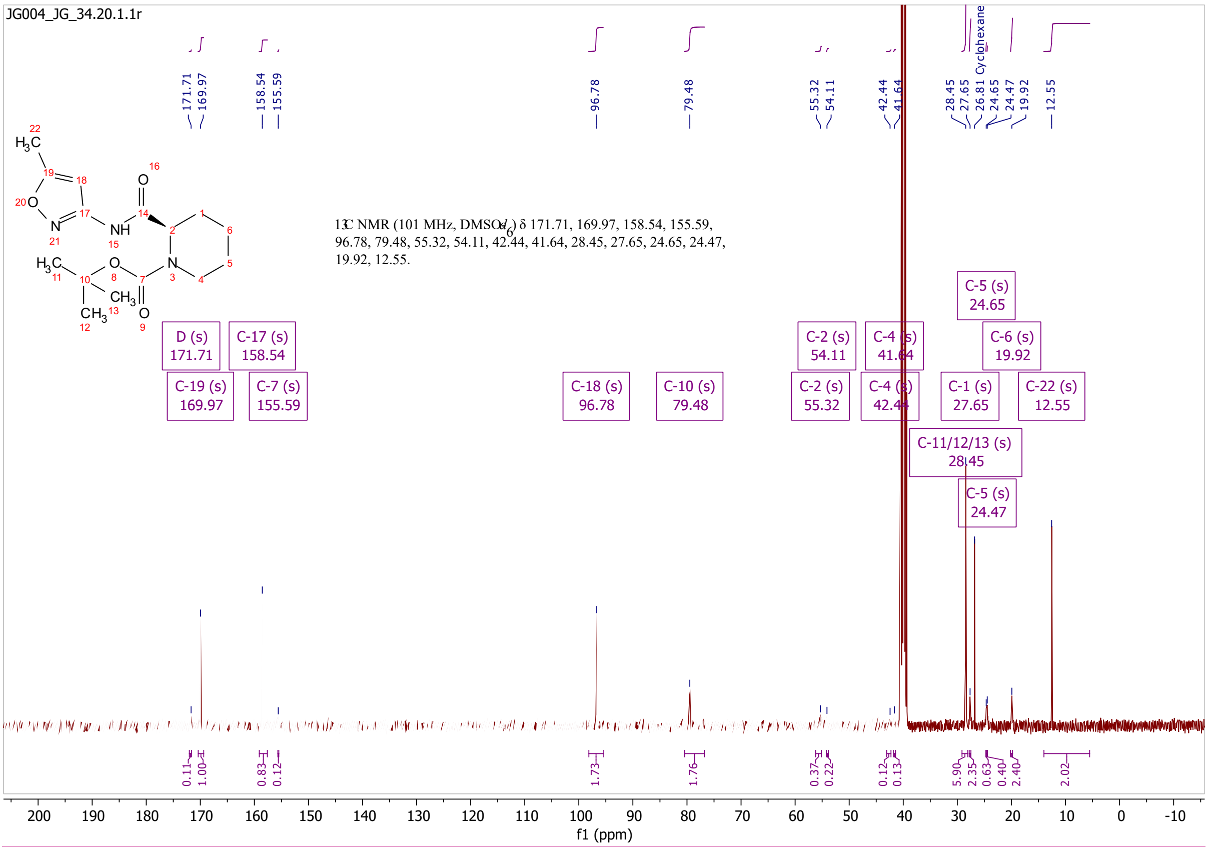
JG004_JG_34.20.1.1r
13C NMR (101 MHz, DMSO-$d_6$) δ 171.71, 169.97, 158.54, 155.59, 96.78, 79.48, 55.32, 54.11, 42.44, 41.64, 28.45, 27.65, 24.65, 24.47, 19.92, 12.55.
D (s) 171.71
C-19 (s) 169.97
C-17 (s) 158.54
C-7 (s) 155.59
C-18 (s) 96.78
C-10 (s) 79.48
C-2 (s) 55.32
C-2 (s) 54.11
C-4 (s) 42.44
C-4 (s) 41.64
C-11/12/13 (s) 28.45
C-1 (s) 27.65
C-5 (s) 24.65
C-5 (s) 24.47
C-6 (s) 19.92
C-22 (s) 12.55
Cyclohexane
f1 (ppm)

# S15

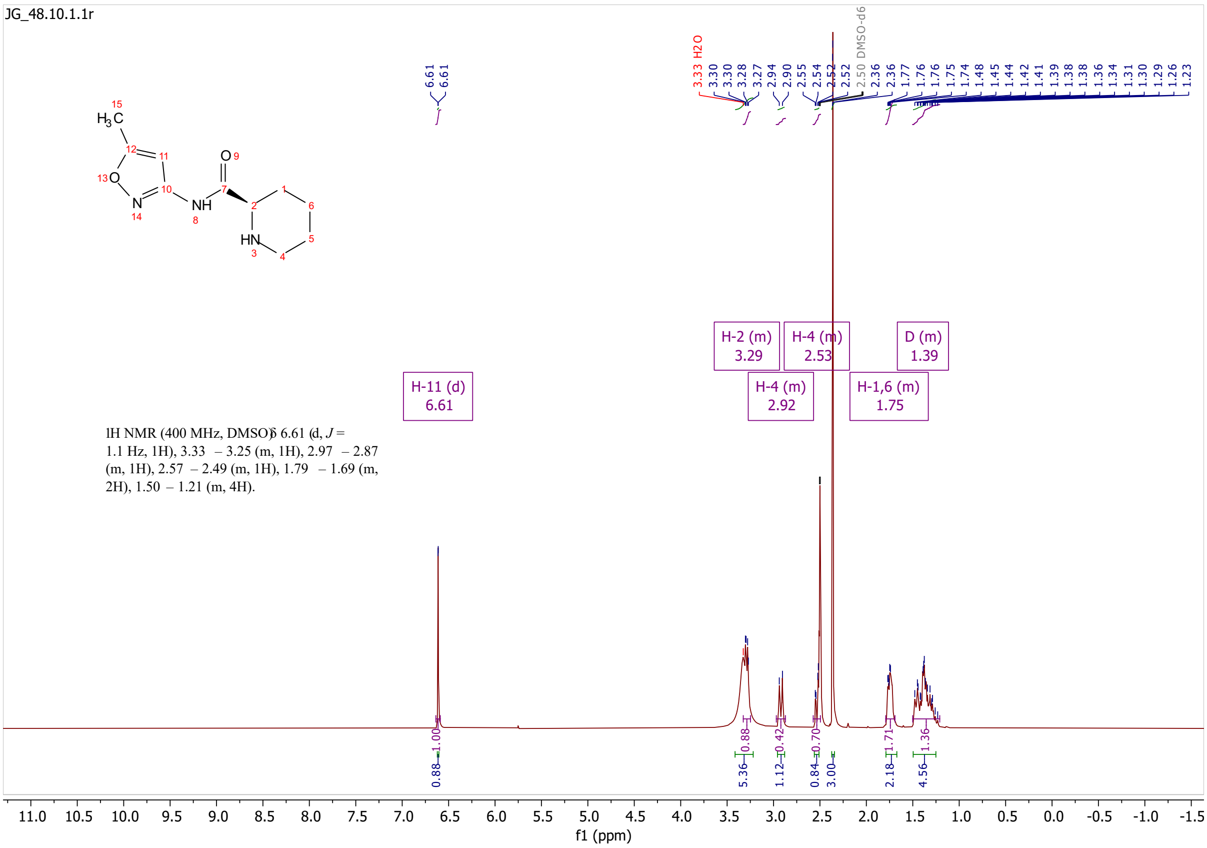
JG_48.10.1.1r
H-11 (d) 6.61
H-2 (m) 3.29
H-4 (m) 2.92
H-4 (m) 2.53
H-1,6 (m) 1.75
D (m) 1.39
1H NMR (400 MHz, DMSO) δ 6.61 (d, J = 1.1 Hz, 1H), 3.33 – 3.25 (m, 1H), 2.97 – 2.87 (m, 1H), 2.57 – 2.49 (m, 1H), 1.79 – 1.69 (m, 2H), 1.50 – 1.21 (m, 4H).
3.33 H2O
2.50 DMSO-d6
f1 (ppm)

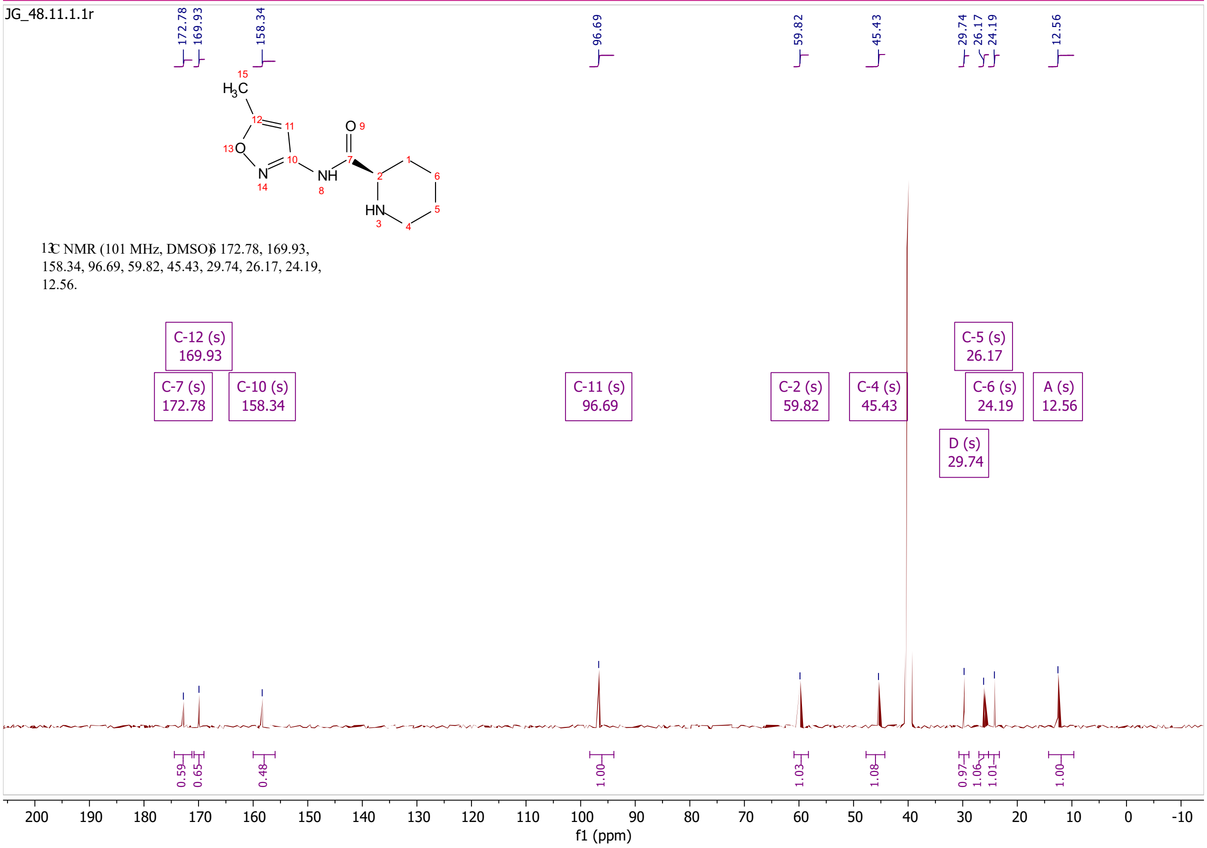
JG_48.11.1.1r
13C NMR (101 MHz, DMSO) δ 172.78, 169.93, 158.34, 96.69, 59.82, 45.43, 29.74, 26.17, 24.19, 12.56.
C-12 (s) 169.93
C-7 (s) 172.78
C-10 (s) 158.34
C-11 (s) 96.69
C-2 (s) 59.82
C-4 (s) 45.43
C-5 (s) 26.17
C-6 (s) 24.19
A (s) 12.56
D (s) 29.74
f1 (ppm)

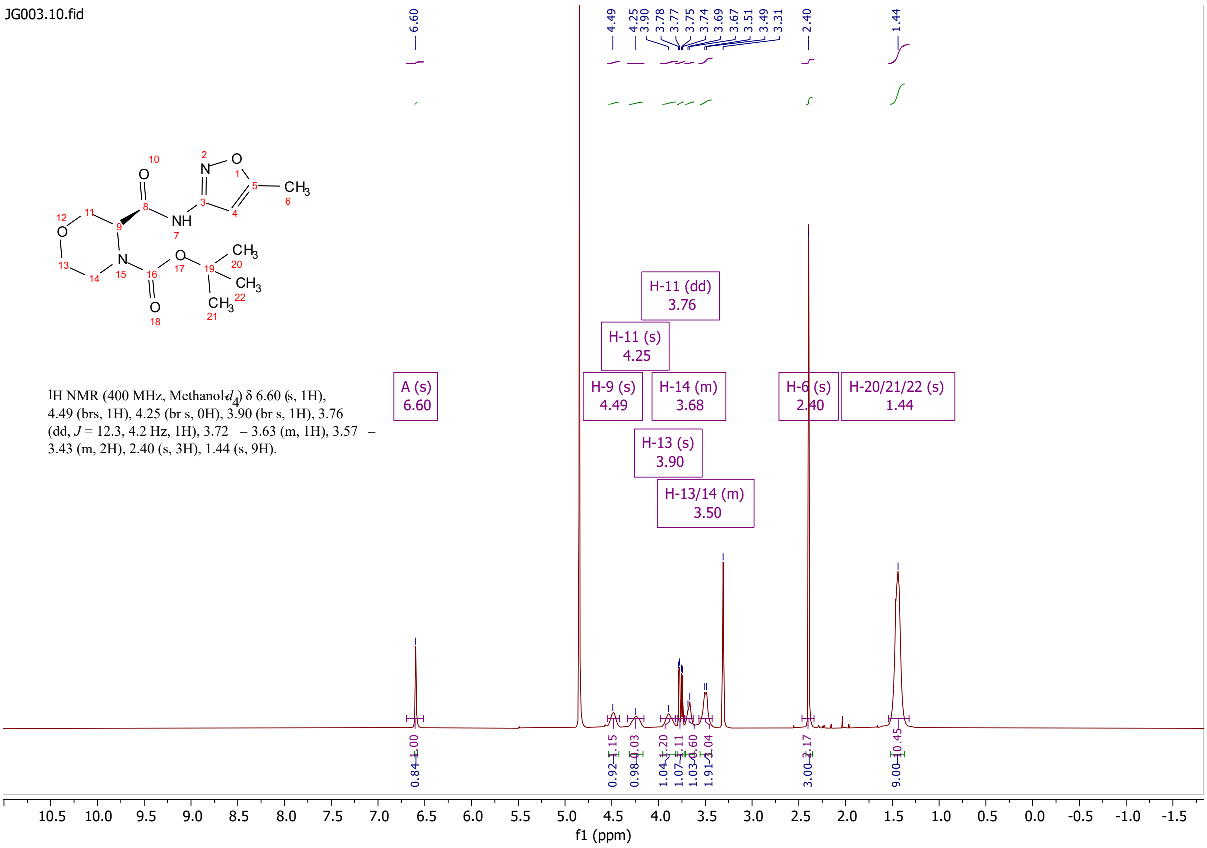
JG003.10.fid
1H NMR (400 MHz, Methanol-$d_4$) δ 6.60 (s, 1H), 4.49 (brs, 1H), 4.25 (br s, 0H), 3.90 (br s, 1H), 3.76 (dd, $J$ = 12.3, 4.2 Hz, 1H), 3.72 – 3.63 (m, 1H), 3.57 – 3.43 (m, 2H), 2.40 (s, 3H), 1.44 (s, 9H).
A (s) 6.60
H-9 (s) 4.49
H-11 (s) 4.25
H-11 (dd) 3.76
H-13 (s) 3.90
H-14 (m) 3.68
H-13/14 (m) 3.50
H-6 (s) 2.40
H-20/21/22 (s) 1.44
f1 (ppm)

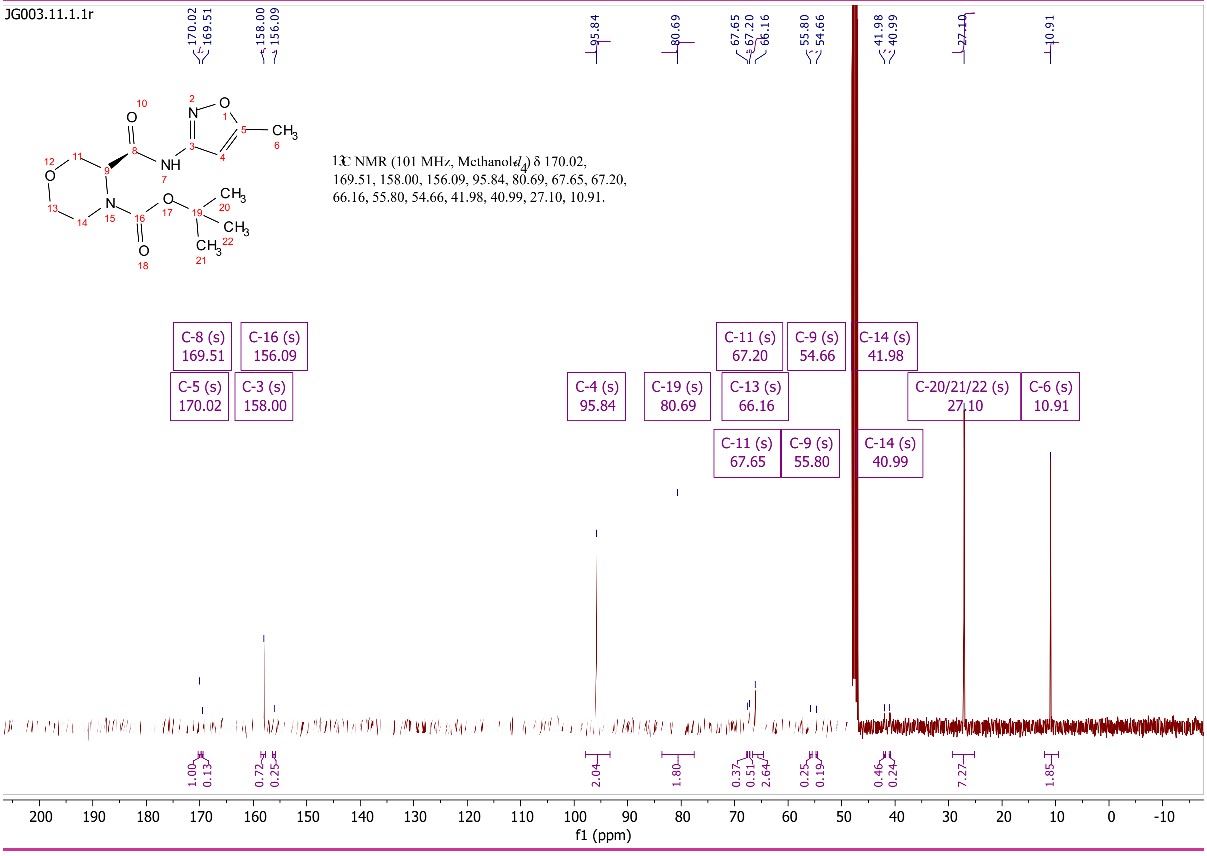
JG003.11.1.1r
13C NMR (101 MHz, Methanol-$d_4$) δ 170.02, 169.51, 158.00, 156.09, 95.84, 80.69, 67.65, 67.20, 66.16, 55.80, 54.66, 41.98, 40.99, 27.10, 10.91.
C-8 (s) 169.51
C-16 (s) 156.09
C-5 (s) 170.02
C-3 (s) 158.00
C-4 (s) 95.84
C-19 (s) 80.69
C-11 (s) 67.20
C-13 (s) 66.16
C-11 (s) 67.65
C-9 (s) 54.66
C-9 (s) 55.80
C-14 (s) 41.98
C-14 (s) 40.99
C-20/21/22 (s) 27.10
C-6 (s) 10.91
f1 (ppm)

# S16

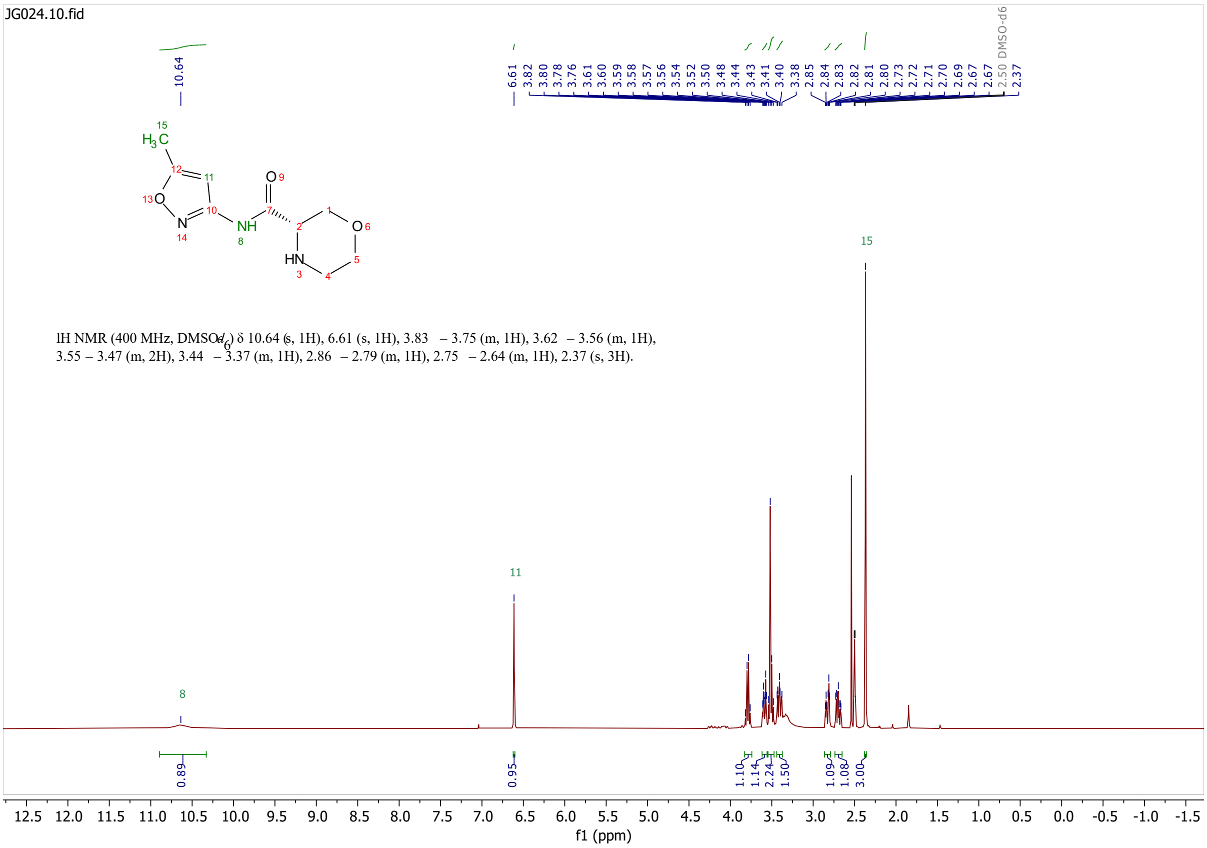
JG024.10.fid
1H NMR (400 MHz, DMSO-$d_6$) δ 10.64 (s, 1H), 6.61 (s, 1H), 3.83 – 3.75 (m, 1H), 3.62 – 3.56 (m, 1H), 3.55 – 3.47 (m, 2H), 3.44 – 3.37 (m, 1H), 2.86 – 2.79 (m, 1H), 2.75 – 2.64 (m, 1H), 2.37 (s, 3H).
f1 (ppm)

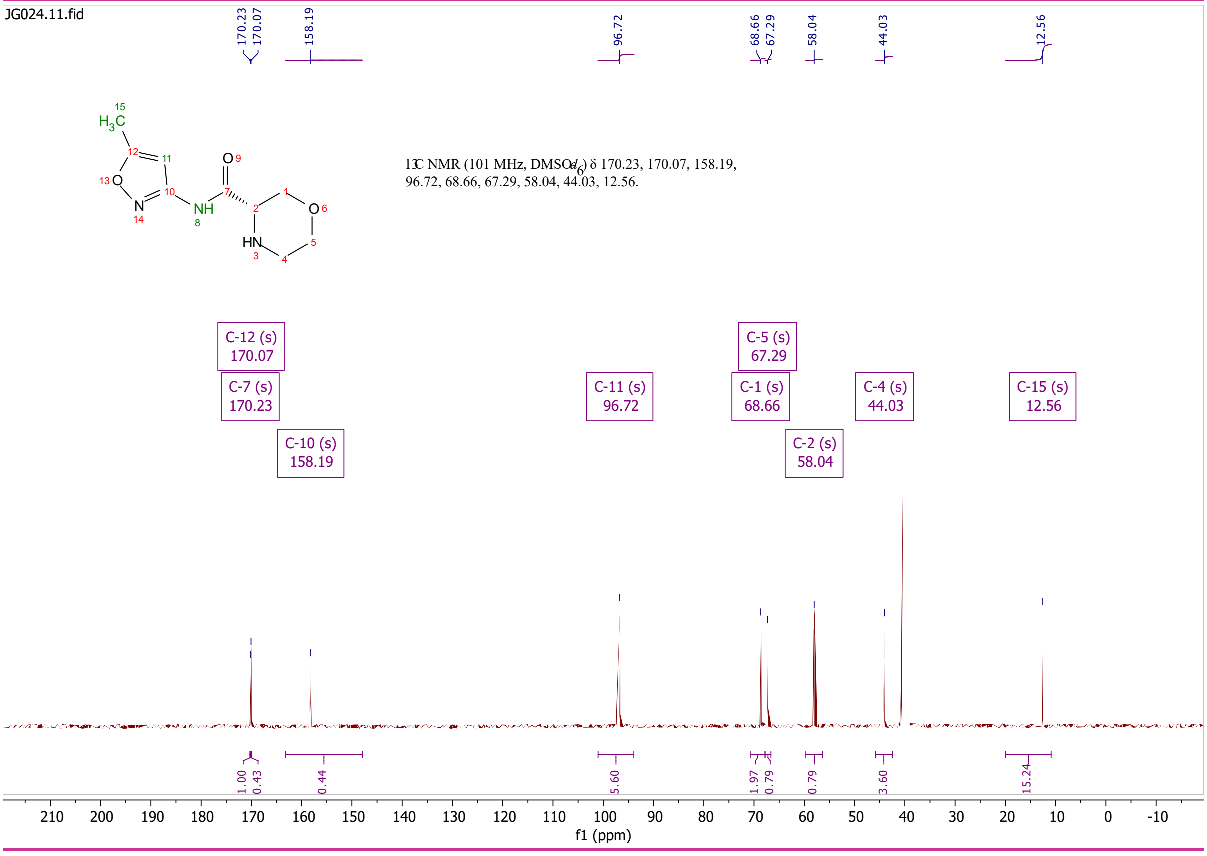
JG024.11.fid
13C NMR (101 MHz, DMSO-$d_6$) δ 170.23, 170.07, 158.19, 96.72, 68.66, 67.29, 58.04, 44.03, 12.56.
C-12 (s) 170.07
C-7 (s) 170.23
C-10 (s) 158.19
C-11 (s) 96.72
C-5 (s) 67.29
C-1 (s) 68.66
C-2 (s) 58.04
C-4 (s) 44.03
C-15 (s) 12.56
f1 (ppm)

# S17

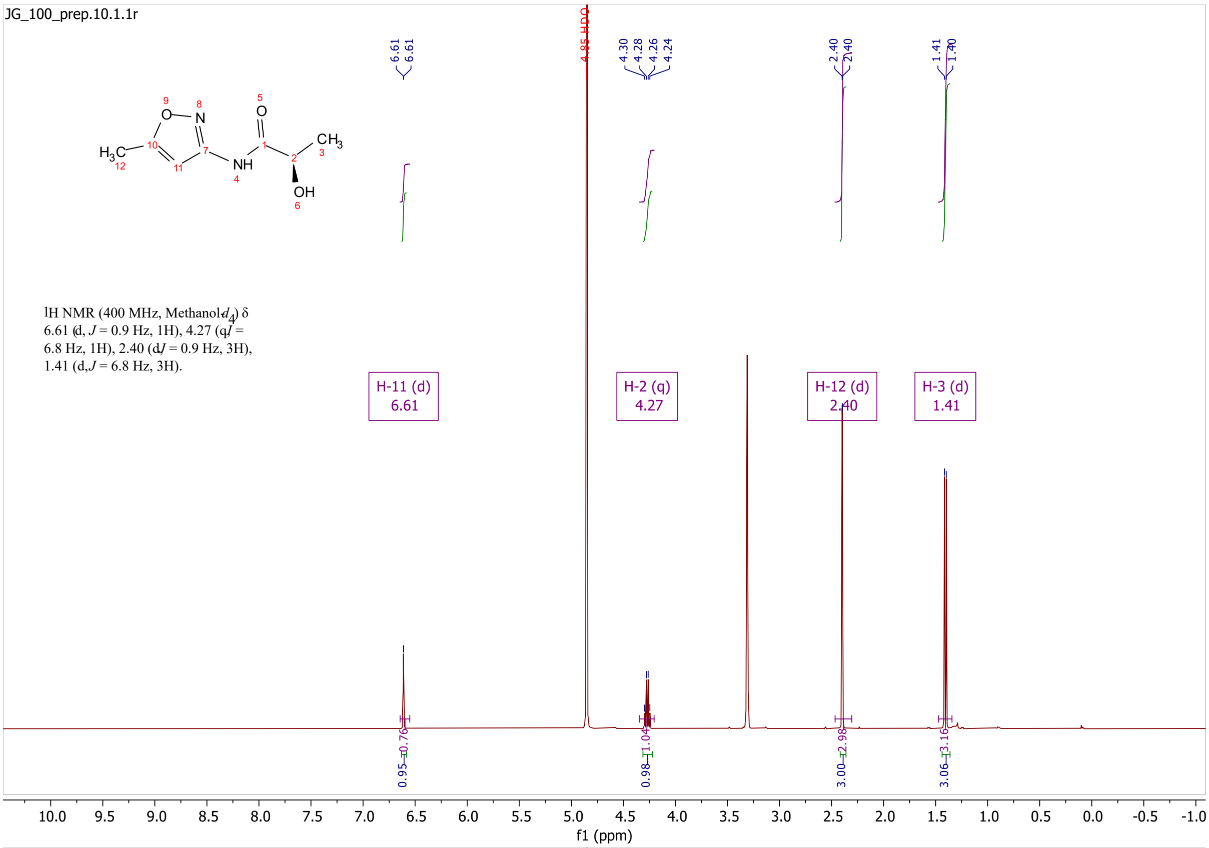
JG_100_prep.10.1.1r
1H NMR (400 MHz, Methanol-$d_4$) δ 6.61 (d, $J$ = 0.9 Hz, 1H), 4.27 (q, $J$ = 6.8 Hz, 1H), 2.40 (d, $J$ = 0.9 Hz, 3H), 1.41 (d, $J$ = 6.8 Hz, 3H).
H-11 (d) 6.61
H-2 (q) 4.27
H-12 (d) 2.40
H-3 (d) 1.41
4.85 HDO
f1 (ppm)

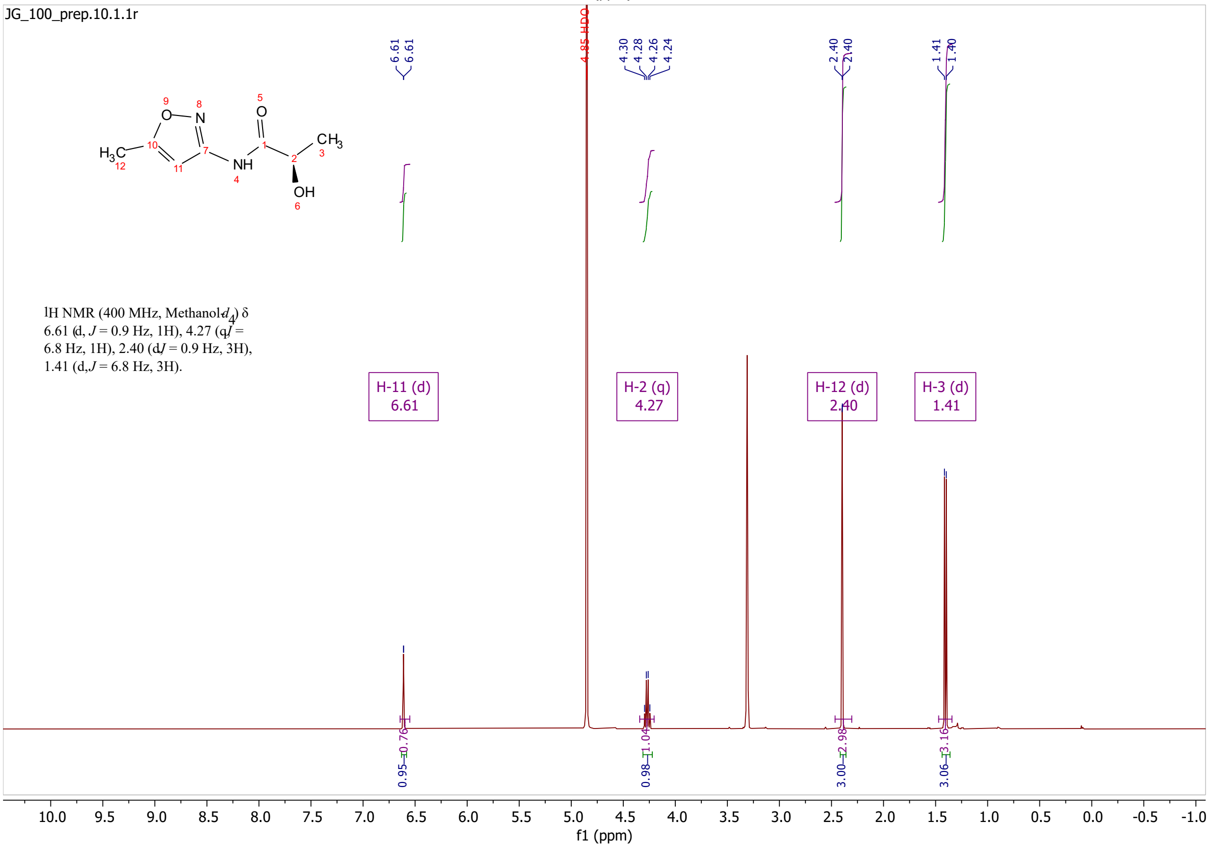
JG_100_prep.10.1.1r
1H NMR (400 MHz, Methanol-$d_4$) δ 6.61 (d, $J$ = 0.9 Hz, 1H), 4.27 (q, $J$ = 6.8 Hz, 1H), 2.40 (d, $J$ = 0.9 Hz, 3H), 1.41 (d, $J$ = 6.8 Hz, 3H).
H-11 (d) 6.61
H-2 (q) 4.27
H-12 (d) 2.40
H-3 (d) 1.41
4.85 HDO
f1 (ppm)

# S18

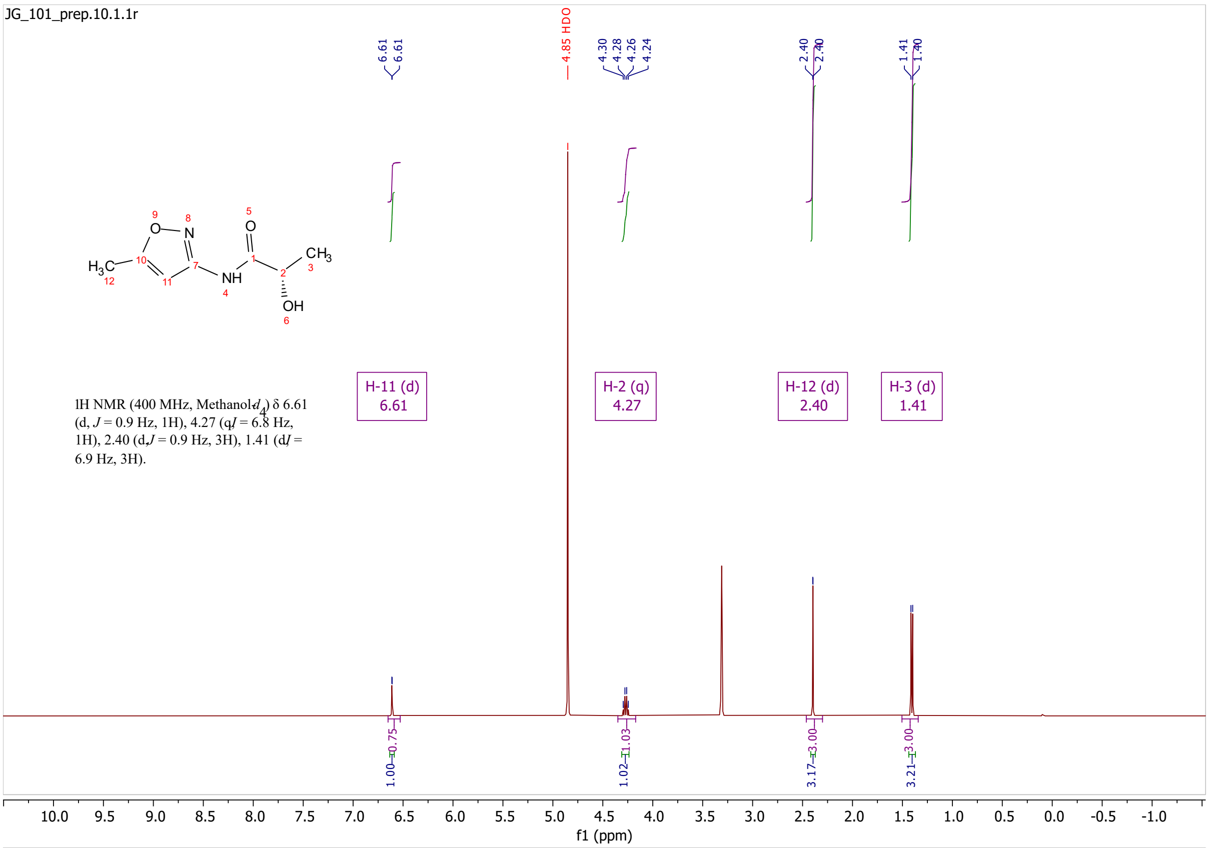

JG_101_prep.10.1.1r
1H NMR (400 MHz, Methanol-$d_4$) δ 6.61 (d, $J$ = 0.9 Hz, 1H), 4.27 (q, $J$ = 6.8 Hz, 1H), 2.40 (d, $J$ = 0.9 Hz, 3H), 1.41 (d, $J$ = 6.9 Hz, 3H).
H-11 (d) 6.61
H-2 (q) 4.27
H-12 (d) 2.40
H-3 (d) 1.41
4.85 HDO
f1 (ppm)

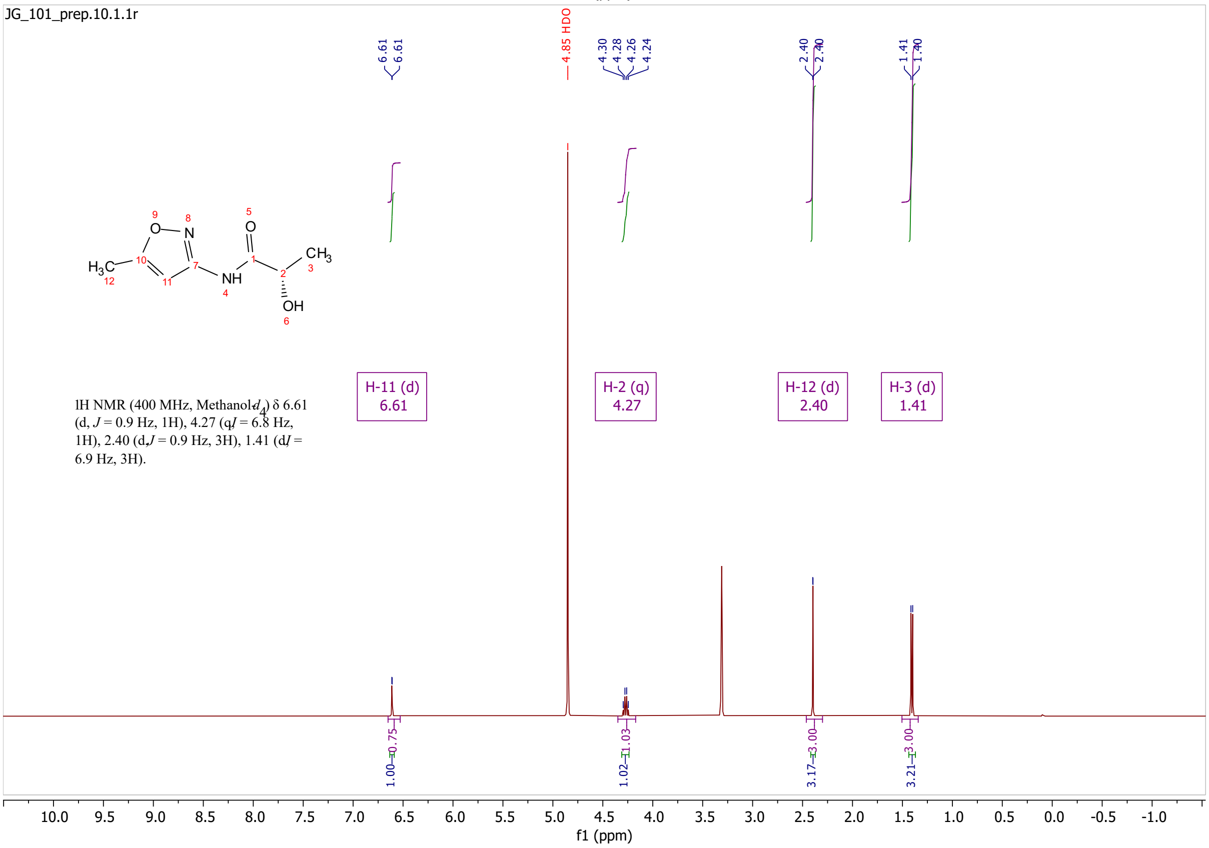

JG_101_prep.10.1.1r
1H NMR (400 MHz, Methanol-$d_4$) δ 6.61 (d, $J$ = 0.9 Hz, 1H), 4.27 (q, $J$ = 6.8 Hz, 1H), 2.40 (d, $J$ = 0.9 Hz, 3H), 1.41 (d, $J$ = 6.9 Hz, 3H).
H-11 (d) 6.61
H-2 (q) 4.27
H-12 (d) 2.40
H-3 (d) 1.41
4.85 HDO
f1 (ppm)

# S19

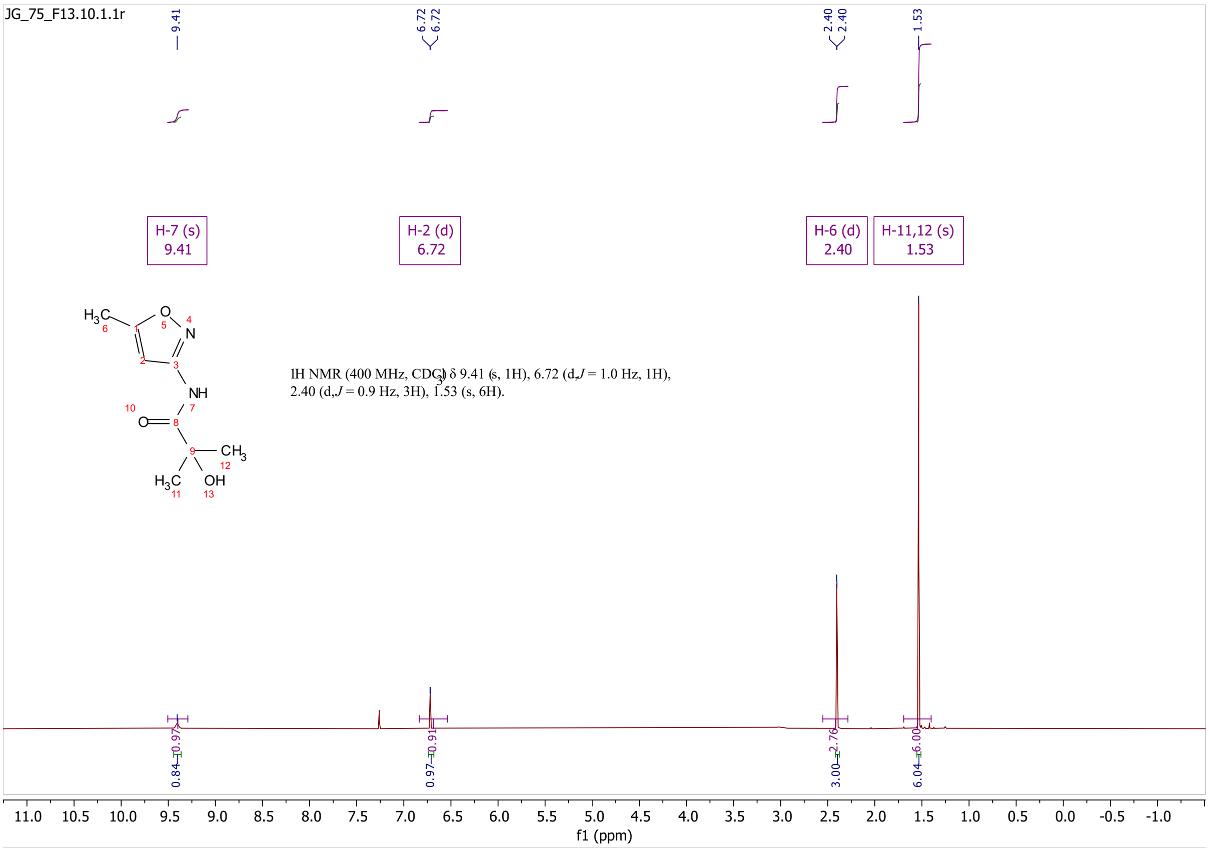
JG_75_F13.10.1.1r
H-7 (s)
9.41
H-2 (d)
6.72
H-6 (d)
2.40
H-11,12 (s)
1.53
1H NMR (400 MHz, $CDCl_3$) δ 9.41 (s, 1H), 6.72 (d, J = 1.0 Hz, 1H),
2.40 (d, J = 0.9 Hz, 3H), 1.53 (s, 6H).
f1 (ppm)

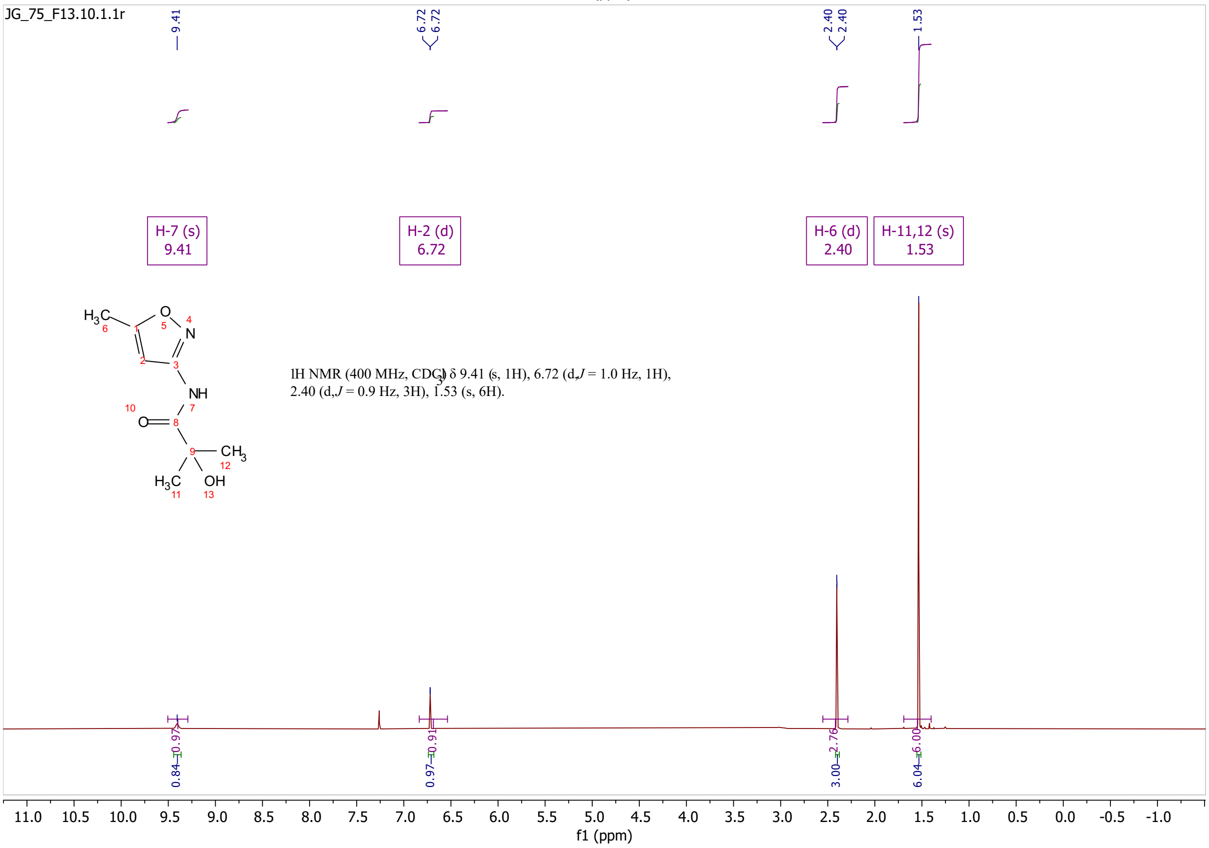
JG_75_F13.10.1.1r
H-7 (s)
9.41
H-2 (d)
6.72
H-6 (d)
2.40
H-11,12 (s)
1.53
1H NMR (400 MHz, $CDCl_3$) δ 9.41 (s, 1H), 6.72 (d, J = 1.0 Hz, 1H),
2.40 (d, J = 0.9 Hz, 3H), 1.53 (s, 6H).
f1 (ppm)

# S20

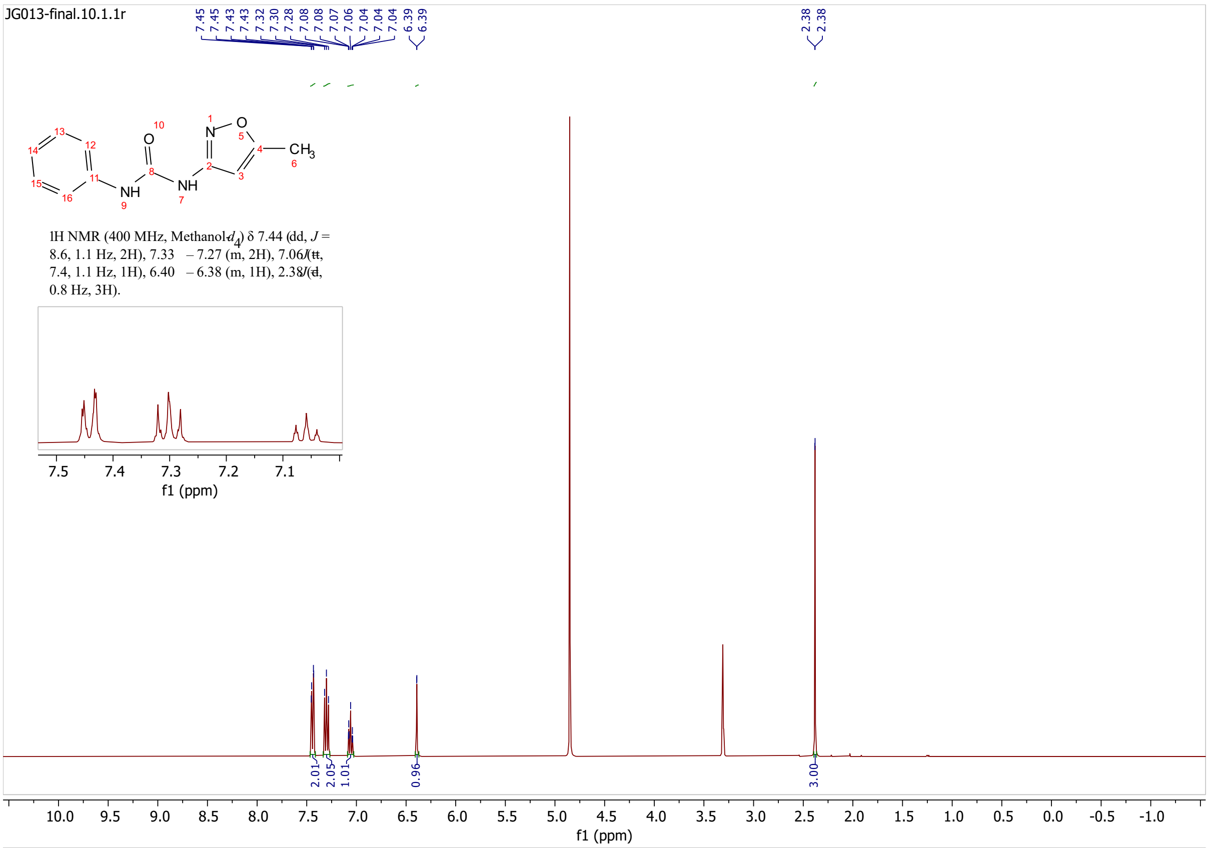
JG013-final.10.1.1r
CH3
NH
NH
1H NMR (400 MHz, Methanol-$d_4$) δ 7.44 (dd, $J$ = 8.6, 1.1 Hz, 2H), 7.33 – 7.27 (m, 2H), 7.06 (tt, 7.4, 1.1 Hz, 1H), 6.40 – 6.38 (m, 1H), 2.38 (d, 0.8 Hz, 3H).
f1 (ppm)
f1 (ppm)

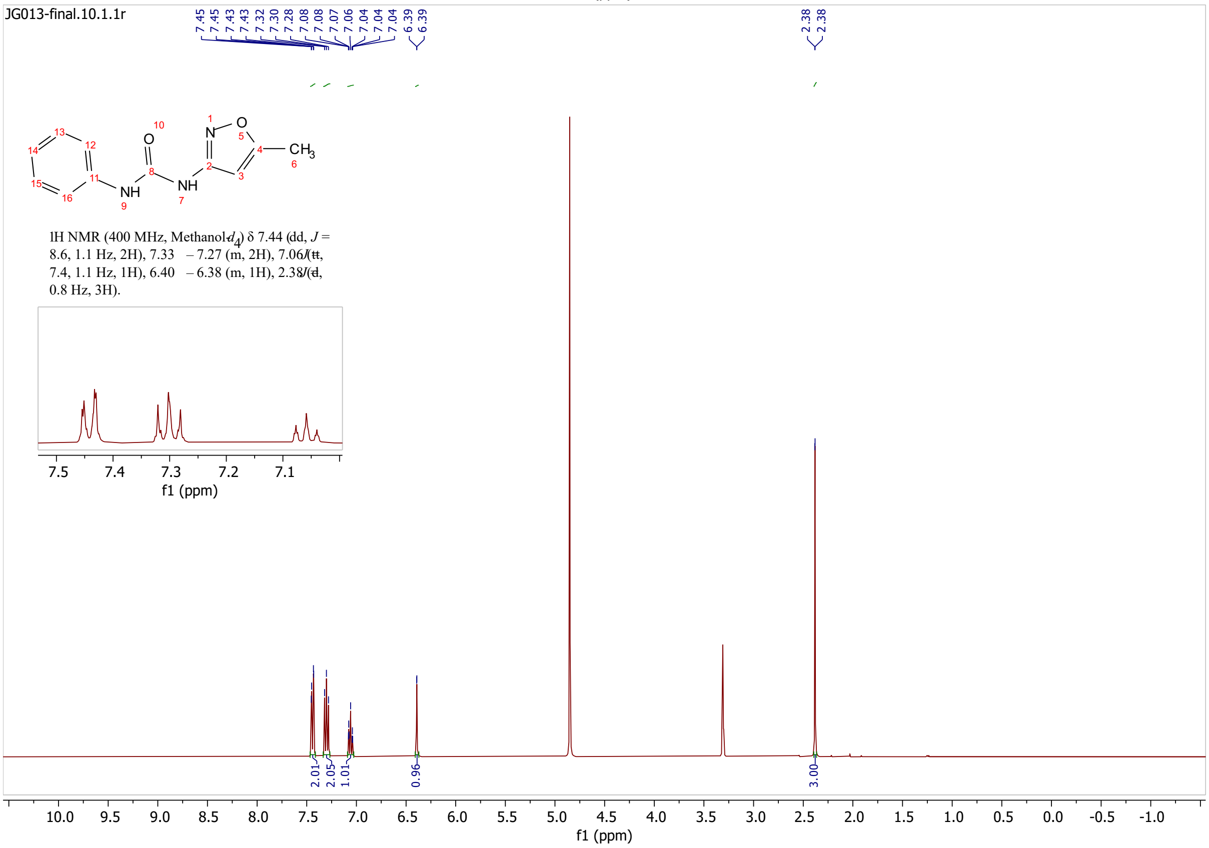
JG013-final.10.1.1r
CH3
NH
NH
1H NMR (400 MHz, Methanol-$d_4$) δ 7.44 (dd, $J$ = 8.6, 1.1 Hz, 2H), 7.33 – 7.27 (m, 2H), 7.06 (tt, 7.4, 1.1 Hz, 1H), 6.40 – 6.38 (m, 1H), 2.38 (d, 0.8 Hz, 3H).
f1 (ppm)
f1 (ppm)

# S21

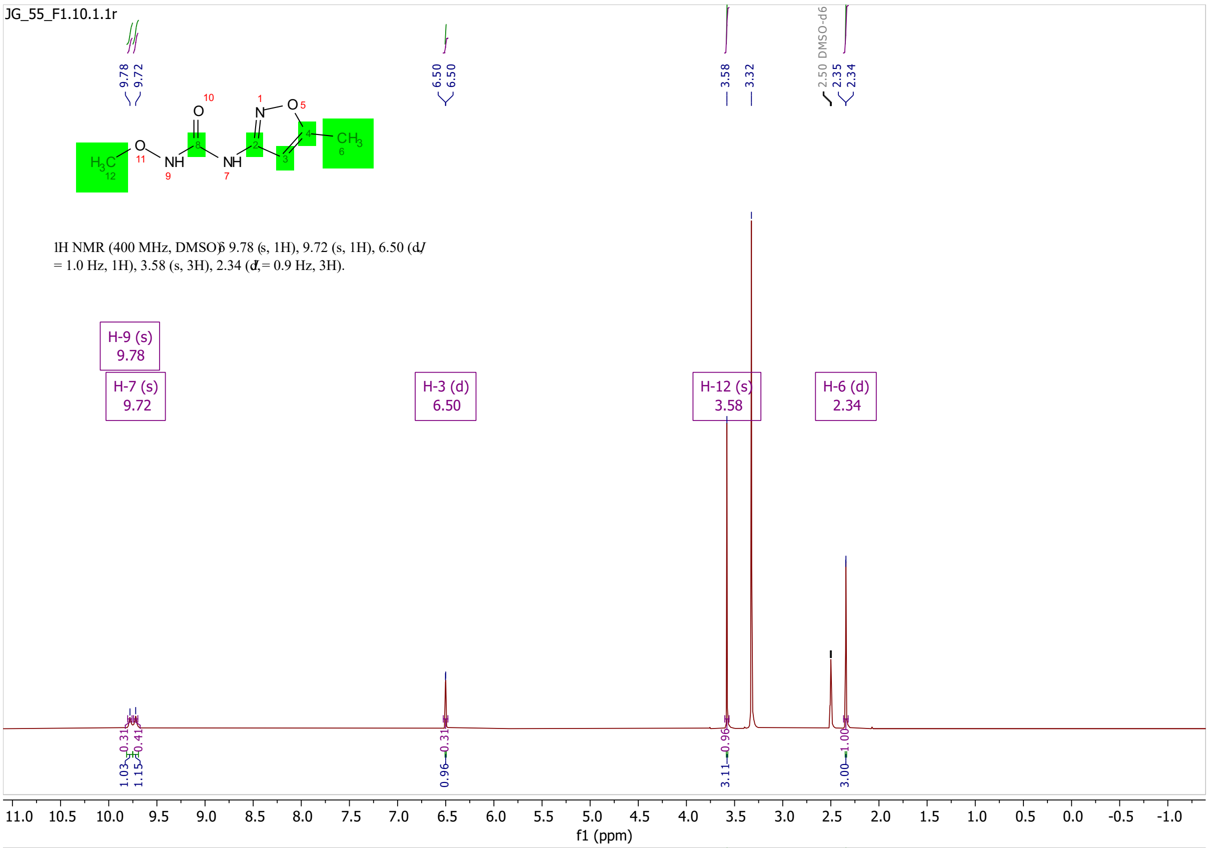

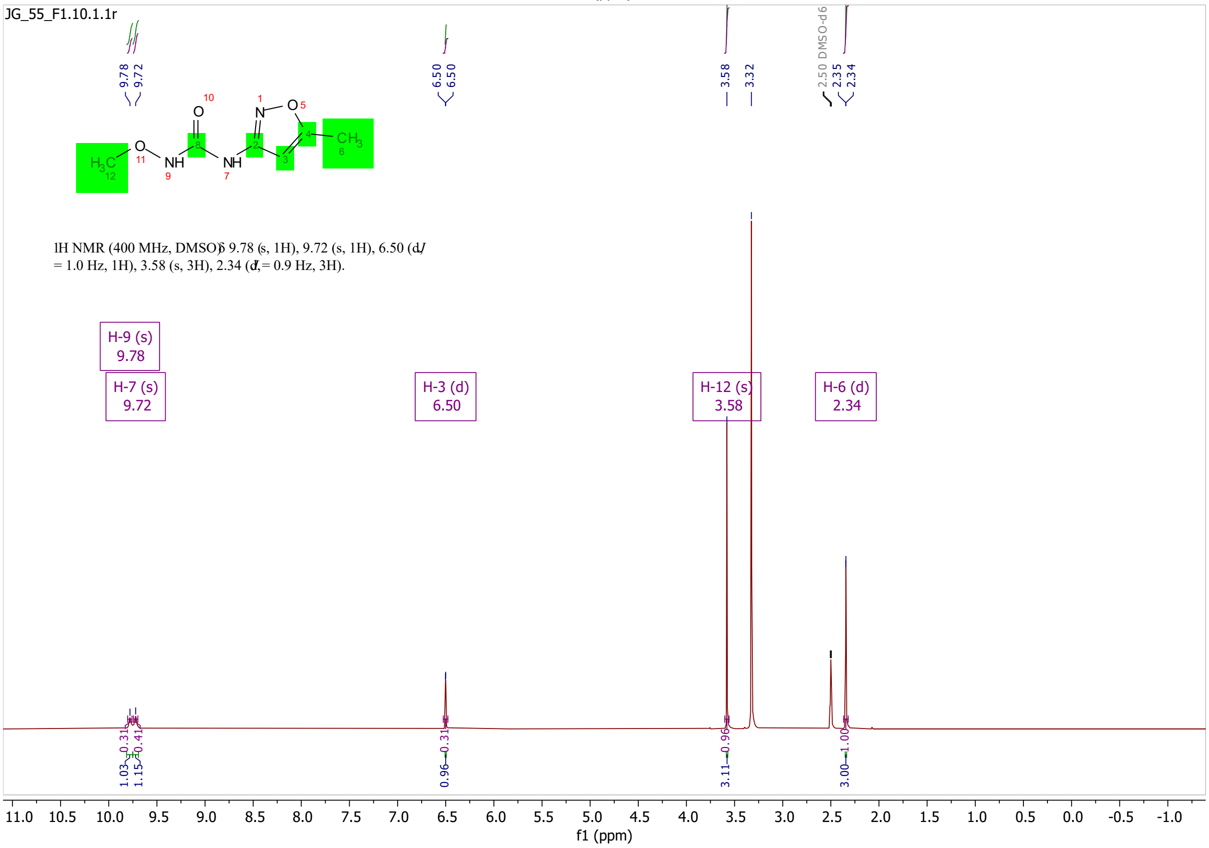
JG_55_F1.10.1.1r
IH NMR (400 MHz, DMSO) δ 9.78 (s, 1H), 9.72 (s, 1H), 6.50 (d, J = 1.0 Hz, 1H), 3.58 (s, 3H), 2.34 (d, J = 0.9 Hz, 3H).
H-9 (s) 9.78
H-7 (s) 9.72
H-3 (d) 6.50
H-12 (s) 3.58
H-6 (d) 2.34
2.50 DMSO-d6
f1 (ppm)

# S22

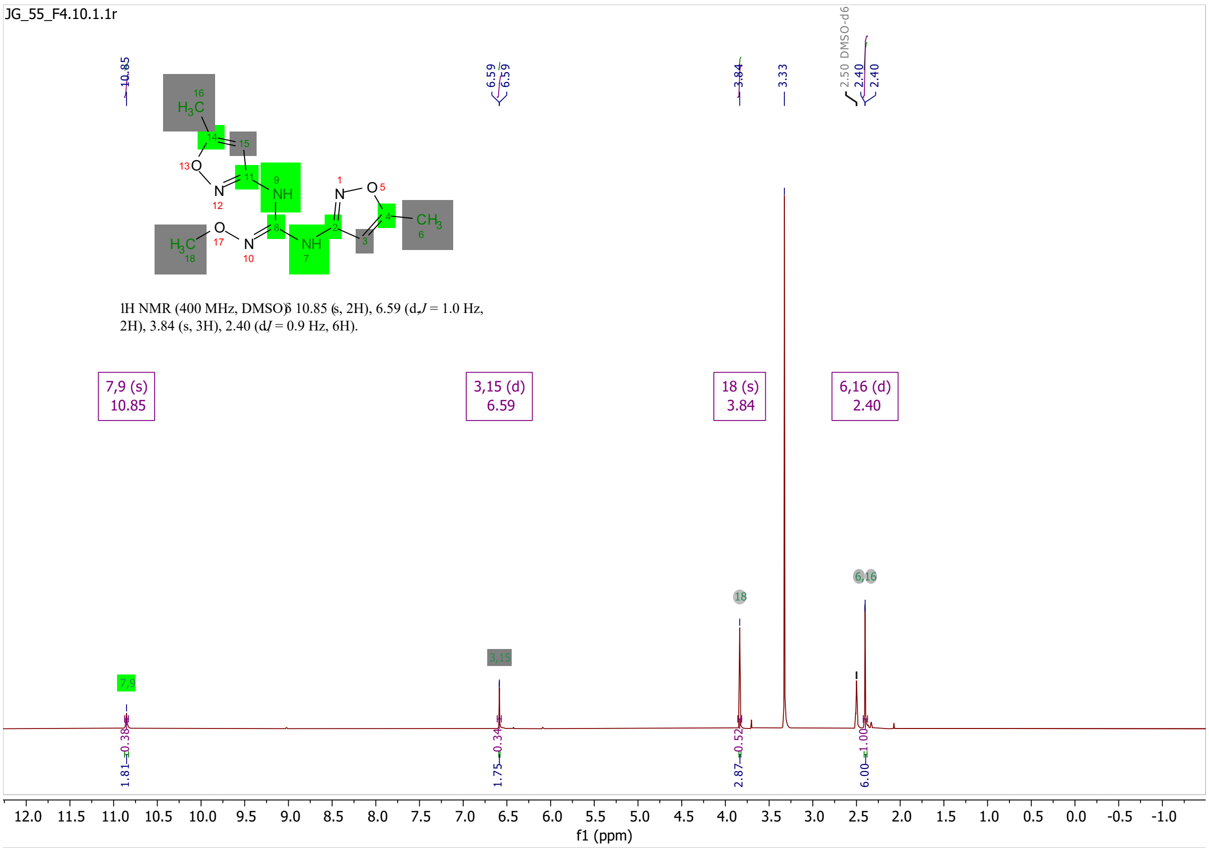
JG_55_F4.10.1.1r
1H NMR (400 MHz, DMSO) δ 10.85 (s, 2H), 6.59 (d, J = 1.0 Hz, 2H), 3.84 (s, 3H), 2.40 (d, J = 0.9 Hz, 6H).
7,9 (s) 10.85
3,15 (d) 6.59
18 (s) 3.84
6,16 (d) 2.40
2.50 DMSO-d6
f1 (ppm)

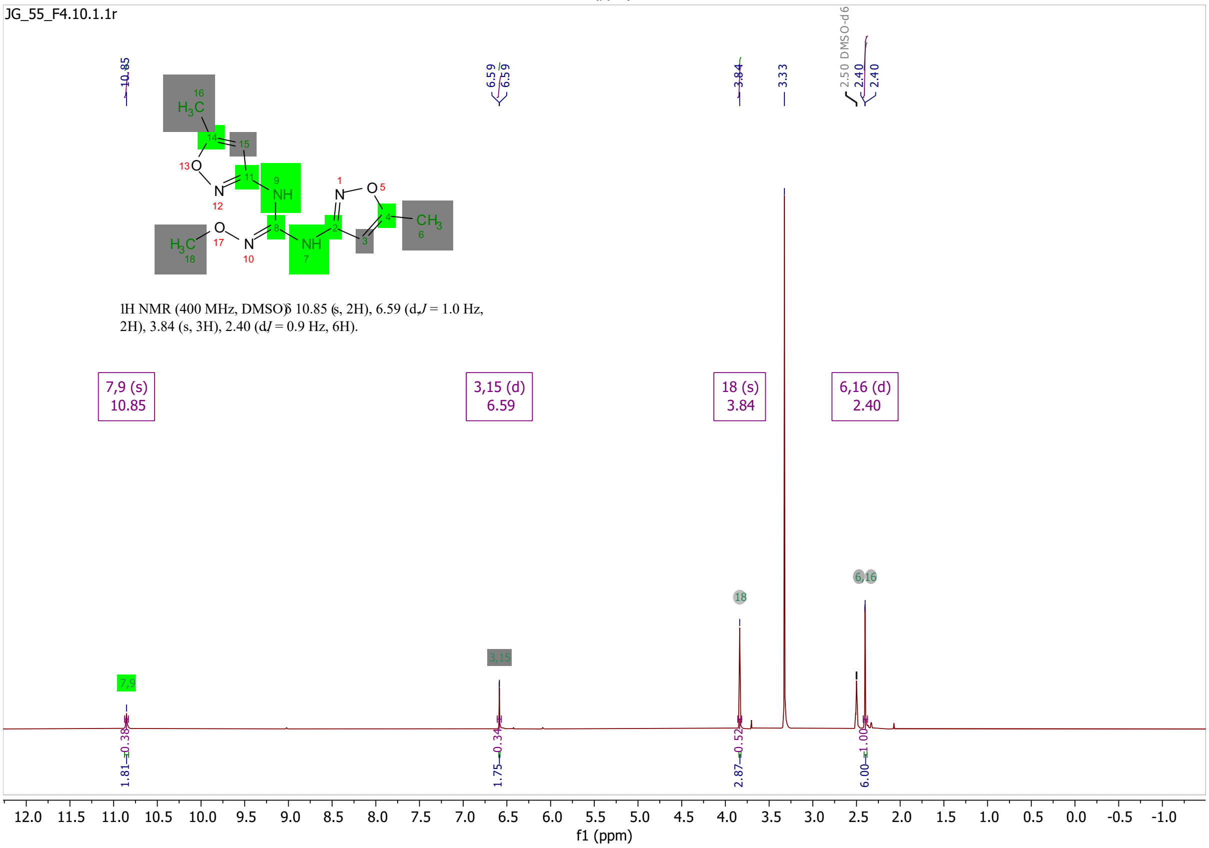
JG_55_F4.10.1.1r
1H NMR (400 MHz, DMSO) δ 10.85 (s, 2H), 6.59 (d, J = 1.0 Hz, 2H), 3.84 (s, 3H), 2.40 (d, J = 0.9 Hz, 6H).
7,9 (s) 10.85
3,15 (d) 6.59
18 (s) 3.84
6,16 (d) 2.40
2.50 DMSO-d6
f1 (ppm)

**S23**

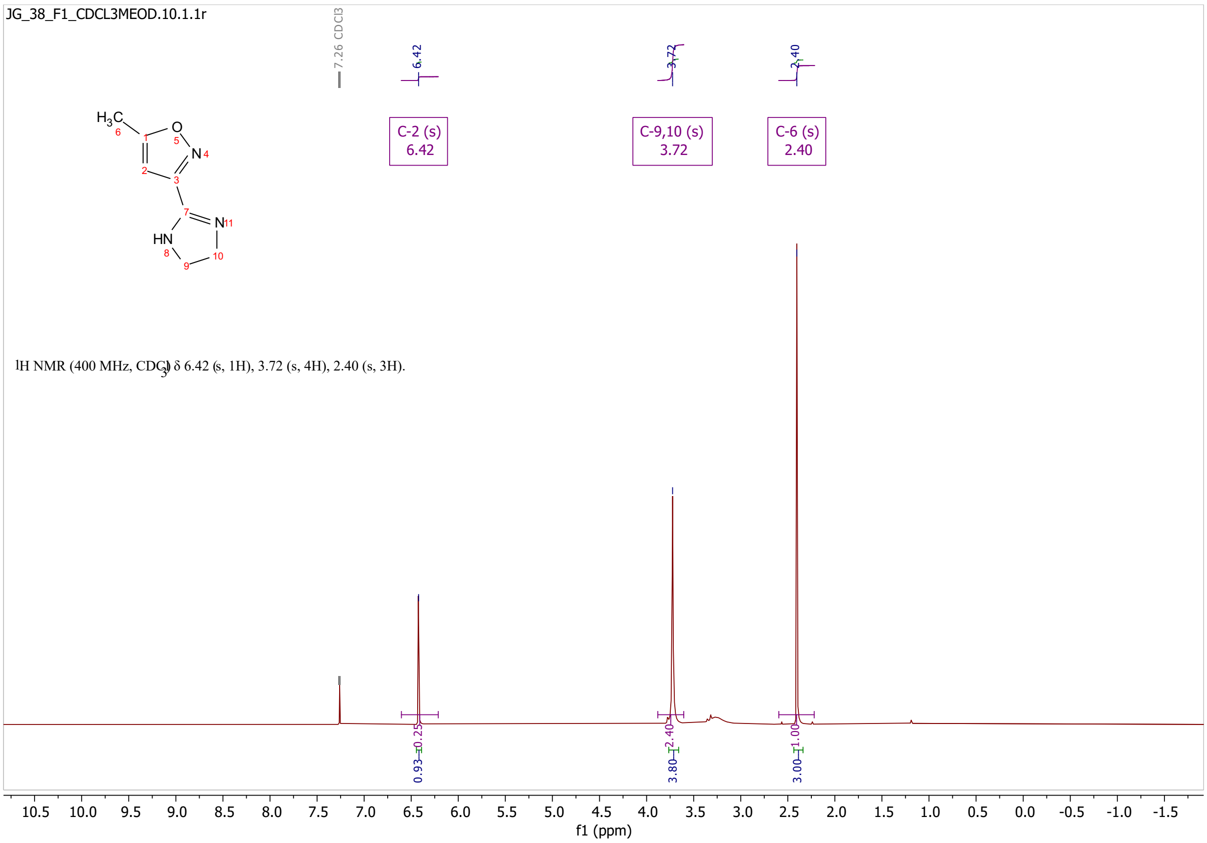
JG_38_F1_CDCL3MEOD.10.1.1r
7.26 CDCl3
6.42
3.72
2.40
C-2 (s)
6.42
C-9,10 (s)
3.72
C-6 (s)
2.40
1H NMR (400 MHz, CDCl3) δ 6.42 (s, 1H), 3.72 (s, 4H), 2.40 (s, 3H).
0.93
0.25
3.80
2.40
3.00
1.00
10.5 10.0 9.5 9.0 8.5 8.0 7.5 7.0 6.5 6.0 5.5 5.0 4.5 4.0 3.5 3.0 2.5 2.0 1.5 1.0 0.5 0.0 -0.5 -1.0 -1.5
f1 (ppm)

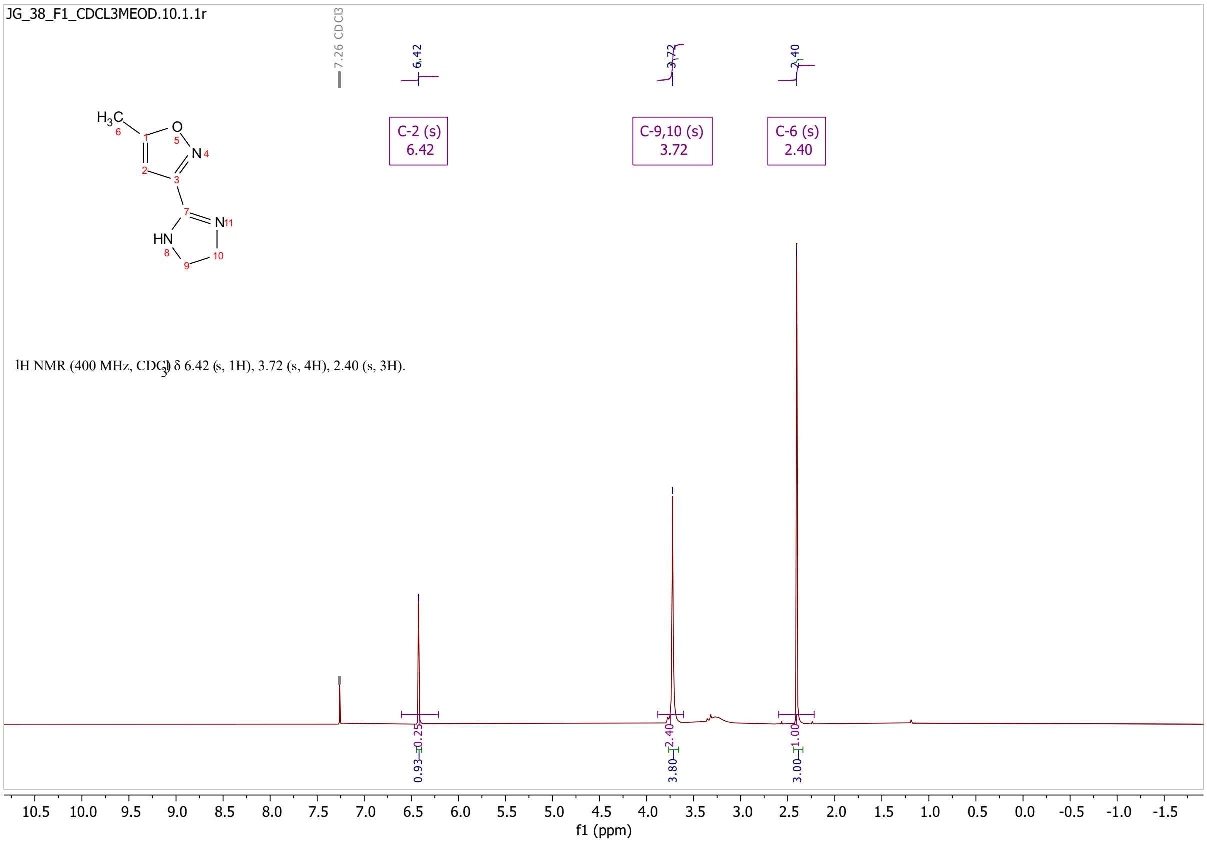
JG_38_F1_CDCL3MEOD.10.1.1r
7.26 CDCl3
6.42
3.72
2.40
C-2 (s)
6.42
C-9,10 (s)
3.72
C-6 (s)
2.40
1H NMR (400 MHz, CDCl3) δ 6.42 (s, 1H), 3.72 (s, 4H), 2.40 (s, 3H).
0.93
0.25
3.80
2.40
3.00
1.00
10.5 10.0 9.5 9.0 8.5 8.0 7.5 7.0 6.5 6.0 5.5 5.0 4.5 4.0 3.5 3.0 2.5 2.0 1.5 1.0 0.5 0.0 -0.5 -1.0 -1.5
f1 (ppm)

# S24

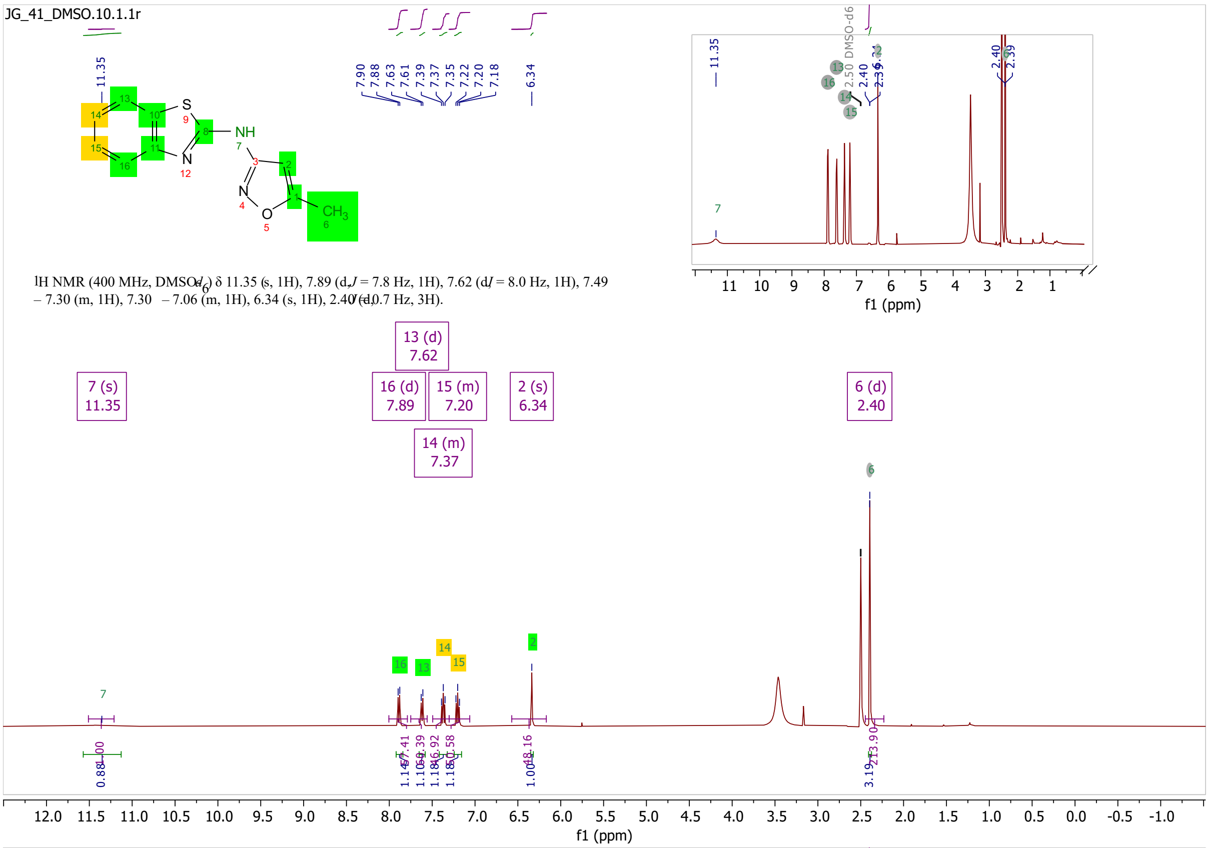
JG_41_DMSO.10.1.1r
$^{1}$H NMR (400 MHz, DMSO-$d_6$) δ 11.35 (s, 1H), 7.89 (d, $J$ = 7.8 Hz, 1H), 7.62 (d, $J$ = 8.0 Hz, 1H), 7.49 – 7.30 (m, 1H), 7.30 – 7.06 (m, 1H), 6.34 (s, 1H), 2.40 (d, $J$ = 0.7 Hz, 3H).
f1 (ppm)

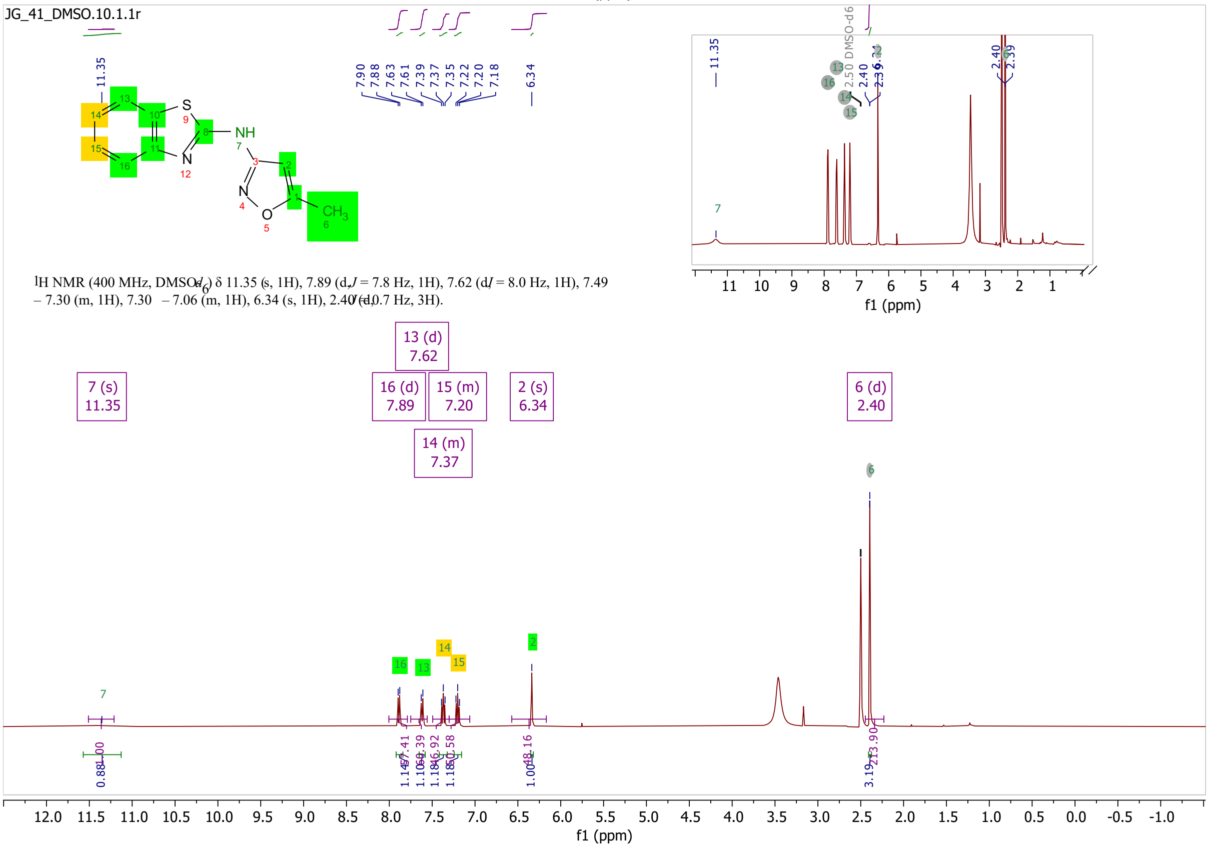

JG_41_DMSO.10.1.1r
$^{1}$H NMR (400 MHz, DMSO-$d_6$) δ 11.35 (s, 1H), 7.89 (d, $J$ = 7.8 Hz, 1H), 7.62 (d, $J$ = 8.0 Hz, 1H), 7.49 – 7.30 (m, 1H), 7.30 – 7.06 (m, 1H), 6.34 (s, 1H), 2.40 (d, $J$ = 0.7 Hz, 3H).
f1 (ppm)

# S54

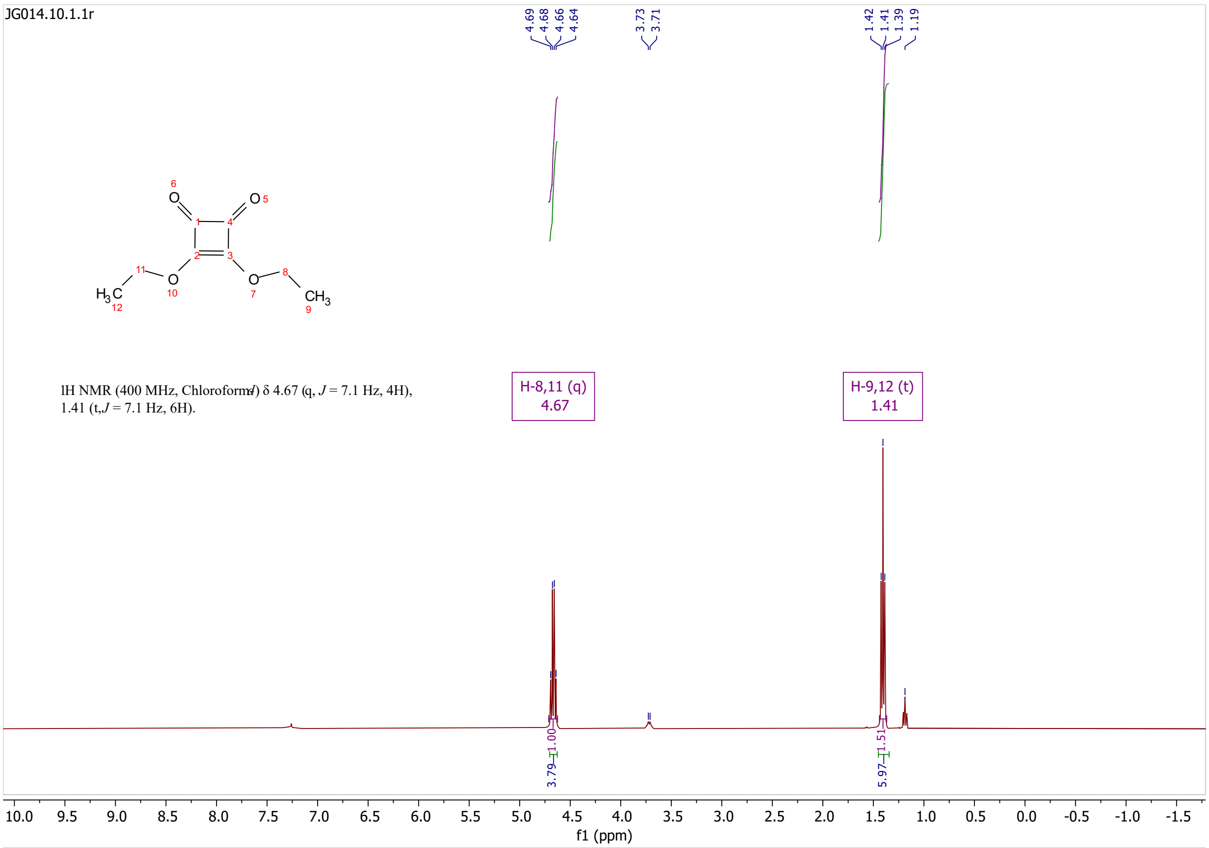

JG014.10.1.1r
4.69
4.68
4.66
4.64
3.73
3.71
1.42
1.41
1.39
1.19
1H NMR (400 MHz, Chloroform-d) δ 4.67 (q, J = 7.1 Hz, 4H), 1.41 (t, J = 7.1 Hz, 6H).
H-8,11 (q)
4.67
H-9,12 (t)
1.41
3.79
1.00
5.97
1.51
10.0 9.5 9.0 8.5 8.0 7.5 7.0 6.5 6.0 5.5 5.0 4.5 4.0 3.5 3.0 2.5 2.0 1.5 1.0 0.5 0.0 -0.5 -1.0 -1.5
f1 (ppm)

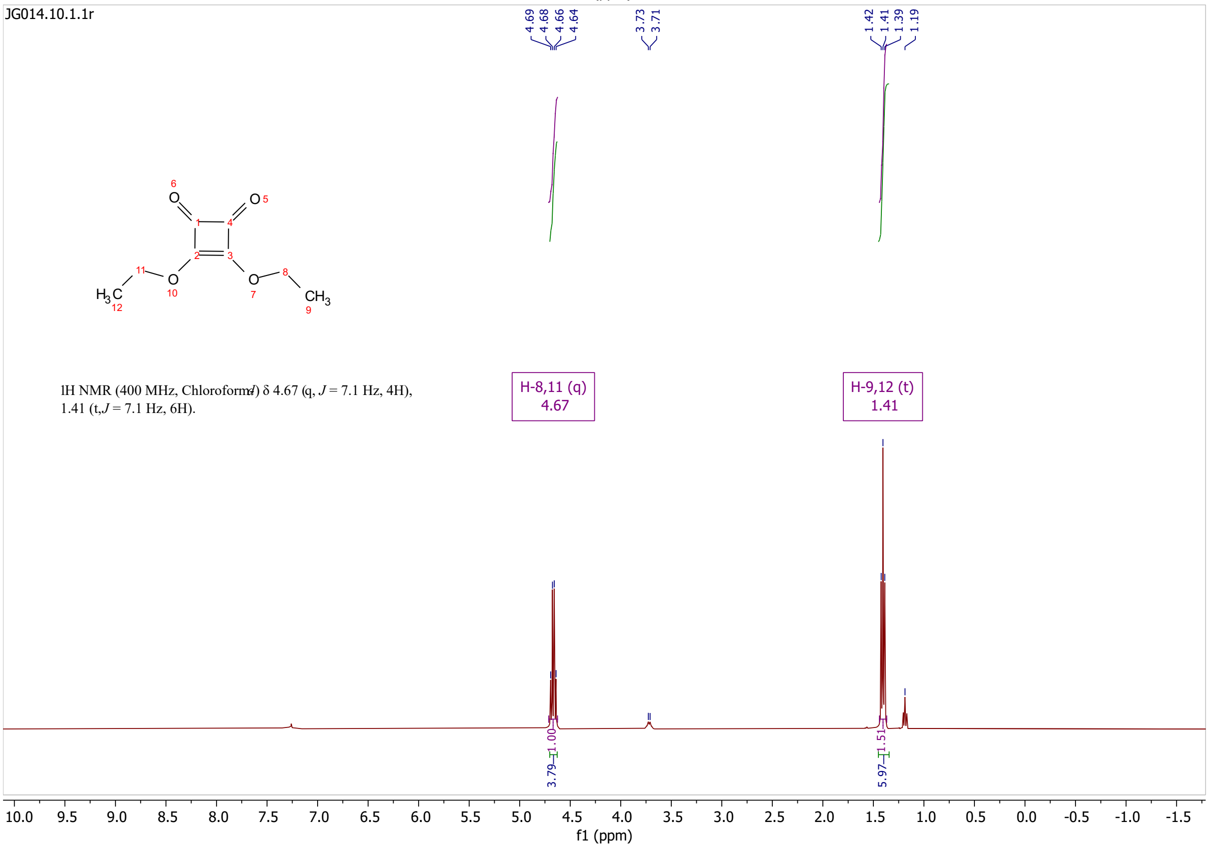

JG014.10.1.1r
1H NMR (400 MHz, Chloroform-d) δ 4.67 (q, J = 7.1 Hz, 4H), 1.41 (t, J = 7.1 Hz, 6H).
H-8,11 (q)
4.67
H-9,12 (t)
1.41
f1 (ppm)

# S55

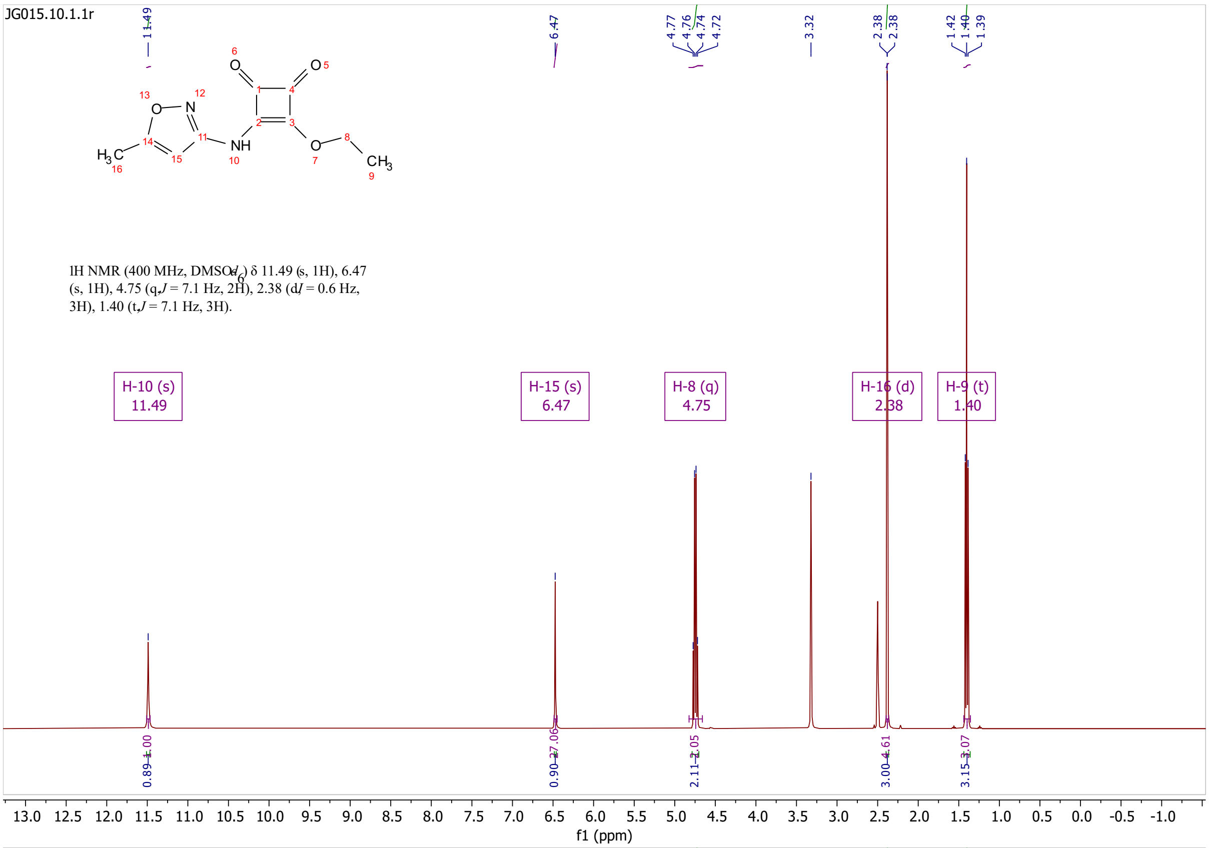
JG015.10.1.1r
1H NMR (400 MHz, DMSO-$d_6$) δ 11.49 (s, 1H), 6.47 (s, 1H), 4.75 (q, J = 7.1 Hz, 2H), 2.38 (d, J = 0.6 Hz, 3H), 1.40 (t, J = 7.1 Hz, 3H).
H-10 (s) 11.49
H-15 (s) 6.47
H-8 (q) 4.75
H-16 (d) 2.38
H-9 (t) 1.40
f1 (ppm)

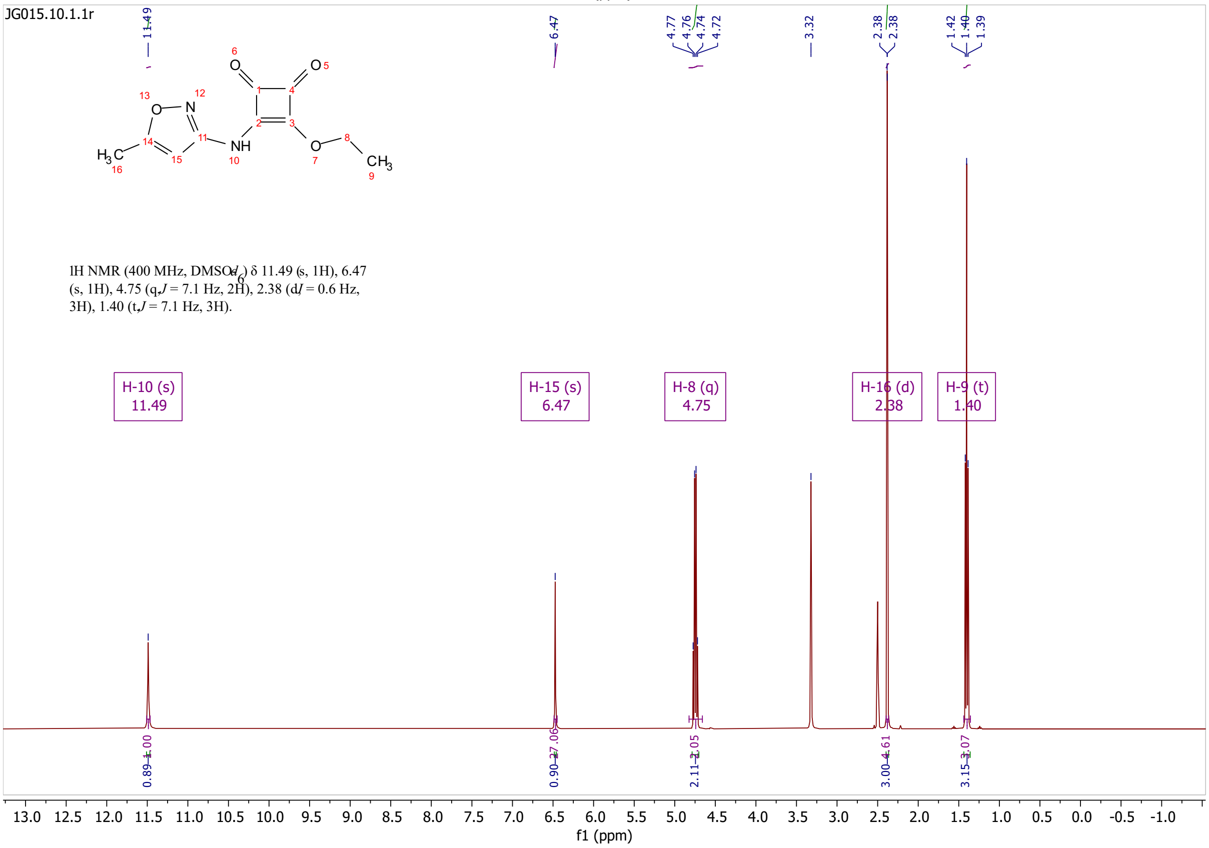
JG015.10.1.1r
1H NMR (400 MHz, DMSO-$d_6$) δ 11.49 (s, 1H), 6.47 (s, 1H), 4.75 (q, J = 7.1 Hz, 2H), 2.38 (d, J = 0.6 Hz, 3H), 1.40 (t, J = 7.1 Hz, 3H).
H-10 (s) 11.49
H-15 (s) 6.47
H-8 (q) 4.75
H-16 (d) 2.38
H-9 (t) 1.40
f1 (ppm)

# S25

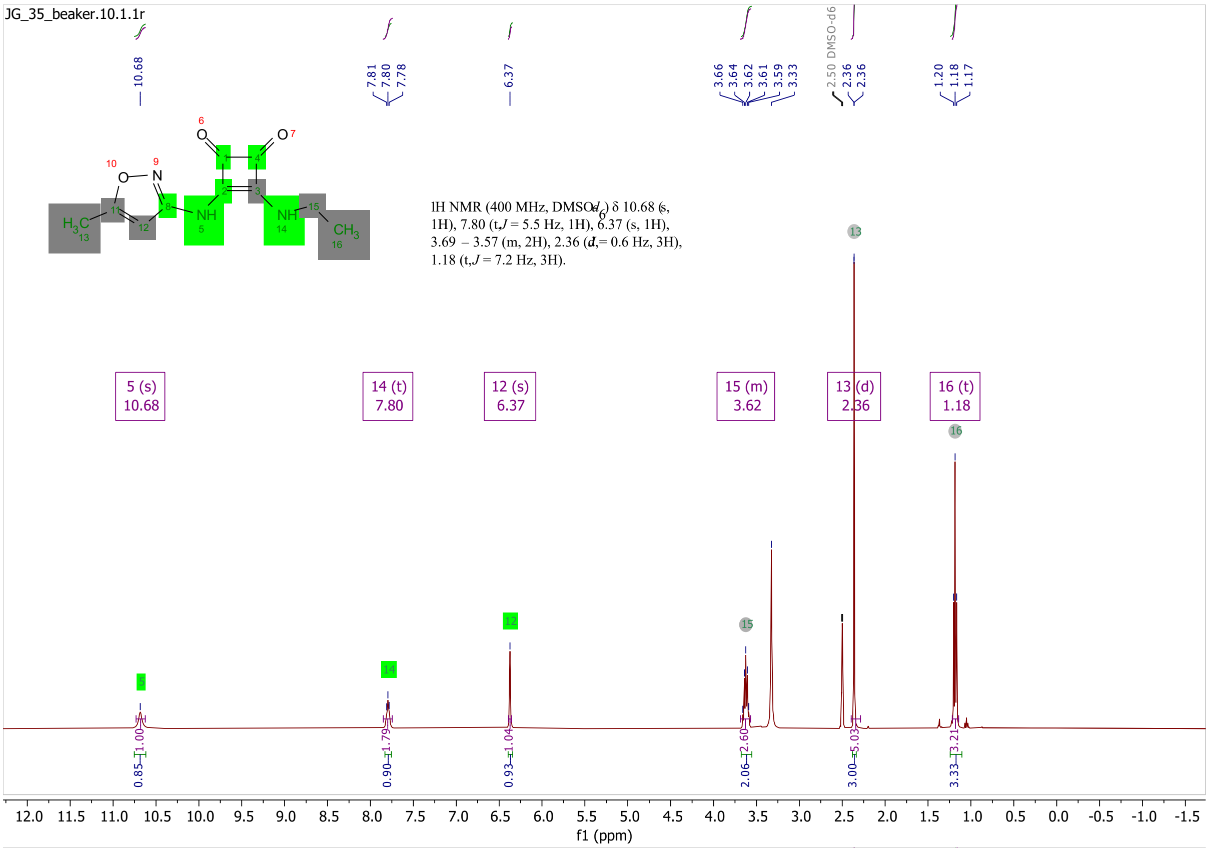
JG_35_beaker.10.1.1r
1H NMR (400 MHz, DMSO-d6) δ 10.68 (s, 1H), 7.80 (t, J = 5.5 Hz, 1H), 6.37 (s, 1H), 3.69 – 3.57 (m, 2H), 2.36 (d, J = 0.6 Hz, 3H), 1.18 (t, J = 7.2 Hz, 3H).
5 (s) 10.68
14 (t) 7.80
12 (s) 6.37
15 (m) 3.62
13 (d) 2.36
16 (t) 1.18
2.50 DMSO-d6
f1 (ppm)

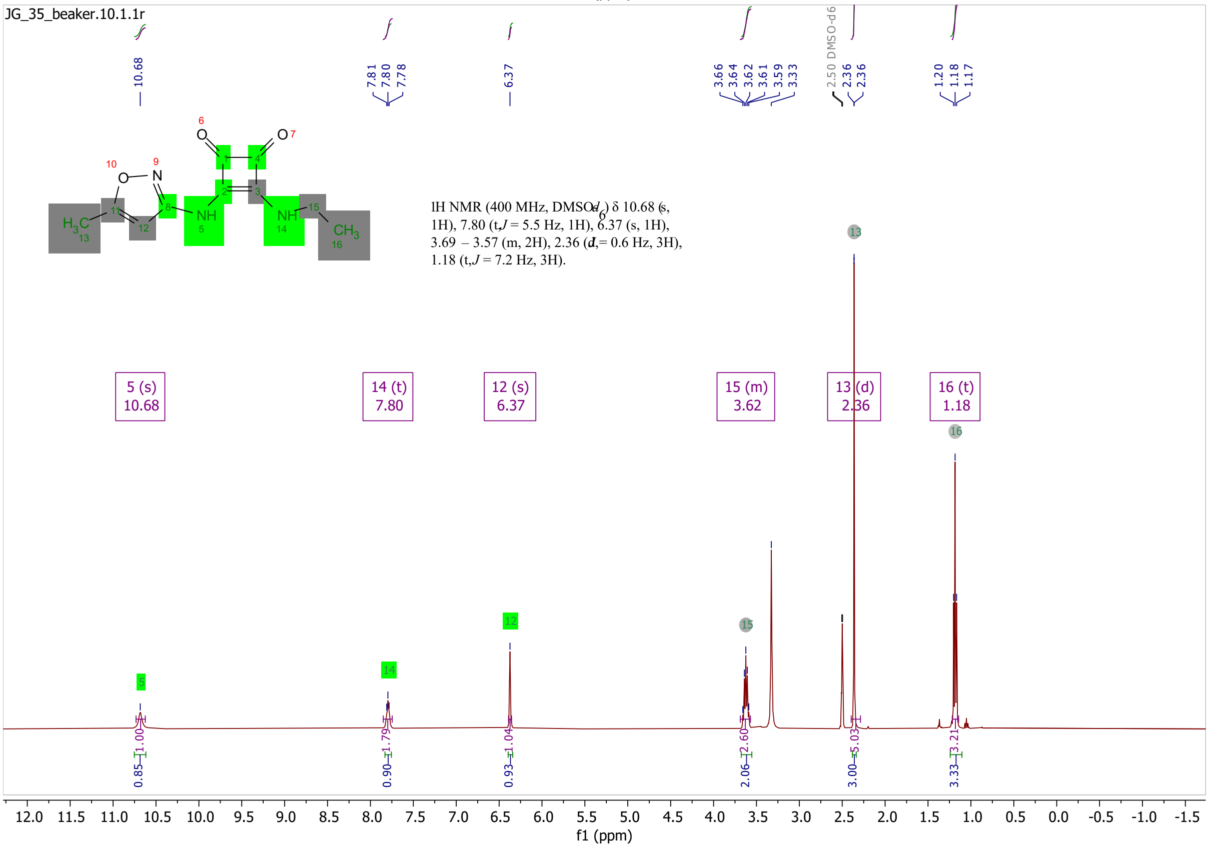
JG_35_beaker.10.1.1r
1H NMR (400 MHz, DMSO-d6) δ 10.68 (s, 1H), 7.80 (t, J = 5.5 Hz, 1H), 6.37 (s, 1H), 3.69 – 3.57 (m, 2H), 2.36 (d, J = 0.6 Hz, 3H), 1.18 (t, J = 7.2 Hz, 3H).
5 (s) 10.68
14 (t) 7.80
12 (s) 6.37
15 (m) 3.62
13 (d) 2.36
16 (t) 1.18
2.50 DMSO-d6
f1 (ppm)

# S56

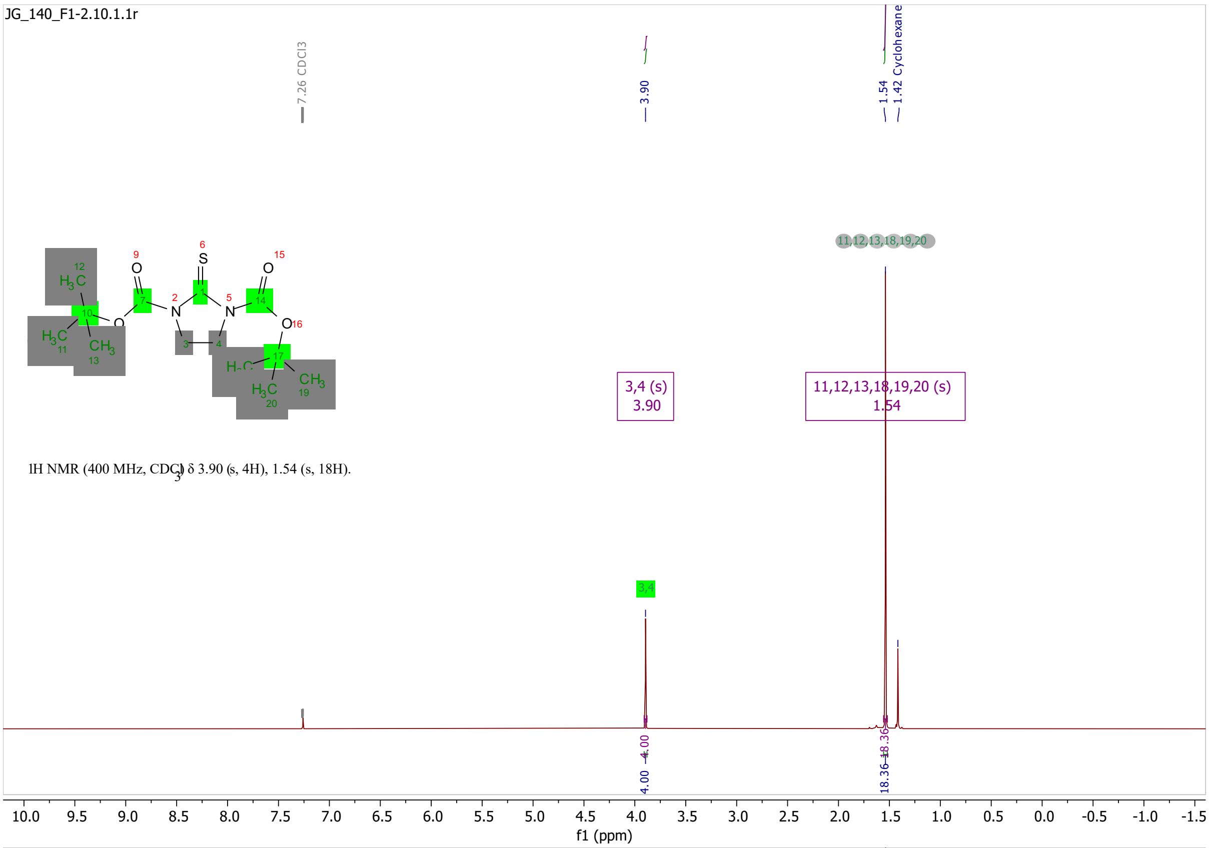
JG_140_F1-2.10.1.1r
7.26 CDCl3
3.90
1.54
1.42 Cyclohexane
3,4 (s)
3.90
11,12,13,18,19,20 (s)
1.54
1H NMR (400 MHz, CDCl3) δ 3.90 (s, 4H), 1.54 (s, 18H).
4.00
18.36
f1 (ppm)

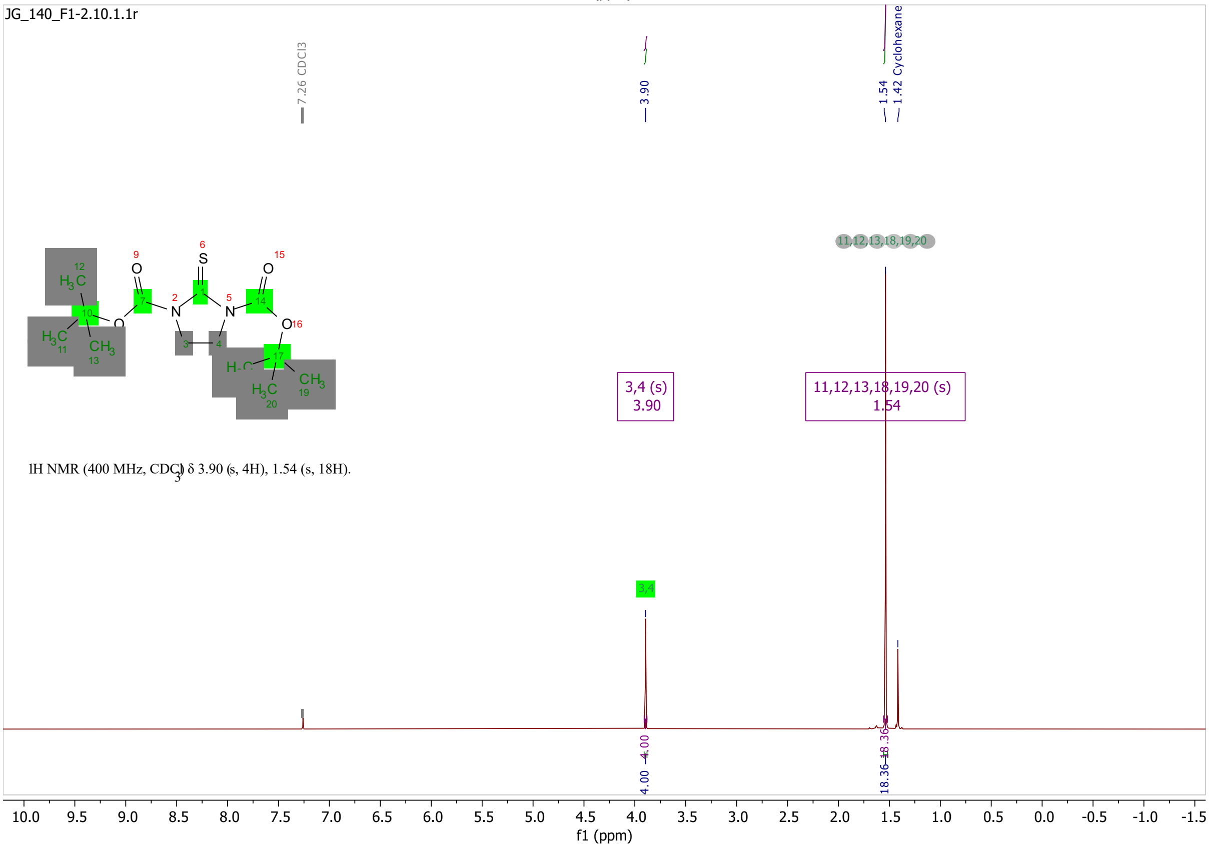
JG_140_F1-2.10.1.1r
7.26 CDCl3
3.90
1.54
1.42 Cyclohexane
3,4 (s)
3.90
11,12,13,18,19,20 (s)
1.54
1H NMR (400 MHz, CDCl3) δ 3.90 (s, 4H), 1.54 (s, 18H).
4.00
18.36
f1 (ppm)

# S57

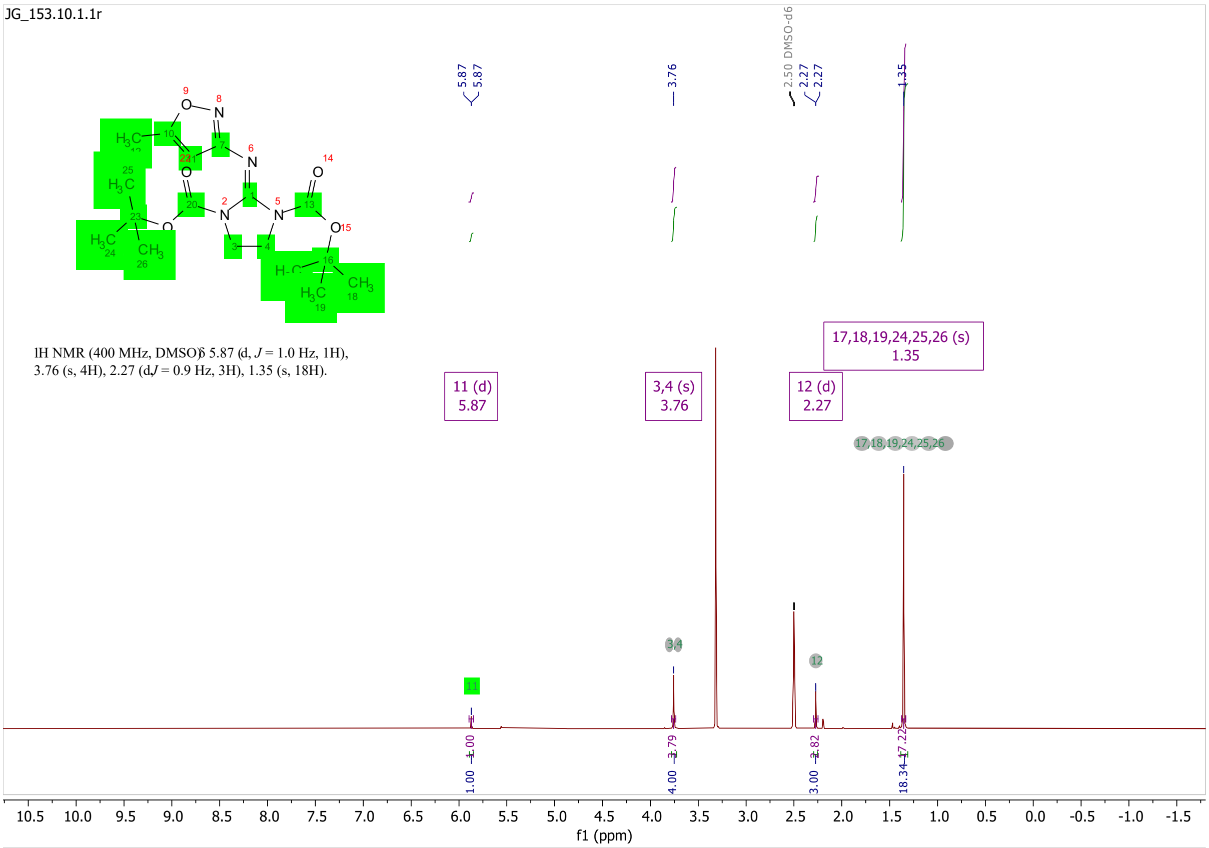
JG_153.10.1.1r
1H NMR (400 MHz, DMSO) δ 5.87 (d, J = 1.0 Hz, 1H), 3.76 (s, 4H), 2.27 (d, J = 0.9 Hz, 3H), 1.35 (s, 18H).
11 (d) 5.87
3,4 (s) 3.76
12 (d) 2.27
17,18,19,24,25,26 (s) 1.35
2.50 DMSO-d6
f1 (ppm)

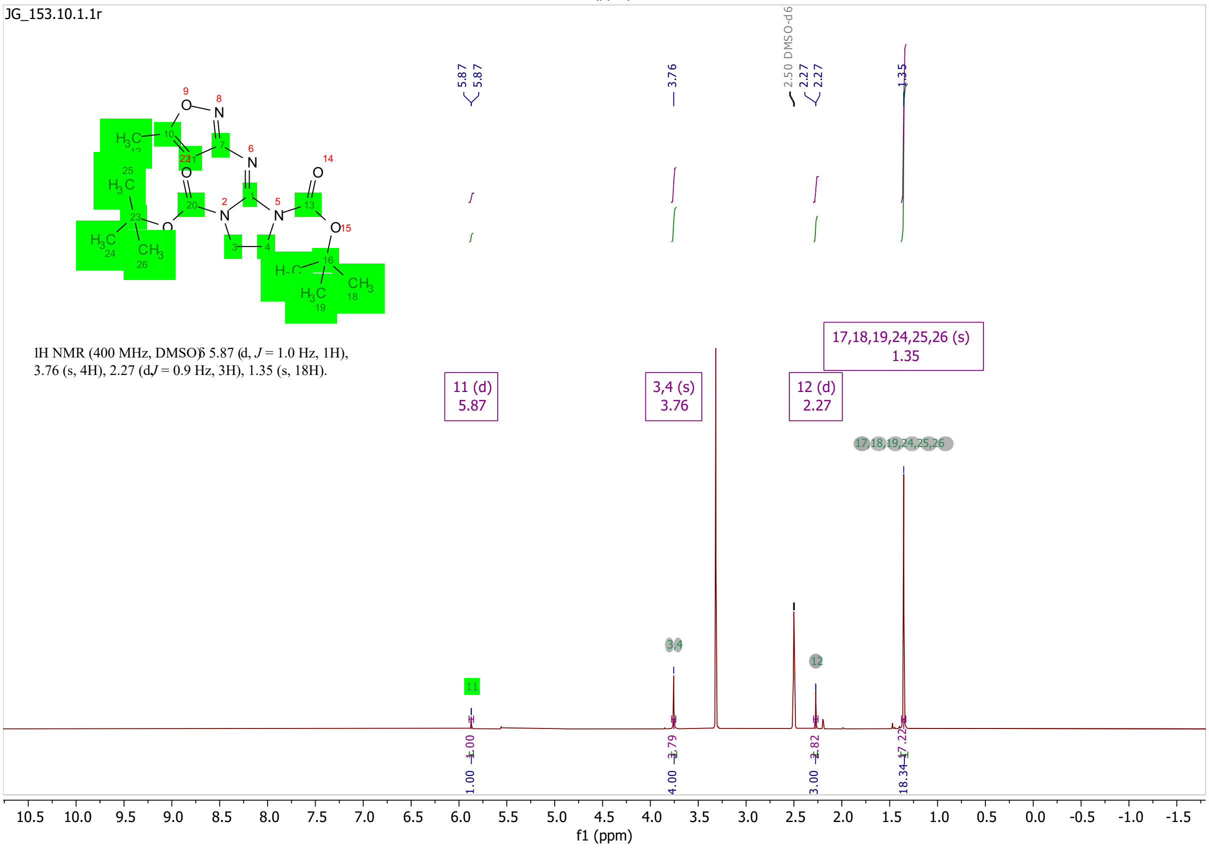
JG_153.10.1.1r
1H NMR (400 MHz, DMSO) δ 5.87 (d, J = 1.0 Hz, 1H), 3.76 (s, 4H), 2.27 (d, J = 0.9 Hz, 3H), 1.35 (s, 18H).
11 (d) 5.87
3,4 (s) 3.76
12 (d) 2.27
17,18,19,24,25,26 (s) 1.35
2.50 DMSO-d6
f1 (ppm)

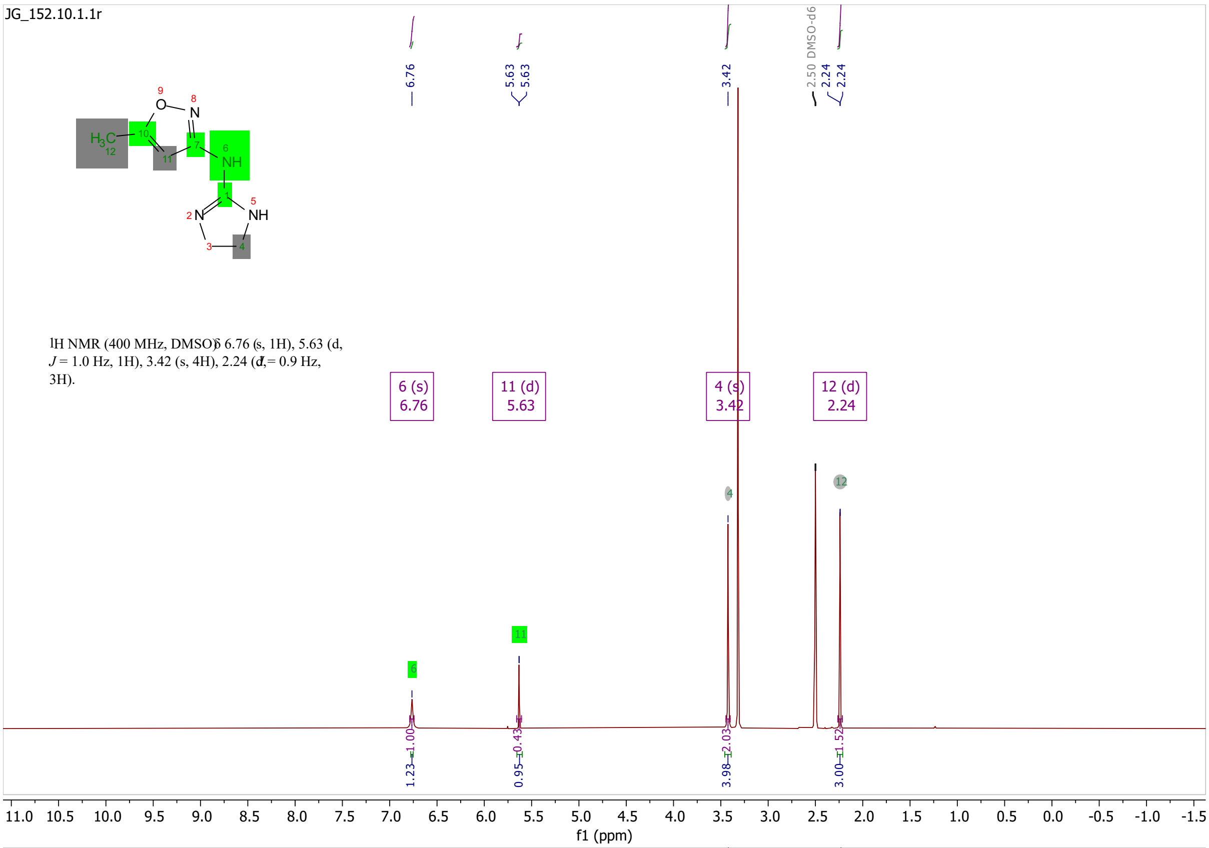
JG_152.10.1.1r
1H NMR (400 MHz, DMSO) δ 6.76 (s, 1H), 5.63 (d, J = 1.0 Hz, 1H), 3.42 (s, 4H), 2.24 (d, J = 0.9 Hz, 3H).
6 (s) 6.76
11 (d) 5.63
4 (s) 3.42
12 (d) 2.24
6.76
5.63
5.63
3.42
2.50 DMSO-d6
2.24
2.24
1.23
1.00
0.95
0.43
3.98
2.03
3.00
1.52
f1 (ppm)

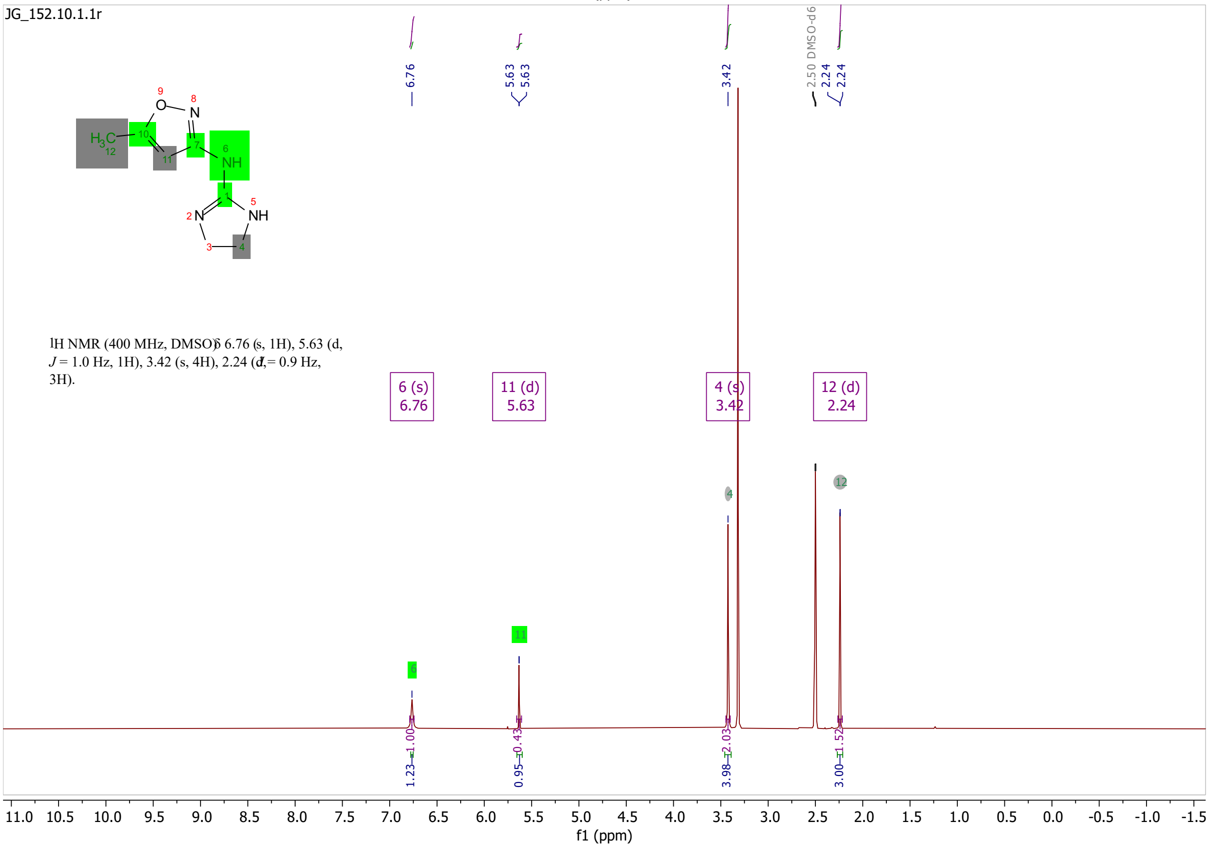
JG_152.10.1.1r
1H NMR (400 MHz, DMSO) δ 6.76 (s, 1H), 5.63 (d, J = 1.0 Hz, 1H), 3.42 (s, 4H), 2.24 (d, J = 0.9 Hz, 3H).
6 (s) 6.76
11 (d) 5.63
4 (s) 3.42
12 (d) 2.24
6.76
5.63
5.63
3.42
2.50 DMSO-d6
2.24
2.24
1.23
1.00
0.95
0.43
3.98
2.03
3.00
1.52
f1 (ppm)

S58

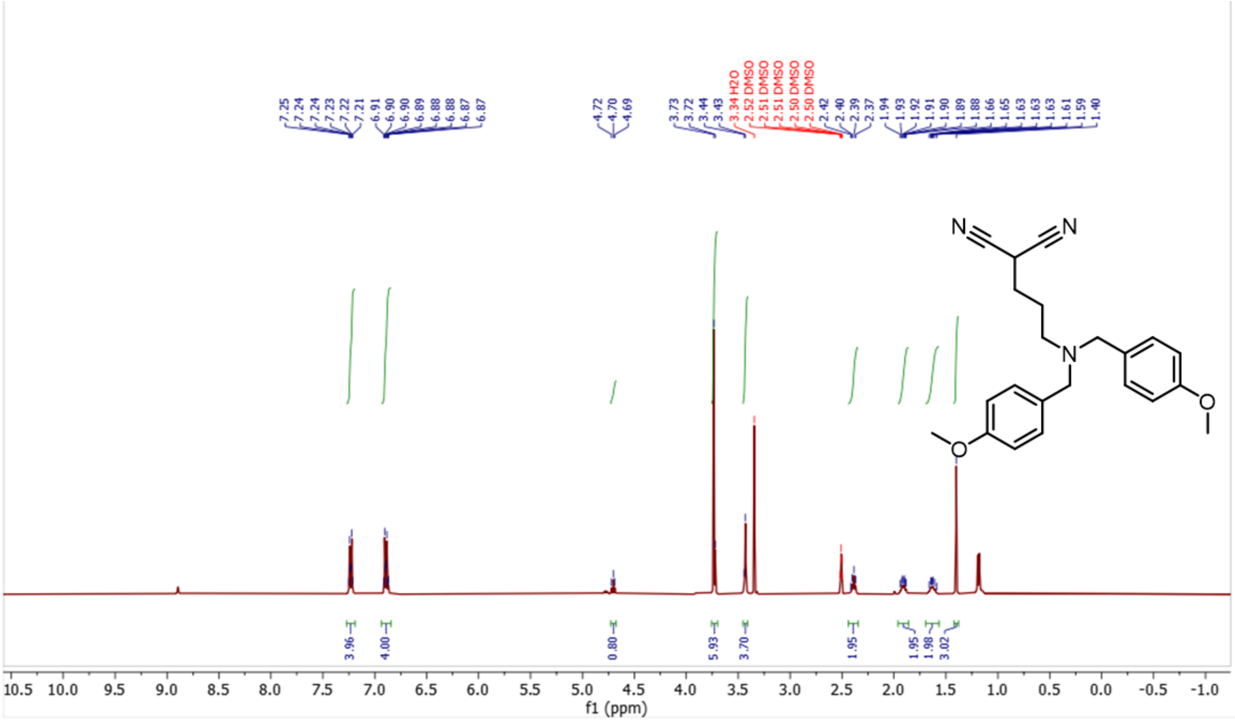

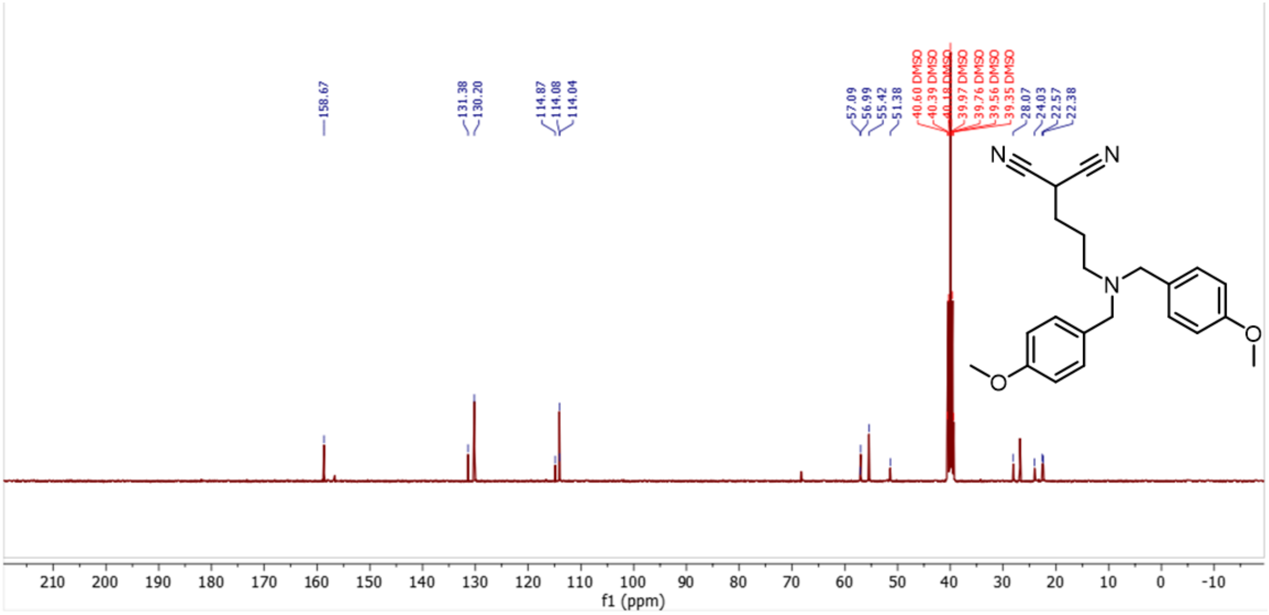

S60

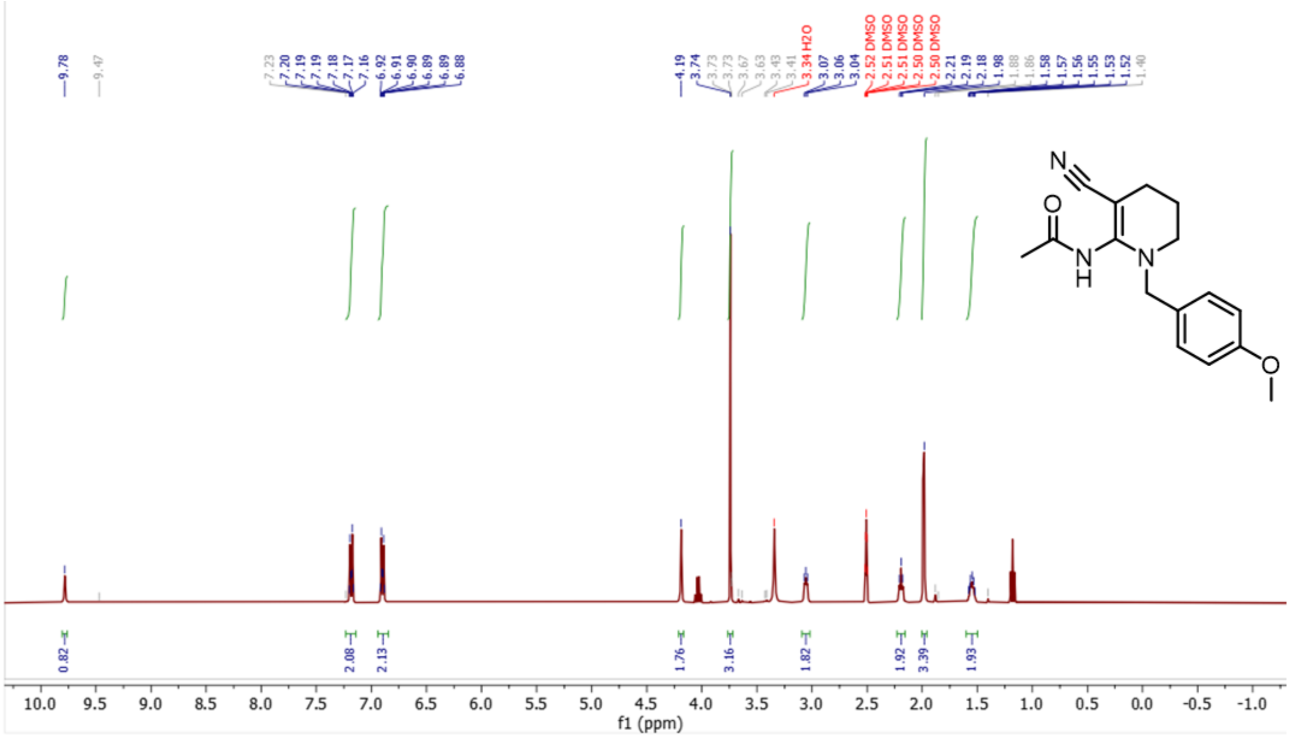

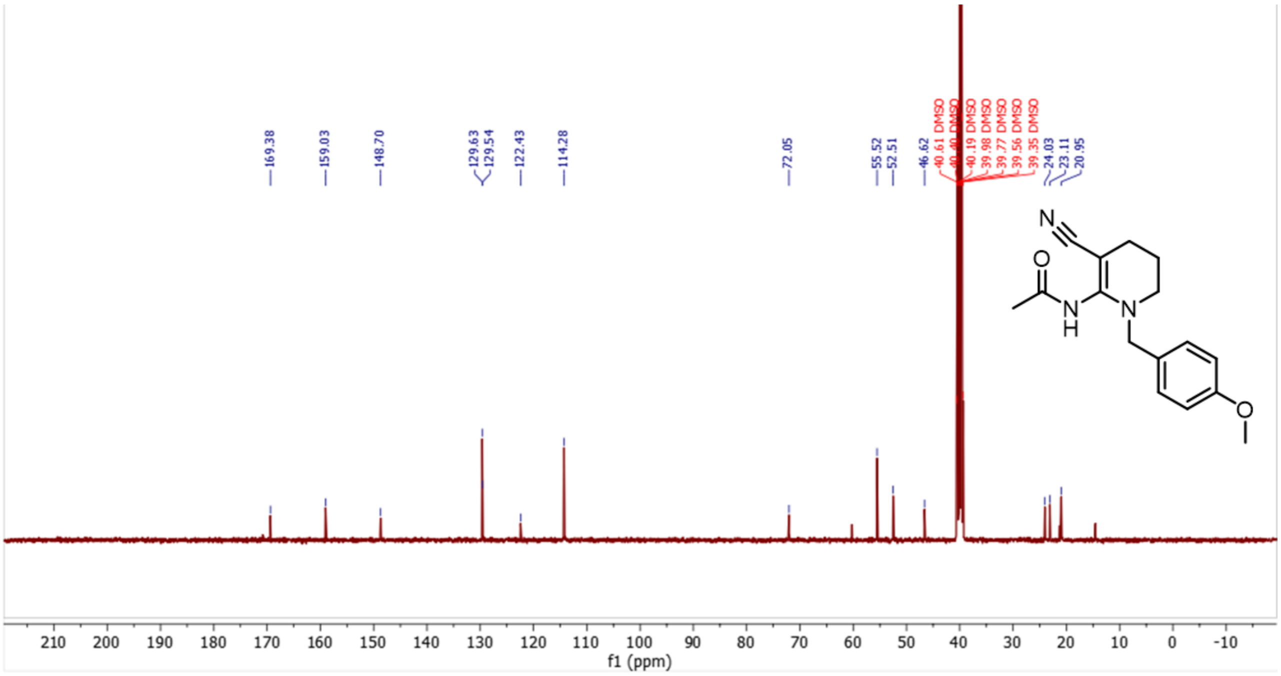

**S61**

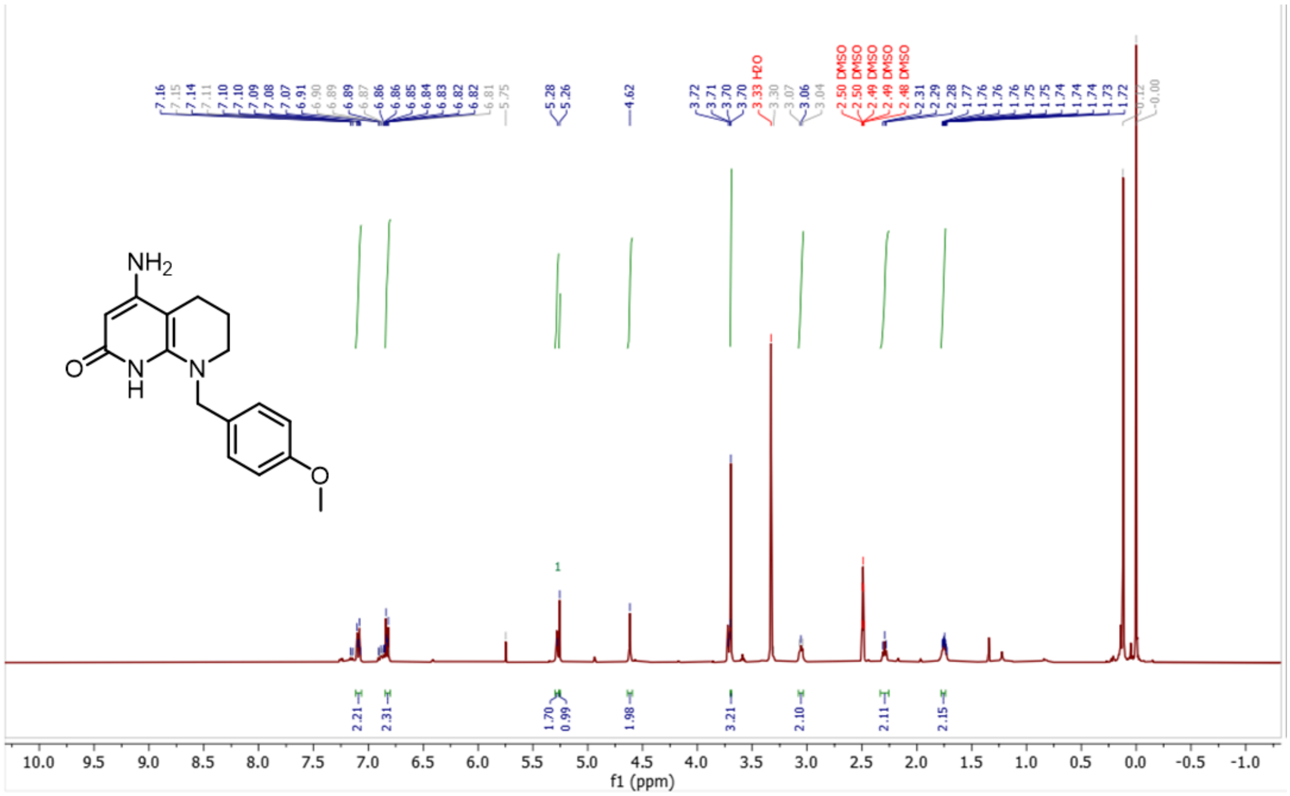

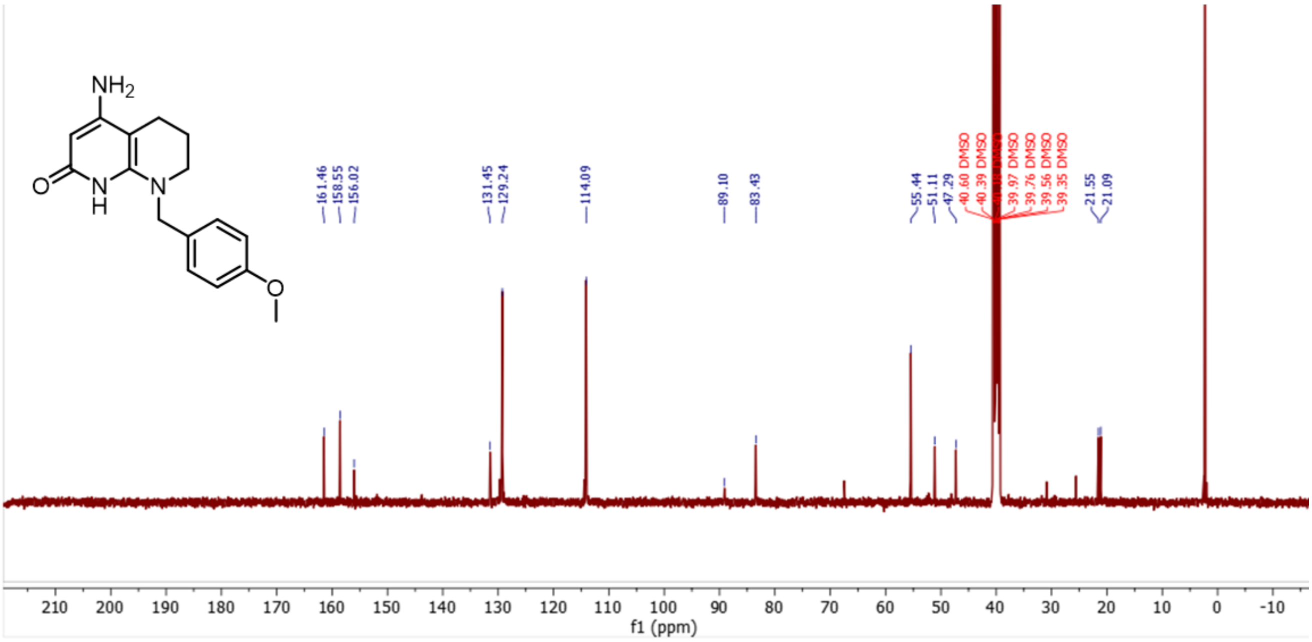

S27

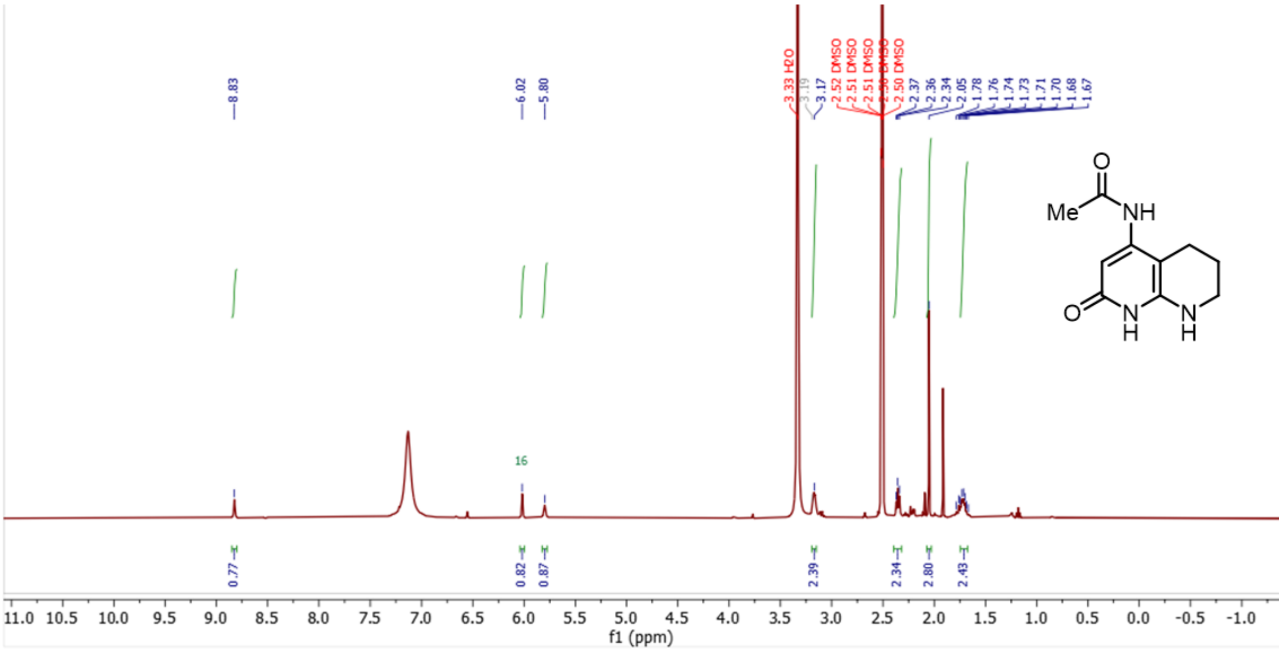

8.83
6.02
5.80
3.33 H2O
3.19
3.17
2.52 DMSO
2.51 DMSO
2.51 DMSO
2.50 DMSO
2.50 DMSO
2.37
2.36
2.34
2.05
1.78
1.76
1.74
1.73
1.71
1.70
1.68
1.67
Me
NH
O
N
H
16
0.77
0.82
0.87
2.39
2.34
2.80
2.43
f1 (ppm)

**S28**

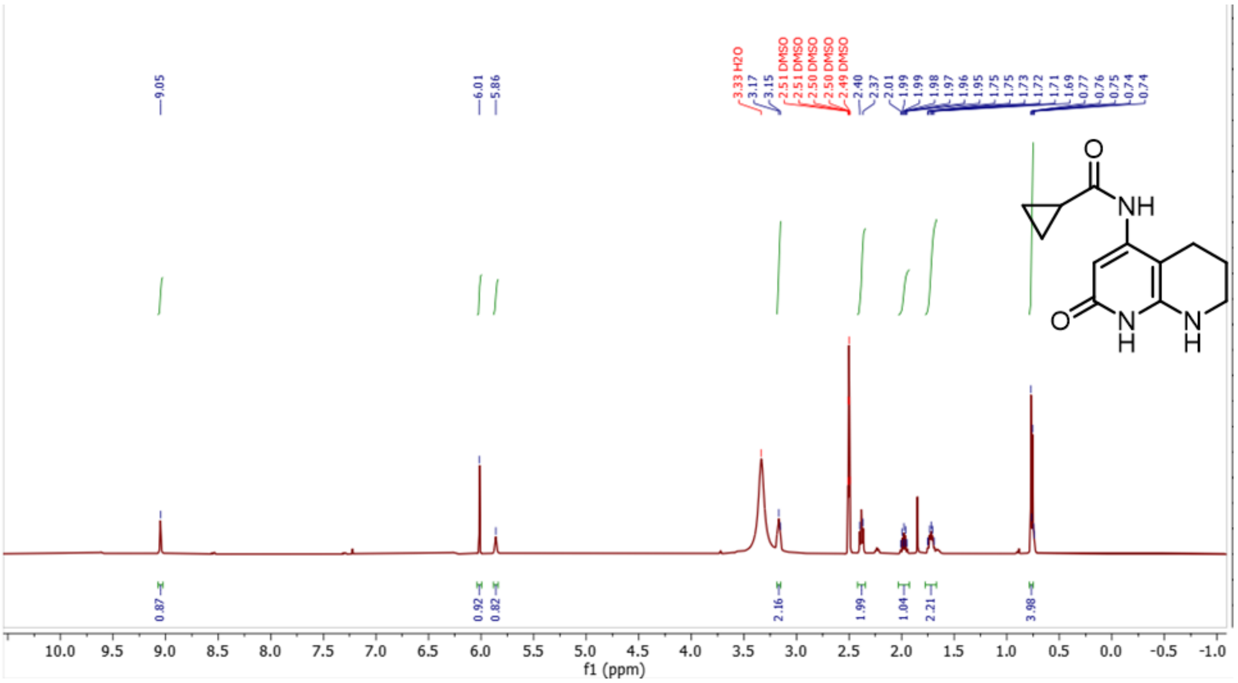

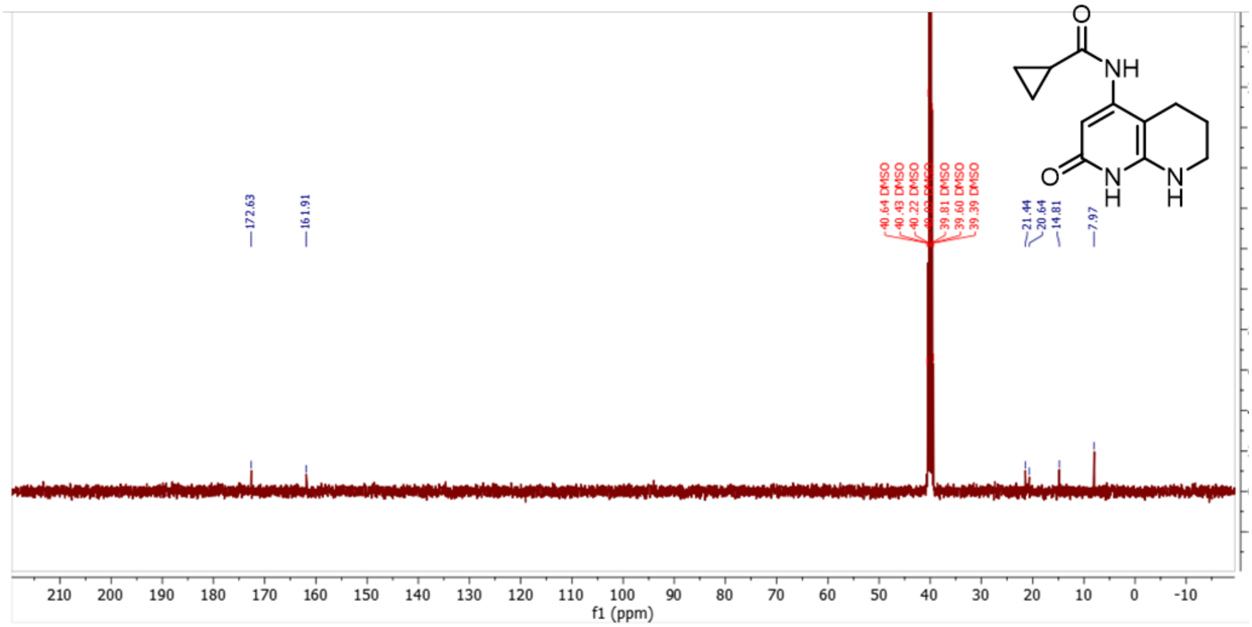

S29

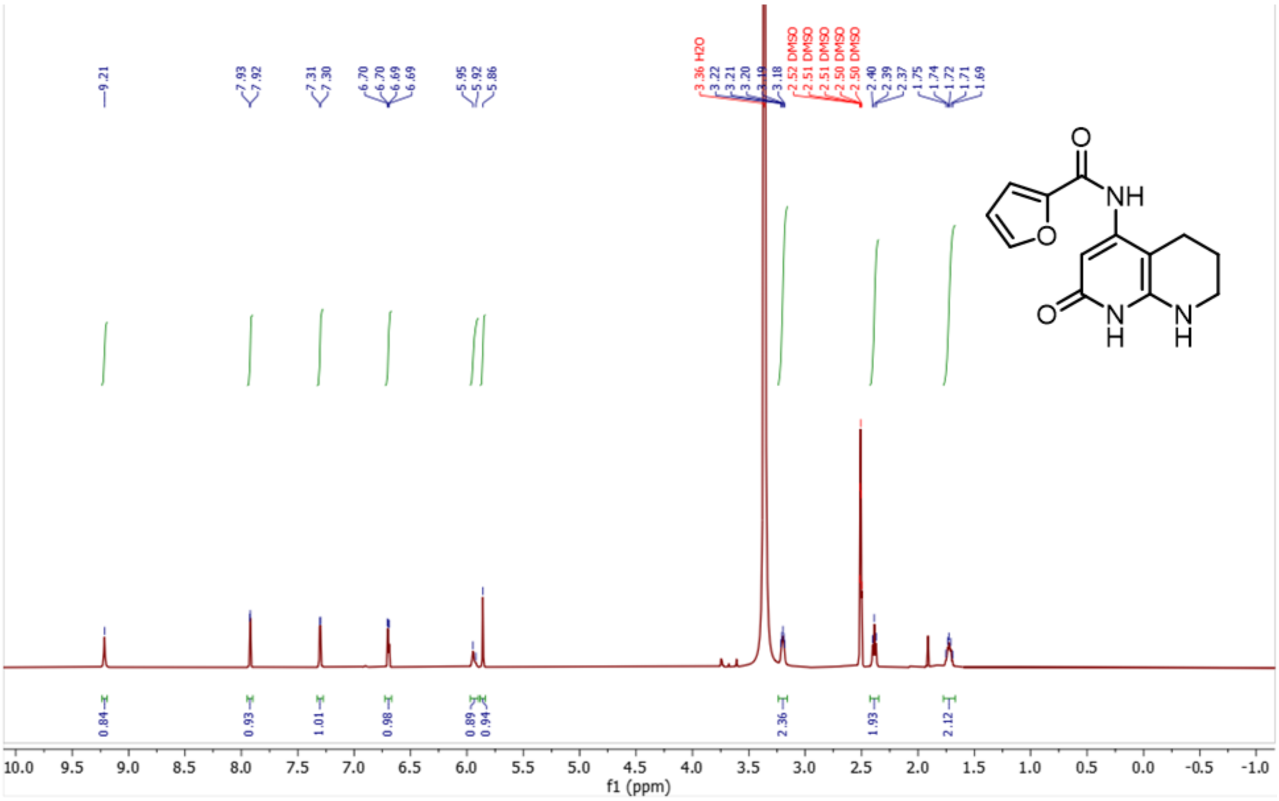
9.21
7.93
7.92
7.31
7.30
6.70
6.70
6.69
6.69
5.95
5.92
5.86
3.36 H2O
3.22
3.21
3.20
3.19
3.18
2.52 DMSO
2.51 DMSO
2.51 DMSO
2.50 DMSO
2.50 DMSO
2.40
2.39
2.37
1.75
1.74
1.72
1.71
1.69
O
NH
O
O
N
H
N
H
0.84
0.93
1.01
0.98
0.89
0.94
2.36
1.93
2.12
f1 (ppm)

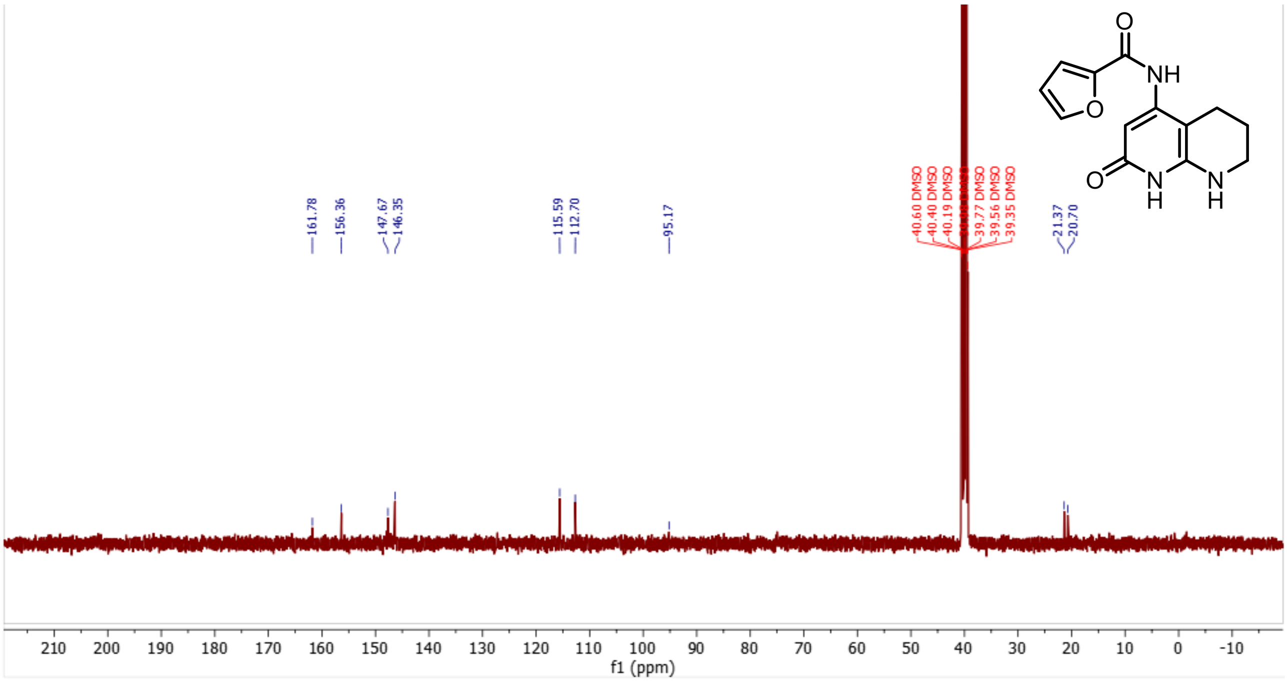
161.78
156.36
147.67
146.35
115.59
112.70
95.17
40.60 DMSO
40.40 DMSO
40.19 DMSO
39.77 DMSO
39.56 DMSO
39.35 DMSO
21.37
20.70
O
NH
O
O
N
H
N
H
f1 (ppm)

**S30**

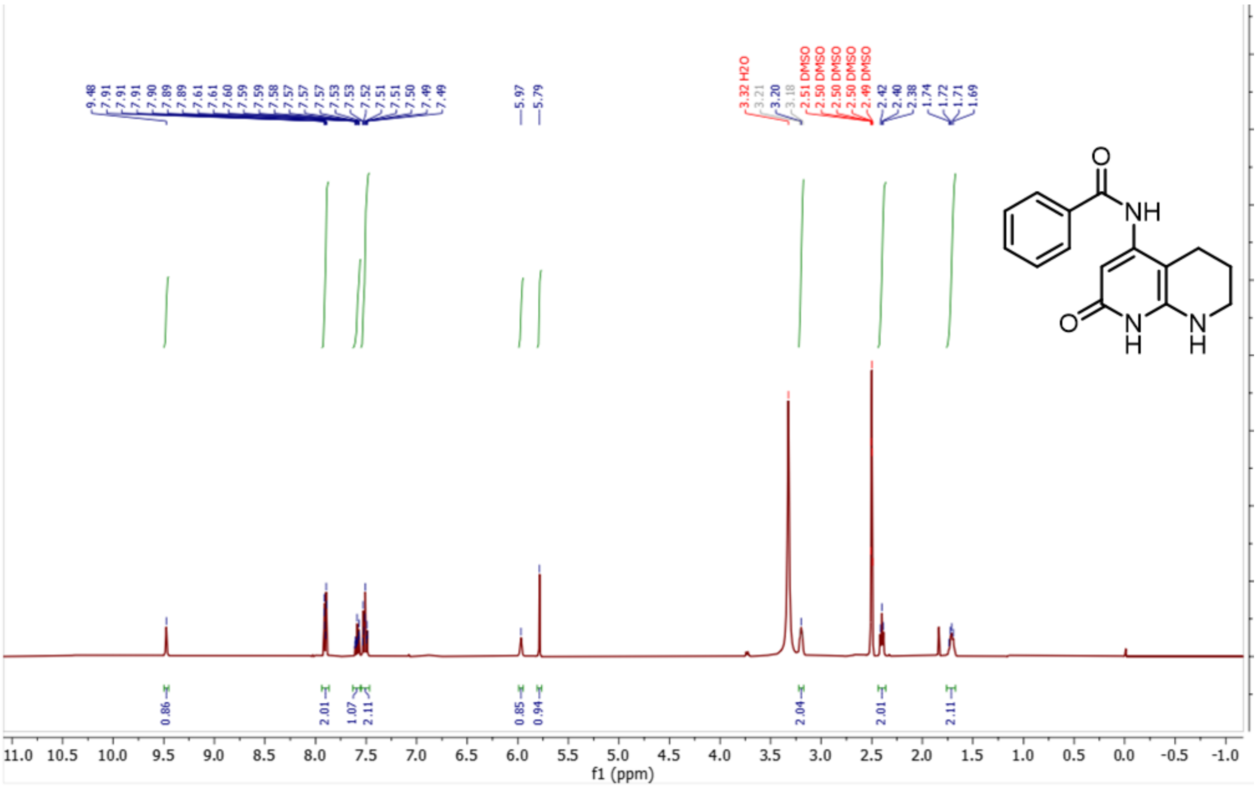

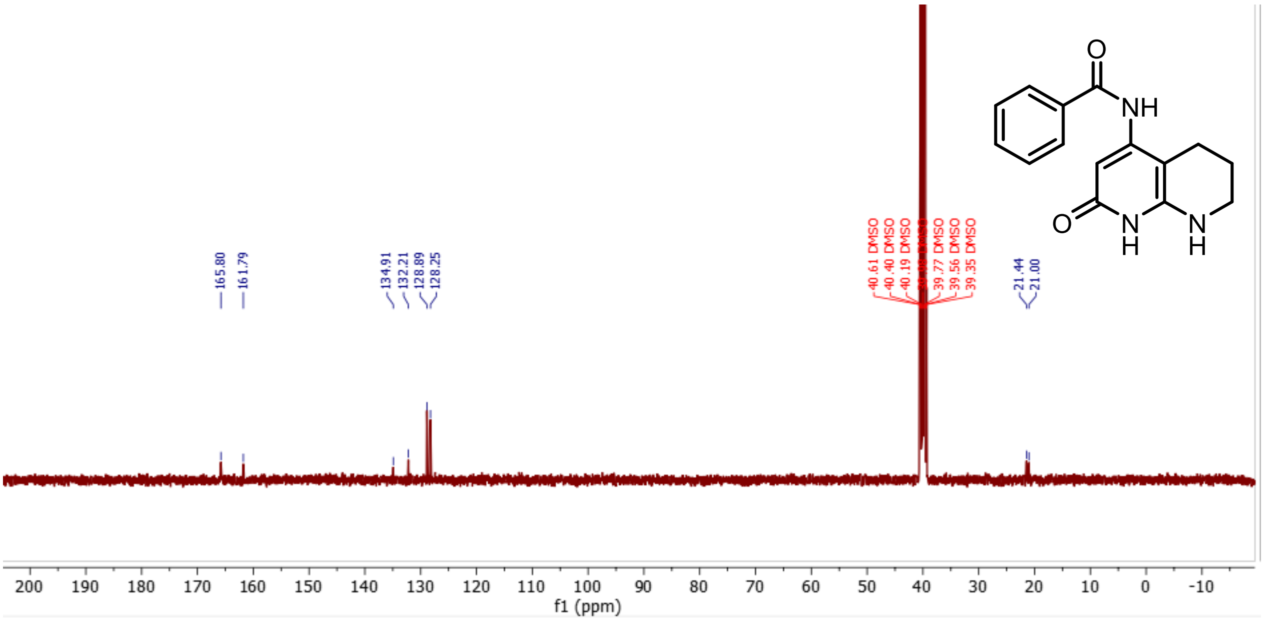

**S31**

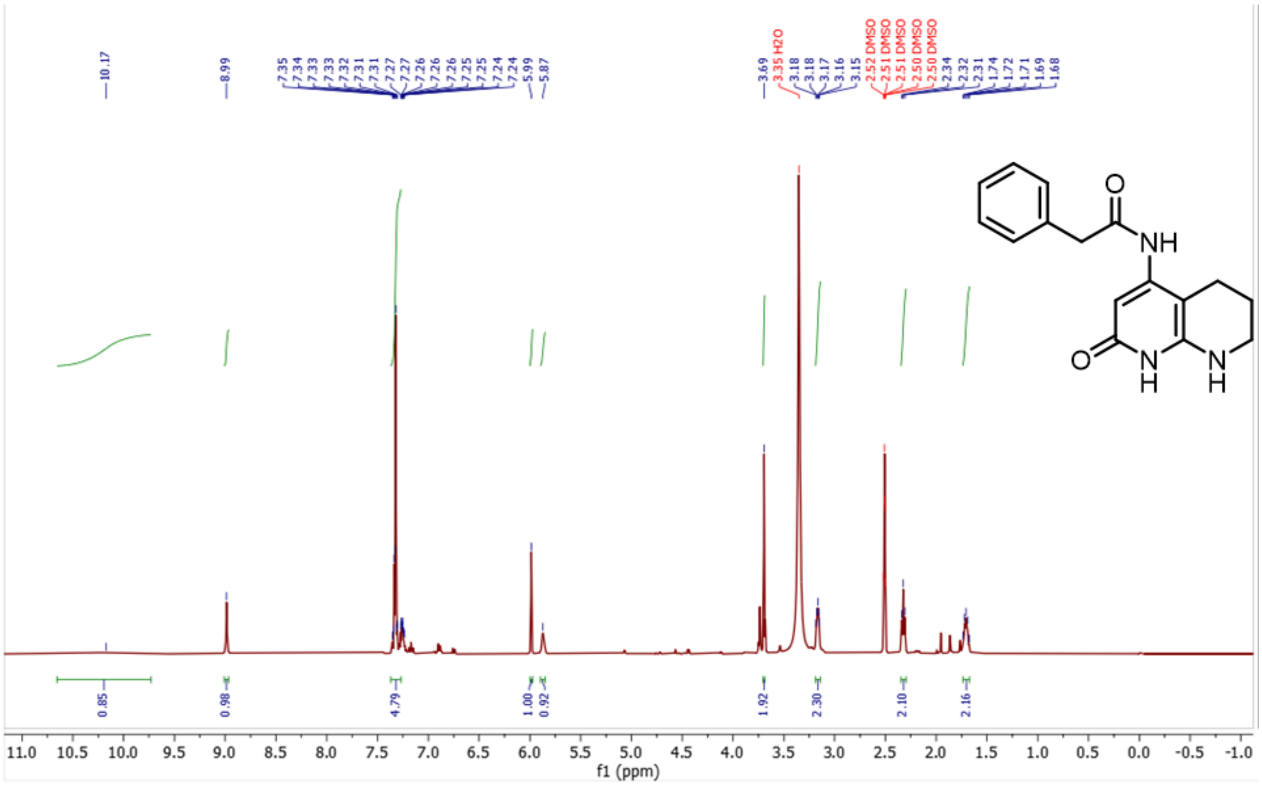
10.17
8.99
7.35
7.34
7.33
7.33
7.32
7.31
7.31
7.27
7.27
7.26
7.26
7.25
7.25
7.24
7.24
5.99
5.87
3.69
3.35 H2O
3.18
3.18
3.17
3.16
3.15
2.52 DMSO
2.51 DMSO
2.51 DMSO
2.50 DMSO
2.50 DMSO
2.34
2.32
2.31
1.74
1.72
1.71
1.69
1.68
0.85
0.98
4.79
1.00
0.92
1.92
2.30
2.10
2.16
f1 (ppm)
O
NH
O
N
H
N
H

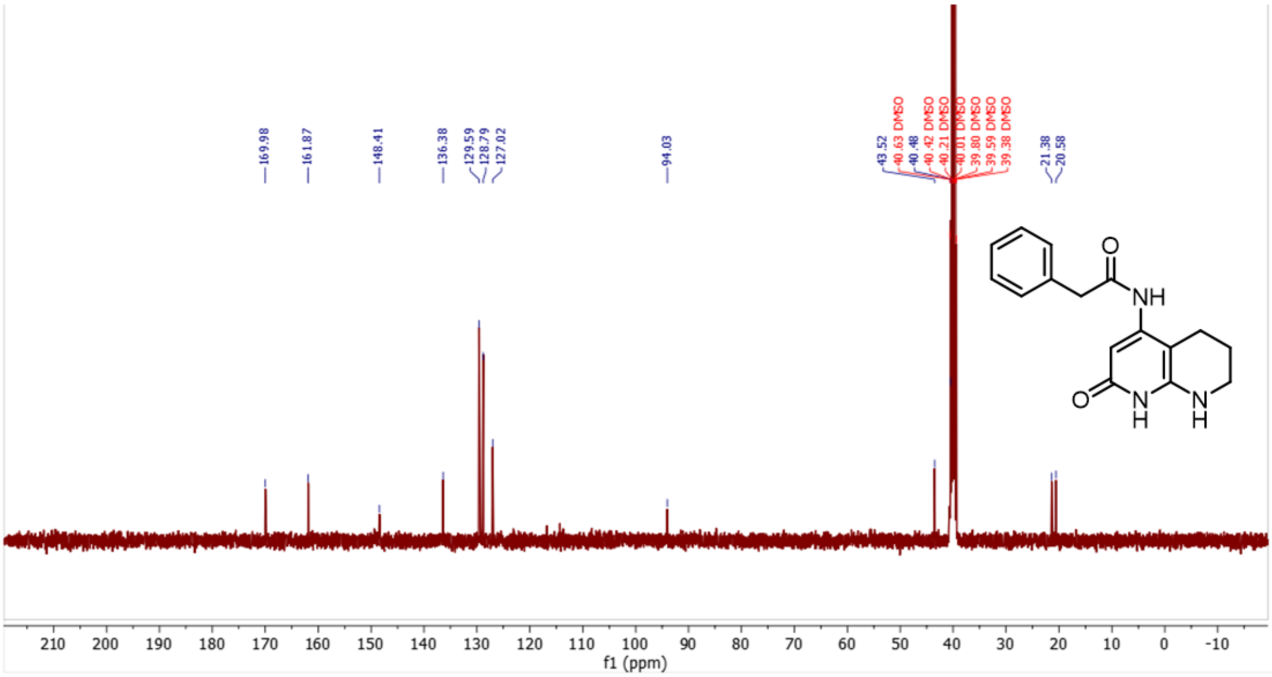
169.98
161.87
148.41
136.38
129.59
128.79
127.02
94.03
43.52
40.63 DMSO
40.48
40.42 DMSO
40.21 DMSO
40.01 DMSO
39.80 DMSO
39.59 DMSO
39.38 DMSO
21.38
20.58
f1 (ppm)
O
NH
O
N
H
N
H

**S63**

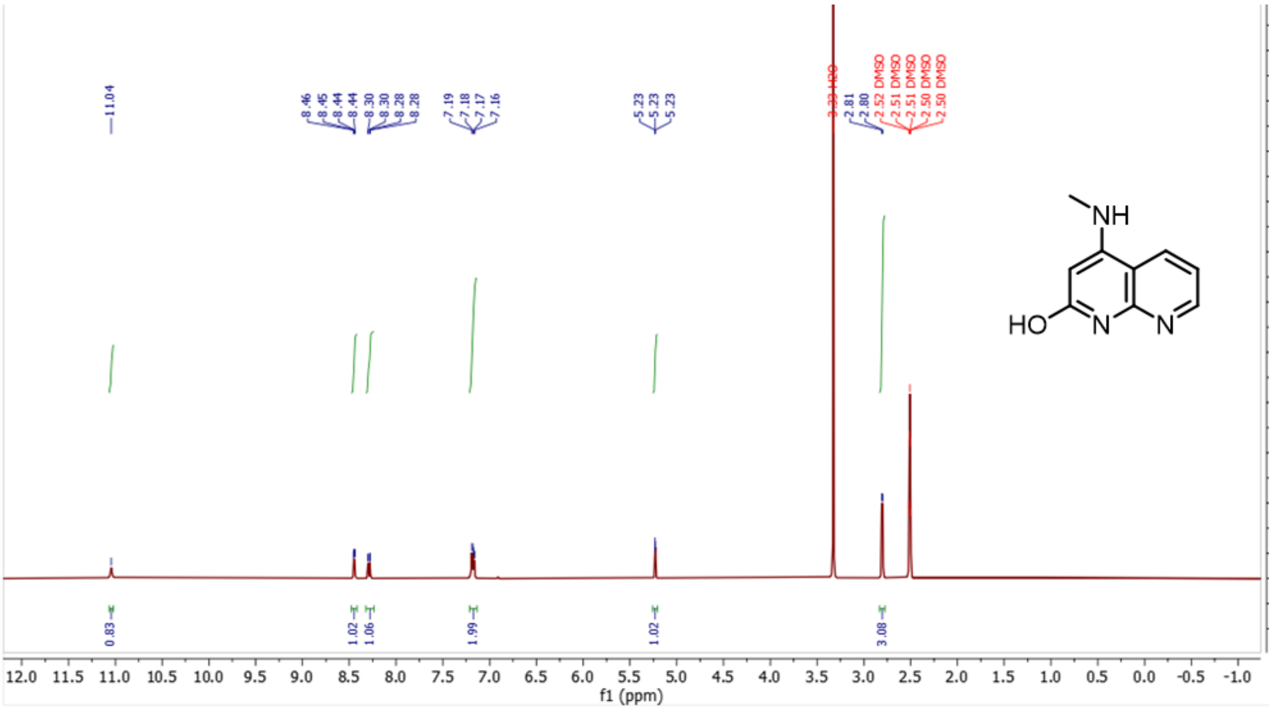

11.04
8.46
8.45
8.44
8.44
8.30
8.30
8.28
8.28
7.19
7.18
7.17
7.16
5.23
5.23
5.23
3.33 H2O
2.81
2.80
2.52 DMSO
2.51 DMSO
2.51 DMSO
2.50 DMSO
2.50 DMSO
NH
HO
N
N
0.83
1.02
1.06
1.99
1.02
3.08
12.0 11.5 11.0 10.5 10.0 9.5 9.0 8.5 8.0 7.5 7.0 6.5 6.0 5.5 5.0 4.5 4.0 3.5 3.0 2.5 2.0 1.5 1.0 0.5 0.0 -0.5 -1.0
f1 (ppm)

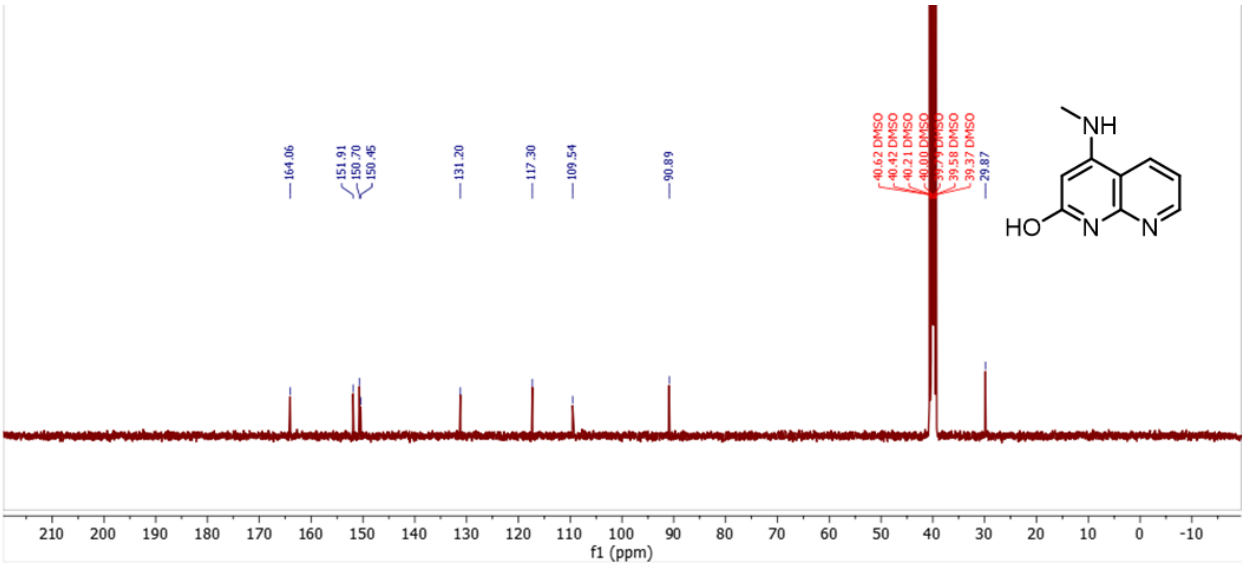

164.06
151.91
150.70
150.45
131.20
117.30
109.54
90.89
40.62 DMSO
40.42 DMSO
40.21 DMSO
40.00 DMSO
39.79 DMSO
39.58 DMSO
39.37 DMSO
29.87
NH
HO
N
N
210 200 190 180 170 160 150 140 130 120 110 100 90 80 70 60 50 40 30 20 10 0 -10
f1 (ppm)

**S64**

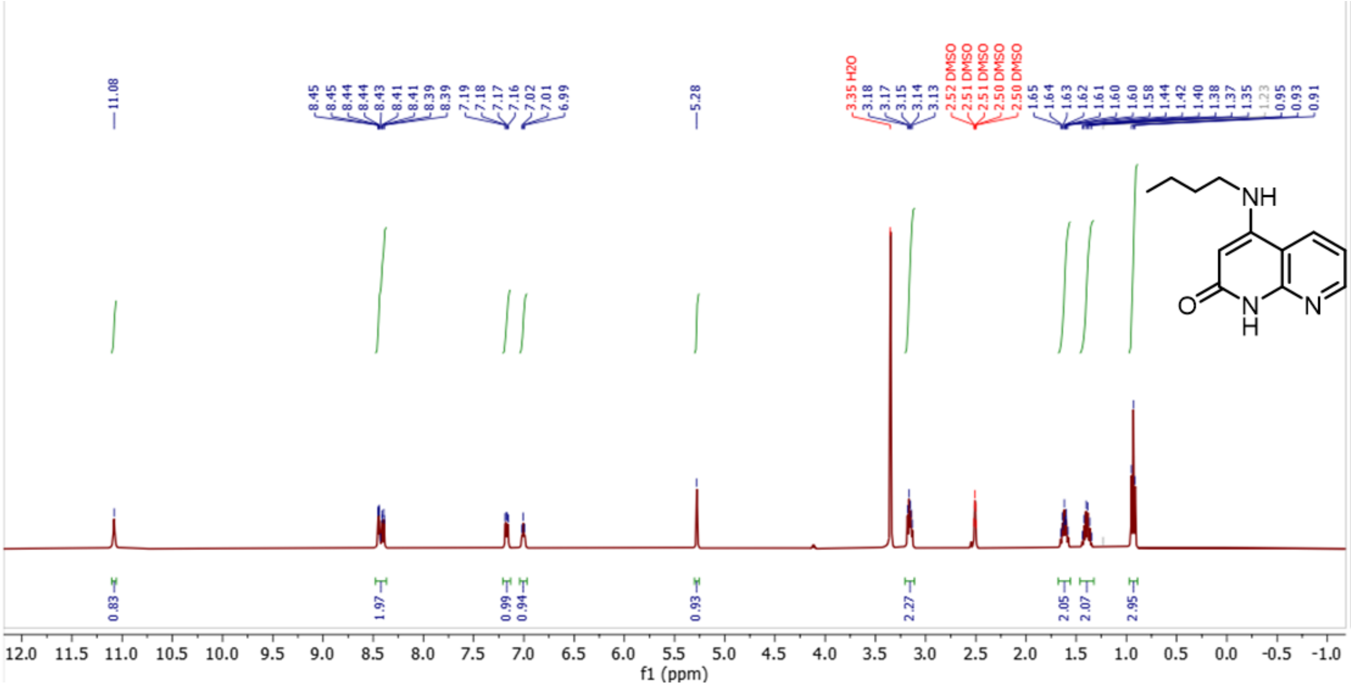
11.08
8.45
8.45
8.44
8.44
8.43
8.41
8.41
8.41
8.39
8.39
7.19
7.18
7.17
7.16
7.02
7.01
6.99
5.28
3.35 H2O
3.18
3.17
3.15
3.14
3.13
2.52 DMSO
2.51 DMSO
2.51 DMSO
2.50 DMSO
2.50 DMSO
1.65
1.64
1.63
1.62
1.61
1.60
1.60
1.58
1.44
1.42
1.40
1.38
1.37
1.35
1.23
0.95
0.93
0.91
NH
O
N
H
N
0.83
1.97
0.99
0.94
0.93
2.27
2.05
2.07
2.95
12.0 11.5 11.0 10.5 10.0 9.5 9.0 8.5 8.0 7.5 7.0 6.5 6.0 5.5 5.0 4.5 4.0 3.5 3.0 2.5 2.0 1.5 1.0 0.5 0.0 -0.5 -1.0
f1 (ppm)

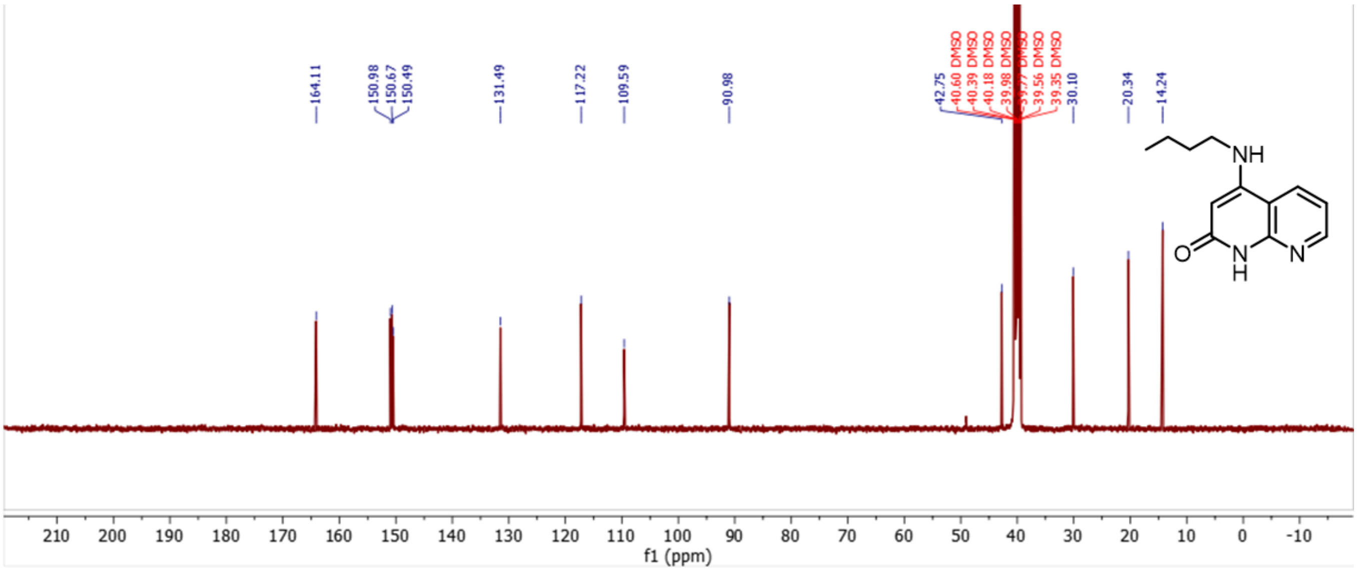
164.11
150.98
150.67
150.49
131.49
117.22
109.59
90.98
42.75
40.60 DMSO
40.39 DMSO
40.18 DMSO
39.98 DMSO
39.77 DMSO
39.56 DMSO
39.35 DMSO
30.10
20.34
14.24
NH
O
N
H
N
210 200 190 180 170 160 150 140 130 120 110 100 90 80 70 60 50 40 30 20 10 0 -10
f1 (ppm)

S65

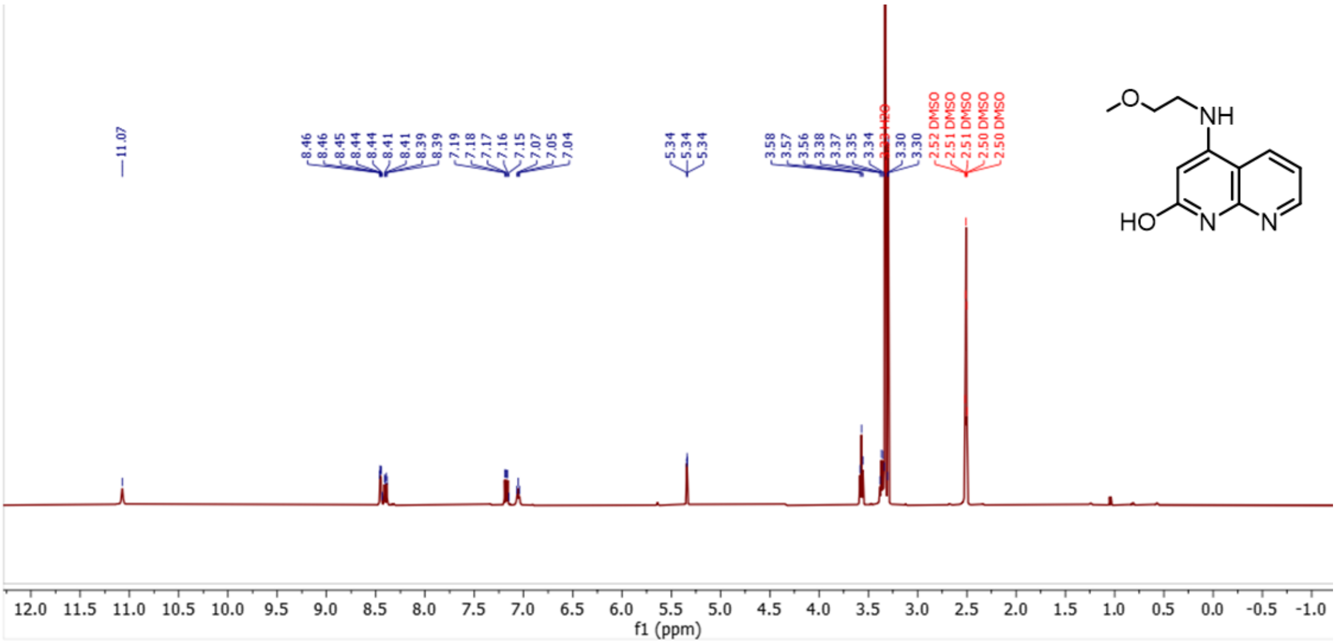

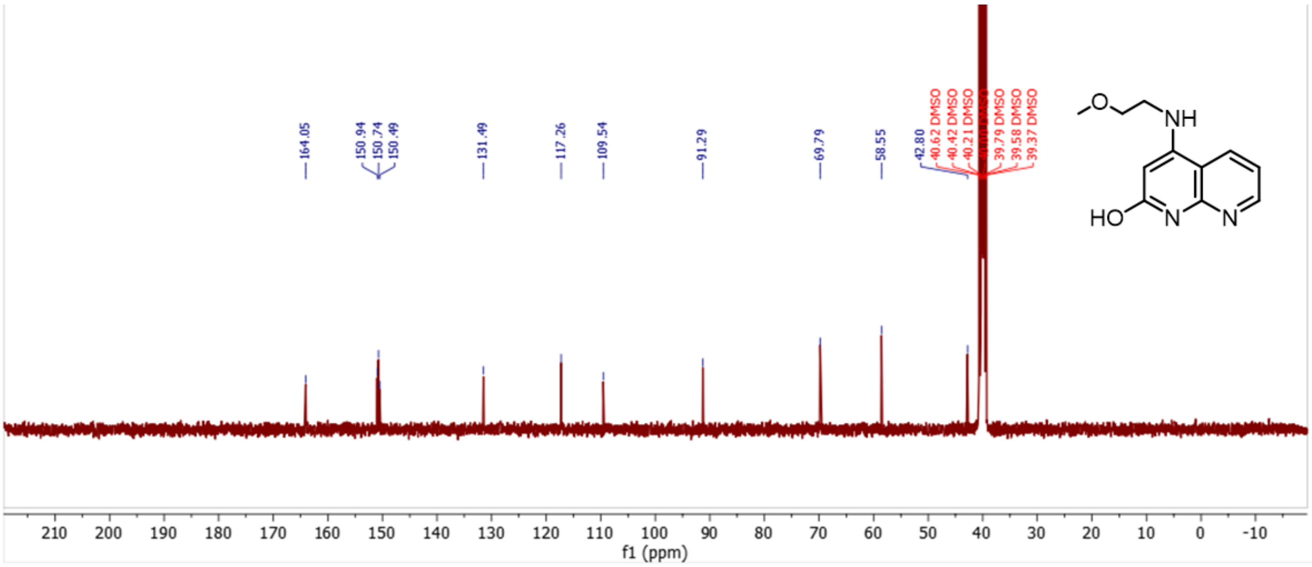

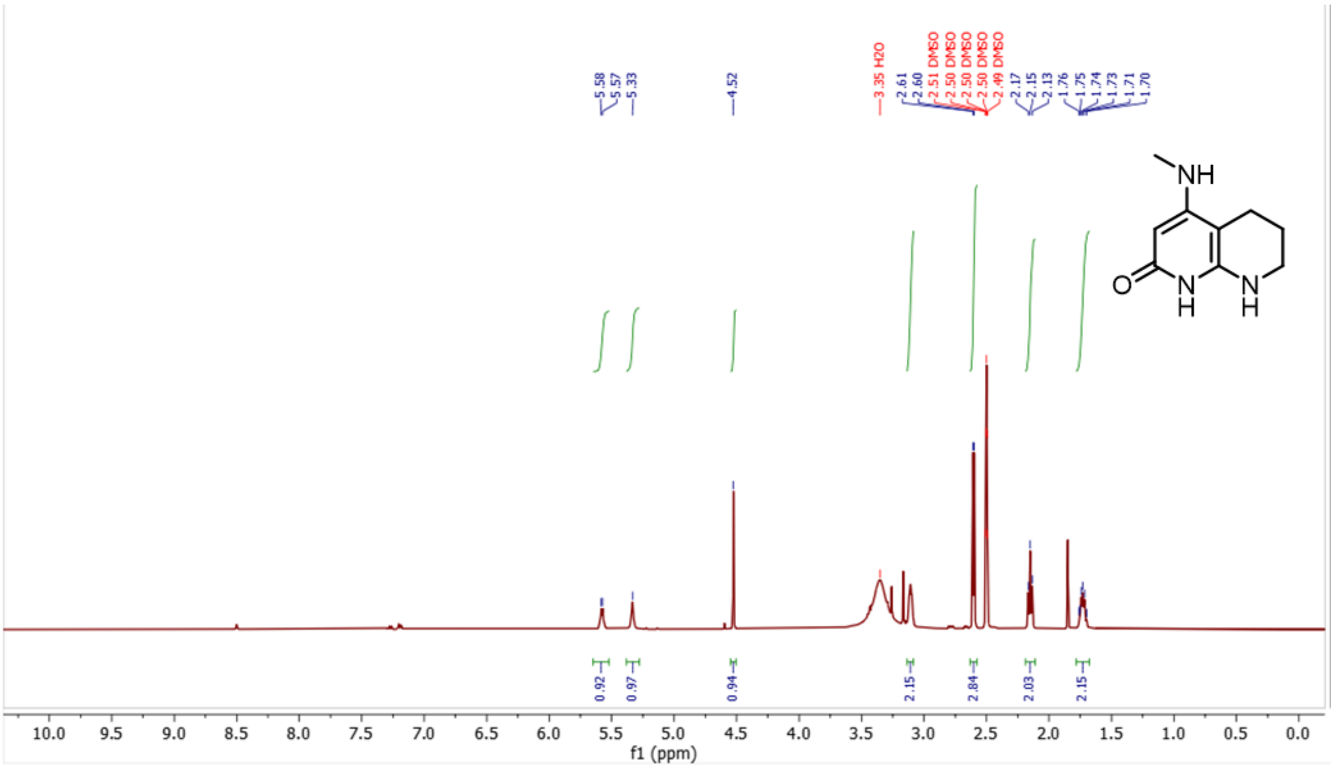
5.58
5.57
5.33
4.52
3.35 H2O
2.61
2.60
2.51 DMSO
2.50 DMSO
2.50 DMSO
2.50 DMSO
2.49 DMSO
2.17
2.15
2.13
1.76
1.75
1.74
1.73
1.71
1.70
NH
O
N
H
N
H
0.92
0.97
0.94
2.15
2.84
2.03
2.15
10.0 9.5 9.0 8.5 8.0 7.5 7.0 6.5 6.0 5.5 5.0 4.5 4.0 3.5 3.0 2.5 2.0 1.5 1.0 0.5 0.0
f1 (ppm)

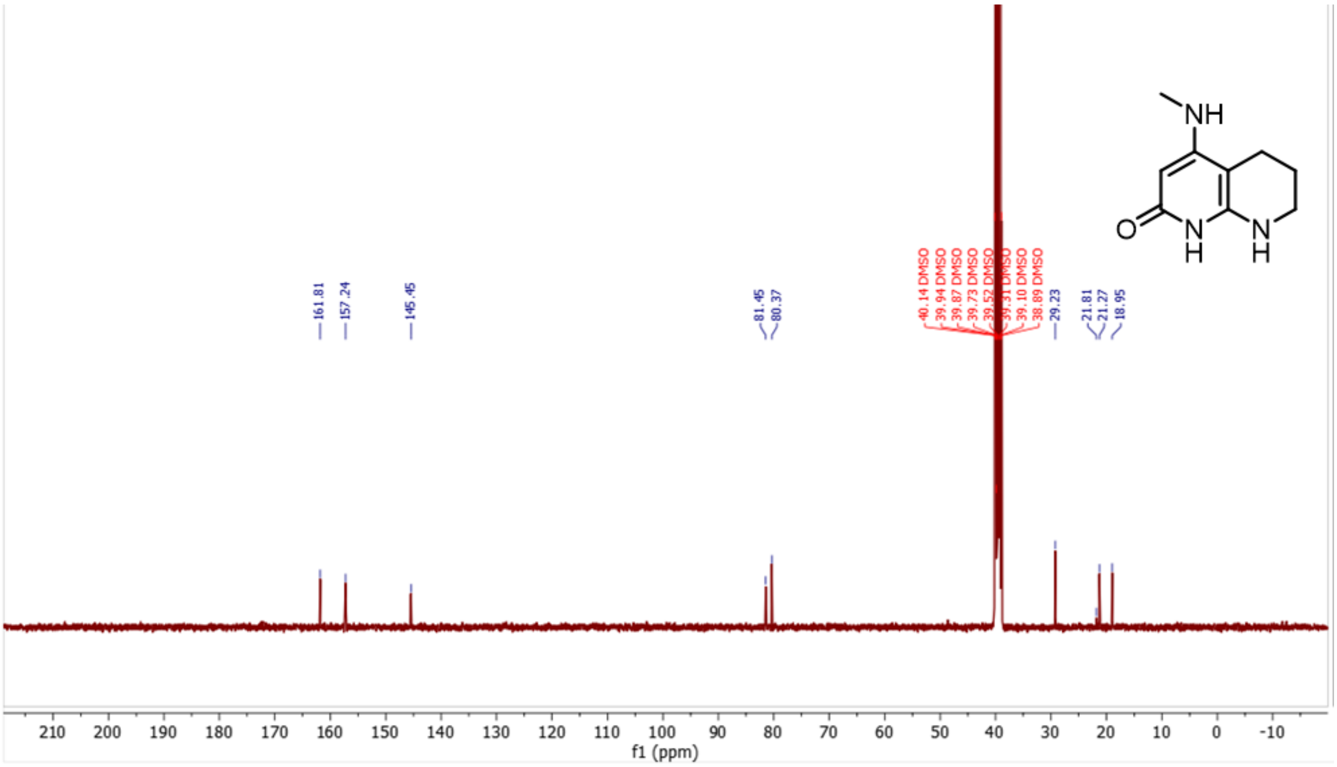
NH
O
N
H
N
H
161.81
157.24
145.45
81.45
80.37
40.14 DMSO
39.94 DMSO
39.87 DMSO
39.73 DMSO
39.52 DMSO
39.31 DMSO
39.10 DMSO
38.89 DMSO
29.23
21.81
21.27
18.95
210 200 190 180 170 160 150 140 130 120 110 100 90 80 70 60 50 40 30 20 10 0 -10
f1 (ppm)

# S34

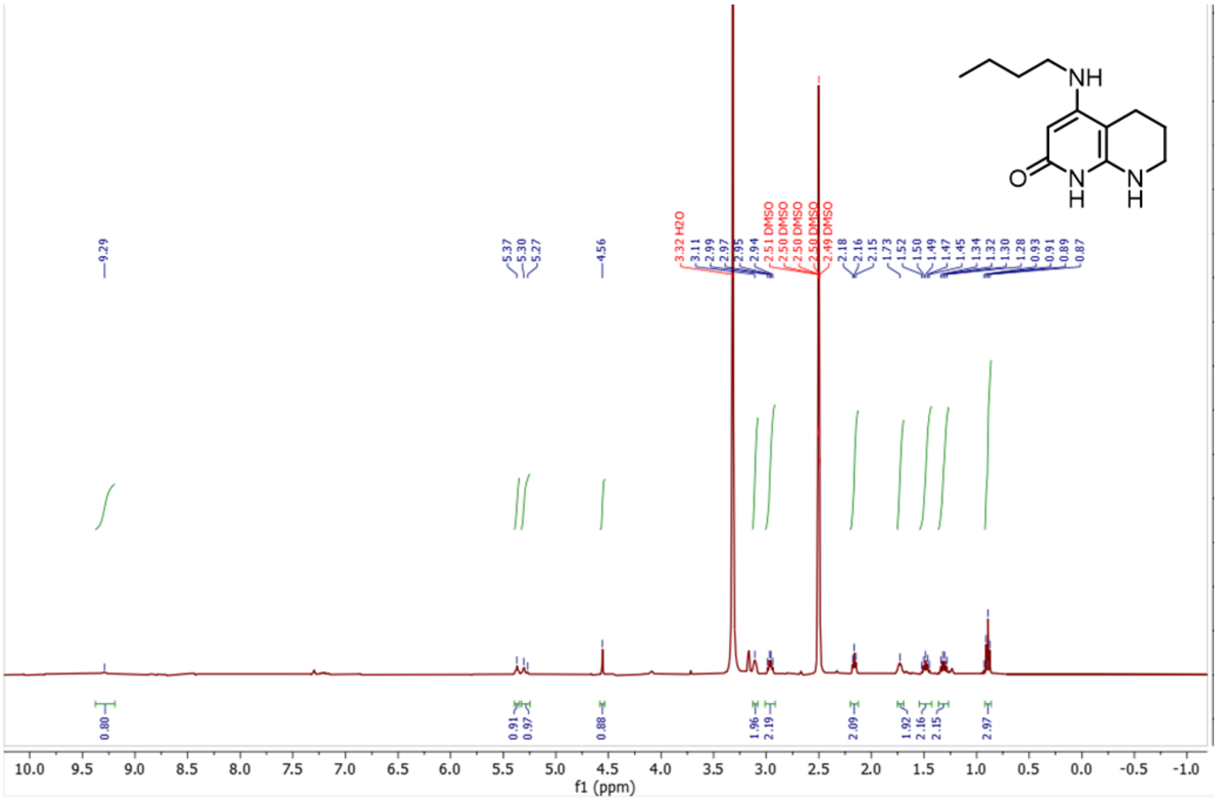

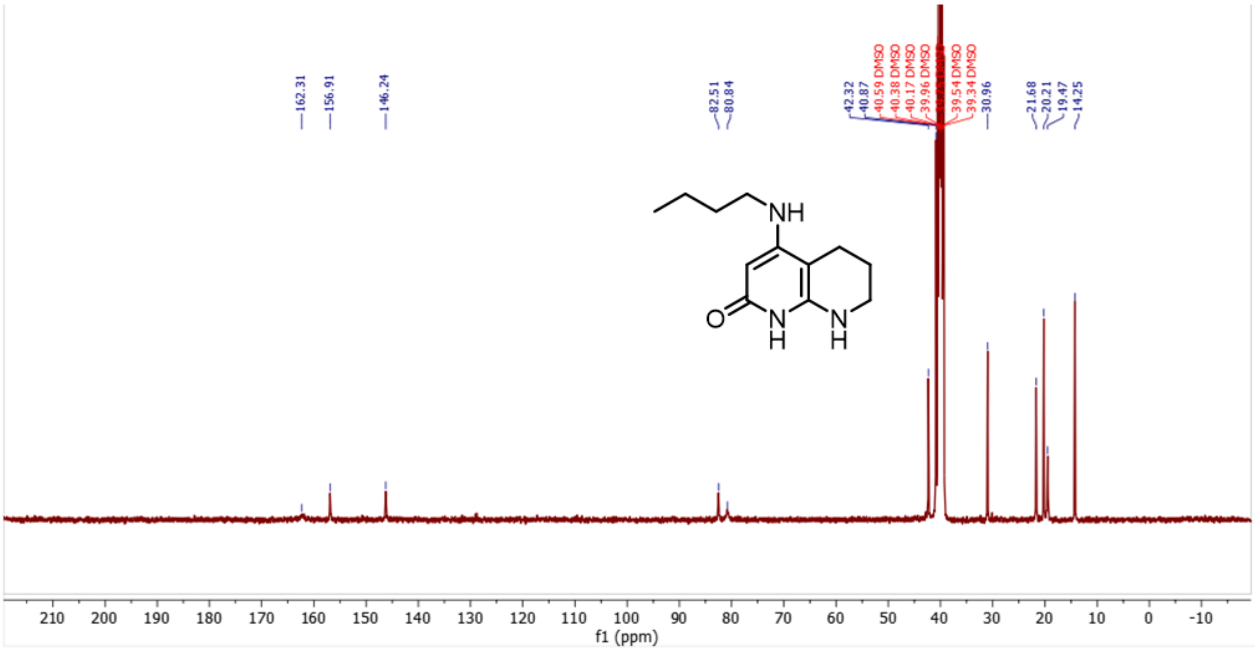

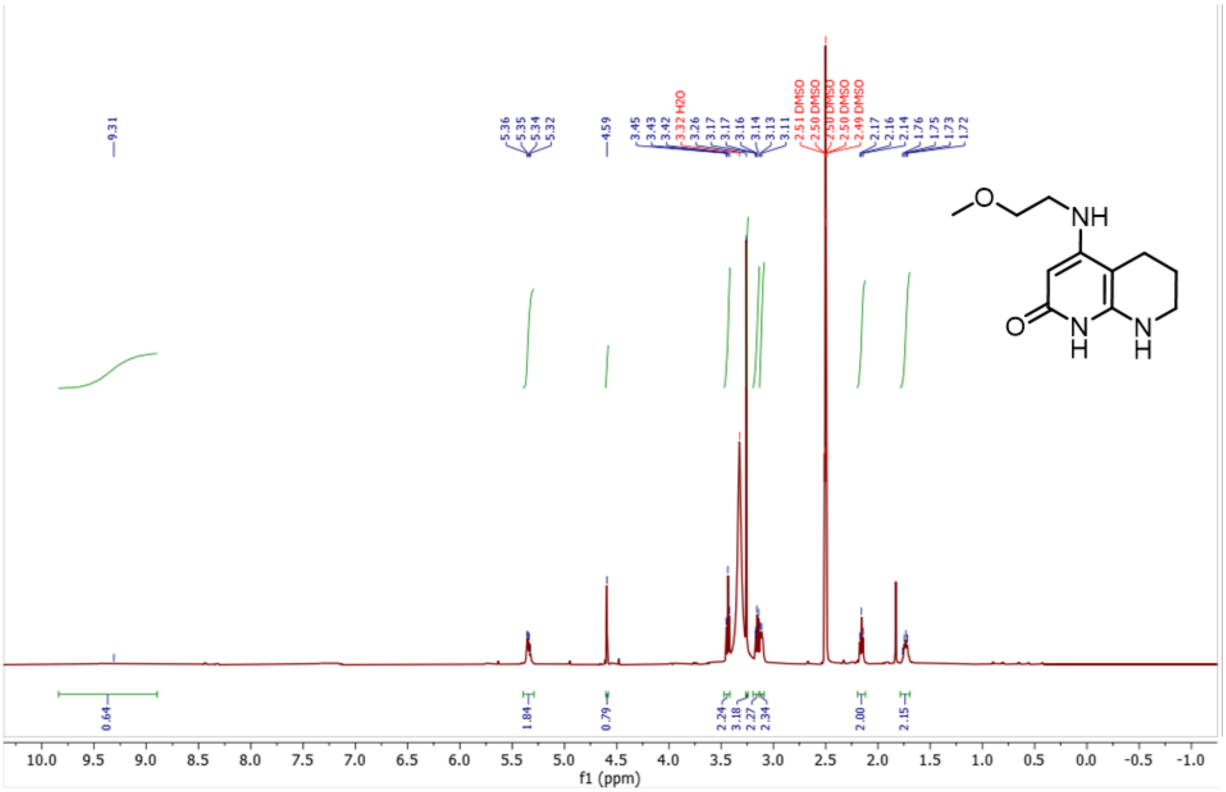

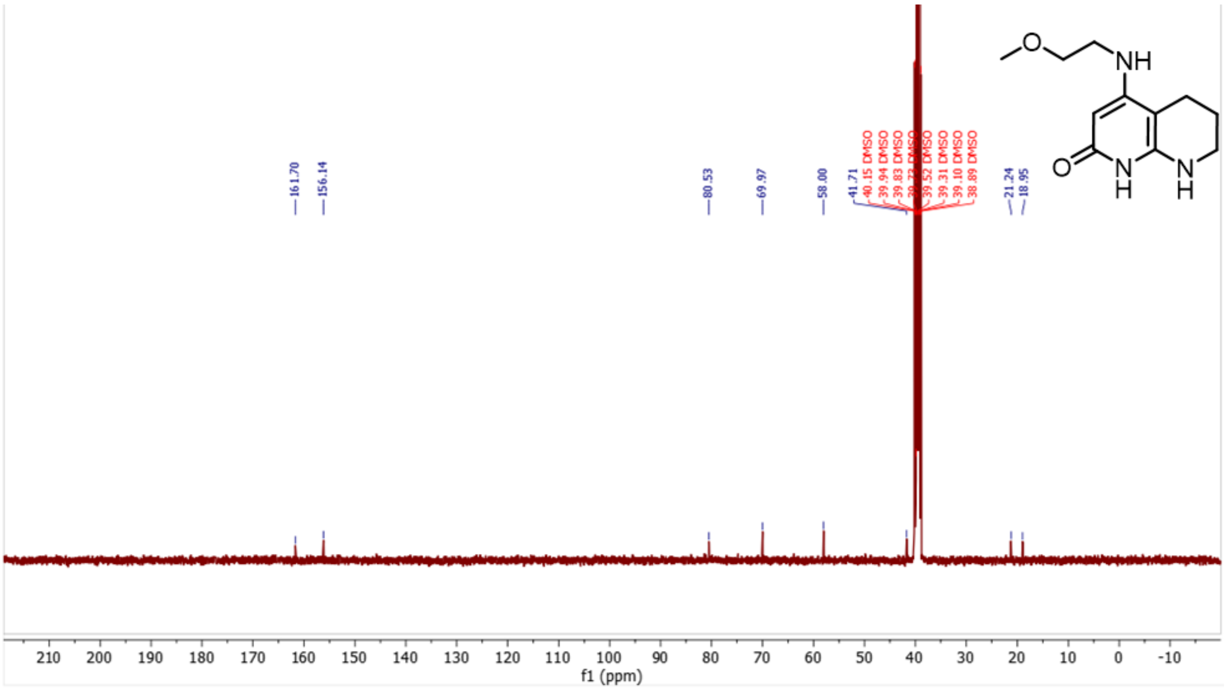

# S67

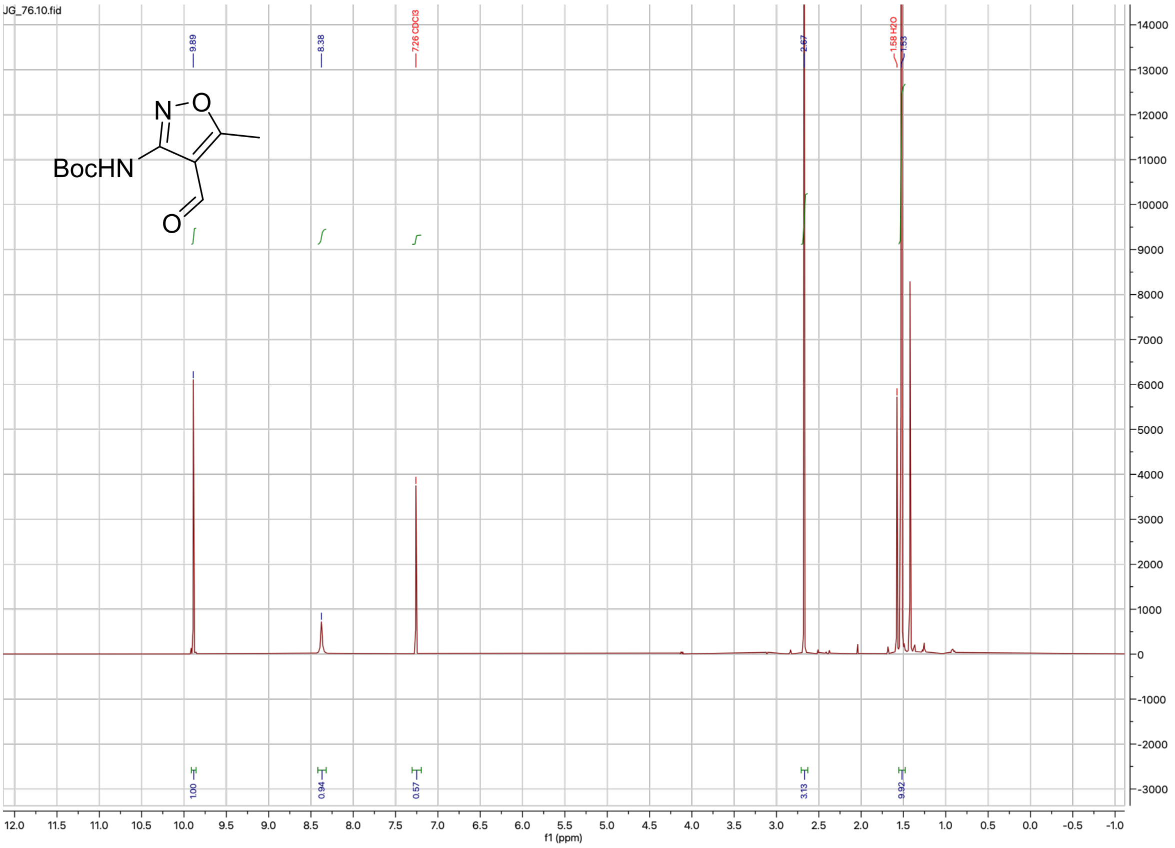
JG_76.10.fid
BocHN
N
O
9.89
8.38
7.26 CDCl3
2.67
1.58 H2O
1.53
1.00
0.94
0.57
3.13
9.92
f1 (ppm)

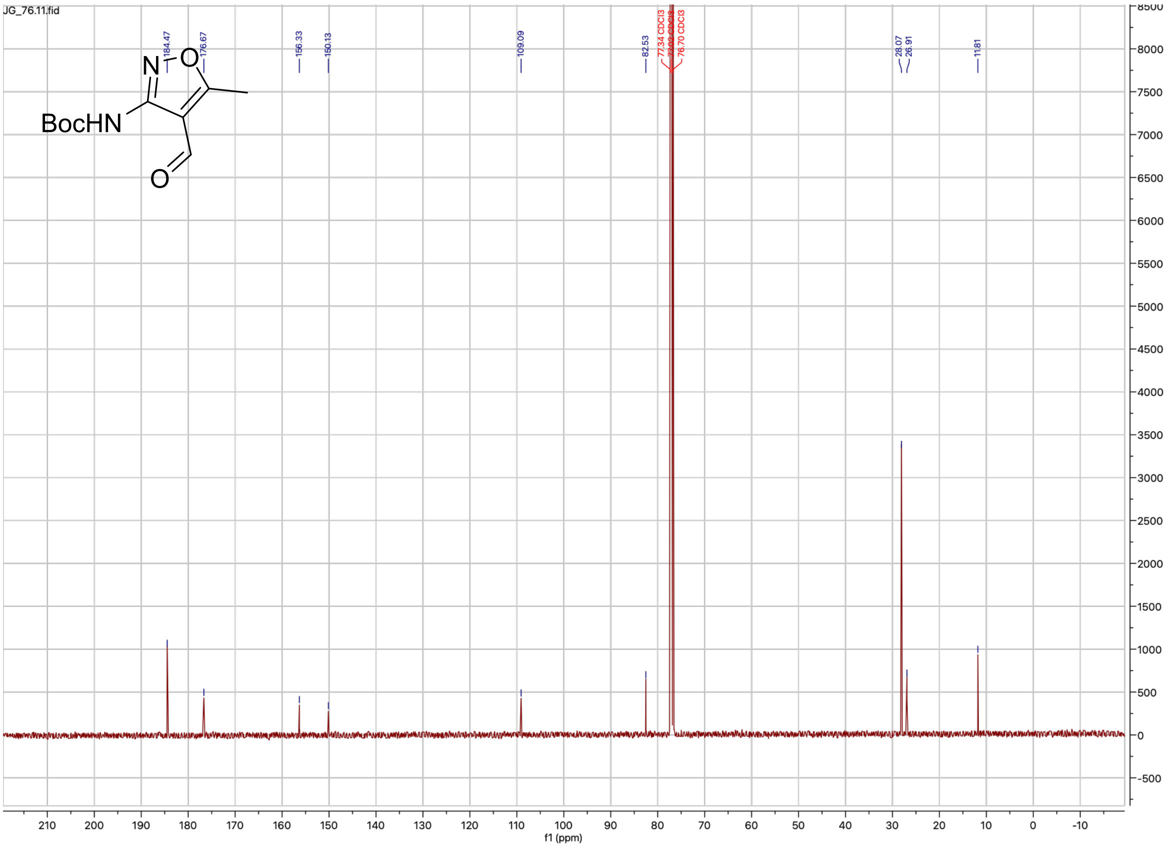
JG_76.11.fid
BocHN
N
O
184.47
176.67
156.33
150.13
109.09
82.53
77.34 CDCl3
76.70 CDCl3
28.07
26.91
11.81
f1 (ppm)

**S68**

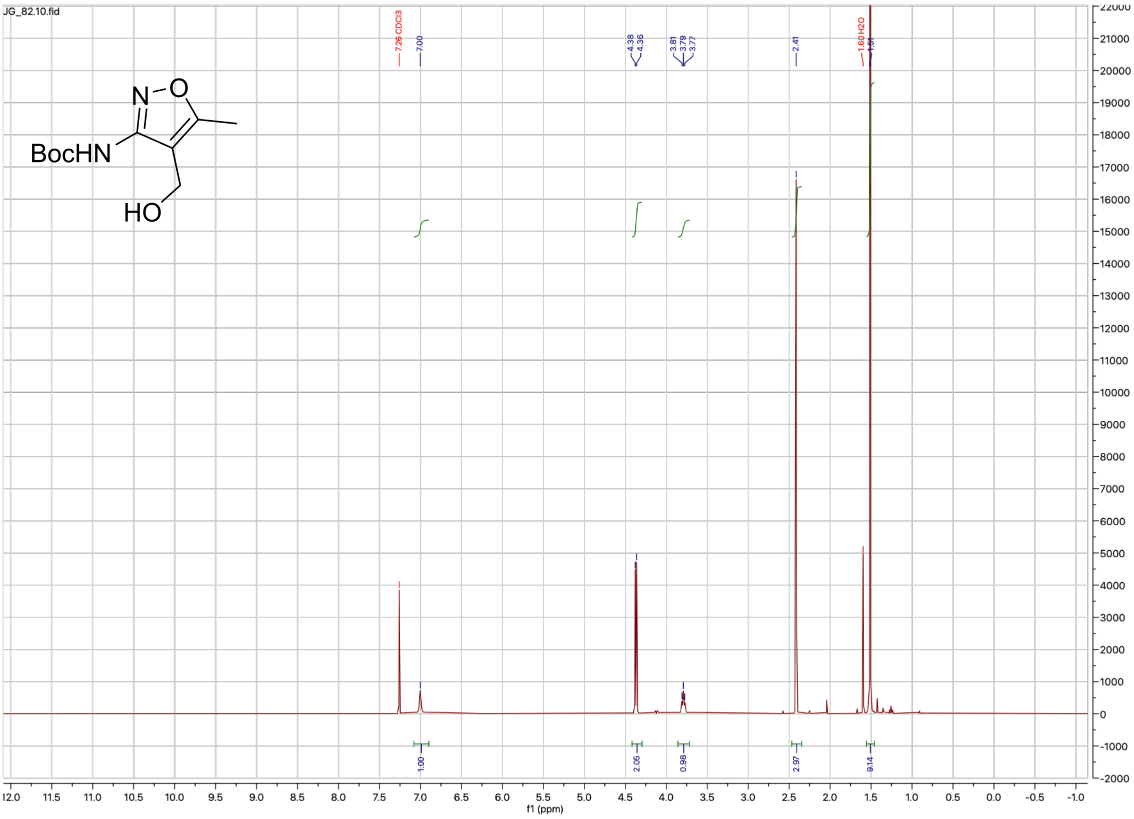
JG_82.10.fid
BocHN
HO
N
O
7.26 CDCl3
7.00
4.38
4.36
3.81
3.79
3.77
2.41
1.60 H2O
1.51
1.00
2.05
0.98
2.97
9.14
f1 (ppm)

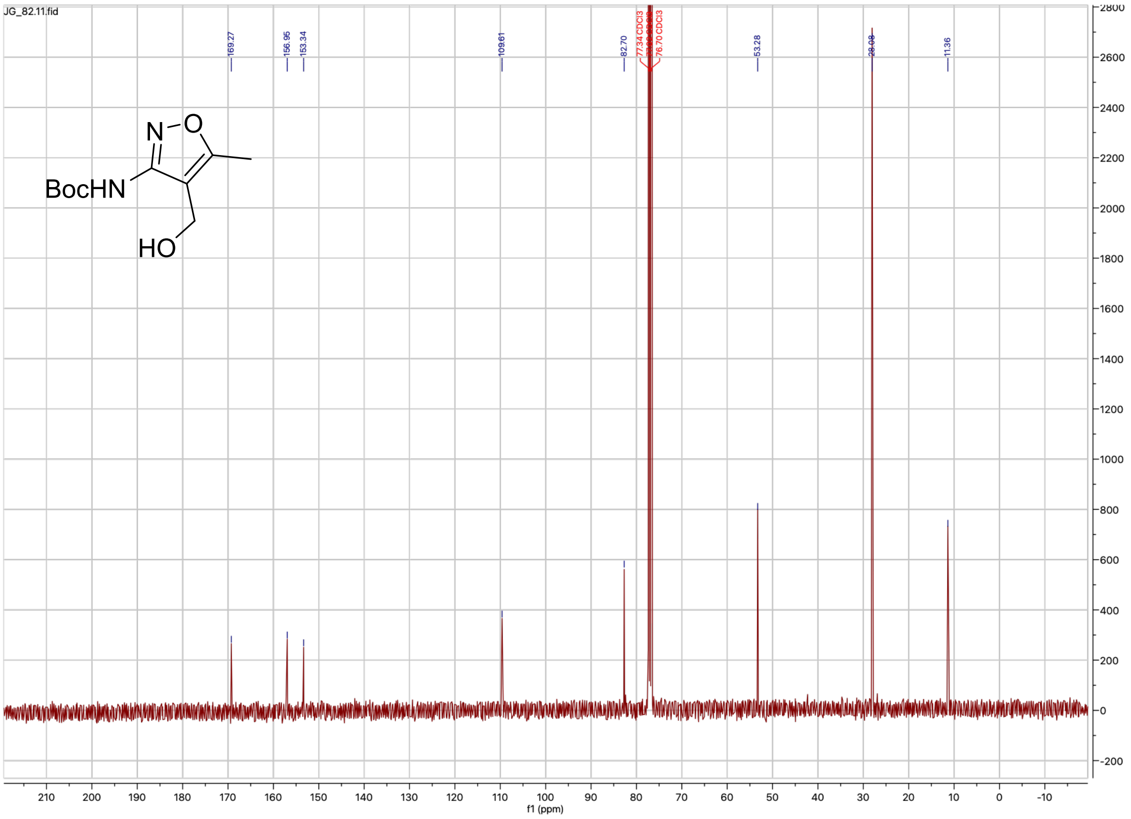
JG_82.11.fid
BocHN
HO
N
O
169.27
156.95
153.34
109.61
82.70
77.34 CDCl3
76.70 CDCl3
53.28
28.08
11.36
f1 (ppm)

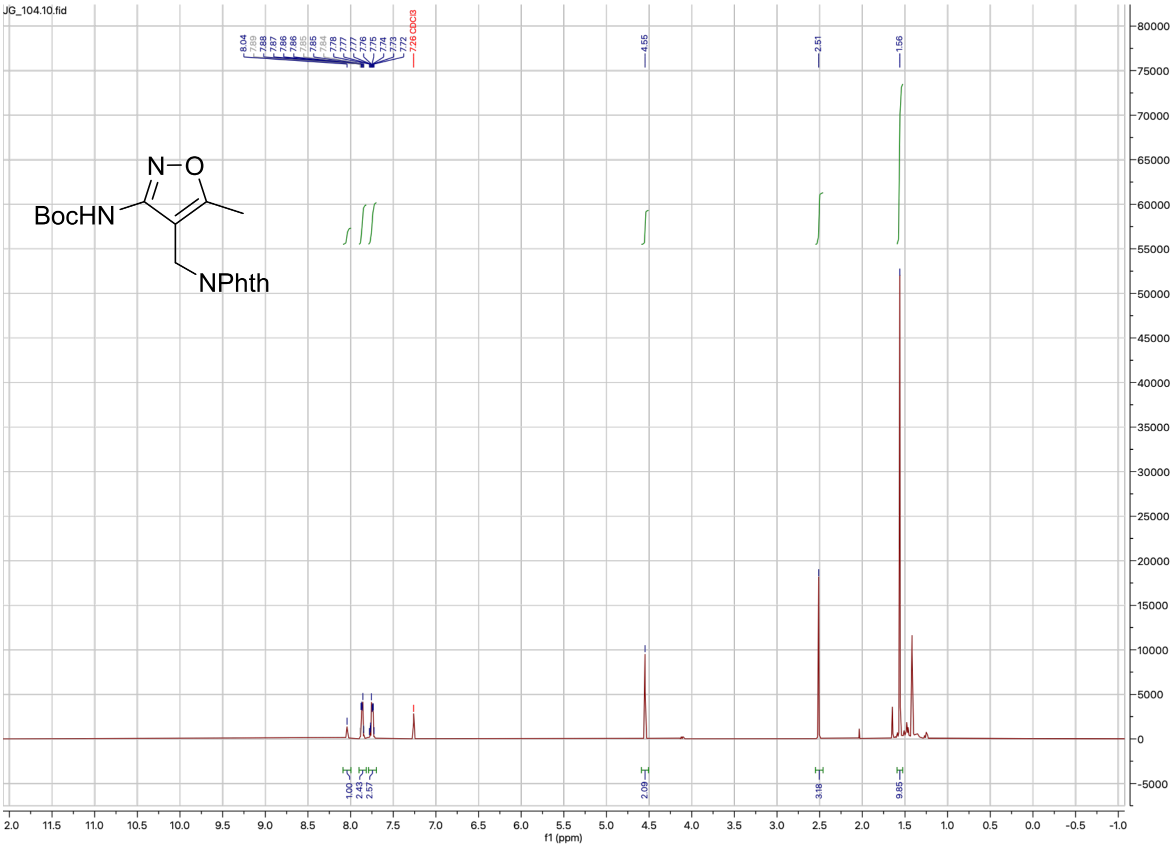
JG_104.10.fid
BocHN
N–O
NPhth
f1 (ppm)

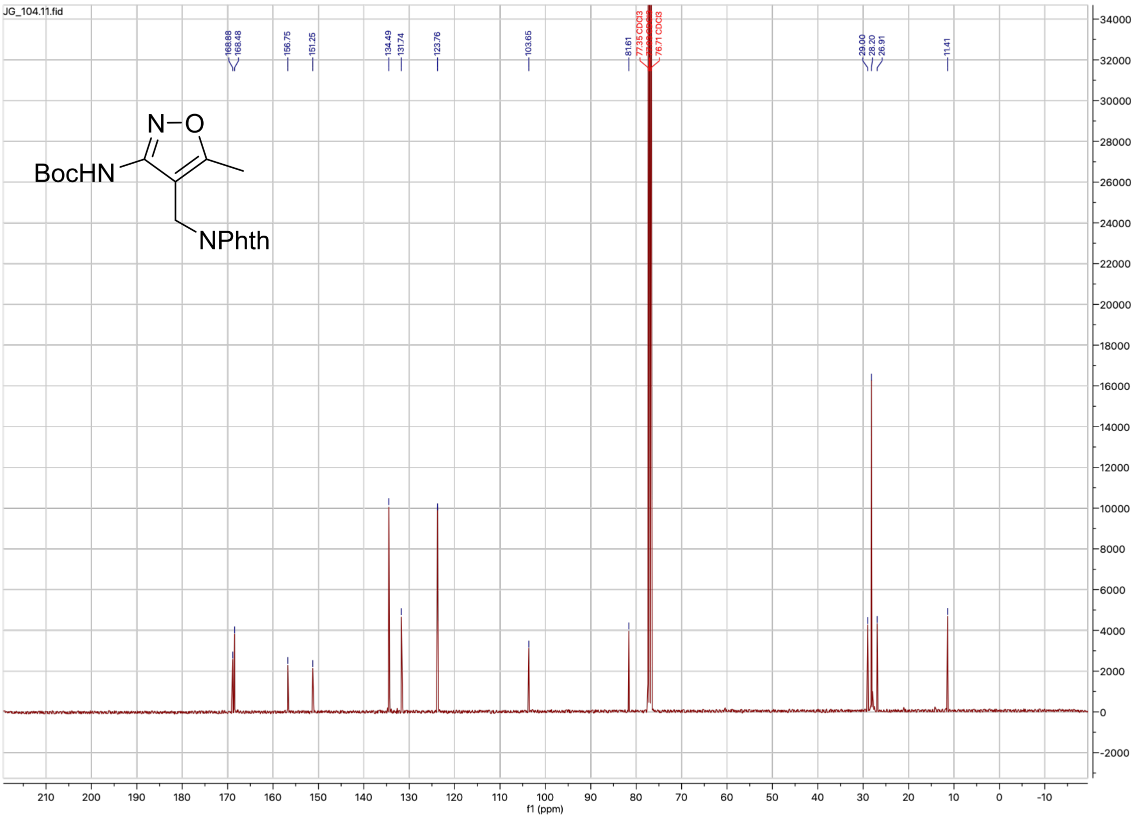
JG_104.11.fid
BocHN
N–O
NPhth
f1 (ppm)

### Compound 3

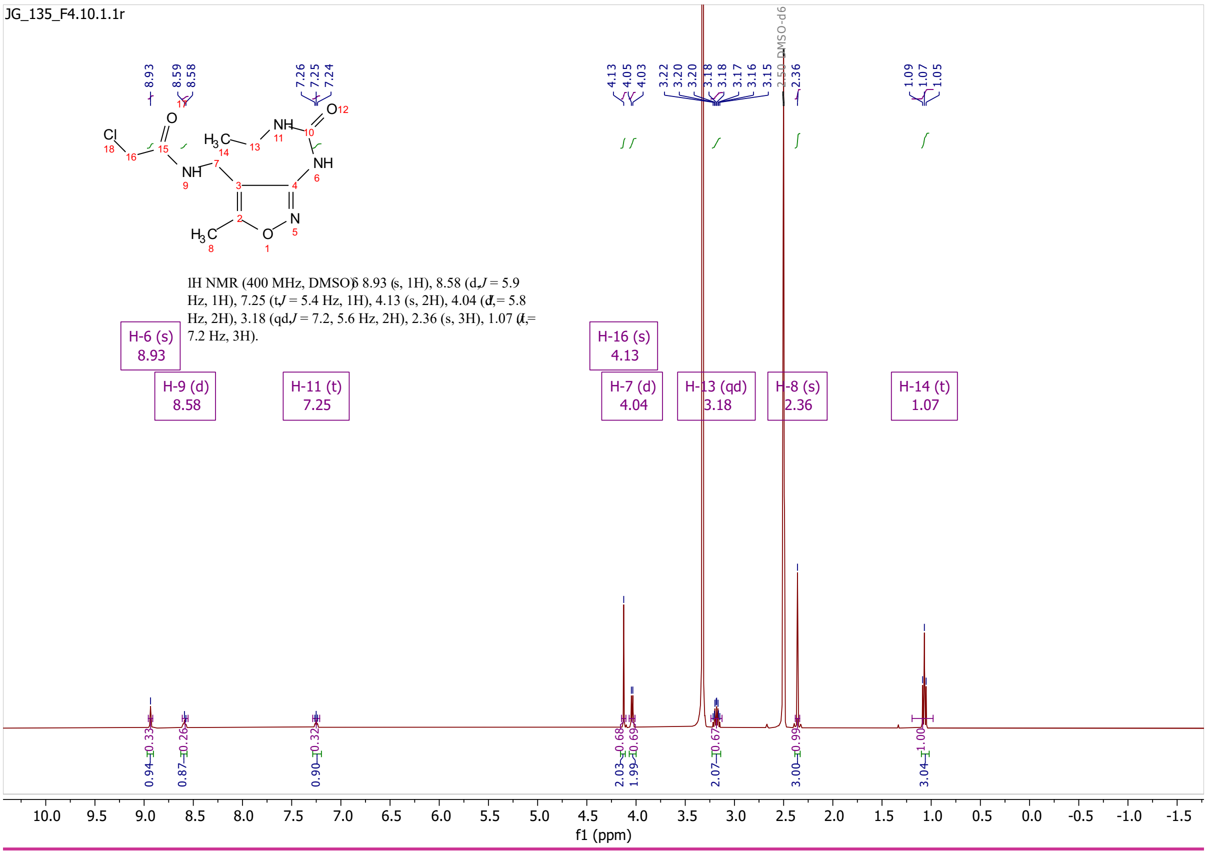
JG_135_F4.10.1.1r
1H NMR (400 MHz, DMSO) δ 8.93 (s, 1H), 8.58 (d, J = 5.9 Hz, 1H), 7.25 (t, J = 5.4 Hz, 1H), 4.13 (s, 2H), 4.04 (d, J = 5.8 Hz, 2H), 3.18 (qd, J = 7.2, 5.6 Hz, 2H), 2.36 (s, 3H), 1.07 (t, J = 7.2 Hz, 3H).
H-6 (s) 8.93
H-9 (d) 8.58
H-11 (t) 7.25
H-16 (s) 4.13
H-7 (d) 4.04
H-13 (qd) 3.18
H-8 (s) 2.36
H-14 (t) 1.07
2.50 DMSO-d6
f1 (ppm)

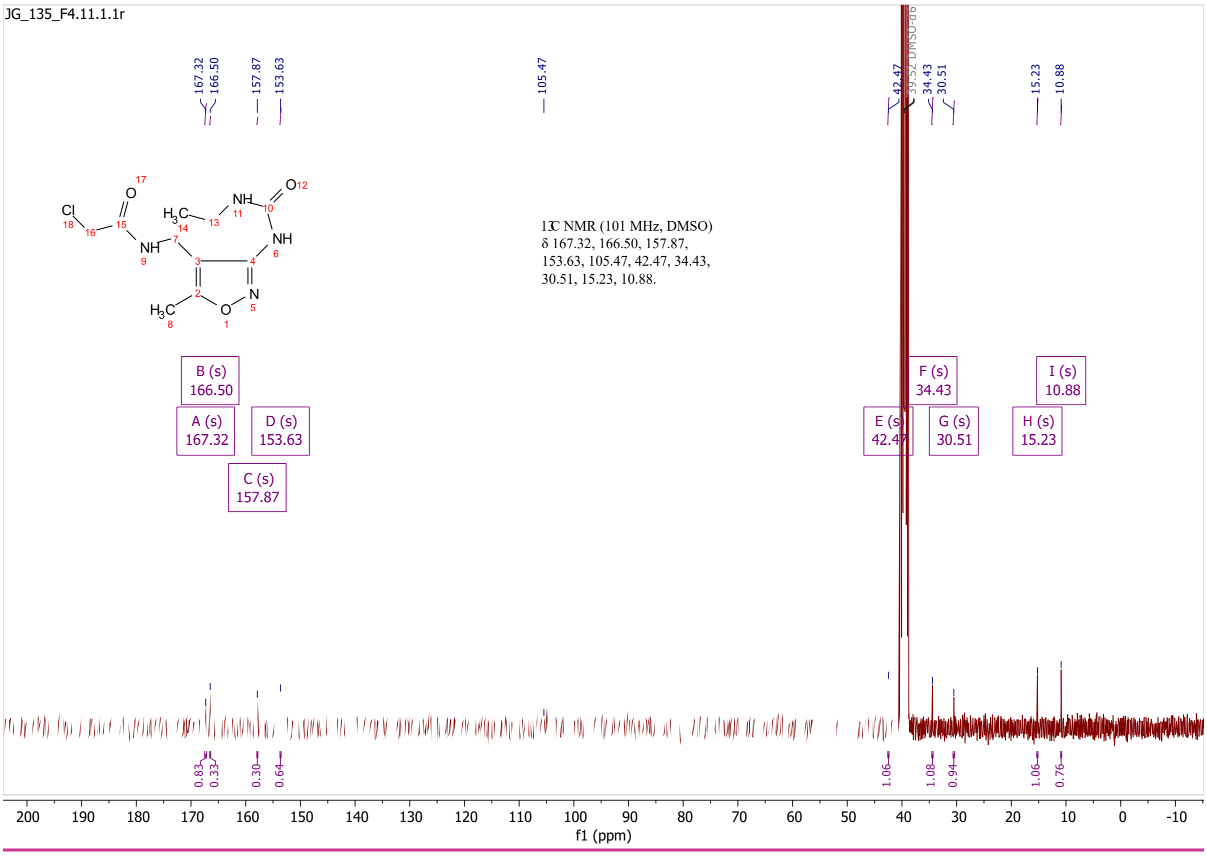
JG_135_F4.11.1.1r
13C NMR (101 MHz, DMSO) δ 167.32, 166.50, 157.87, 153.63, 105.47, 42.47, 34.43, 30.51, 15.23, 10.88.
A (s) 167.32
B (s) 166.50
C (s) 157.87
D (s) 153.63
E (s) 42.47
F (s) 34.43
G (s) 30.51
H (s) 15.23
I (s) 10.88
39.52 DMSO-d6
f1 (ppm)

# S74

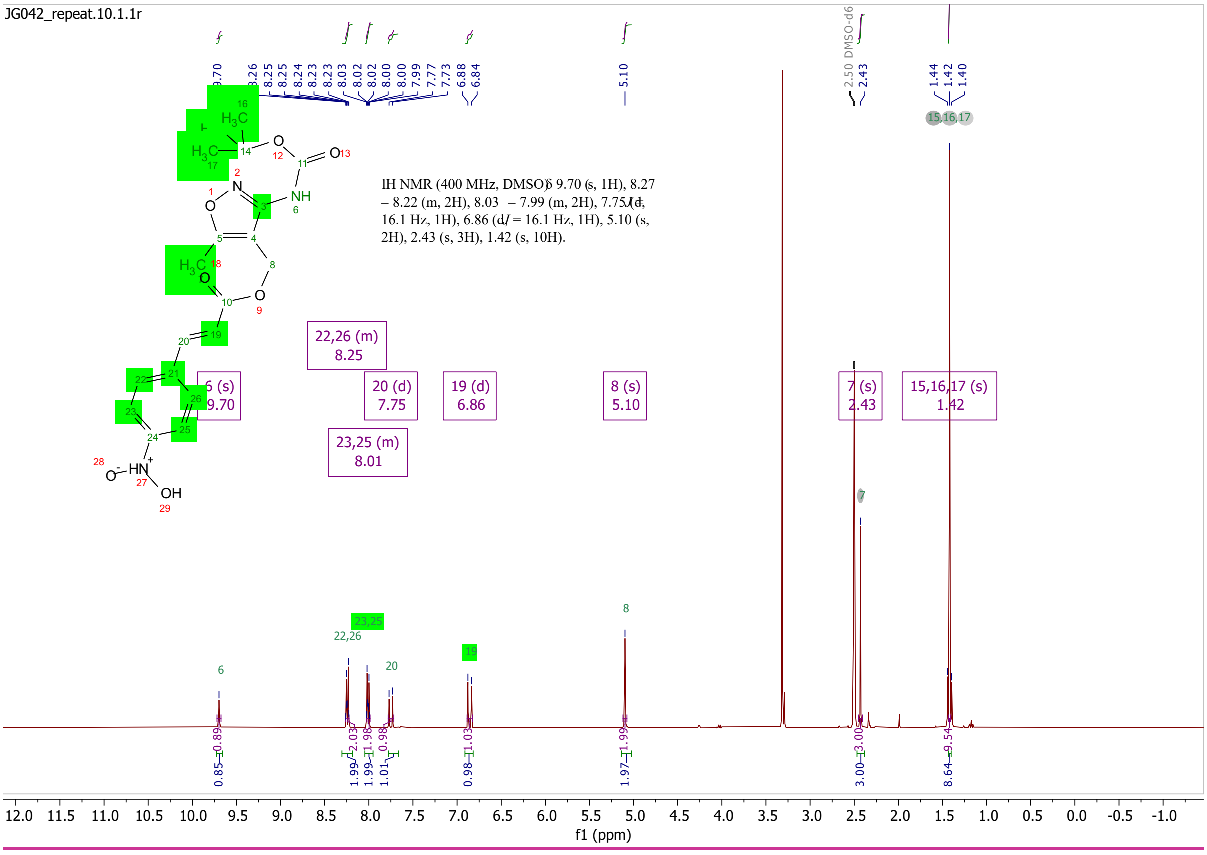
JG042_repeat.10.1.1r
1H NMR (400 MHz, DMSO) δ 9.70 (s, 1H), 8.27 – 8.22 (m, 2H), 8.03 – 7.99 (m, 2H), 7.75 (d, J = 16.1 Hz, 1H), 6.86 (d, J = 16.1 Hz, 1H), 5.10 (s, 2H), 2.43 (s, 3H), 1.42 (s, 10H).
22,26 (m) 8.25
6 (s) 9.70
20 (d) 7.75
19 (d) 6.86
8 (s) 5.10
7 (s) 2.43
15,16,17 (s) 1.42
23,25 (m) 8.01
2.50 DMSO-d6
f1 (ppm)

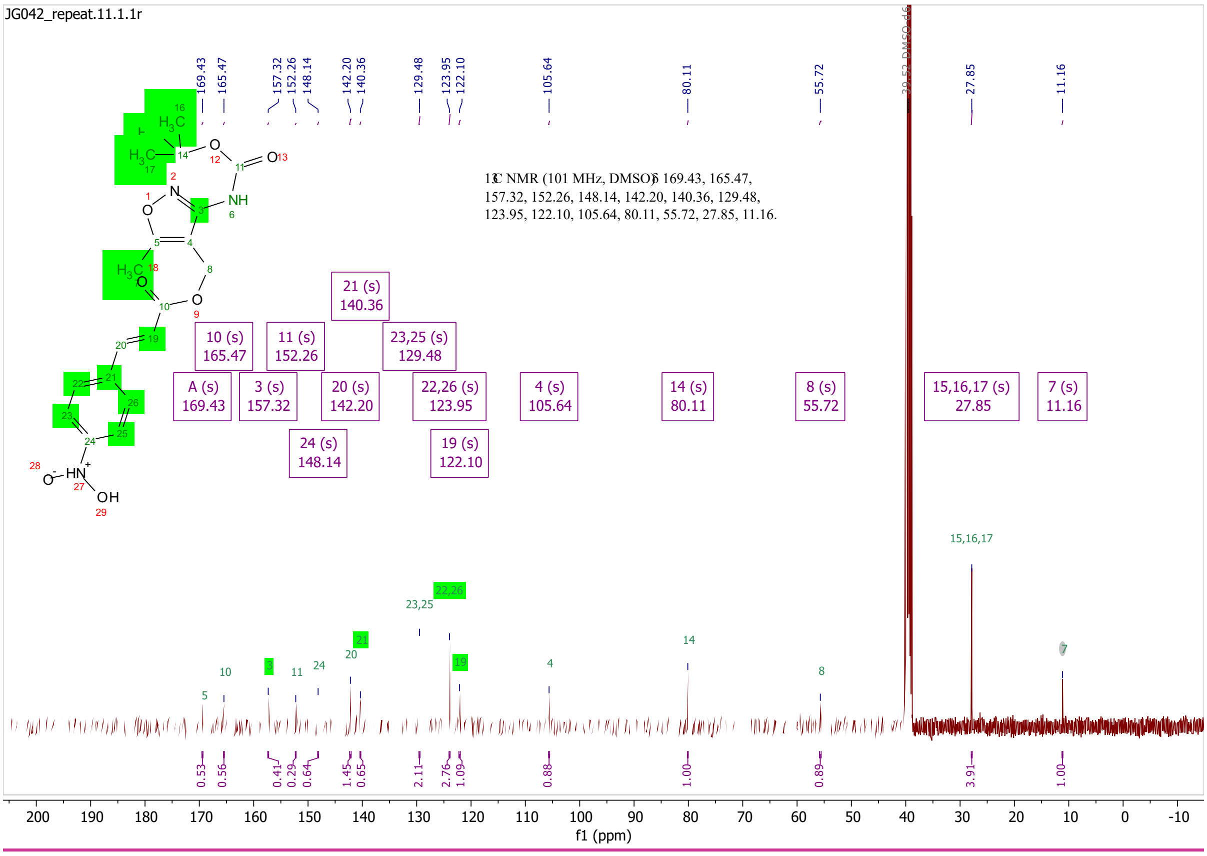
JG042_repeat.11.1.1r
13C NMR (101 MHz, DMSO) δ 169.43, 165.47, 157.32, 152.26, 148.14, 142.20, 140.36, 129.48, 123.95, 122.10, 105.64, 80.11, 55.72, 27.85, 11.16.
21 (s) 140.36
10 (s) 165.47
11 (s) 152.26
23,25 (s) 129.48
A (s) 169.43
3 (s) 157.32
20 (s) 142.20
22,26 (s) 123.95
4 (s) 105.64
14 (s) 80.11
8 (s) 55.72
15,16,17 (s) 27.85
7 (s) 11.16
24 (s) 148.14
19 (s) 122.10
f1 (ppm)

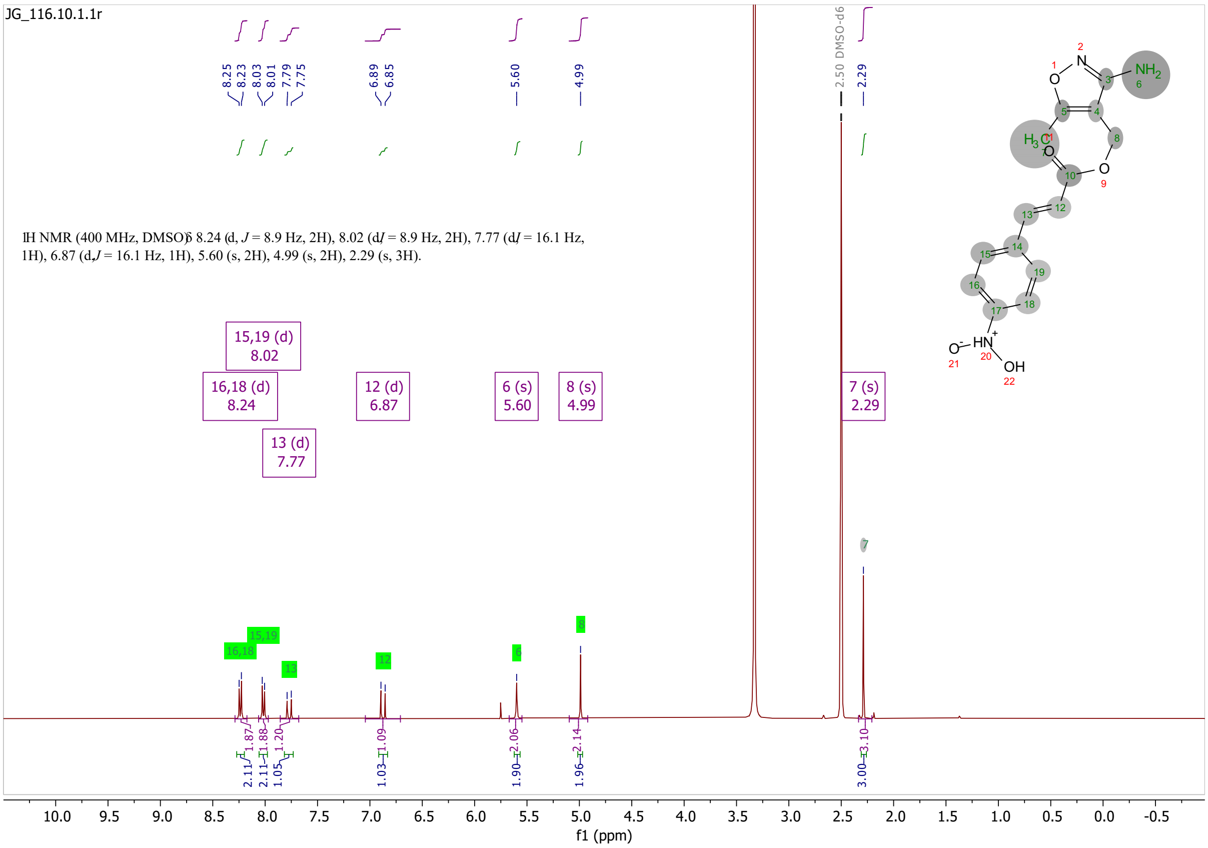
JG_116.10.1.1r
1H NMR (400 MHz, DMSO) δ 8.24 (d, J = 8.9 Hz, 2H), 8.02 (d, J = 8.9 Hz, 2H), 7.77 (d, J = 16.1 Hz, 1H), 6.87 (d, J = 16.1 Hz, 1H), 5.60 (s, 2H), 4.99 (s, 2H), 2.29 (s, 3H).
15,19 (d) 8.02
16,18 (d) 8.24
13 (d) 7.77
12 (d) 6.87
6 (s) 5.60
8 (s) 4.99
7 (s) 2.29
2.50 DMSO-d6
f1 (ppm)

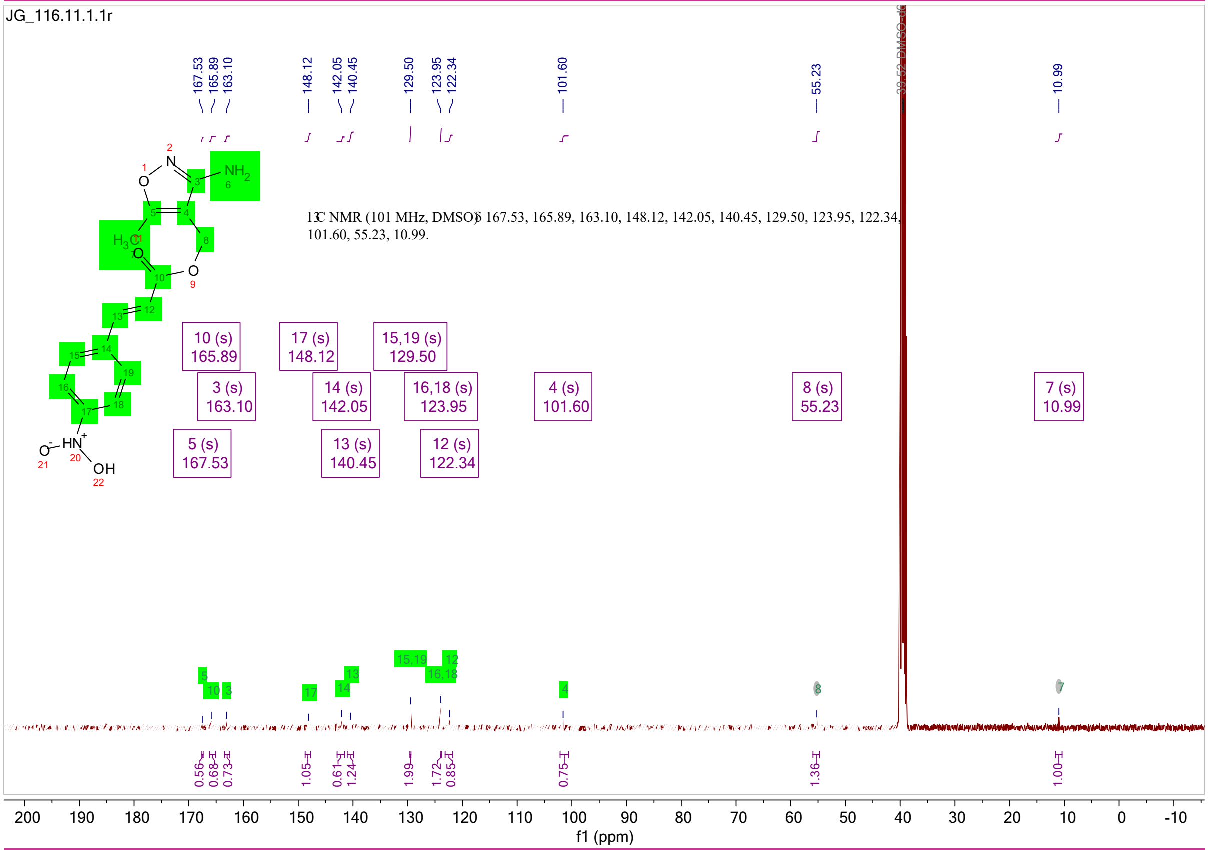
JG_116.11.1.1r
13C NMR (101 MHz, DMSO) δ 167.53, 165.89, 163.10, 148.12, 142.05, 140.45, 129.50, 123.95, 122.34, 101.60, 55.23, 10.99.
10 (s) 165.89
3 (s) 163.10
5 (s) 167.53
17 (s) 148.12
14 (s) 142.05
13 (s) 140.45
15,19 (s) 129.50
16,18 (s) 123.95
12 (s) 122.34
4 (s) 101.60
8 (s) 55.23
7 (s) 10.99
f1 (ppm)

# 4

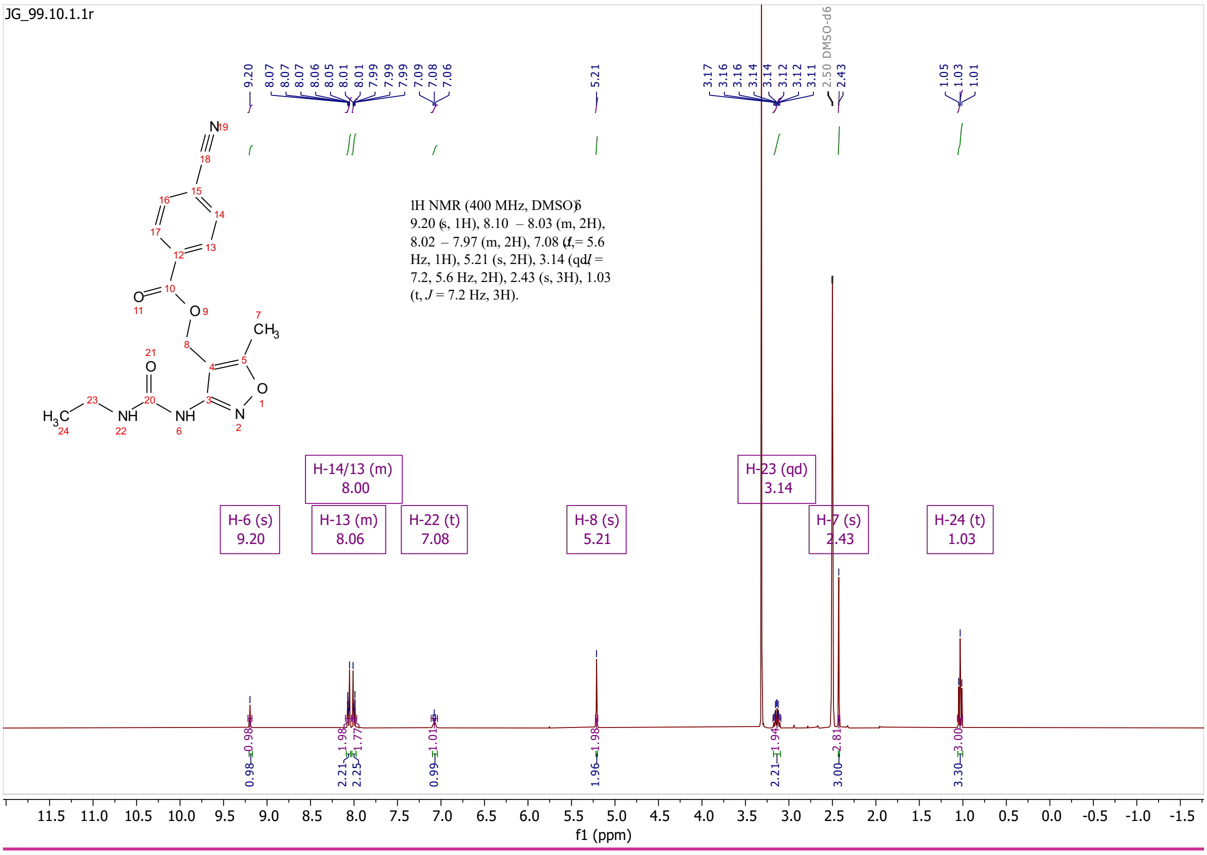
JG_99.10.1.1r
1H NMR (400 MHz, DMSO) δ 9.20 (s, 1H), 8.10 – 8.03 (m, 2H), 8.02 – 7.97 (m, 2H), 7.08 (t, J = 5.6 Hz, 1H), 5.21 (s, 2H), 3.14 (qd, J = 7.2, 5.6 Hz, 2H), 2.43 (s, 3H), 1.03 (t, J = 7.2 Hz, 3H).
H-6 (s) 9.20
H-14/13 (m) 8.00
H-13 (m) 8.06
H-22 (t) 7.08
H-8 (s) 5.21
H-23 (qd) 3.14
H-7 (s) 2.43
H-24 (t) 1.03
2.50 DMSO-d6
f1 (ppm)

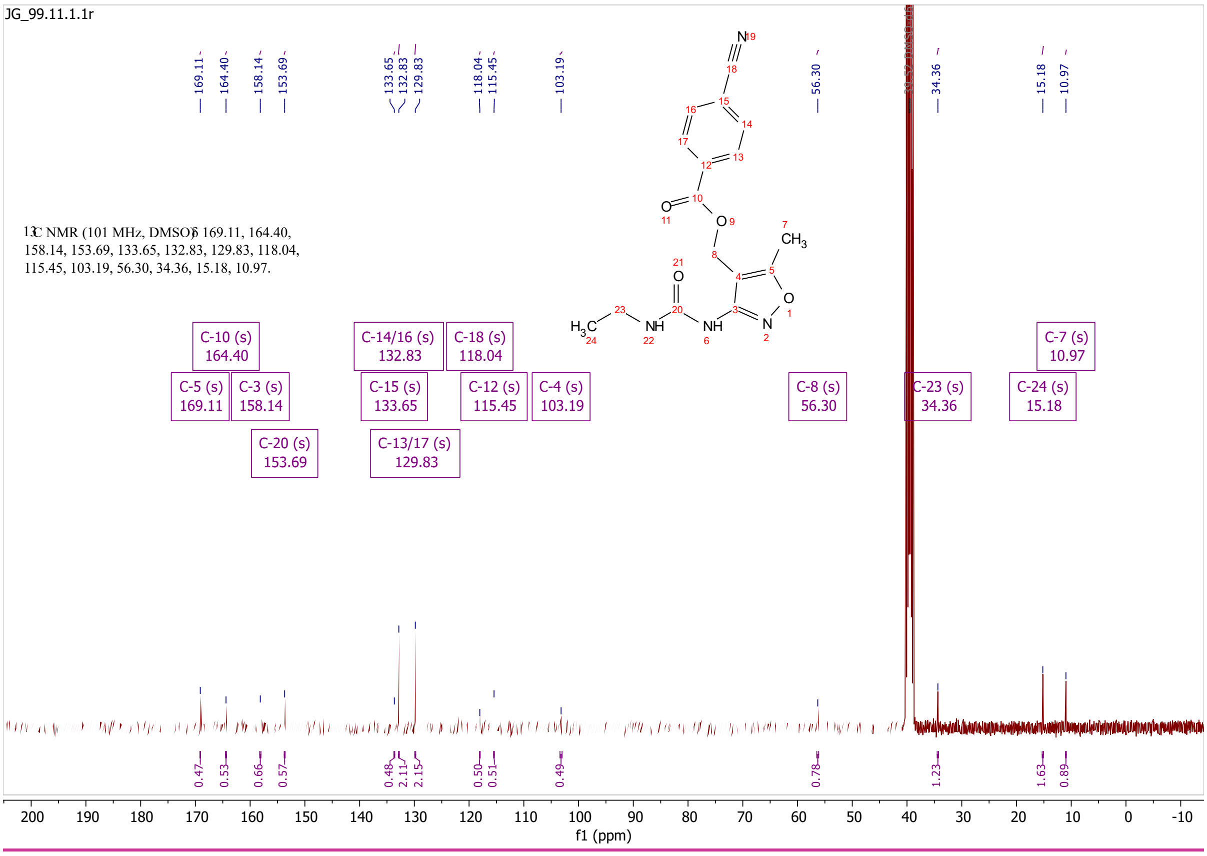
JG_99.11.1.1r
13C NMR (101 MHz, DMSO) δ 169.11, 164.40, 158.14, 153.69, 133.65, 132.83, 129.83, 118.04, 115.45, 103.19, 56.30, 34.36, 15.18, 10.97.
C-10 (s) 164.40
C-5 (s) 169.11
C-3 (s) 158.14
C-20 (s) 153.69
C-14/16 (s) 132.83
C-15 (s) 133.65
C-13/17 (s) 129.83
C-18 (s) 118.04
C-12 (s) 115.45
C-4 (s) 103.19
C-8 (s) 56.30
C-23 (s) 34.36
C-24 (s) 15.18
C-7 (s) 10.97
f1 (ppm)

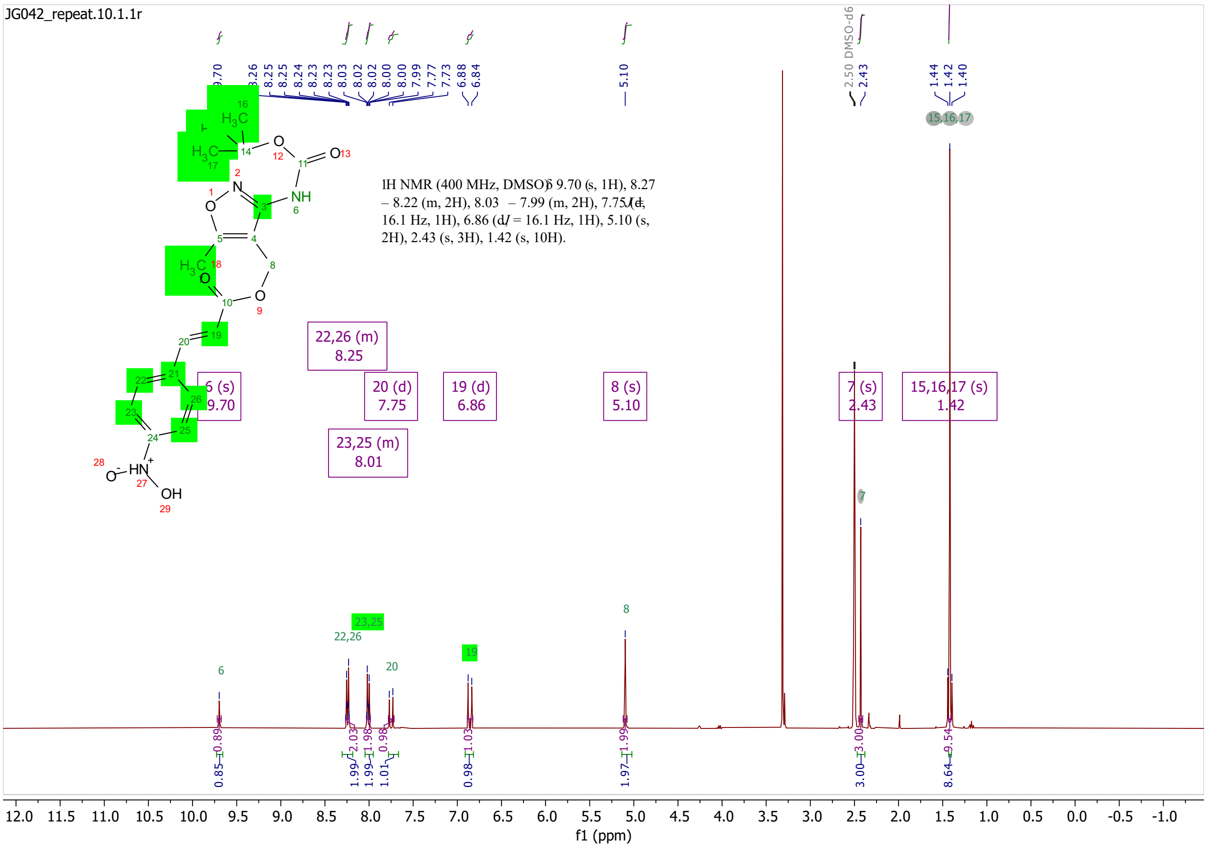
JG042_repeat.10.1.1r
1H NMR (400 MHz, DMSO) δ 9.70 (s, 1H), 8.27 – 8.22 (m, 2H), 8.03 – 7.99 (m, 2H), 7.75 (d, J = 16.1 Hz, 1H), 6.86 (d, J = 16.1 Hz, 1H), 5.10 (s, 2H), 2.43 (s, 3H), 1.42 (s, 10H).
f1 (ppm)

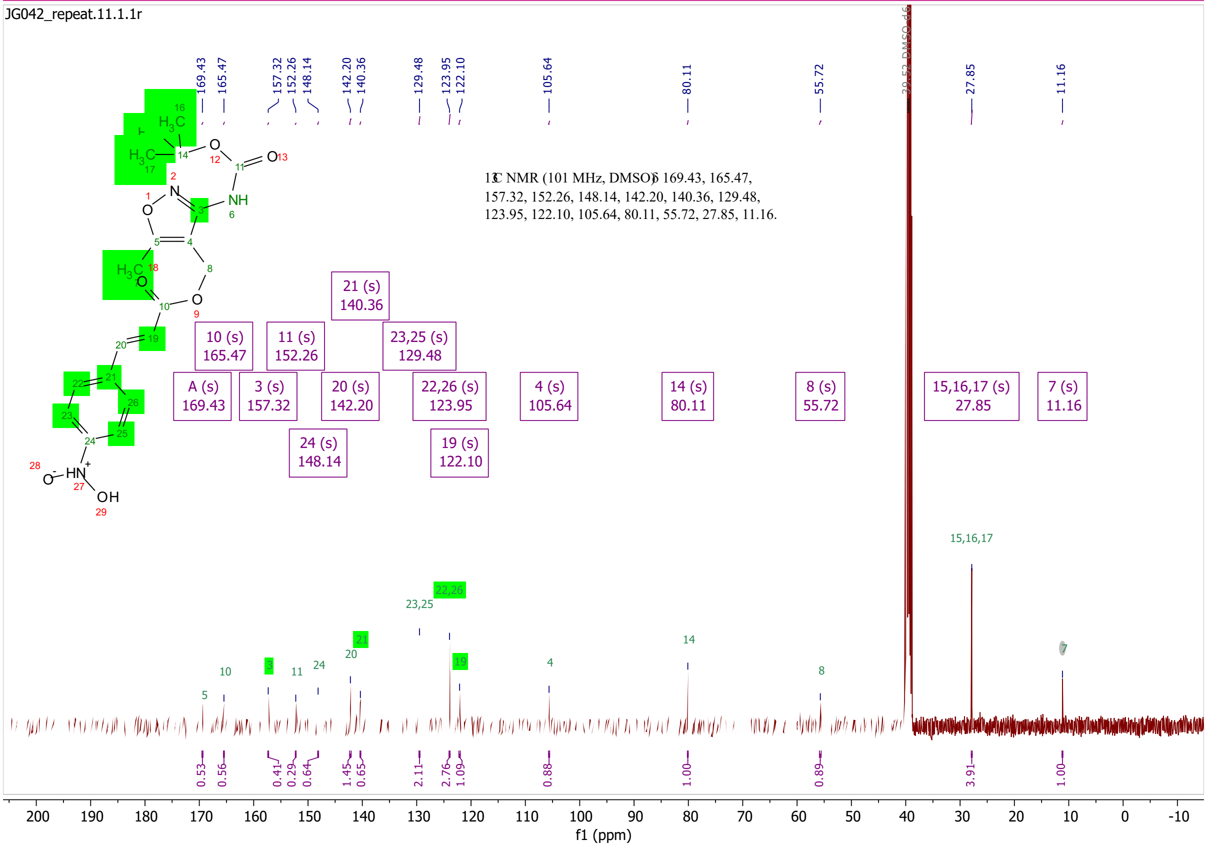
JG042_repeat.11.1.1r
13C NMR (101 MHz, DMSO) δ 169.43, 165.47, 157.32, 152.26, 148.14, 142.20, 140.36, 129.48, 123.95, 122.10, 105.64, 80.11, 55.72, 27.85, 11.16.
f1 (ppm)

# S77

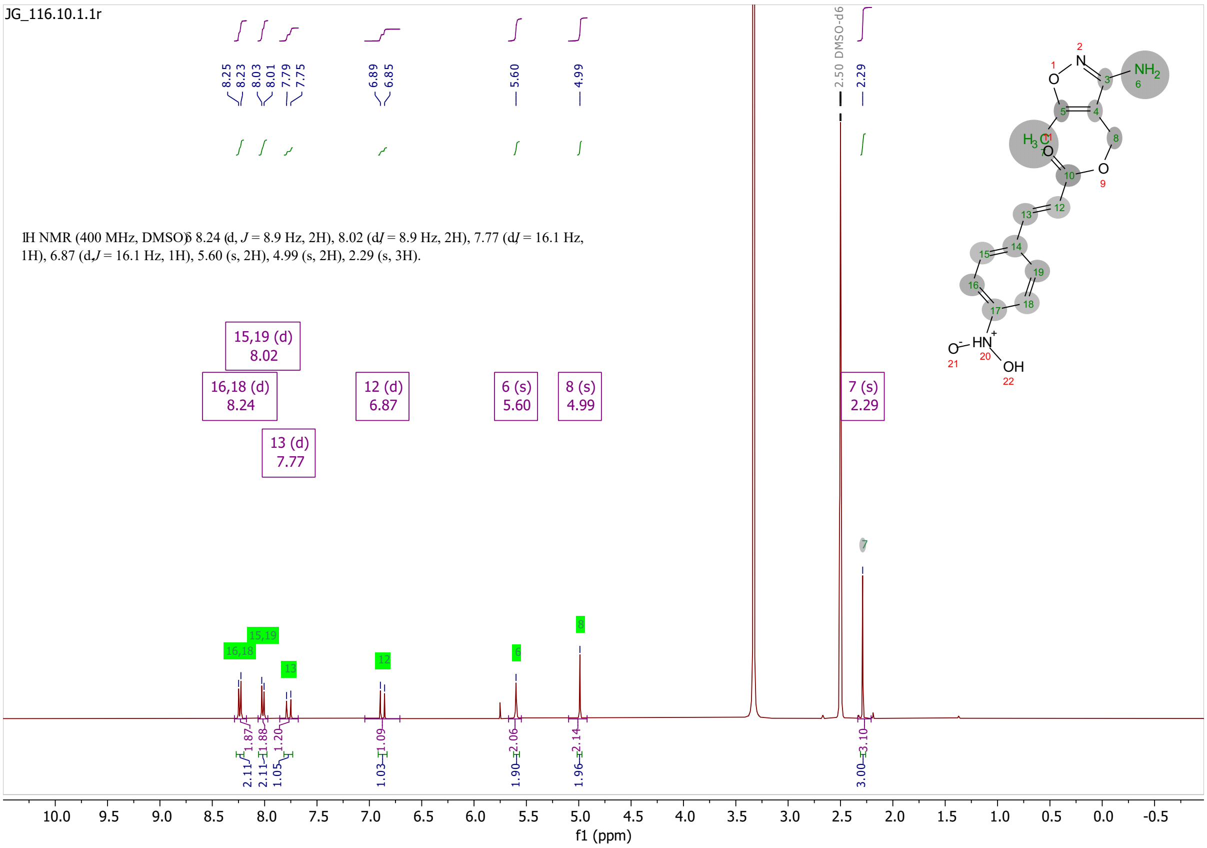
JG_116.10.1.1r
1H NMR (400 MHz, DMSO) δ 8.24 (d, J = 8.9 Hz, 2H), 8.02 (d, J = 8.9 Hz, 2H), 7.77 (d, J = 16.1 Hz, 1H), 6.87 (d, J = 16.1 Hz, 1H), 5.60 (s, 2H), 4.99 (s, 2H), 2.29 (s, 3H).
15,19 (d) 8.02
16,18 (d) 8.24
13 (d) 7.77
12 (d) 6.87
6 (s) 5.60
8 (s) 4.99
7 (s) 2.29
2.50 DMSO-d6
f1 (ppm)

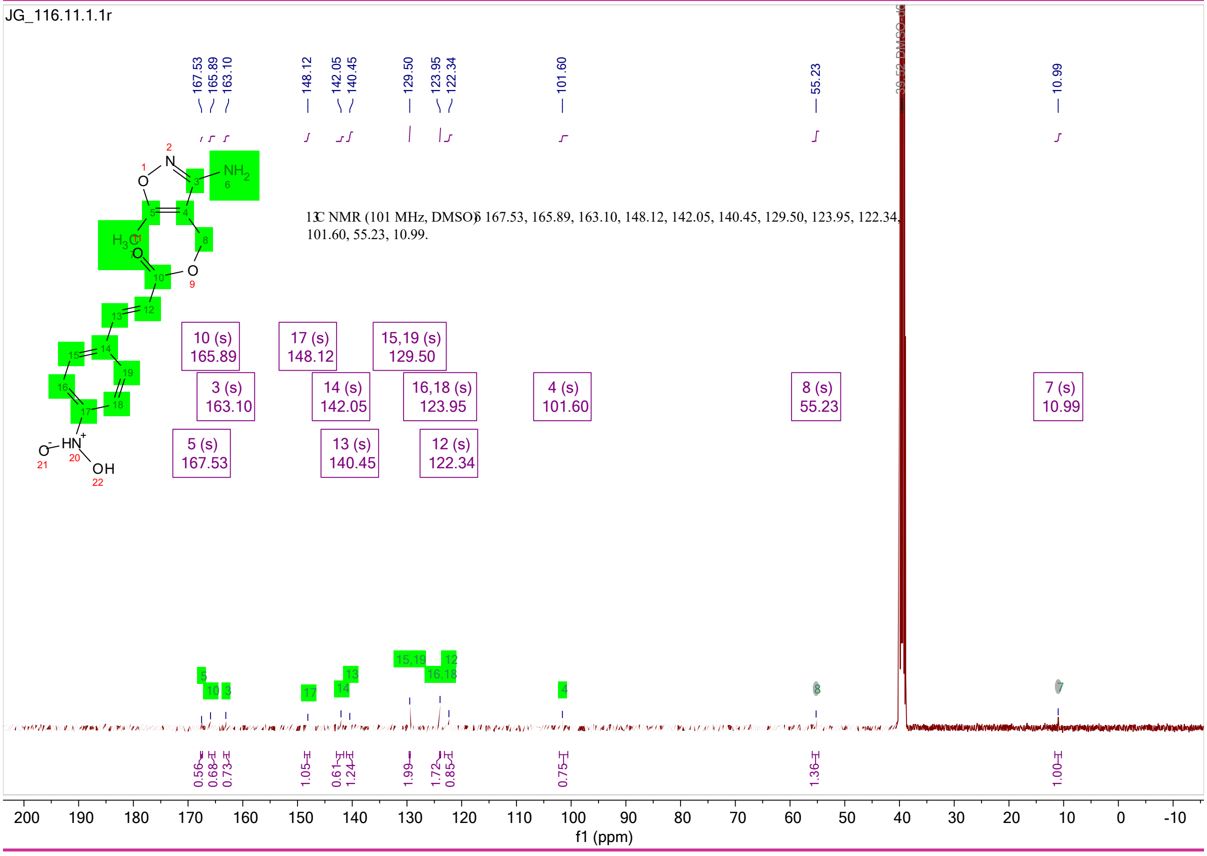
JG_116.11.1.1r
13C NMR (101 MHz, DMSO) δ 167.53, 165.89, 163.10, 148.12, 142.05, 140.45, 129.50, 123.95, 122.34, 101.60, 55.23, 10.99.
10 (s) 165.89
17 (s) 148.12
15,19 (s) 129.50
3 (s) 163.10
14 (s) 142.05
16,18 (s) 123.95
4 (s) 101.60
8 (s) 55.23
7 (s) 10.99
5 (s) 167.53
13 (s) 140.45
12 (s) 122.34
f1 (ppm)

# 5

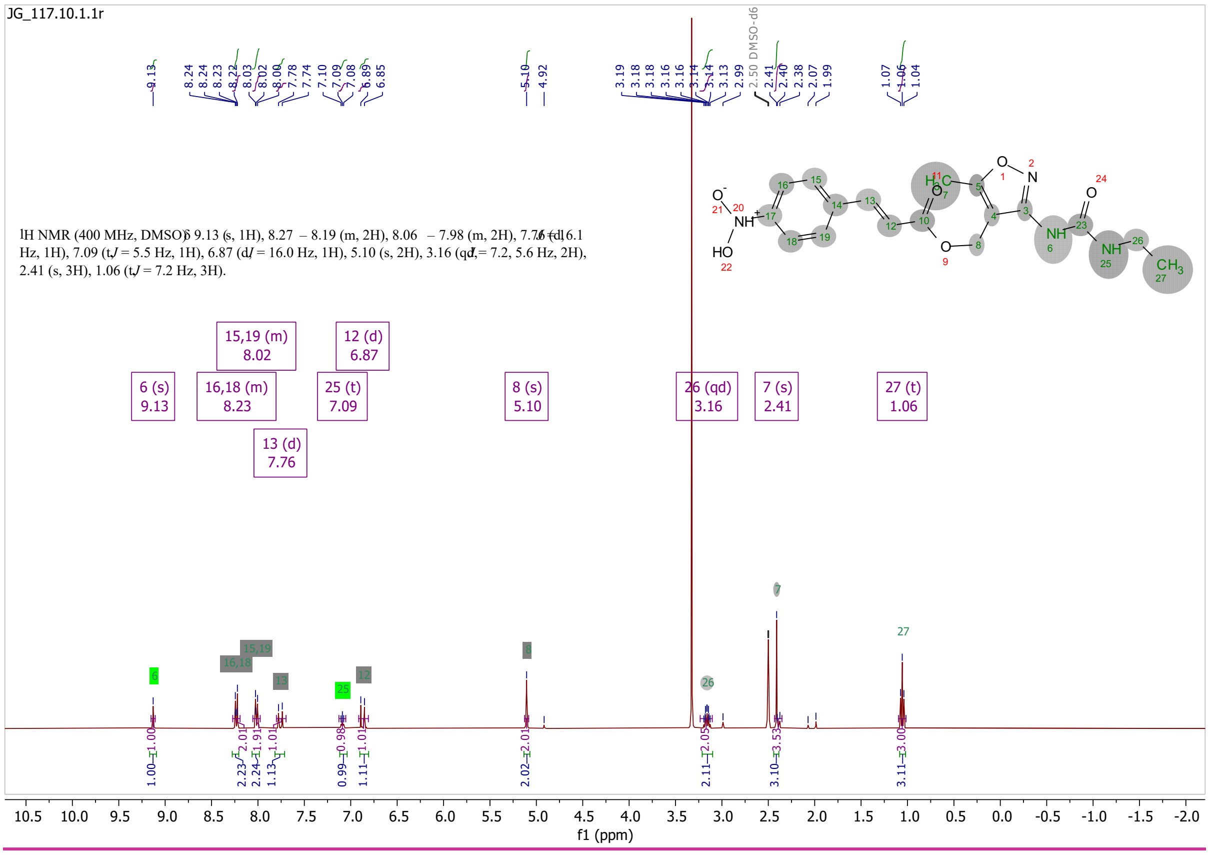
JG_117.10.1.1r
1H NMR (400 MHz, DMSO) δ 9.13 (s, 1H), 8.27 – 8.19 (m, 2H), 8.06 – 7.98 (m, 2H), 7.76 (d, J = 16.1 Hz, 1H), 7.09 (t, J = 5.5 Hz, 1H), 6.87 (d, J = 16.0 Hz, 1H), 5.10 (s, 2H), 3.16 (qd, J = 7.2, 5.6 Hz, 2H), 2.41 (s, 3H), 1.06 (t, J = 7.2 Hz, 3H).
f1 (ppm)

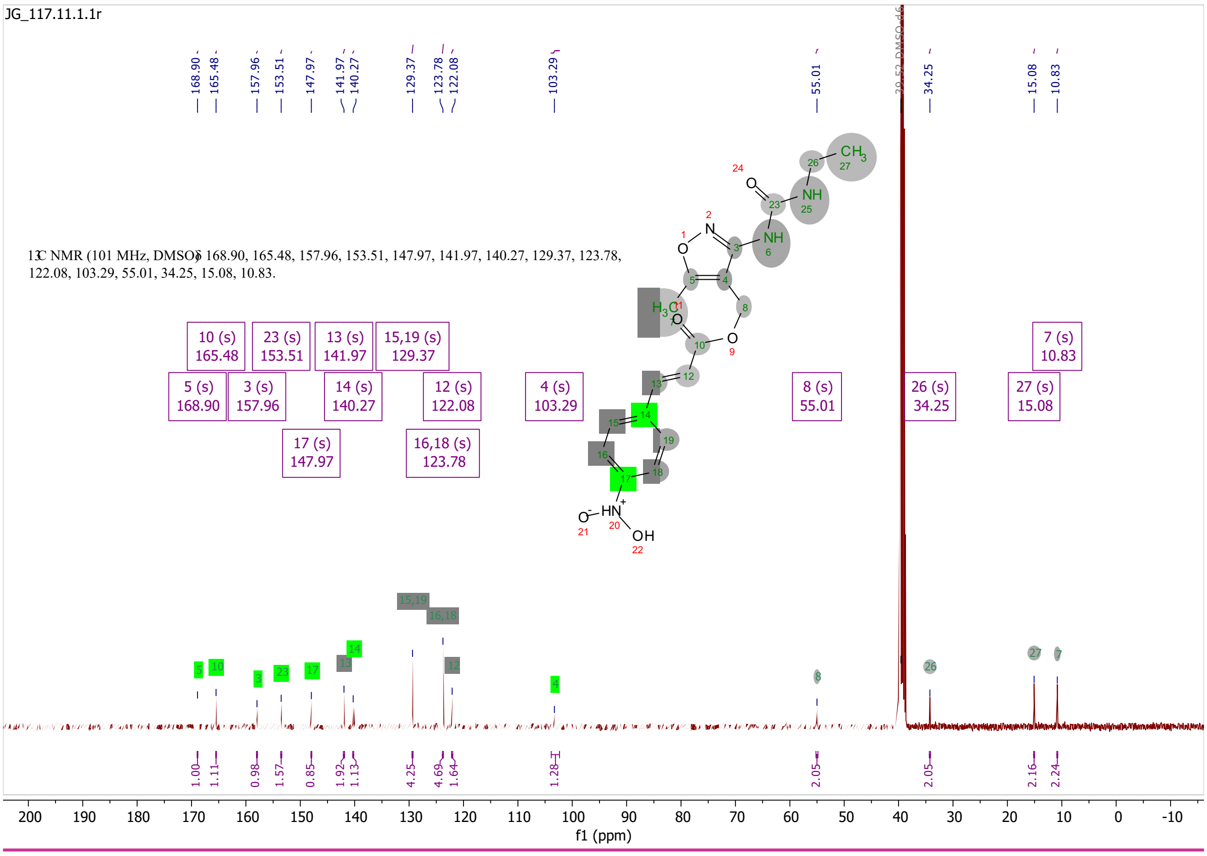
JG_117.11.1.1r
13C NMR (101 MHz, DMSO) δ 168.90, 165.48, 157.96, 153.51, 147.97, 141.97, 140.27, 129.37, 123.78, 122.08, 103.29, 55.01, 34.25, 15.08, 10.83.
f1 (ppm)
